# Rare Genetic Variants Correlate with Better Processing Speed

**DOI:** 10.1101/2022.01.12.476030

**Authors:** Zeyuan Song, Anastasia Gurinovich, Marianne Nygaard, Jonas Mengel-From, Stacy Andersen, Stephanie Cosentino, Nicole Schupfs, Joseph Lee, Joseph Zmuda, Svetlana Ukraintseva, Konstantin Arbeev, Kaare Christensen, Thomas Perls, Paola Sebastiani

## Abstract

We conducted a genome-wide association study (GWAS) of Digit Symbol Substitution Test (DSST) scores administered in 4207 family members of the Long Life Family Study (LLFS). Genotype data were imputed to the HRC panel of 64,940 haplotypes resulting in ~15M genetic variants with quality score > 0.7. The results were replicated using genetic data imputed to the 1000 Genomes phase 3 reference panel from two Danish twin cohorts: the study of Middle Aged Danish Twins and the Longitudinal Study of Aging Danish Twins. The GWAS in LLFS discovered 20 rare genetic variants (minor allele frequency (MAF) < 1.0%) that reached genome-wide significance (p-value < 5×10^−8^). Among these, 18 variants had large protective effects on the processing speed, including rs7623455, rs9821776, rs9821587, rs78704059 on chromosome 3, which were replicated in the combined Danish twin cohort. These SNPs are located in/near two genes, *THRB* and *RARB*, that belonged to thyroid hormone receptors family that may influence speed of metabolism and cognitive aging. The gene-level tests in LLFS confirmed that these two genes are associated with processing speed.

## Introduction

Human cognitive functions change with age, both normally and pathologically. Some cognitive abilities, such as vocabulary, are resilient to brain aging, while other abilities such as conceptual reasoning, memory, and processing speed, decline with age [Harada 2013]. Age-related cognitive impairment is an increasing problem for society. As the global population aged over 65 years continue to increase, and the number of older persons is projected to double to 1.5 billion in 2050 [United Nations 2019], it is expected that the clinical and economic burden of cognitive impairment on society will increase. Understanding cognitive aging and its possible therapeutic targets should thus be an important research focus to help delay or prevent age-related cognitive impairment.

Cognitive functions are usually assessed by administering a battery of neuropsychological tests. For example, verbal episodic memory can be assessed by logical memory tests for immediate and delayed recall. Working memory can be assessed by administering the digit span forward and backward tests, and processing speed and executive control can be assessed by administering the Digit Symbol Substitution Test (DSST). Several of these tests are under strong genetic control with heritability ranging from 16% to 68% [Cirulli 2010], and over the past years several genomewide association studies (GWAS) have identified common genetic variants associated with cognitive functions [Trampush 2017; Davies 2015]. A recent GWAS of 300,486 individuals identified 148 independent genetic loci that associate with general cognitive function defined as the first principal component of multiple cognitive test scores [Davies 2019].

The heritability of the DSST score is strong, ranging from 0.36-0.68 [Cirulli 2010] but, despite such a high heritability, previous genome-wide association studies failed to identify and replicate common genetic variants associated with DSST score [Luciano 2011; Ibrahim-Verbaas 2015]. Previous studies focused attention on the association between common genetic variants and DSST scores in the general population. Here, we leverage the unique family-based design and the enrichment for healthy agers of the Long Life Family Study (LLFS) to identify rare variants potentially associated with preservation of DSST scores with aging. The LLFS includes very old subjects (90+ years old) who maintained good DSST scores compared to individuals without familial longevity. The study administered the DSST at enrollment and approximately 8 years after and, by studying the longitudinal changes of DSST, we showed that there are individuals whose DSST scores decline slower compared to others [Sebastiani 2020]. In addition, we showed that the heritability of DSST at baseline and its change over time was over 40% [Wojczynski 2019]. These results suggest that those extremely long lived individuals with normal cognitive function may carry protective genetic variants that make them resilient to cognitive decline as they age. We therefore conducted a genome-wide association study of DSST scores measured in subjects enrolled in the LLFS. The genetic findings were further replicated in middle-aged and elderly twins from the Danish Twin Registry.

## Method

### Study populations

#### Long Life Family Study (LLFS)

The LLFS is a multicenter, multinational longitudinal two generation family study of longevity and heathy aging [Wojczynski 2019]. A total of 4953 subjects from 539 families were enrolled between 2006 and 2009, and carefully phenotyped in person. Participants were enrolled at three American field centers (Boston, Pittsburgh and New York), and a European field center in Denmark. Potential probands were recruited based on older age, capacity to understand the study and their Family Longevity Selection Score (FLoSS). The FLoSS quantifies the degree of familial longevity using sex and birth-year cohort survival probabilities of the proband and their siblings [Sebastiani 2009]. Eligibility of sibships for the study was based on a FLoSS score >7 and having at least one living sibling and at least one offspring willing to be enrolled in the study. Socio-demographic, medical history data, current medical conditions and medications, physical and cognitive function data, and blood samples were collected via in-person visits and phone questionnaires for all subjects at the time of enrollment as described elsewhere [Newman 2011; Elo 2013]. Participants have been followed-up annually to track vital and health status. All subjects provided informed consent approved by the field centers IRB.

#### The Danish Twin Registry

The Danish Twin Registry sample included 1432 individuals who completed the Symbol Digit Substitution Task (SDST) and were collected as part of the study of Middle Aged Danish Twins (MADT, N=1146) and the Longitudinal Study of Aging Danish Twins (LSADT, N=286). MADT was initiated in 1998 and includes 4,314 twins randomly chosen from the birth years 1931-1952. Surviving participants were revisited from 2008 to 2011, where the blood samples used in the present study were collected. The survey data used in the present study was obtained from the intake survey undertaken in 1998 [Pedersen 2019]. LSADT was initiated in 1995 and includes twins aged 70 years and older. Follow-up assessments were conducted every second year until 2005. The individuals included here all participated in the 1997 assessment, where blood samples were donated by same sex twin pairs, and in 1999 participants completed the processing speed test used in the present study [Pedersen 2019]. Written informed consents were obtained from all participants. Collection and use of biological material, survey and registry information were approved by the Regional Scientific Ethical Committees for Southern Denmark (MADT: S-VF-19980072, LSADT: S-VF-20040241), and the study is registered in SDU’s internal list (notification no. 11.108) and complies with the rules in the General Data Protection Regulation.

### Genetic data

DNA samples of 4577 LLFS participants were genotyped at the Center for Inherited Disease Research (CIDR) using the Illumina Omni 2.5 platform, and genotype calls were cleaned following a strict quality control process described in [Bae 2013]. The genotype data were imputed with Michigan Imputation Server to the HRC panel (version r1.1 2016) of 64,940 haplotypes with 39,635,008 sites. [Das et al. 2016]. Genome-wide genotype data are available from dbGaP (**dbGaP Study Accession:** phs000397.v1.p1). _The Danish Twin Registry samples were genotyped using the Illumina Infinium PsychArray (Illumina San Diego, CA, USA). Pre-imputation quality control included filtering SNPs on genotype call rate <98%, HWE *P*<10^−6^, and MAF = 0, and individuals on sample call rate <99%, relatedness and gender mismatch. Prephasing and imputation to the 1000 Genomes phase 3 reference panel was performed using IMPUTE2 version 2.3.2.

### Assessment of Processing speed

The assessment of processing speed in participants in LLFS was conducted by administering the Digital Symbol Substitution Test (DSST) during the in-person visit at enrollment (2006—2009) and the second in-person visit conducted between 2014 and 2017. The test consists of recoding numbers into symbols using a look-up table, and the test score represents the number of substitutions completed in 90 seconds. The test score ranges from 0 to 100. Table 1 provides a summary table of the participants in LLFS included in the final analysis. The administration of the cognitive speed test in the two Danish cohorts were completed in 1998 (MADT) and 1999 (LSADT). Processing speed in the Danish twins was assessed by using a cognitive speed test similar to the DSST, in which symbols were to be replaced with numbers (Symbol Digit Substitution Task (SDST)) within 90 seconds, and the test score represents the number of symbol substitutions. Table 1 provides a summary table of the two Danish cohorts included in the replication analysis.

**Table 1.** Demographic characteristics of the LLFS cohort at baseline measurement and at the second visit, and the MADT and the LSADT cohorts.

|  | LLFS baseline<br>(N=4207) | LLFS offspring baseline<br>(N=2188) | LLFS offspring spouse baseline<br>(N=733) | LLFS proband baseline<br>(N=1122) | LLFS proband spouse baseline<br>(N=164) |
| --- | --- | --- | --- | --- | --- |
| DSST score, mean ± sd<br>[min, max] | 43.94 ± 15.94<br>[0, 93] | 51.00 ± 12.36<br>[0, 93] | 48.69 ± 12.97<br>[1, 85] | 28.42 ± 12.36<br>[0, 73] | 34.65 ± 13.43<br>[0, 67] |
| Age, mean ± sd<br>[min, max] | 69.86 ± 15.20<br>[25, 109] | 61.09 ± 8.20<br>[32, 88] | 61.71 ± 8.54<br>[25, 88] | 90.29 ± 6.39<br>[49, 109] | 83.68 ± 7.24<br>[55, 98] |
| Female, n (%) | 2291 (54.46) | 1268 (57.95) | 337 (45.98) | 560 (49.91) | 126 (76.83) |
| Education years, mean ± sd<br>[min, max] | 11.75 ± 3.48<br>[1, 17] | 12.57 ± 3.01<br>[2, 17] | 12.01 ± 3.35<br>[3, 17] | 10.15 ± 3.78<br>[1, 17] | 10.49 ± 3.64<br>[2, 16] |

|  | LLFS 2nd visit<br>(N=2160) | LLFS offspring 2nd visit<br>(N=1417) | LLFS offspring spouse 2nd visit<br>(N=441) | LLFS proband 2nd visit<br>(N=253) | LLFS proband spouse 2nd visit<br>(N=49) |
| --- | --- | --- | --- | --- | --- |
| DSST score, mean ± sd<br>[min, max] | 44.88 ± 14.13<br>[0, 90] | 48.43 ± 12.23<br>[0, 90] | 45.16 ± 12.22<br>[4, 78] | 26.71 ± 12.45<br>[0, 68] | 33.73 ± 12.14<br>[0, 59] |
| Age, mean ± sd<br>[min, max] | 72.30 ± 11.18<br>[43, 106] | 68.76 ± 7.77<br>[43, 94] | 69.90 ± 7.81<br>[47, 90] | 93.32 ± 6.52<br>[57, 106] | 88.02 ± 6.90<br>[71, 99] |
| Female, n (%) | 1196 (55.37) | 810 (57.16) | 206 (46.71) | 137 (54.15) | 43 (87.76) |
| Education years, mean ± sd<br>[min, max] | 12.25 ± 3.23<br>[2, 17] | 12.68 ± 2.99<br>[2, 17] | 12.09 ± 3.40<br>[3, 17] | 10.51 ± 3.40<br>[3, 17] | 10.35 ± 3.62<br>[2, 15] |

|  | MADT<br>(N=1146) | LSADT<br>(N=286) | Danish combined (MADT+LSADT)<br>(N=1432) |
| --- | --- | --- | --- |
| SDST score, mean ± sd<br>[min, max] | 49.14 ± 10.53<br>[8, 81] | 28.43 ± 12.67<br>[0, 61] | 45.00 ± 13.76<br>[0, 81] |
| Age, mean ± sd<br>[min, max] | 55.68 ± 5.97<br>[45, 68] | 79.98 ± 3.72<br>[75, 96] | 60.54 ± 11.21<br>[45, 96] |
| Female, n (%) | 550 (47.99) | 190 (66.43) | 740 (51.68) |
| Education, n (%) |  |  |  |
| <7 years - pro training | 0 (0) | 22 (7.89) | 22 (1.57) |
| 7+ years -pro training | 234 (20.86) | 113 (40.50) | 347 (24.77) |
| 8-9 years + pro training | 299 (26.65) | 65 (23.30) | 364 (25.98) |
| 9+ years + pro training | 262 (23.35) | 21 (7.53) | 283 (20.20) |
| short-medium college/UNI | 249 (22.19) | 47 (16.85) | 296 (21.13) |
| long college/UNI | 78 (6.95) | 11 (3.94) | 89 (6.35) |

### Genome-wide association study

We removed duplicate, monomorphic and ambiguous SNPs (A/T, C/G) and selected SNPs with high imputation quality (Rsq > 0.8), which resulted in 15,346,220 clean SNPs. The associations between baseline DSST scores and dosages of SNPs were tested using a linear mixed effect regression model to account for cryptic relationships. The genetic relations and population stratification were analyzed by the R/3.6.0 packages PC-Relate and PC-Air following the method by Conomos [Conomos 2015 and 2016]. The regression analysis was adjusted by age at enrollment, sex, education, and the first four genome-wide principal components. Significance of the associations was tested using the Wald test and the score test. The association analyses were conducted using the R/3.6.0 package GENESIS [Gogarten 2019] and the results were annotated using ANNOVAR [Wang 2010]. The baseline analysis as described was performed by a recently developed GWAS pipeline [Song 2021]. The associations between DSST scores at both visit 1 and visit 2 were tested using a linear mixed effect model to account for both cryptic relationships and repeated measures as implemented in the R/3.6.0 package GMMAT [Chen 2016]. This analysis was adjusted by age at enrollment, age differences between two visits, sex, education, and the first four genome-wide principal components. The score test was used for each SNP using genotype count first, and the Wald test was used on the top associated SNPs using expected dosages to obtain comparable effect sizes to the baseline GWAS.

In the replication study, the analyses were carried out using a linear regression model adjusted by sex, interview age, education, cohort (LSADT=1, MADT=2), and twin pair number as a random effect to account for the correlation between twins from the same pair. Analyses were only done on the combined sample. The regression was not adjusted by principal components since the Danish population is highly homogeneous. Among the 20 genome-wide significant SNPs discovered by the LLFS GWAS, six SNPs with imputation quality score > 0.3 were tested.

### Gene-based analysis

Burden tests were performed on the two genes Retinoic Acid Receptor Beta (*RARB*) and Thyroid Hormone Receptor Beta (*THRB*), tagged by 17 protective rare genetic variants on chromosome 3 in LLFS following the method by Madsen [Madsen 2009]. 3029 rare variants within *RARB* and 1356 rare variants within *THRB* with minor allele frequency < 0.01 were aggregated by weighted sum. The weights were assigned considering the effect direction using the following formula:

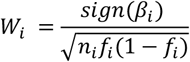

where *β_i_* is the effect of *SNP_i_*, *n_i_* is the total number of subjects, and *f_i_* is the MAF of *SNP_i_*. The summary score of one gene for an individual was calculated as:

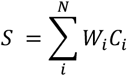

where *C_i_* is the number of copies of *SNP_i_* on that gene.

Summary scores for *RARB* and *THRB* genes can be found in **Supplement Table 4**. The associations between the scores and DSST were tested using the same model as the single SNP association test at baseline.

## Results

### Demographic characteristics

We identified 4207 subjects in LLFS with genotype data, and valid results of the DSST who were included in the GWAS of baseline DSST. Only 2160 of these subjects participated in the second assessment, due to mortality or dropout (88% in the oldest generation and 14% in the youngest generation). **Table 1** displays the characteristics of the 4207 LLFS participants included in the analysis of DSST at baseline, and the 2160 participants at the second visit. The enrollment age of LLFS participants ranged from 25 to 109 years with mean age of 70 years old. This group included 1286 participants from the oldest generation, with 1122 members from long-lived families (mean age = 90), and 164 spousal controls (mean age = 84). The LLFS group also included 2921 participant from the offspring generation, with 2188 members from long-lived families (mean age = 61) and 733 spousal controls (mean age = 62). The MADT included only middle-aged twins with age ranging between 45 and 68 years. The LSADT included older twins with ages 75-96 years old. The average speed score was 49.14 measured by SDST in MADT, 43.94 by DSST in LLFS, and 28.43 by SDST in LSADT, and it declined with increasing average age. The pooled Danish cohort (MADT+LSADT) with average age 60 and speed score 45.00 was comparable to the speed score in the LLFS cohort.

### Genome-wide association Study

Analyses of the baseline DSST scores in the LLFS identified 20 SNPs in four distinct genomic regions that passed the genome-wide significance threshold with p-value < 5×10^−8^ (**Table 2**). The complete list of SNPs with “suggestive” level of significance (p-value < 5×10^−6^) is in **Supplement Table 1**. All the 20 genome-wide level significant SNPs were rare variants with MAF < 0.01. 18 SNPs showed a positive effect on the DSST score, and carriers of the rare alleles had a DSST score greater than 20 points. 17 SNPs were within a region of 1.5 MB on chromosome 3 that includes two genes that belong to a family of thyroid hormone receptors: Retinoic Acid Receptor Beta (*RARB*) and Thyroid Hormone Receptor Beta (*THRB*), as well as miR-4792 which may regulate a number of mitochondria-related genes, among other targets [Liu 2019]. An additional rare protective variant rs58169119 was in an intergenic region on chromosome 8, and the rare allele of this variant was associated with an average increase of 28 points in DSST. The two deleterious rare SNPs also showed very large negative effects, whereby the rare alleles were associated with a decrease in DSST score > 50, specifically: SNP rs146299120 located in an intergenic region on chromosome 9, and a SNP without rs ID located on chromosome 18 in an intergenic region of the Deleted in Colorectal Cancer (*DCC*) gene, which encodes the netrin 1 receptor, that is involved in nervous system development [Boyer and Gupton 2018; Duman-Scheel M 2009]. Six of the rare variants with imputation score R^2^ > 0.3 were analyzed in the combined Danish twin cohort (**Table 2; Supplement Table 3**). The analyses confirmed the association of four SNPs on chromosome 3 with SDST score after Bonferroni correction for 3 independent loci (p-value < 0.017). The SNP rs7623455 had the lowest p-value = 2.7×10^−18^ with large positive effect size (β=10.7) in the combined Danish cohort. All the rare variants that replicated in the Danish cohort also showed positive effects and increase SDST scores by more than 8 points.

**Table 2.**
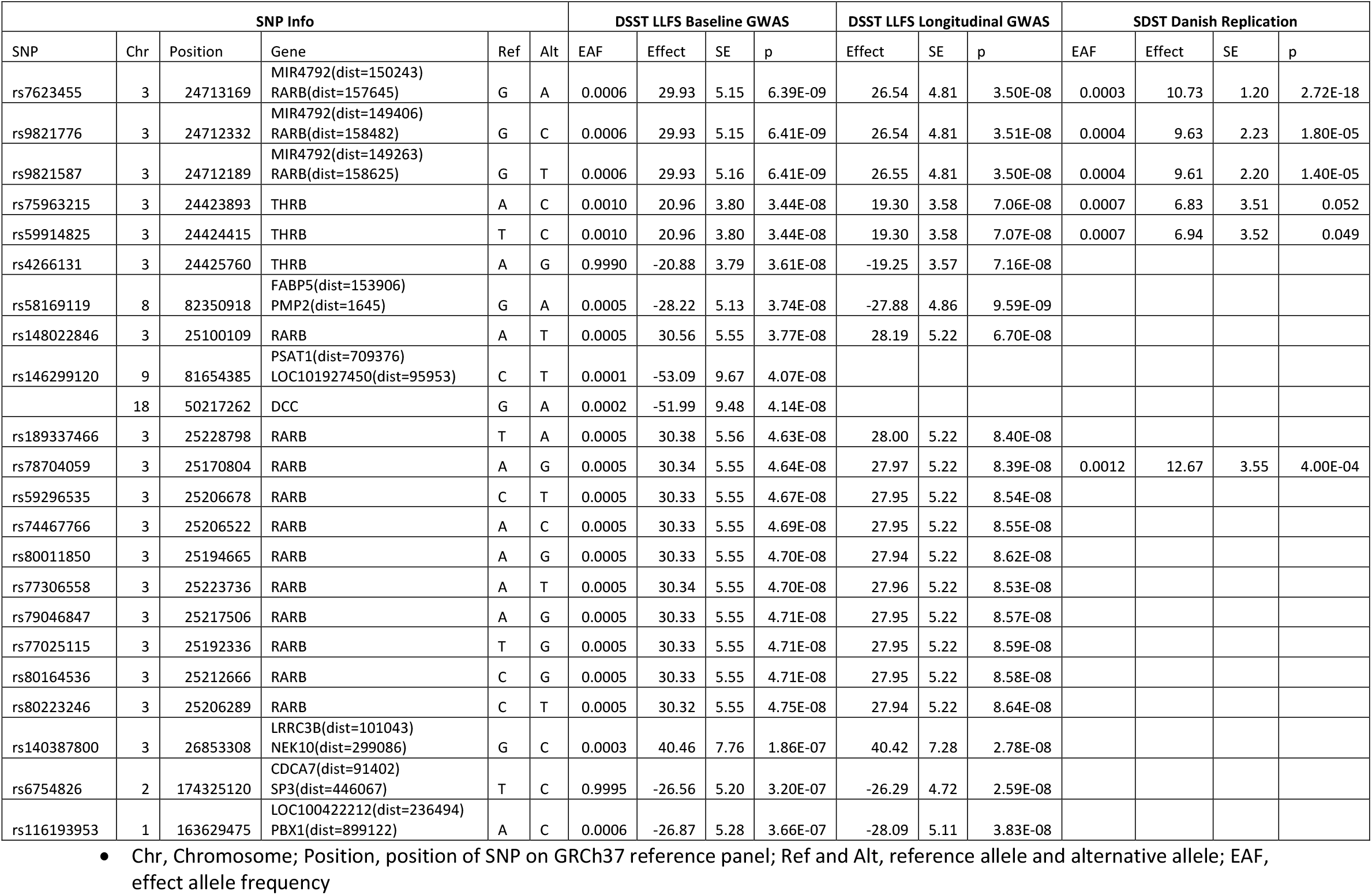
SNPs with p-values < 5×10^−8^ from the baseline or longitudinal GWAS in LLFS and replication in the Danish combined cohort.

| SNP Info |  |  |  |  |  | DSST LLFS Baseline GWAS |  |  |  | DSST LLFS Longitudinal GWAS |  |  | SDST Danish Replication |  |  |  |
| --- | --- | --- | --- | --- | --- | --- | --- | --- | --- | --- | --- | --- | --- | --- | --- | --- |
| SNP | Chr | Position | Gene | Ref | Alt | EAF | Effect | SE | p | Effect | SE | p | EAF | Effect | SE | p |
| rs7623455 | 3 | 24713169 | MIR4792(dist=150243)<br>RARB(dist=157645) | G | A | 0.0006 | 29.93 | 5.15 | 6.39E-09 | 26.54 | 4.81 | 3.50E-08 | 0.0003 | 10.73 | 1.20 | 2.72E-18 |
| rs9821776 | 3 | 24712332 | MIR4792(dist=149406)<br>RARB(dist=158482) | G | C | 0.0006 | 29.93 | 5.15 | 6.41E-09 | 26.54 | 4.81 | 3.51E-08 | 0.0004 | 9.63 | 2.23 | 1.80E-05 |
| rs9821587 | 3 | 24712189 | MIR4792(dist=149263)<br>RARB(dist=158625) | G | T | 0.0006 | 29.93 | 5.16 | 6.41E-09 | 26.55 | 4.81 | 3.50E-08 | 0.0004 | 9.61 | 2.20 | 1.40E-05 |
| rs75963215 | 3 | 24423893 | THRB | A | C | 0.0010 | 20.96 | 3.80 | 3.44E-08 | 19.30 | 3.58 | 7.06E-08 | 0.0007 | 6.83 | 3.51 | 0.052 |
| rs59914825 | 3 | 24424415 | THRB | T | C | 0.0010 | 20.96 | 3.80 | 3.44E-08 | 19.30 | 3.58 | 7.07E-08 | 0.0007 | 6.94 | 3.52 | 0.049 |
| rs4266131 | 3 | 24425760 | THRB | A | G | 0.9990 | -20.88 | 3.79 | 3.61E-08 | -19.25 | 3.57 | 7.16E-08 |  |  |  |  |
| rs58169119 | 8 | 82350918 | FABP5(dist=153906)<br>PMP2(dist=1645) | G | A | 0.0005 | -28.22 | 5.13 | 3.74E-08 | -27.88 | 4.86 | 9.59E-09 |  |  |  |  |
| rs148022846 | 3 | 25100109 | RARB | A | T | 0.0005 | 30.56 | 5.55 | 3.77E-08 | 28.19 | 5.22 | 6.70E-08 |  |  |  |  |
| rs146299120 | 9 | 81654385 | PSAT1(dist=709376)<br>LOC101927450(dist=95953) | C | T | 0.0001 | -53.09 | 9.67 | 4.07E-08 |  |  |  |  |  |  |  |
|  | 18 | 50217262 | DCC | G | A | 0.0002 | -51.99 | 9.48 | 4.14E-08 |  |  |  |  |  |  |  |
| rs189337466 | 3 | 25228798 | RARB | T | A | 0.0005 | 30.38 | 5.56 | 4.63E-08 | 28.00 | 5.22 | 8.40E-08 |  |  |  |  |
| rs78704059 | 3 | 25170804 | RARB | A | G | 0.0005 | 30.34 | 5.55 | 4.64E-08 | 27.97 | 5.22 | 8.39E-08 | 0.0012 | 12.67 | 3.55 | 4.00E-04 |
| rs59296535 | 3 | 25206678 | RARB | C | T | 0.0005 | 30.33 | 5.55 | 4.67E-08 | 27.95 | 5.22 | 8.54E-08 |  |  |  |  |
| rs74467766 | 3 | 25206522 | RARB | A | C | 0.0005 | 30.33 | 5.55 | 4.69E-08 | 27.95 | 5.22 | 8.55E-08 |  |  |  |  |
| rs80011850 | 3 | 25194665 | RARB | A | G | 0.0005 | 30.33 | 5.55 | 4.70E-08 | 27.94 | 5.22 | 8.62E-08 |  |  |  |  |
| rs77306558 | 3 | 25223736 | RARB | A | T | 0.0005 | 30.34 | 5.55 | 4.70E-08 | 27.96 | 5.22 | 8.53E-08 |  |  |  |  |
| rs79046847 | 3 | 25217506 | RARB | A | G | 0.0005 | 30.33 | 5.55 | 4.71E-08 | 27.95 | 5.22 | 8.57E-08 |  |  |  |  |
| rs77025115 | 3 | 25192336 | RARB | T | G | 0.0005 | 30.33 | 5.55 | 4.71E-08 | 27.95 | 5.22 | 8.59E-08 |  |  |  |  |
| rs80164536 | 3 | 25212666 | RARB | C | G | 0.0005 | 30.33 | 5.55 | 4.71E-08 | 27.95 | 5.22 | 8.58E-08 |  |  |  |  |
| rs80223246 | 3 | 25206289 | RARB | C | T | 0.0005 | 30.32 | 5.55 | 4.75E-08 | 27.94 | 5.22 | 8.64E-08 |  |  |  |  |
| rs140387800 | 3 | 26853308 | LRRC3B(dist=101043)<br>NEK10(dist=299086) | G | C | 0.0003 | 40.46 | 7.76 | 1.86E-07 | 40.42 | 7.28 | 2.78E-08 |  |  |  |  |
| rs6754826 | 2 | 174325120 | CDCA7(dist=91402)<br>SP3(dist=446067) | T | C | 0.9995 | -26.56 | 5.20 | 3.20E-07 | -26.29 | 4.72 | 2.59E-08 |  |  |  |  |
| rs116193953 | 1 | 163629475 | LOC100422212(dist=236494)<br>PBX1(dist=899122) | A | C | 0.0006 | -26.87 | 5.28 | 3.66E-07 | -28.09 | 5.11 | 3.83E-08 |  |  |  |  |

- Chr, Chromosome; Position, position of SNP on GRCh37 reference panel; Ref and Alt, reference allele and alternative allele; EAF, effect allele frequency

The GWAS of the repeated assessment of DSST identified 7 SNPs reaching genome-wide significance level (**Table 2; Supplement Table 2**). Four SNPs were among the 20 SNPs identified in the baseline analysis. The remaining three SNPs with p-values < 5×10^−8^ in the longitudinal analysis all had p-values < 5×10^−7^ in the baseline analysis (**Table 2**). SNP rs140387800 was in an intergenic region on chromosome 3 with large protective effect size (β=40.5). SNP rs6754826 was in an intergenic region on chromosome 2 and the rare allele was associated with a 26.6-point increase in DSST. The SNP rs116193953 was in an intergenic region on chromosome 1 and the rare allele was associated with an average decrease of 28 points in DSST.

### Gene-based analysis

The GWAS results showed strong evidence that two genes, *RARB* and *THRB*, on chromosome 3 were potentially related to processing speed. We therefore performed Burden tests on these two genes in LLFS. The weighted score for *RARB* ranged from −3.29 to 12.98, with mean 0.052 by aggregating 3029 rare alleles as described in the Methods section. The weighted score for *THRB* ranged from –3.26 to 30.50, with mean 0.016 by aggregating 1356 rare alleles. Both gene scores were associated with DSST scores with p-value < 0.025 after Bonferroni correction for two tests. The two genes also showed overall protective effects on processing speed. Each unit increase in *RARB* score increased DSST by 2.13 points (p-value = 7.7×10^−17^) and each unit increase in *THRB* scores increased DSST by 0.96 points (p-value = 1.3×10^−8^) (**Table 3**).

**Table 3.** Association results of the two gene summary scores in LLFS.

| Gene | Chr | start | end | Number of rare variants (MAF < 0.01) | Effect | SE | p |
| --- | --- | --- | --- | --- | --- | --- | --- |
| RARB | 3 | 24829321 | 25597932 | 3029 | 2.13 | 0.25 | $7.68 \times 10^{-17}$ |
| THRB | 3 | 24117153 | 24495708 | 1356 | 0.96 | 0.17 | $1.28 \times 10^{-8}$ |

### Replication of Published Results

We looked at a replication of the recently published GWAS of global cognition [Davies 2019]. Out of the11,600 genome-wide level significant SNPs in that study, 11,564 SNPs were present in the LLFS, and only 12 SNPs passed the Bonferroni corrected p-value threshold and show concordant effects (1.15×10^−4^, 434 independent significant SNPs). These 12 SNPs were within a 10kb intergenic region on chromosome 1 and their associations in LLFS are summarized in **Table 4**. None of our results showed a genome-wide significant association with global cognition.

**Table 4.** SNPs replicated in LLFS baseline GWAS from published results.

| SNP Info |  |  |  |  |  | GCF as Discovery |  |  | DSST LLFS GWAS as Replication |  |  |  |
| --- | --- | --- | --- | --- | --- | --- | --- | --- | --- | --- | --- | --- |
| SNP | Chr | Position | Gene | Ref | Alt | EAF | Z_Score | P | EAF_LLFS | Effect | SE | p |
| rs6659327 | 1 | 183405853 | NMNAT2(dist=18219),SMG7-AS1(dist=24158) | G | A | 0.4858 | 5.571 | 2.53E-08 | 0.5112 | 0.985 | 0.235 | 2.71E-05 |
| rs6684015 | 1 | 183405794 | NMNAT2(dist=18160),SMG7-AS1(dist=24217) | G | A | 0.5141 | -5.55 | 2.86E-08 | 0.4889 | -0.984 | 0.235 | 2.77E-05 |
| rs6424896 | 1 | 183405648 | NMNAT2(dist=18014),SMG7-AS1(dist=24363) | C | T | 0.4858 | 5.562 | 2.67E-08 | 0.5110 | 0.983 | 0.235 | 2.84E-05 |
| rs4047801 | 1 | 183415318 | NMNAT2(dist=27684),SMG7-AS1(dist=14693) | G | A | 0.4859 | 5.539 | 3.04E-08 | 0.5112 | 0.969 | 0.235 | 3.71E-05 |
| rs7550777 | 1 | 183425047 | NMNAT2(dist=37413),SMG7-AS1(dist=4964) | C | T | 0.4452 | -5.968 | 2.41E-09 | 0.4196 | -0.962 | 0.237 | 4.83E-05 |
| rs4652802 | 1 | 183426758 | NMNAT2(dist=39124),SMG7-AS1(dist=3253) | G | A | 0.555 | 5.981 | 2.22E-09 | 0.5806 | 0.961 | 0.237 | 4.89E-05 |
| rs3965889 | 1 | 183407953 | NMNAT2(dist=20319),SMG7-AS1(dist=22058) | G | A | 0.5535 | 5.928 | 3.06E-09 | 0.5795 | 0.938 | 0.236 | 7.18E-05 |
| rs10911334 | 1 | 183405382 | NMNAT2(dist=17748),SMG7-AS1(dist=24629) | G | C | 0.4465 | -5.923 | 3.16E-09 | 0.4205 | -0.937 | 0.237 | 7.63E-05 |
| rs3856126 | 1 | 183416374 | NMNAT2(dist=28740),SMG7-AS1(dist=13637) | G | T | 0.4473 | -5.978 | 2.26E-09 | 0.4202 | -0.935 | 0.237 | 7.69E-05 |
| rs12039578 | 1 | 183413886 | NMNAT2(dist=26252),SMG7-AS1(dist=16125) | G | A | 0.4466 | -5.978 | 2.26E-09 | 0.4206 | -0.930 | 0.237 | 8.39E-05 |
| rs11808408 | 1 | 183415589 | NMNAT2(dist=27955),SMG7-AS1(dist=14422) | G | A | 0.4464 | -5.98 | 2.23E-09 | 0.4206 | -0.929 | 0.237 | 8.60E-05 |
| rs17558314 | 1 | 183415188 | NMNAT2(dist=27554),SMG7-AS1(dist=14823) | C | T | 0.4463 | -5.961 | 2.51E-09 | 0.4206 | -0.929 | 0.237 | 8.64E-05 |

- Chr, Chromosome; Position, position of SNP on GRCh37 reference panel; Ref and Alt, reference allele and alternative allele; EAF, effect allele frequency

## Discussion

Our GWAS of the processing speed (based on DSST scores) measured in 4207 LLFS participants, discovered 20 genome-wide significant SNPs (p-value < 5×10^−8^) on chromosomes 3, 8, 9 and 18. Sixteen of these SNPs formed a cluster of rare protective variation on chromosome 3. These rare variants showed very large effect sizes compared to common variants that reached only suggestive significance threshold (p-value < 5×10^−6^) [Supplement Table 1]. All sixteen rare SNPs were confirmed in the longitudinal analysis with comparable effect sizes [Table 2]. We also replicated the association of four of the sixteen variants, rs7623455, rs9821776, rs9821587 and rs78704059, in an independent sample of Danish ancestry (p-values < 0.017). Note that SNPs shown in Table 2 were rare and independent in White participants; however, the same SNPs may potentially be common and in LD in different populations/race groups. E.g., rs9821776 and rs9821587 from Table 2 are in complete LD in individuals of African ancestry, according to LDlink, NIH supported portal.

We also investigated the association of 11,564 SNPs that reached a genome-wide significant association with global cognition in a recent GWAS with 300,486 individuals [Davies 2019]. Although our analysis focused on the specific cognitive domain of processing speed and attention assessed by the DSST score, we replicated the association of 12 common SNPs in an intergenic region on chromosome 1 with concordant effect directions [Table 4]. None of the rare variants we found associated with DSST were reported in the analysis by [Davies 2019], but their study dropped SNPs with minor allele count < 25.

Another GWAS of 30,576 individuals of European ancestry and 5758 individuals of African ancestry investigated the associations of rare variants with processing speed but did not identify any genome-wide significant associations [Bressler 2021]. Their gene-based analysis adjusted for age and gender discovered one genome-wide significant result: ring finger protein 19A (*RNF19A*). We attempted replication of this association in our data using our method and adjusting for age, gender, education and first 4 PCs. The association reached a p-value of 7.6×10^−4^ for this gene, consistent with their result. A possible reason that our study of only 4207 individuals in LLFS was able to identify genome-wide significant rare variants could be that the family based design produced a dataset enriched for protective rare variants, thus increasing our discovery power. The allele counts per 10,000 alleles of the top rare variants were doubled or tripled comparing to the general European population reported on dbSNP.

Although the most significant variants were very rare, they were notably concentrated in/near two functionally related genes on chromosome 3, *RARB* and *THRB.* The *RARB* and *THRB* belong to the family of thyroid hormone receptors, which could potentially contribute to their associations with the processing speed. For example, *THRB* may mediate the biological activities of thyroid hormone and through this influence the speed of metabolism, which in turn may influence the processing speed. Some studied also linked reduced thyroid function during aging with a decline in cognition [Bégin 2008; Jasim 2017]. Additional biological effects of these genes could play a role in their connection to the processing speed as well. E.g., the *RARB* encodes a receptor binding the retinoic acid, the biologically active form of vitamin A. The deficiency of vitamin A was associated with the cognitive decline in aging [Pallet 2015; Wołoszynowska-Fraser 2020]. The *RARB* rare variant burden was associated with severe cognitive deficits in schizophrenia [Reay 2020]. The GTEX portal shows that there are several expression quantitative trait loci in the brain frontal cortex for the *RARB* gene, warranting further exploration of the observed associations. The gene-based analysis also confirmed that *RARB* and *THRB* genes are both associated with DSST scores and have an overall beneficial effect on the processing speed.

In addition, the three top SNPs on chromosome 3 shown in Table 2 are close to miR-4792. Targets of this microRNA include a number of mitochondria-related genes [Liu 2019]. This points to a possible role of miR-4792 in energy metabolism and ATP production, which in turn may also contribute to the processing speed. This connection is, however, underexplored in the literature, and is purely hypothetical.

A potential limitation of our results is that we used imputed data for rare variants. We were extremely conservative and only SNPs with a high quality score were retained in the analysis. The LLFS has recently completed Whole Genome Sequencing (WGS) data in approximately 3,500 participants. We compared the top imputed SNPs to the WGS data in the common set of individuals with both data types, and observed a perfect concordance. Therefore, these rare variants imputed to the HRC panel with high quality appear to be trustworthy.

In summary, our study provided new evidence that rare genetic variants can play an important role in cognitive aging. We discovered 20 rare protective SNPs located in/near two genes on chromosome 3, *RARB* and *THRB*, belonging to the family of thyroid hormone receptors that may help aging individuals maintain younger processing speed, potentially through preserving the speed of metabolism. This explanation, however, needs to be thoroughly explored before these genes and relevant biological pathways could become new targets for therapies helping older adults to maintain the processing speed without undesirable trade-offs, such as depletion of stem cell reserves, damage accumulation, etc. [reviewed in Ukraintseva et al. 2021]. In future research, the gene-based analysis scanning through the whole genome might detect more genomic regions associated with the processing speed with increased power.

**Figure 1.**
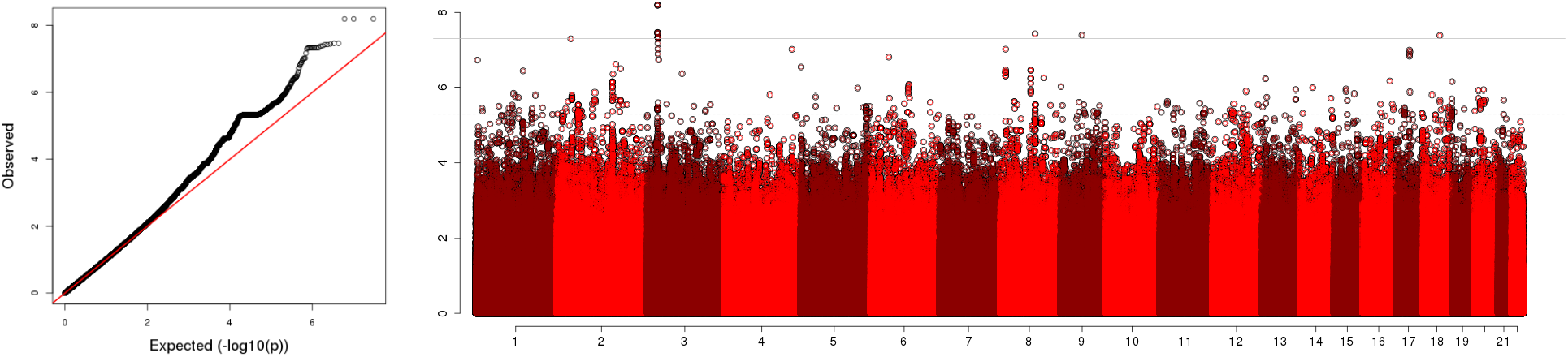
QQ-plot (left) and Manhattan plot (right) of the genome-wide association with baseline DSST scores in LLFS. The x-axis of QQ-plot reports the expected −log10 (p-value) and the y-axis reports the observed −log10 (p-value). The x-axis of Manhattan plot reports chromosomes and coordinates within chromosomes. The y-axis reports the −log10 (p-value). The genomic control parameter is 1.04, indicating population stratification addressed properly.

**Figure 2.**
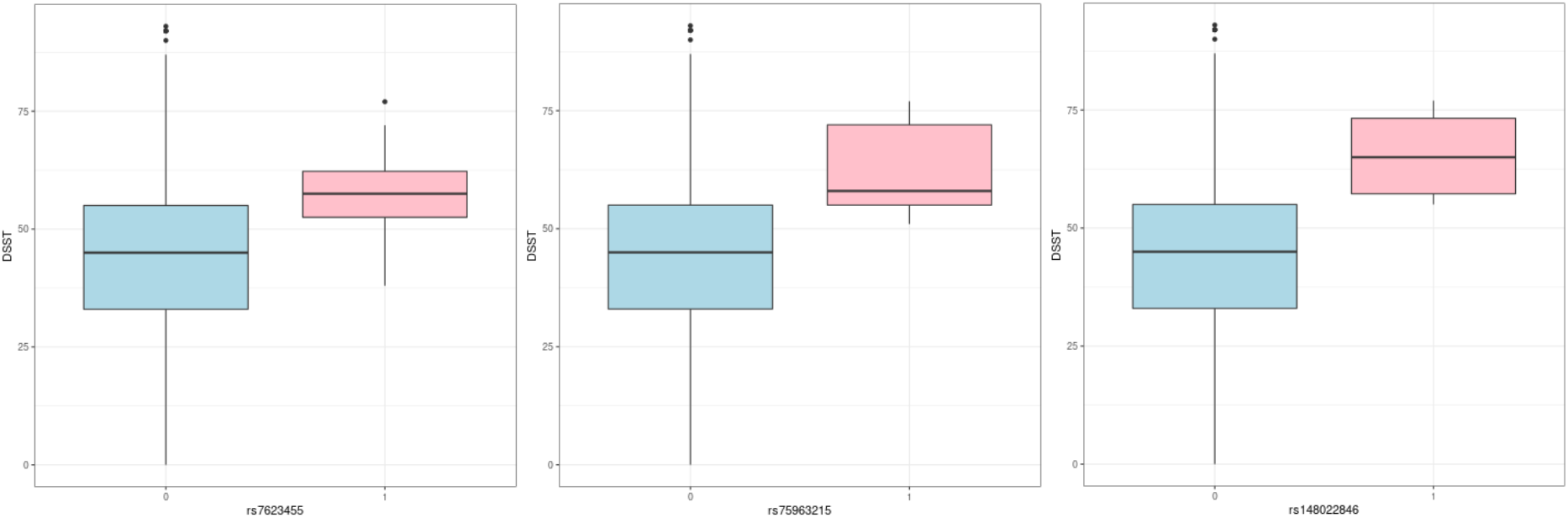
Examples of boxplots of baseline DSST scores by SNP status. The x-axis reports the number of SNPs carried by individuals in this group. The y-axis reports the baseline DSST scores. Carriers of all three SNPs show better average performance in the DSST assessment.

## Acknowledgments

Supported by the National Institute on Aging (K01AG057798 to S.L.A., 5U19AG063893 5U01AG023749 to S.C., 5U01AG023755 to T.T.P., U19AG063893 R01AG062623 to S.U. and K.A., 5U01AG023712, 5U01AG023744, 5U01AG023746); Genotyping of the Danish twins was conducted by the SNP&SEQ Technology Platform, Science for Life Laboratory, Uppsala, Sweden (http://snpseq.medsci.uu.se/genotyping/snp-services/) and supported by NIH R01 AG037985 (Pedersen).

