## Supplemental_table1 for "Rare Genetic Variants Correlate with Better Processing Speed"

**Supplement Table1. SNPs with Wald p-values < $1\times10^{-6}$ from the baseline GWAS in LLFS.**

Chr, Chromosome; Position, position of SNP on GRCh37 reference panel; Ref and Alt, reference allele and alternative allele; EAF, effect allele frequency; MAC, minor allele count; Wald.p, p-values from Wald test; Score.p, p-values from Score test

| SNP | Chr | Position | Info | Gene | Ref | Alt | EAF | MAC | Effect | SE | Wald.p | Score.p |
|---|---|---|---|---|---|---|---|---|---|---|---|---|
| rs7623455 | 3 | 24713169 | IMPUTED | MIR4792(dist=150243),RARB(dist=157645) | G | A | 0.0005624 | 5 | 29.9275872 | 5.15449032 | 6.39E-09 | 7.30E-09 |
| rs9821776 | 3 | 24712332 | IMPUTED | MIR4792(dist=149406),RARB(dist=158482) | G | C | 0.0005624 | 5 | 29.9255668 | 5.15449047 | 6.41E-09 | 7.32E-09 |
| rs9821587 | 3 | 24712189 | IMPUTED | MIR4792(dist=149263),RARB(dist=158625) | G | T | 0.00056204 | 5 | 29.9310431 | 5.15549071 | 6.41E-09 | 7.32E-09 |
| rs75963215 | 3 | 24423893 | IMPUTED | THRB | A | C | 0.00095139 | 8 | 20.9604586 | 3.79900493 | 3.44E-08 | 3.84E-08 |
| rs59914825 | 3 | 24424415 | GENOTYPED | THRB | T | C | 0.00095127 | 8 | 20.9604289 | 3.79901212 | 3.44E-08 | 3.84E-08 |
| rs4266131 | 3 | 24425760 | IMPUTED | THRB | A | G | 0.99902817 | 8 | -20.876462 | 3.78950526 | 3.61E-08 | 4.02E-08 |
| rs58169119 | 8 | 82350918 | IMPUTED | FABP5(dist=153906),PMP2(dist=1645) | G | A | 0.00047076 | 4 | -28.215725 | 5.12758589 | 3.74E-08 | 4.16E-08 |
| rs148022846 | 3 | 25100109 | IMPUTED | RARB | A | T | 0.00048205 | 4 | 30.5589111 | 5.55472799 | 3.77E-08 | 4.19E-08 |
| rs146299120 | 9 | 81654385 | IMPUTED | PSAT1(dist=709376),LOC101927450(dist=95953) | C | T | 0.00014832 | 1 | -53.09392 | 9.67454874 | 4.07E-08 | 4.52E-08 |
| | 18 | 50217262 | IMPUTED | DCC | G | A | 0.00016306 | 1 | -51.991479 | 9.4790259 | 4.14E-08 | 4.60E-08 |
| rs189337466 | 3 | 25228798 | IMPUTED | RARB | T | A | 0.00047504 | 4 | 30.3770174 | 5.55859522 | 4.63E-08 | 5.14E-08 |
| rs78704059 | 3 | 25170804 | IMPUTED | RARB | A | G | 0.00047611 | 4 | 30.340652 | 5.55230542 | 4.64E-08 | 5.15E-08 |
| rs59296535 | 3 | 25206678 | IMPUTED | RARB | C | T | 0.00047659 | 4 | 30.3334109 | 5.55222203 | 4.67E-08 | 5.19E-08 |
| rs74467766 | 3 | 25206522 | IMPUTED | RARB | A | C | 0.00047682 | 4 | 30.3303663 | 5.55222055 | 4.69E-08 | 5.20E-08 |
| rs80011850 | 3 | 25194665 | IMPUTED | RARB | A | G | 0.00047742 | 4 | 30.3287208 | 5.5521981 | 4.70E-08 | 5.21E-08 |
| rs77306558 | 3 | 25223736 | IMPUTED | RARB | A | T | 0.00047659 | 4 | 30.3378677 | 5.553928 | 4.70E-08 | 5.21E-08 |
| rs79046847 | 3 | 25217506 | IMPUTED | RARB | A | G | 0.00047682 | 4 | 30.3266684 | 5.55223637 | 4.71E-08 | 5.22E-08 |
| rs77025115 | 3 | 25192336 | IMPUTED | RARB | T | G | 0.00047778 | 4 | 30.3260331 | 5.552182 | 4.71E-08 | 5.22E-08 |
| rs80164536 | 3 | 25212666 | IMPUTED | RARB | C | G | 0.00047694 | 4 | 30.3252941 | 5.55222455 | 4.71E-08 | 5.23E-08 |
| rs80223246 | 3 | 25206289 | IMPUTED | RARB | C | T | 0.0004773 | 4 | 30.31773 | 5.55223107 | 4.75E-08 | 5.27E-08 |
| rs183090477 | 2 | 36794491 | IMPUTED | FEZ2 | T | G | 0.00073247 | 6 | 24.3678355 | 4.47231884 | 5.08E-08 | 5.63E-08 |
| rs75060076 | 3 | 25235889 | IMPUTED | RARB | C | T | 0.00045698 | 4 | 31.3298548 | 5.80464282 | 6.76E-08 | 7.47E-08 |
| rs77473871 | 3 | 25279859 | IMPUTED | RARB | T | C | 0.00049465 | 4 | 29.6057311 | 5.54769463 | 9.47E-08 | 1.04E-07 |
| rs78188599 | 8 | 9141514 | GENOTYPED | LOC101929128(dist=81148),LOC157273(dist=41047) | G | A | 0.02427098 | 204 | 4.00600023 | 0.75112975 | 9.64E-08 | 1.06E-07 |
| rs774879104 | 4 | 174458371 | IMPUTED | HAND2-AS1 | G | A | 0.00013549 | 1 | -51.744176 | 9.7046374 | 9.72E-08 | 1.07E-07 |
| rs762794965 | 17 | 45930212 | IMPUTED | SP6 | G | A | 0.00011944 | 1 | -51.402889 | 9.65660328 | 1.02E-07 | 1.12E-07 |
| rs180699920 | 17 | 47263875 | IMPUTED | B4GALNT2(dist=16524),GNGT2(dist=19721) | G | A | 0.00013632 | 1 | -51.184561 | 9.6504767 | 1.13E-07 | 1.24E-07 |
| rs76789477 | 3 | 25378703 | IMPUTED | RARB | C | G | 0.00050297 | 4 | 29.0947362 | 5.5101991 | 1.29E-07 | 1.41E-07 |
| rs138691552 | 17 | 46185570 | IMPUTED | SNX11 | C | T | 0.00013347 | 1 | -50.937448 | 9.65355344 | 1.32E-07 | 1.44E-07 |
| rs539056950 | 17 | 46094904 | IMPUTED | CDK5RAP3(dist=35752),COPZ2(dist=8629) | C | G | 0.0001362 | 1 | -50.727797 | 9.6545549 | 1.49E-07 | 1.62E-07 |
| rs563992891 | 6 | 40903799 | IMPUTED | LOC101929555 | C | T | 0.00099643 | 8 | -20.369024 | 3.88431519 | 1.57E-07 | 1.72E-07 |
| rs140387800 | 3 | 26853308 | IMPUTED | LRRC3B(dist=101043),NEK10(dist=299086) | G | C | 0.00027419 | 2 | 40.4605301 | 7.76113119 | 1.86E-07 | 2.02E-07 |
| rs61764931 | 1 | 4986475 | IMPUTED | AJAP1(dist=142624),MIR4417(dist=637656) | A | G | 0.01469907 | 124 | -5.2862152 | 1.01461048 | 1.89E-07 | 2.06E-07 |
| rs146691363 | 2 | 160767097 | IMPUTED | LY75-CD302(dist=5830),PLA2R1(dist=30163) | G | C | 0.00072463 | 6 | -24.737095 | 4.79107714 | 2.43E-07 | 2.64E-07 |
| rs183648689 | 5 | 3316585 | IMPUTED | LINC01377(dist=135239),LINC01019(dist=100681) | T | C | 0.00027026 | 2 | -35.417642 | 6.90273334 | 2.88E-07 | 3.13E-07 |
| rs6754826 | 2 | 174325120 | IMPUTED | CDCA7(dist=91402),SP3(dist=446067) | T | C | 0.99945436 | 5 | -26.562719 | 5.19708931 | 3.20E-07 | 3.47E-07 |
| rs77189115 | 8 | 9143416 | IMPUTED | LOC101929128(dist=83050),LOC157273(dist=39145) | C | T | 0.02435346 | 205 | 3.83010569 | 0.75151469 | 3.46E-07 | 3.74E-07 |
| rs111681509 | 8 | 71131078 | IMPUTED | NCOA2 | A | G | 0.00025386 | 2 | -36.110299 | 7.08705225 | 3.48E-07 | 3.77E-07 |
| rs16936886 | 8 | 71207356 | IMPUTED | NCOA2 | C | T | 0.00023461 | 2 | -36.888662 | 7.24484104 | 3.55E-07 | 3.84E-07 |
| rs116193953 | 1 | 163629475 | IMPUTED | LOC100422212(dist=236494),PBX1(dist=899122) | A | C | 0.00055764 | 5 | -26.868162 | 5.2831779 | 3.66E-07 | 3.96E-07 |
| rs719408 | 8 | 9151138 | IMPUTED | LOC101929128(dist=90772),LOC157273(dist=31423) | G | T | 0.02462111 | 207 | 3.79127918 | 0.74610251 | 3.75E-07 | 4.05E-07 |
| rs111975479 | 8 | 9148463 | IMPUTED | LOC101929128(dist=88097),LOC157273(dist=34098) | G | C | 0.02459698 | 207 | 3.77954041 | 0.74579018 | 4.02E-07 | 4.35E-07 |
| rs111795201 | 8 | 9147367 | IMPUTED | LOC101929128(dist=87001),LOC157273(dist=35194) | G | A | 0.02460007 | 207 | 3.77811397 | 0.74571584 | 4.05E-07 | 4.38E-07 |
| rs75170749 | 8 | 9149731 | IMPUTED | LOC101929128(dist=89365),LOC157273(dist=32830) | C | T | 0.02460304 | 207 | 3.77660622 | 0.74563978 | 4.09E-07 | 4.41E-07 |
| rs76745642 | 8 | 9151712 | IMPUTED | LOC101929128(dist=91346),LOC157273(dist=30849) | C | T | 0.02458272 | 207 | 3.78643695 | 0.74783197 | 4.12E-07 | 4.45E-07 |
| rs986373288 | 3 | 87267145 | IMPUTED | LINC00506(dist=60926),MIR4795(dist=8194) | G | C | 0.00022522 | 2 | -39.613333 | 7.83925381 | 4.34E-07 | 4.69E-07 |
| rs527468128 | 2 | 153245962 | IMPUTED | FMNL2 | A | G | 0.00033682 | 3 | 32.9365057 | 6.52492189 | 4.47E-07 | 4.82E-07 |
| rs75702282 | 8 | 9150378 | GENOTYPED | LOC101929128(dist=90012),LOC157273(dist=32183) | T | C | 0.02483955 | 209 | 3.72933798 | 0.74187587 | 4.98E-07 | 5.37E-07 |
| rs77444611 | 8 | 9150131 | GENOTYPED | LOC101929128(dist=89765),LOC157273(dist=32430) | C | T | 0.0248392 | 209 | 3.72930901 | 0.74188776 | 4.99E-07 | 5.37E-07 |

| | | | | | | | | | | | | |
|---|---|---|---|---|---|---|---|---|---|---|---|---|
| rs180993022 | 8 | 106167096 | IMPUTED | LRP12(dist=565844),ZFPM2(dist=164051) | A | C | 0.00053459 | 4 | 26.4478606 | 5.28555538 | 5.62E-07 | 6.05E-07 |
| rs11987438 | 8 | 70996125 | IMPUTED | PRDM14(dist=12563),NCOA2(dist=25872) | C | T | 0.00019634 | 2 | -44.824805 | 8.96638793 | 5.76E-07 | 6.19E-07 |
| rs9694166 | 8 | 70993882 | IMPUTED | PRDM14(dist=10320),NCOA2(dist=28115) | T | A | 0.00021512 | 2 | -44.081107 | 8.81846138 | 5.77E-07 | 6.21E-07 |
| rs577475538 | 13 | 29950912 | IMPUTED | MTUS2 | G | C | 0.0005101 | 4 | -26.56502 | 5.31999679 | 5.93E-07 | 6.38E-07 |
| rs370514761 | 16 | 81922041 | IMPUTED | PLCG2 | A | G | 0.00077953 | 7 | 23.0548297 | 4.64207346 | 6.82E-07 | 7.32E-07 |
| rs187064111 | 19 | 1072247 | IMPUTED | ARHGAP45 | C | T | 0.00277502 | 23 | 11.5323771 | 2.32560421 | 7.09E-07 | 7.61E-07 |
| rs77977386 | 2 | 151550092 | IMPUTED | LOC101929282(dist=58221),RBM43(dist=554636) | G | A | 0.00046637 | 4 | -27.152343 | 5.4779313 | 7.17E-07 | 7.69E-07 |
| rs74397915 | 2 | 151554303 | IMPUTED | LOC101929282(dist=62432),RBM43(dist=550425) | A | G | 0.00046637 | 4 | -27.152343 | 5.4779313 | 7.17E-07 | 7.69E-07 |
| rs79907700 | 2 | 151544901 | GENOTYPED | LOC101929282(dist=53030),RBM43(dist=559827) | C | T | 0.00046541 | 4 | -27.19154 | 5.48590713 | 7.17E-07 | 7.70E-07 |
| rs74761575 | 2 | 151541269 | IMPUTED | LOC101929282(dist=49398),RBM43(dist=563459) | A | T | 0.00046589 | 4 | -27.180506 | 5.48644757 | 7.27E-07 | 7.79E-07 |
| rs113378776 | 2 | 151540798 | IMPUTED | LOC101929282(dist=48927),RBM43(dist=563930) | T | C | 0.00046007 | 4 | -27.522878 | 5.56290838 | 7.51E-07 | 8.06E-07 |
| rs147648162 | 8 | 70998109 | IMPUTED | PRDM14(dist=14547),NCOA2(dist=23888) | A | C | 0.00017946 | 2 | -44.651135 | 9.04366207 | 7.92E-07 | 8.49E-07 |
| rs1047911201 | 6 | 94594504 | IMPUTED | TSG1(dist=108305),MANEA-AS1(dist=1413468) | C | A | 0.00059246 | 5 | 23.8911756 | 4.85190593 | 8.48E-07 | 9.08E-07 |
| rs553961701 | 6 | 93833544 | IMPUTED | CASC6(dist=1433398),EPHA7(dist=116196) | C | T | 0.00060233 | 5 | 23.788558 | 4.8365617 | 8.72E-07 | 9.34E-07 |
| rs112905701 | 2 | 151546989 | IMPUTED | LOC101929282(dist=55118),RBM43(dist=557739) | C | T | 0.00044046 | 4 | -28.488153 | 5.81214832 | 9.51E-07 | 1.02E-06 |
| rs75464434 | 2 | 151548853 | IMPUTED | LOC101929282(dist=56982),RBM43(dist=555875) | C | A | 0.00043986 | 4 | -28.51266 | 5.81967739 | 9.62E-07 | 1.03E-06 |
| rs575578613 | 9 | 2759716 | IMPUTED | KCNV2(dist=29679),PUM3(dist=44439) | A | G | 0.00060019 | 5 | -23.2725 | 4.75135263 | 9.68E-07 | 1.03E-06 |
| rs1017740900 | 6 | 92831492 | IMPUTED | CASC6(dist=431346),EPHA7(dist=1118248) | A | G | 0.00071892 | 6 | 21.9082834 | 4.48094567 | 1.01E-06 | 1.08E-06 |
| rs114219304 | 2 | 151583525 | IMPUTED | LOC101929282(dist=91654),RBM43(dist=521203) | G | A | 0.00043725 | 4 | -28.569688 | 5.84664157 | 1.03E-06 | 1.10E-06 |
| rs142516872 | 14 | 55235501 | IMPUTED | SAMD4A | G | A | 0.00024899 | 2 | -39.3711 | 8.05939603 | 1.03E-06 | 1.10E-06 |
| rs577268629 | 5 | 150322147 | IMPUTED | ZNF300P1 | A | T | 0.00053625 | 5 | 27.0875166 | 5.54990228 | 1.06E-06 | 1.13E-06 |
| rs1024983013 | 6 | 92781358 | IMPUTED | CASC6(dist=381212),EPHA7(dist=1168382) | A | G | 0.00070692 | 6 | 22.2060134 | 4.55797119 | 1.11E-06 | 1.18E-06 |
| rs181989792 | 15 | 61401267 | IMPUTED | RORA | C | T | 0.00126753 | 11 | 16.0671759 | 3.29857012 | 1.11E-06 | 1.19E-06 |
| rs35899516 | 13 | 107450854 | IMPUTED | LINC00443(dist=126326),FAM155A(dist=370025) | T | G | 0.00054528 | 5 | 24.5718495 | 5.05294957 | 1.16E-06 | 1.23E-06 |
| rs554910598 | 20 | 25474435 | IMPUTED | NINL | G | A | 0.00034894 | 3 | -29.493755 | 6.06725523 | 1.17E-06 | 1.25E-06 |
| rs11985947 | 8 | 70999631 | IMPUTED | PRDM14(dist=16069),NCOA2(dist=22366) | C | T | 0.00020311 | 2 | -43.29721 | 8.91089386 | 1.18E-06 | 1.26E-06 |
| rs139275277 | 20 | 13137998 | IMPUTED | SPTLC3 | C | T | 0.00038686 | 3 | 32.1435352 | 6.621196 | 1.21E-06 | 1.29E-06 |
| rs185050080 | 15 | 61400356 | IMPUTED | RORA | G | A | 0.0012497 | 11 | 16.0054341 | 3.30240284 | 1.26E-06 | 1.34E-06 |
| rs74596618 | 2 | 151544939 | IMPUTED | LOC101929282(dist=53068),RBM43(dist=559789) | C | T | 0.00043404 | 4 | -28.733013 | 5.93049864 | 1.27E-06 | 1.35E-06 |
| rs73225625 | 12 | 95598185 | IMPUTED | FGD6 | C | T | 0.00124994 | 11 | 17.2613957 | 3.56316371 | 1.27E-06 | 1.35E-06 |
| rs902845237 | 6 | 93232345 | IMPUTED | CASC6(dist=832199),EPHA7(dist=717395) | G | T | 0.0007251 | 6 | 21.2311609 | 4.3828145 | 1.27E-06 | 1.36E-06 |
| rs145878596 | 8 | 71000054 | IMPUTED | PRDM14(dist=16492),NCOA2(dist=21943) | A | T | 0.00020276 | 2 | -43.094492 | 8.90414385 | 1.30E-06 | 1.39E-06 |
| rs184927009 | 17 | 47245198 | IMPUTED | B4GALNT2 | C | G | 0.00276836 | 23 | -11.32581 | 2.34151225 | 1.32E-06 | 1.41E-06 |
| rs140760764 | 2 | 103757580 | IMPUTED | LINC01935(dist=156693),LOC100287010(dist=1237728) | C | G | 0.00038377 | 3 | 32.9999815 | 6.83161556 | 1.36E-06 | 1.45E-06 |
| rs138495502 | 8 | 71000055 | IMPUTED | PRDM14(dist=16493),NCOA2(dist=21942) | C | T | 0.00020097 | 2 | -42.935844 | 8.89249178 | 1.38E-06 | 1.47E-06 |
| rs931770839 | 6 | 92645910 | IMPUTED | CASC6(dist=245764),EPHA7(dist=1303830) | A | G | 0.0007421 | 6 | 21.1275954 | 4.38192484 | 1.42E-06 | 1.52E-06 |
| rs534689729 | 1 | 108007520 | IMPUTED | NTNG1 | G | A | 0.00078928 | 7 | 22.3404908 | 4.6371435 | 1.45E-06 | 1.55E-06 |
| rs181742183 | 15 | 85216399 | IMPUTED | SEC11A | G | A | 0.00039315 | 3 | -28.234418 | 5.86289485 | 1.47E-06 | 1.56E-06 |
| rs190154072 | 4 | 117381972 | IMPUTED | MIR1973(dist=161048),TRAM1L1(dist=622738) | A | G | 0.00022284 | 2 | -37.16641 | 7.73492394 | 1.55E-06 | 1.65E-06 |
| rs531471021 | 18 | 29650959 | IMPUTED | RNF125(NM_017831:c.*26120>0) | G | A | 0.00048158 | 4 | -27.133314 | 5.64714397 | 1.55E-06 | 1.65E-06 |
| rs551686288 | 1 | 196714448 | IMPUTED | CFH | A | G | 0.00153173 | 13 | 15.6035053 | 3.25093567 | 1.59E-06 | 1.69E-06 |
| rs113543017 | 2 | 151564112 | IMPUTED | LOC101929282(dist=72241),RBM43(dist=540616) | G | A | 0.00045496 | 4 | -27.935046 | 5.82094211 | 1.59E-06 | 1.70E-06 |
| rs117717189 | 2 | 39154242 | IMPUTED | ARHGEF33 | T | C | 0.00073199 | 6 | 20.9787855 | 4.37374713 | 1.61E-06 | 1.72E-06 |
| rs74585063 | 2 | 39151832 | IMPUTED | ARHGEF33 | G | C | 0.00072926 | 6 | 20.9706842 | 4.37566268 | 1.65E-06 | 1.75E-06 |
| rs759739760 | 3 | 103915969 | IMPUTED | MIR548A3 | A | T | 2.25E-05 | | -685.65837 | 143.148329 | 1.67E-06 | |
| rs758322952 | 1 | 113233395 | IMPUTED | MOV10 | C | T | 0.00026836 | 2 | -34.352545 | 7.18012268 | 1.72E-06 | 1.82E-06 |
| rs1005986274 | 5 | 39226382 | IMPUTED | FYB | G | A | 0.00020751 | 2 | -39.693796 | 8.30105582 | 1.74E-06 | 1.85E-06 |
| rs1050739172 | 2 | 156651436 | IMPUTED | KCNJ3(dist=936572),LINC01876(dist=225611) | A | G | 0.00165023 | 14 | 15.4623182 | 3.23486936 | 1.75E-06 | 1.86E-06 |
| rs74967419 | 13 | 33583387 | GENOTYPED | LINC00423(dist=97597),KL(dist=7184) | C | A | 0.0002377 | 2 | 33.2186174 | 6.95235901 | 1.77E-06 | 1.88E-06 |
| rs782769776 | 1 | 147130330 | IMPUTED | ACP6,NBPF19 | A | G | 0.00048657 | 4 | -26.688628 | 5.58905266 | 1.80E-06 | 1.91E-06 |
| rs79803408 | 2 | 151539121 | IMPUTED | LOC101929282(dist=47250),RBM43(dist=565607) | C | A | 0.00041633 | 4 | -29.126411 | 6.10095708 | 1.81E-06 | 1.92E-06 |
| rs778464311 | 6 | 95898881 | IMPUTED | TSG1(dist=1412682),MANEA-AS1(dist=109091) | C | A | 0.00076159 | 6 | 21.041015 | 4.40894513 | 1.82E-06 | 1.93E-06 |

| | | | | | | | | | | | | |
|---|---|---|---|---|---|---|---|---|---|---|---|---|
| rs150294795 | 2 | 39156337 | IMPUTED | ARHGEF33 | C | T | 0.00071952 | 6 | 20.8966453 | 4.3820524 | 1.85E-06 | 1.97E-06 |
| rs115845689 | 2 | 39155706 | IMPUTED | ARHGEF33 | G | A | 0.0007194 | 6 | 20.8942157 | 4.38205584 | 1.86E-06 | 1.97E-06 |
| rs114962859 | 2 | 39177653 | IMPUTED | ARHGEF33 | C | T | 0.00075042 | 6 | 21.0172455 | 4.4085856 | 1.87E-06 | 1.98E-06 |
| rs554008645 | 6 | 42346928 | IMPUTED | TRERF1 | C | T | 0.00074483 | 6 | 21.8212544 | 4.57746296 | 1.87E-06 | 1.98E-06 |
| rs538595234 | 14 | 101390810 | IMPUTED | MEG8 | A | C | 0.00121702 | 10 | 17.1012638 | 3.58772955 | 1.87E-06 | 1.99E-06 |
| rs544558253 | 20 | 13123525 | IMPUTED | SPTLC3 | G | A | 0.00037628 | 3 | 31.5929752 | 6.62997261 | 1.89E-06 | 2.00E-06 |
| rs543850521 | 19 | 495019 | IMPUTED | ODF3L2(dist=20036),MADCAM1(dist=1471) | G | A | 0.00035393 | 3 | 33.3186897 | 7.00138043 | 1.95E-06 | 2.07E-06 |
| rs9301165 | 13 | 107437444 | IMPUTED | LINC00443(dist=112916),FAM155A(dist=383435) | T | C | 0.00049762 | 4 | 25.1098779 | 5.27860242 | 1.97E-06 | 2.09E-06 |
| rs78744226 | 2 | 39163630 | IMPUTED | ARHGEF33 | A | G | 0.00071464 | 6 | 20.848015 | 4.38300805 | 1.97E-06 | 2.09E-06 |
| rs144258224 | 2 | 39160588 | IMPUTED | ARHGEF33 | G | T | 0.00071322 | 6 | 20.829206 | 4.38313012 | 2.01E-06 | 2.14E-06 |
| rs79315504 | 2 | 39149289 | IMPUTED | ARHGEF33 | T | C | 0.0007131 | 6 | 20.8269254 | 4.38313015 | 2.02E-06 | 2.14E-06 |
| rs79229305 | 2 | 39149783 | GENOTYPED | ARHGEF33 | C | T | 0.0007131 | 6 | 20.8269254 | 4.38313015 | 2.02E-06 | 2.14E-06 |
| rs79141371 | 2 | 39149911 | IMPUTED | ARHGEF33 | A | C | 0.0007131 | 6 | 20.8269254 | 4.38313015 | 2.02E-06 | 2.14E-06 |
| rs75939897 | 2 | 39161250 | IMPUTED | ARHGEF33 | A | T | 0.0007131 | 6 | 20.8269254 | 4.38313015 | 2.02E-06 | 2.14E-06 |
| rs76593331 | 2 | 39163177 | GENOTYPED | ARHGEF33 | G | A | 0.0007131 | 6 | 20.8269254 | 4.38313015 | 2.02E-06 | 2.14E-06 |
| rs80270650 | 2 | 39163519 | IMPUTED | ARHGEF33 | C | T | 0.0007131 | 6 | 20.8269254 | 4.38313015 | 2.02E-06 | 2.14E-06 |
| rs115901879 | 2 | 39171796 | GENOTYPED | ARHGEF33 | C | G | 0.0007131 | 6 | 20.8269254 | 4.38313015 | 2.02E-06 | 2.14E-06 |
| rs114990055 | 2 | 39145286 | IMPUTED | MORN2(dist=35436),ARHGEF33(dist=1218) | T | C | 0.00071488 | 6 | 20.82536 | 4.38313945 | 2.02E-06 | 2.15E-06 |
| rs79778788 | 2 | 39145206 | IMPUTED | MORN2(dist=35356),ARHGEF33(dist=1298) | G | A | 0.00071357 | 6 | 20.8248578 | 4.38312822 | 2.02E-06 | 2.15E-06 |
| rs78786041 | 2 | 160469060 | IMPUTED | BAZ2B | C | T | 0.00046886 | 4 | -29.67656 | 6.24760335 | 2.03E-06 | 2.16E-06 |
| rs59494028 | 13 | 107436325 | IMPUTED | LINC00443(dist=111797),FAM155A(dist=384554) | C | T | 0.0004773 | 4 | 25.1084302 | 5.28737022 | 2.05E-06 | 2.17E-06 |
| rs144763189 | 2 | 103366466 | IMPUTED | TMEM182 | A | G | 0.00050452 | 4 | 26.497962 | 5.58195964 | 2.06E-06 | 2.19E-06 |
| rs80103983 | 20 | 34350913 | IMPUTED | RBM39(dist=20655),PHF20(dist=9010) | C | T | 0.00025981 | 2 | 34.5238427 | 7.27745245 | 2.10E-06 | 2.22E-06 |
| rs4972570 | 2 | 174325744 | IMPUTED | CDCA7(dist=92026),SP3(dist=445443) | A | T | 0.99952341 | 4 | -25.241139 | 5.32258807 | 2.11E-06 | 2.24E-06 |
| rs939168442 | 18 | 50317044 | IMPUTED | DCC | T | G | 0.00024471 | 2 | -36.515083 | 7.70013065 | 2.11E-06 | 2.24E-06 |
| rs142156809 | 21 | 31517942 | IMPUTED | GRIK1(dist=205572),CLDN17(dist=20299) | A | C | 0.00039018 | 3 | 31.0679021 | 6.55191734 | 2.12E-06 | 2.25E-06 |
| rs188948126 | 20 | 25316590 | IMPUTED | ABHD12 | T | C | 0.00049334 | 4 | -24.288082 | 5.1221843 | 2.12E-06 | 2.25E-06 |
| rs141219189 | 2 | 103429638 | IMPUTED | TMEM182 | T | G | 0.00049418 | 4 | 26.5048204 | 5.59092471 | 2.13E-06 | 2.26E-06 |
| rs76432074 | 2 | 151565341 | IMPUTED | LOC101929282(dist=73470),RBM43(dist=539387) | A | G | 0.00044925 | 4 | -28.118598 | 5.93466744 | 2.16E-06 | 2.29E-06 |
| rs145423064 | 15 | 64643725 | IMPUTED | CSNK1G1 | A | G | 0.00834407 | 70 | -6.4138713 | 1.35405004 | 2.17E-06 | 2.30E-06 |
| rs74649461 | 2 | 151564122 | IMPUTED | LOC101929282(dist=72251),RBM43(dist=540606) | G | C | 0.00044937 | 4 | -28.110286 | 5.93464453 | 2.17E-06 | 2.30E-06 |
| rs1405229 | 2 | 174324895 | IMPUTED | CDCA7(dist=91177),SP3(dist=446292) | A | G | 0.99951854 | 4 | -25.197357 | 5.32060439 | 2.18E-06 | 2.31E-06 |
| rs181020921 | 20 | 25373429 | IMPUTED | ABHD12(dist=1811),GINS1(dist=14890) | A | G | 0.00049394 | 4 | -24.200657 | 5.11232812 | 2.20E-06 | 2.34E-06 |
| rs76625767 | 2 | 103393185 | IMPUTED | TMEM182 | G | A | 0.0004975 | 4 | 26.4319382 | 5.58662715 | 2.23E-06 | 2.37E-06 |
| rs4972402 | 2 | 174327221 | IMPUTED | CDCA7(dist=93503),SP3(dist=443966) | A | G | 0.99950844 | 4 | -25.180547 | 5.32252896 | 2.23E-06 | 2.37E-06 |
| rs537853682 | 2 | 25263621 | IMPUTED | DNAJC27-AS1(dist=1058),EFR3B(dist=1352) | C | T | 0.00010019 | 1 | -55.610322 | 11.7605498 | 2.26E-06 | 2.40E-06 |
| rs59733052 | 11 | 20779968 | IMPUTED | NELL1 | C | T | 0.00012966 | 1 | -47.202377 | 9.98547428 | 2.28E-06 | 2.41E-06 |
| rs6044663 | 20 | 17158967 | GENOTYPED | OTOR(dist=426158),PCSK2(dist=47785) | G | T | 0.00038377 | 3 | -25.229028 | 5.33716853 | 2.28E-06 | 2.41E-06 |
| rs13282027 | 8 | 25004229 | IMPUTED | NEFL(dist=189846),DOCK5(dist=38009) | G | A | 0.33414595 | 2812 | -1.1617147 | 0.24588367 | 2.31E-06 | 2.44E-06 |
| rs536311319 | 2 | 30897792 | IMPUTED | LCLAT1(dist=30701),CAPN13(dist=47846) | A | T | 0.00308783 | 26 | 10.91449 | 2.31037676 | 2.31E-06 | 2.45E-06 |
| rs1340931 | 20 | 17157920 | IMPUTED | OTOR(dist=425111),PCSK2(dist=48832) | G | A | 0.00038258 | 3 | -25.293914 | 5.35550493 | 2.32E-06 | 2.46E-06 |
| rs13250561 | 8 | 24991003 | GENOTYPED | NEFL(dist=176620),DOCK5(dist=51235) | T | C | 0.33245353 | 2797 | -1.163967 | 0.24644884 | 2.32E-06 | 2.46E-06 |
| rs75946017 | 18 | 50376400 | IMPUTED | DCC | T | G | 0.01601569 | 135 | -4.3721691 | 0.92674808 | 2.38E-06 | 2.53E-06 |
| rs185185799 | 9 | 74187011 | IMPUTED | TRPM3(dist=450497),TMEM2(dist=111271) | A | T | 0.0018922 | 16 | 13.2686983 | 2.81372942 | 2.41E-06 | 2.55E-06 |
| rs529459930 | 6 | 151843034 | IMPUTED | CCDC170 | C | T | 0.00044367 | 4 | 30.6953198 | 6.51167496 | 2.43E-06 | 2.57E-06 |
| rs535373912 | 20 | 29991638 | IMPUTED | DEFB119(dist=13186),DEFB121(dist=1010) | G | A | 0.00047255 | 4 | -24.347779 | 5.16562823 | 2.44E-06 | 2.58E-06 |
| rs117326470 | 2 | 103722596 | IMPUTED | LINC01935(dist=121709),LOC100287010(dist=1272712) | A | G | 0.00047195 | 4 | 27.0142965 | 5.73389467 | 2.46E-06 | 2.61E-06 |
| rs115962163 | 3 | 25402205 | IMPUTED | RARB | C | T | 0.00067423 | 6 | 22.3518497 | 4.74521864 | 2.47E-06 | 2.62E-06 |
| rs6034762 | 20 | 17160348 | IMPUTED | OTOR(dist=427539),PCSK2(dist=46404) | G | A | 0.00037901 | 3 | -25.485243 | 5.41073729 | 2.48E-06 | 2.62E-06 |
| rs529668978 | 20 | 29526419 | IMPUTED | LINC01597(dist=5206),LINC01598(dist=32069) | G | T | 0.00047064 | 4 | -24.306621 | 5.1605934 | 2.48E-06 | 2.62E-06 |
| rs114711347 | 20 | 17160919 | IMPUTED | OTOR(dist=428110),PCSK2(dist=45833) | C | A | 0.00037758 | 3 | -25.560277 | 5.43291746 | 2.54E-06 | 2.69E-06 |
| rs10415006 | 19 | 439393 | IMPUTED | SHC2 | C | T | 0.00051723 | 4 | 25.9550729 | 5.51744164 | 2.55E-06 | 2.70E-06 |

| | | | | | | | | | | | | |
|---|---|---|---|---|---|---|---|---|---|---|---|---|
| rs760275844 | 11 | 20797858 | IMPUTED | NELL1 | G | A | 0.00013501 | 1 | -47.052442 | 10.0033212 | 2.56E-06 | 2.70E-06 |
| rs530523771 | 11 | 106342138 | IMPUTED | LOC101928535(dist=206506),GUCY1A2(dist=202600) | G | T | 0.00387675 | 33 | 8.88135766 | 1.8882546 | 2.56E-06 | 2.71E-06 |
| rs564522958 | 3 | 155871086 | IMPUTED | KCNAB1 | G | C | 0.00011255 | 1 | -49.208301 | 10.4639548 | 2.57E-06 | 2.72E-06 |
| rs542332660 | 13 | 30487023 | IMPUTED | LINC00297(dist=24481),LINC00572(dist=5761) | G | T | 0.00053993 | 5 | 25.3777224 | 5.39904559 | 2.60E-06 | 2.75E-06 |
| rs80134155 | 20 | 17161694 | IMPUTED | OTOR(dist=428885),PCSK2(dist=45058) | A | G | 0.00037628 | 3 | -25.628253 | 5.45328976 | 2.61E-06 | 2.76E-06 |
| rs141622093 | 6 | 83194457 | IMPUTED | TPBG(dist=117324),UBE3D(dist=407660) | G | A | 0.00062016 | 5 | 22.2966133 | 4.74557131 | 2.62E-06 | 2.78E-06 |
| rs532779237 | 12 | 63527786 | IMPUTED | PPM1H(dist=199121),AVPR1A(dist=8753) | T | C | 0.00039696 | 3 | 30.8113726 | 6.5622559 | 2.66E-06 | 2.82E-06 |
| rs541726732 | 20 | 29962442 | IMPUTED | DEFB118(dist=737) | G | T | 0.00048835 | 4 | -24.057433 | 5.12414834 | 2.67E-06 | 2.82E-06 |
| rs113368668 | 11 | 67815327 | IMPUTED | TCIRG1 | G | A | 0.00011612 | 1 | -50.414049 | 10.7386615 | 2.67E-06 | 2.83E-06 |
| rs544725047 | 2 | 96828129 | IMPUTED | DUSP2(dist=16950),STARD7(dist=22474) | C | T | 0.00138864 | 12 | 15.7627588 | 3.35866995 | 2.69E-06 | 2.85E-06 |
| rs772818834 | 3 | 47423512 | IMPUTED | PTPN23 | G | T | 0.00012741 | 1 | -47.186891 | 10.0544282 | 2.69E-06 | 2.85E-06 |
| rs563560528 | 16 | 85361133 | IMPUTED | MIR5093(dist=21202),GSE1(dist=283896) | C | A | 0.00012467 | 1 | -46.267332 | 9.86236967 | 2.71E-06 | 2.87E-06 |
| rs188505066 | 16 | 60432138 | IMPUTED | LOC729159(dist=38441),MIR4426(dist=657473) | C | G | 0.0003909 | 3 | -28.81086 | 6.14165936 | 2.72E-06 | 2.88E-06 |
| rs6034764 | 20 | 17163074 | IMPUTED | OTOR(dist=430265),PCSK2(dist=43678) | A | G | 0.00037652 | 3 | -25.762519 | 5.49407331 | 2.74E-06 | 2.90E-06 |
| rs184187225 | 2 | 191383876 | IMPUTED | NEMP2 | C | T | 0.00013763 | 1 | -45.796831 | 9.77118242 | 2.77E-06 | 2.93E-06 |
| rs144757912 | 11 | 20292346 | IMPUTED | DBX1(dist=110476),HTATIP2(dist=92885) | C | T | 0.00012871 | 1 | -45.769697 | 9.76577359 | 2.78E-06 | 2.94E-06 |
| rs779410455 | 8 | 83381356 | IMPUTED | SNX16(dist=626835),LOC101927141(dist=442983) | G | A | 0.0001198 | 1 | -47.220118 | 10.0755872 | 2.78E-06 | 2.94E-06 |
| rs146709819 | 3 | 164358149 | IMPUTED | MIR1263(dist=468805),LINC01324(dist=73734) | G | A | 0.0001242 | 1 | -48.482216 | 10.347856 | 2.80E-06 | 2.96E-06 |
| rs567855665 | 9 | 86225665 | IMPUTED | FRMD3(dist=72317),IDNK(dist=12299) | G | A | 0.00063977 | 5 | -24.277589 | 5.18195859 | 2.80E-06 | 2.96E-06 |
| rs562565330 | 12 | 127167561 | IMPUTED | LOC100996671 | C | G | 0.00129914 | 11 | 15.2547791 | 3.25609066 | 2.80E-06 | 2.96E-06 |
| rs535347456 | 8 | 32294257 | IMPUTED | NRG1 | G | C | 0.00092786 | 8 | 20.2222393 | 4.31640018 | 2.80E-06 | 2.96E-06 |
| rs530481997 | 8 | 82900022 | IMPUTED | SNX16(dist=145501),LOC101927141(dist=924317) | C | T | 0.00012277 | 1 | -45.824575 | 9.78213407 | 2.81E-06 | 2.97E-06 |
| rs949980187 | 17 | 73475525 | IMPUTED | TMEM94 | C | T | 0.00011707 | 1 | -49.174008 | 10.4974341 | 2.81E-06 | 2.97E-06 |
| rs549491075 | 2 | 52691266 | IMPUTED | LOC730100(dist=56211),MIR4431(dist=238394) | G | A | 0.00036713 | 3 | 30.9790432 | 6.61342196 | 2.81E-06 | 2.97E-06 |
| rs1034233648 | 1 | 231611902 | IMPUTED | SNRPD2P2 | G | C | 0.00010958 | 1 | -50.741165 | 10.8342789 | 2.82E-06 | 2.98E-06 |
| rs527764081 | 2 | 52691238 | IMPUTED | LOC730100(dist=56183),MIR4431(dist=238422) | G | A | 0.00036701 | 3 | 30.9730137 | 6.6137007 | 2.82E-06 | 2.99E-06 |
| rs773229669 | 2 | 180006844 | IMPUTED | SESTD1 | A | T | 0.00012337 | 1 | -45.76558 | 9.77302774 | 2.83E-06 | 2.99E-06 |
| rs188491002 | 8 | 82935147 | IMPUTED | SNX16(dist=180626),LOC101927141(dist=889192) | G | A | 0.00012039 | 1 | -45.813088 | 9.78392049 | 2.83E-06 | 3.00E-06 |
| rs149603239 | 1 | 104863243 | IMPUTED | LOC100129138(dist=243550),LINC01676(dist=1269073) | G | A | 0.00036582 | 3 | 31.4144817 | 6.71083131 | 2.85E-06 | 3.02E-06 |
| rs372998427 | 19 | 9006337 | IMPUTED | MUC16 | G | A | 0.00011647 | 1 | -48.98661 | 10.465608 | 2.86E-06 | 3.02E-06 |
| rs1918085 | 2 | 52700429 | IMPUTED | LOC730100(dist=65374),MIR4431(dist=229231) | A | G | 0.00036808 | 3 | 30.9483906 | 6.61443296 | 2.88E-06 | 3.05E-06 |
| rs191005731 | 8 | 124557018 | IMPUTED | FBXO32(dist=3525),KLHL38(dist=100897) | T | C | 0.00012123 | 1 | -45.768477 | 9.78389571 | 2.90E-06 | 3.06E-06 |
| rs115309670 | 2 | 32981307 | IMPUTED | TTC27 | C | T | 0.00010494 | 1 | -51.940089 | 11.107166 | 2.92E-06 | 3.09E-06 |
| rs111405156 | 9 | 5628556 | IMPUTED | RIC1(dist=563) | T | C | 0.0091368 | 77 | 5.70222231 | 1.21959114 | 2.93E-06 | 3.10E-06 |
| rs556795906 | 3 | 155835138 | IMPUTED | GMPS(dist=179618),KCNAB1(dist=3199) | G | A | 0.00011136 | 1 | -48.928051 | 10.4648854 | 2.93E-06 | 3.10E-06 |
| rs529238197 | 16 | 87301319 | IMPUTED | LOC101928708(dist=41284),LOC101928682(dist=4193) | G | A | 0.00011873 | 1 | -45.768382 | 9.79386445 | 2.97E-06 | 3.14E-06 |
| rs776691922 | 2 | 169243036 | IMPUTED | STK39(dist=138931),CERS6(dist=69723) | A | G | 0.00221452 | 19 | 12.861402 | 2.75268145 | 2.98E-06 | 3.15E-06 |
| rs60396700 | 2 | 32980913 | IMPUTED | TTC27 | A | T | 0.00010435 | 1 | -52.426738 | 11.2219084 | 2.99E-06 | 3.16E-06 |
| rs114018484 | 20 | 17165073 | IMPUTED | OTOR(dist=432264),PCSK2(dist=41679) | A | G | 0.00037259 | 3 | -25.953204 | 5.55540054 | 2.99E-06 | 3.16E-06 |
| rs576061242 | 9 | 127362503 | IMPUTED | NR6A1 | T | C | 0.00012301 | 1 | -45.700569 | 9.78376027 | 3.00E-06 | 3.17E-06 |
| rs778109765 | 20 | 24866728 | IMPUTED | SYNDIG1(dist=219475),CST7(dist=63138) | G | C | 0.00011837 | 1 | -47.014032 | 10.0662597 | 3.01E-06 | 3.18E-06 |
| rs74799151 | 2 | 32979083 | IMPUTED | TTC27 | A | G | 0.00011208 | 1 | -48.718173 | 10.4314113 | 3.01E-06 | 3.18E-06 |
| rs186307384 | 2 | 32979871 | IMPUTED | TTC27 | C | T | 0.00010435 | 1 | -52.409777 | 11.2218592 | 3.01E-06 | 3.18E-06 |
| rs553246003 | 3 | 120171375 | IMPUTED | FSTL1(dist=1457),NDUFB4(dist=143753) | G | A | 0.00014749 | 1 | -46.199377 | 9.89931382 | 3.06E-06 | 3.23E-06 |
| rs575655339 | 1 | 67401582 | IMPUTED | MIER1 | A | G | 0.00012729 | 1 | -45.690086 | 9.79149362 | 3.07E-06 | 3.24E-06 |
| rs533019949 | 5 | 39083282 | IMPUTED | RICTOR(dist=8772),FYB(dist=22072) | G | T | 0.00021714 | 2 | -37.310902 | 7.995824 | 3.07E-06 | 3.24E-06 |
| rs567613074 | 8 | 105643811 | IMPUTED | LRP12(dist=42559),ZFPM2(dist=687336) | T | C | 0.00011528 | 1 | -49.482673 | 10.6104389 | 3.11E-06 | 3.28E-06 |
| rs59766339 | 5 | 171259114 | IMPUTED | SMIM23(dist=41022),FBXW11(dist=29442) | T | C | 0.0003594 | 3 | -29.292399 | 6.28148646 | 3.11E-06 | 3.29E-06 |
| rs546326444 | 5 | 180708080 | IMPUTED | TRIM52-AS1(dist=8772),LOC100133331(dist=42427) | T | C | 0.00041657 | 4 | 30.2571574 | 6.48857813 | 3.11E-06 | 3.29E-06 |
| rs7844172 | 8 | 105321247 | IMPUTED | RIMS2(dist=54591),DCSTAMP(dist=30777) | C | T | 0.00011398 | 1 | -50.394228 | 10.8106414 | 3.14E-06 | 3.32E-06 |
| rs80172391 | 10 | 132702249 | IMPUTED | GLRX3(dist=723603),MIR378C(dist=58602) | C | A | 0.00012301 | 1 | -45.58488 | 9.78354742 | 3.17E-06 | 3.35E-06 |
| rs73328046 | 5 | 171260585 | IMPUTED | SMIM23(dist=42493),FBXW11(dist=27971) | A | C | 0.0003714 | 3 | -28.09392 | 6.02998104 | 3.18E-06 | 3.36E-06 |

| | | | | | | | | | | | | |
|---|---|---|---|---|---|---|---|---|---|---|---|---|
| rs528890470 | 13 | 23427662 | IMPUTED | LINC00540(dist=577003),LINC00621(dist=22869) | A | C | 0.00012313 | 1 | -45.745514 | 9.8219539 | 3.20E-06 | 3.38E-06 |
| rs761025544 | 18 | 72134950 | IMPUTED | FAM69C(dist=10447),CNDP2(dist=28550) | C | T | 0.001285 | 11 | 15.8473635 | 3.40261182 | 3.20E-06 | 3.38E-06 |
| rs760213983 | 1 | 112731736 | IMPUTED | LINC01750(dist=190273),CTTNBP2NL(dist=207064) | T | C | 0.00011469 | 1 | -50.643099 | 10.8810077 | 3.25E-06 | 3.43E-06 |
| rs13183308 | 5 | 163996922 | IMPUTED | LOC102546299(dist=26933),NONE(dist=NONE) | C | G | 0.20474745 | 1723 | 1.36071379 | 0.29257783 | 3.31E-06 | 3.49E-06 |
| rs73328041 | 5 | 171260244 | IMPUTED | SMIM23(dist=42152),FBXW11(dist=28312) | A | G | 0.00036808 | 3 | -28.016478 | 6.02412177 | 3.31E-06 | 3.49E-06 |
| rs73328042 | 5 | 171260437 | IMPUTED | SMIM23(dist=42345),FBXW11(dist=28119) | A | G | 0.00036808 | 3 | -28.016478 | 6.02412177 | 3.31E-06 | 3.49E-06 |
| rs570594505 | 16 | 87663525 | IMPUTED | JPH3 | T | C | 0.00012372 | 1 | -45.439956 | 9.77171458 | 3.32E-06 | 3.50E-06 |
| rs114883353 | 17 | 30026956 | IMPUTED | MIR365B(dist=124416),COPRS(dist=151928) | G | A | 0.00040231 | 3 | 30.4706675 | 6.55505036 | 3.34E-06 | 3.53E-06 |
| rs190509573 | 1 | 104890979 | IMPUTED | LOC100129138(dist=271286),LINC01676(dist=1241337) | A | T | 0.00035857 | 3 | 31.4533234 | 6.76862225 | 3.37E-06 | 3.56E-06 |
| rs569160882 | 17 | 68788764 | IMPUTED | KCNJ2(dist=612581),CASC17(dist=305151) | C | T | 5.84E-05 | | -97.648713 | 21.0167516 | 3.38E-06 | |
| rs527262521 | 11 | 16866327 | IMPUTED | PLEKHA7 | A | G | 9.50E-05 | 1 | -63.597487 | 13.6879911 | 3.38E-06 | 3.57E-06 |
| rs141237895 | 1 | 215308848 | IMPUTED | KCNK2 | A | C | 0.00019277 | 2 | -41.286866 | 8.88692677 | 3.39E-06 | 3.58E-06 |
| rs753298408 | 18 | 62608115 | IMPUTED | LINC01924(dist=517288),CDH7(dist=809373) | G | C | 0.00013787 | 1 | -46.112412 | 9.92635313 | 3.39E-06 | 3.58E-06 |
| rs9841732 | 3 | 24698792 | IMPUTED | MIR4792(dist=135866),RARB(dist=172022) | A | G | 0.00071322 | 6 | 20.1294509 | 4.33361574 | 3.40E-06 | 3.59E-06 |
| rs9877614 | 3 | 24703565 | IMPUTED | MIR4792(dist=140639),RARB(dist=167249) | C | G | 0.00071322 | 6 | 20.1294509 | 4.33361574 | 3.40E-06 | 3.59E-06 |
| rs9812320 | 3 | 24680869 | IMPUTED | MIR4792(dist=117943),RARB(dist=189945) | G | A | 0.0007131 | 6 | 20.1239536 | 4.33263058 | 3.41E-06 | 3.59E-06 |
| rs9853786 | 3 | 24688359 | IMPUTED | MIR4792(dist=125433),RARB(dist=182455) | C | T | 0.0007131 | 6 | 20.1239536 | 4.33263058 | 3.41E-06 | 3.59E-06 |
| rs9834142 | 3 | 24702416 | IMPUTED | MIR4792(dist=139490),RARB(dist=168398) | A | G | 0.00071333 | 6 | 20.1270004 | 4.33361816 | 3.41E-06 | 3.60E-06 |
| rs9843065 | 3 | 24707247 | IMPUTED | MIR4792(dist=144321),RARB(dist=163567) | T | C | 0.00071322 | 6 | 20.1261189 | 4.3343475 | 3.43E-06 | 3.62E-06 |
| rs187469016 | 5 | 100024484 | IMPUTED | FAM174A(dist=102040),ST8SIA4(dist=118155) | G | A | 0.00276004 | 23 | -10.383076 | 2.23651559 | 3.44E-06 | 3.63E-06 |
| rs531086075 | 1 | 96278703 | IMPUTED | LOC100996635(dist=297683),LOC102723661(dist=178921) | T | G | 0.00052365 | 4 | 25.5174039 | 5.4965276 | 3.44E-06 | 3.63E-06 |
| rs879852680 | 11 | 45482976 | IMPUTED | LOC399886(dist=72917),CHST1(dist=186263) | G | A | 0.00012515 | 1 | -45.366736 | 9.77346282 | 3.45E-06 | 3.65E-06 |
| rs534827699 | 1 | 104801983 | IMPUTED | LOC100129138(dist=182290),LINC01676(dist=1330333) | C | A | 0.00037129 | 3 | 30.8707829 | 6.65087613 | 3.46E-06 | 3.65E-06 |
| rs902095714 | 1 | 18195234 | IMPUTED | ACTL8(dist=41676),LINC01654(dist=196917) | G | A | 0.0002062 | 2 | -40.687879 | 8.76866267 | 3.48E-06 | 3.67E-06 |
| rs151046299 | 20 | 17166317 | IMPUTED | OTOR(dist=433508),PCSK2(dist=40435) | A | G | 0.00036332 | 3 | -26.254153 | 5.65843531 | 3.49E-06 | 3.68E-06 |
| rs760246630 | 2 | 24976439 | IMPUTED | NCOA1 | A | G | 0.00012978 | 1 | -45.607691 | 9.82989487 | 3.49E-06 | 3.68E-06 |
| rs557024846 | 11 | 39763763 | IMPUTED | LINC01493(dist=1086964),LRRC4C(dist=371988) | G | A | 0.00013513 | 1 | -45.826183 | 9.87839019 | 3.50E-06 | 3.69E-06 |
| rs180916150 | 6 | 53757816 | IMPUTED | LRRC1 | C | T | 0.00079189 | 7 | 20.4459459 | 4.40841659 | 3.52E-06 | 3.71E-06 |
| rs114438921 | 19 | 549382 | IMPUTED | GZMM | G | T | 0.00039256 | 3 | 30.6365738 | 6.60570674 | 3.52E-06 | 3.71E-06 |
| rs75407235 | 1 | 188102370 | IMPUTED | LINC01037(dist=656016),BRINP3(dist=1964427) | A | T | 0.00022058 | 2 | -37.8598 | 8.16403952 | 3.53E-06 | 3.72E-06 |
| rs1014454058 | 12 | 87220823 | IMPUTED | MGAT4C | A | G | 0.00041336 | 3 | 31.1301555 | 6.71299054 | 3.53E-06 | 3.73E-06 |
| rs577850512 | 8 | 25888477 | IMPUTED | EBF2 | A | T | 0.00059484 | 5 | 24.5230451 | 5.28912331 | 3.54E-06 | 3.74E-06 |
| rs146565413 | 17 | 30027922 | IMPUTED | MIR365B(dist=125382),COPRS(dist=150962) | A | G | 0.00038864 | 3 | 30.4530342 | 6.57309183 | 3.60E-06 | 3.80E-06 |
| rs1052356610 | 11 | 120408875 | IMPUTED | GRIK4 | C | T | 0.00038127 | 3 | 30.7228502 | 6.63319798 | 3.63E-06 | 3.83E-06 |
| rs143685503 | 19 | 35088376 | IMPUTED | SCGB1B2P | C | T | 0.00242667 | 20 | 12.1502717 | 2.62413016 | 3.65E-06 | 3.85E-06 |
| rs192971782 | 2 | 96901964 | IMPUTED | STARD7-AS1 | A | C | 0.00179629 | 15 | 13.1956914 | 2.85062883 | 3.67E-06 | 3.88E-06 |
| rs545083715 | 18 | 72851707 | IMPUTED | ZNF407(dist=74079),ZADH2(dist=55358) | C | T | 0.00041669 | 4 | 30.1266592 | 6.5088483 | 3.68E-06 | 3.88E-06 |
| rs7637378 | 3 | 24665688 | IMPUTED | MIR4792(dist=102762),RARB(dist=205126) | T | A | 0.00074329 | 6 | 20.0331451 | 4.32885157 | 3.70E-06 | 3.90E-06 |
| rs142776406 | 20 | 17166309 | IMPUTED | OTOR(dist=433500),PCSK2(dist=40443) | G | A | 0.00036071 | 3 | -26.36277 | 5.69791167 | 3.71E-06 | 3.92E-06 |
| rs9834456 | 3 | 24672478 | IMPUTED | MIR4792(dist=109552),RARB(dist=198336) | A | G | 0.00071702 | 6 | 20.0291614 | 4.33223493 | 3.78E-06 | 3.98E-06 |
| rs55723486 | 3 | 24671717 | IMPUTED | MIR4792(dist=108791),RARB(dist=199097) | G | C | 0.0007194 | 6 | 20.0260897 | 4.33214515 | 3.79E-06 | 4.00E-06 |
| rs377455263 | 16 | 87365020 | IMPUTED | FBXO31 | G | A | 0.00012467 | 1 | -45.117843 | 9.76211301 | 3.81E-06 | 4.01E-06 |
| rs55999886 | 3 | 24692886 | IMPUTED | MIR4792(dist=129960),RARB(dist=177928) | G | A | 0.00073176 | 6 | 20.0201255 | 4.33215242 | 3.81E-06 | 4.02E-06 |
| rs113920675 | 8 | 82440603 | IMPUTED | FABP12 | C | T | 0.00030307 | 3 | -31.498871 | 6.81657991 | 3.82E-06 | 4.03E-06 |
| rs75425170 | 2 | 46057870 | IMPUTED | PRKCE | C | A | 0.00101331 | 9 | 18.9312021 | 4.09745075 | 3.83E-06 | 4.04E-06 |
| rs190128772 | 18 | 53360845 | IMPUTED | TCF4(dist=57621),LINC01415(dist=80987) | C | T | 0.00034015 | 3 | -33.591212 | 7.27060337 | 3.83E-06 | 4.04E-06 |
| rs142018669 | 12 | 53411481 | IMPUTED | EIF4B | G | A | 0.00023283 | 2 | -36.641189 | 7.93551955 | 3.89E-06 | 4.10E-06 |
| rs190625552 | 8 | 82753377 | IMPUTED | SNX16 | G | T | 0.00013834 | 1 | -45.050319 | 9.75731179 | 3.89E-06 | 4.10E-06 |
| rs9861176 | 3 | 24670141 | IMPUTED | MIR4792(dist=107215),RARB(dist=200673) | C | G | 0.00073092 | 6 | 19.9860983 | 4.32923906 | 3.90E-06 | 4.11E-06 |
| rs560162233 | 2 | 52690443 | IMPUTED | LOC730100(dist=55388),MIR4431(dist=239217) | A | G | 0.00036059 | 3 | 30.5772449 | 6.62496195 | 3.92E-06 | 4.13E-06 |
| rs552544727 | 19 | 460642 | IMPUTED | SHC2 | C | A | 0.00040991 | 3 | 30.4933591 | 6.60730772 | 3.93E-06 | 4.14E-06 |
| rs543535159 | 3 | 151581475 | IMPUTED | AADACL2-AS1 | C | T | 0.00054338 | 5 | -24.316008 | 5.26923475 | 3.94E-06 | 4.15E-06 |

| | | | | | | | | | | | | |
|---|---|---|---|---|---|---|---|---|---|---|---|---|
| rs58604397 | 5 | 171259455 | IMPUTED | SMIM23(dist=41363),FBXW11(dist=29101) | A | C | 0.00035548 | 3 | -28.075567 | 6.08406168 | 3.94E-06 | 4.15E-06 |
| rs780871194 | 14 | 103486835 | IMPUTED | CDC42BPB | C | T | 0.00037556 | 3 | 30.5194543 | 6.61441448 | 3.95E-06 | 4.16E-06 |
| rs6883878 | 5 | 171271875 | IMPUTED | SMIM23(dist=53783),FBXW11(dist=16681) | C | T | 0.00035441 | 3 | -28.129904 | 6.09656251 | 3.95E-06 | 4.16E-06 |
| rs73328039 | 5 | 171260229 | IMPUTED | SMIM23(dist=42137),FBXW11(dist=28327) | G | A | 0.00035702 | 3 | -27.831079 | 6.03551502 | 4.00E-06 | 4.22E-06 |
| rs192882754 | 1 | 152019397 | IMPUTED | S100A11(dist=9886),LOC100131107(dist=32492) | G | A | 0.00465248 | 39 | 9.09471042 | 1.97243115 | 4.01E-06 | 4.23E-06 |
| rs139487541 | 2 | 52526697 | IMPUTED | LOC730100 | C | T | 0.00036095 | 3 | 30.5481115 | 6.6253629 | 4.01E-06 | 4.23E-06 |
| rs57657657 | 5 | 171259464 | GENOTYPED | SMIM23(dist=41372),FBXW11(dist=29092) | A | C | 0.00035655 | 3 | -27.822063 | 6.03551043 | 4.03E-06 | 4.25E-06 |
| rs73328032 | 5 | 171259886 | IMPUTED | SMIM23(dist=41794),FBXW11(dist=28670) | C | G | 0.00035655 | 3 | -27.822063 | 6.03551043 | 4.03E-06 | 4.25E-06 |
| rs73328034 | 5 | 171259906 | IMPUTED | SMIM23(dist=41814),FBXW11(dist=28650) | C | A | 0.00035655 | 3 | -27.822063 | 6.03551043 | 4.03E-06 | 4.25E-06 |
| rs73328036 | 5 | 171260179 | GENOTYPED | SMIM23(dist=42087),FBXW11(dist=28377) | C | T | 0.00035655 | 3 | -27.822063 | 6.03551043 | 4.03E-06 | 4.25E-06 |
| rs57685840 | 5 | 171264540 | IMPUTED | SMIM23(dist=46448),FBXW11(dist=24016) | T | C | 0.00036 | 3 | -27.908317 | 6.05491812 | 4.04E-06 | 4.26E-06 |
| rs572790669 | 9 | 86148883 | IMPUTED | FRMD3 | T | C | 0.00032791 | 3 | -30.41727 | 6.59934107 | 4.04E-06 | 4.26E-06 |
| rs115700753 | 17 | 30029860 | IMPUTED | MIR365B(dist=127320),COPRS(dist=149024) | T | C | 0.00038091 | 3 | 30.4728391 | 6.611845 | 4.05E-06 | 4.27E-06 |
| rs6555974 | 5 | 171263439 | IMPUTED | SMIM23(dist=45347),FBXW11(dist=25117) | T | C | 0.00035738 | 3 | -27.870529 | 6.04725548 | 4.05E-06 | 4.27E-06 |
| rs73328044 | 5 | 171260492 | IMPUTED | SMIM23(dist=42400),FBXW11(dist=28064) | C | T | 0.00035738 | 3 | -27.814486 | 6.03550683 | 4.06E-06 | 4.27E-06 |
| rs560100734 | 2 | 52700082 | IMPUTED | LOC730100(dist=65027),MIR4431(dist=229578) | A | G | 0.0003594 | 3 | 30.5278739 | 6.62543613 | 4.07E-06 | 4.29E-06 |
| rs74095773 | 12 | 60226569 | IMPUTED | SLC16A7(dist=42934),FAM19A2(dist=1875460) | C | T | 0.00039244 | 3 | 30.4976571 | 6.62072279 | 4.10E-06 | 4.32E-06 |
| rs189412822 | 2 | 59920678 | IMPUTED | LINC01793(dist=414143),MIR4432HG(dist=665673) | A | G | 0.00042025 | 4 | 30.2814231 | 6.57408488 | 4.10E-06 | 4.32E-06 |
| rs769885627 | 20 | 3820067 | IMPUTED | AP5S1(dist=14113),MAVS(dist=7379) | T | C | 0.00013418 | 1 | -45.228208 | 9.8192466 | 4.10E-06 | 4.32E-06 |
| rs184293531 | 2 | 59037301 | IMPUTED | LINC01122 | T | C | 0.00036237 | 3 | 30.532403 | 6.63033564 | 4.13E-06 | 4.35E-06 |
| rs543405722 | 1 | 19207966 | IMPUTED | ALDH4A1 | A | G | 0.00014904 | 1 | 44.3815404 | 9.63844765 | 4.13E-06 | 4.35E-06 |
| rs766959296 | 17 | 16996395 | IMPUTED | MPRIP | G | A | 0.00037224 | 3 | 30.4949435 | 6.62282658 | 4.13E-06 | 4.36E-06 |
| rs569238584 | 11 | 133008828 | IMPUTED | OPCML | C | T | 0.00040207 | 3 | 30.3082809 | 6.58300339 | 4.14E-06 | 4.37E-06 |
| rs73328048 | 5 | 171263665 | IMPUTED | SMIM23(dist=45573),FBXW11(dist=24891) | G | T | 0.00035881 | 3 | -27.851716 | 6.05105789 | 4.17E-06 | 4.39E-06 |
| rs143690756 | 9 | 112065921 | IMPUTED | EPB41L4B | C | A | 0.0092112 | 78 | -5.7062344 | 1.23982989 | 4.18E-06 | 4.40E-06 |
| rs56005337 | 5 | 171263728 | IMPUTED | SMIM23(dist=45636),FBXW11(dist=24828) | G | A | 0.0004092 | 3 | -27.160673 | 5.90568189 | 4.24E-06 | 4.47E-06 |
| rs76113994 | 2 | 151595775 | IMPUTED | LOC101929282(dist=103904),RBM43(dist=508953) | C | T | 0.00041883 | 4 | -28.506986 | 6.19877512 | 4.25E-06 | 4.48E-06 |
| rs561772098 | 2 | 53168754 | IMPUTED | MIR4431(dist=239001),ASB3(dist=728363) | G | A | 0.00036118 | 3 | 30.4643865 | 6.62530857 | 4.26E-06 | 4.49E-06 |
| rs567670762 | 2 | 52692097 | IMPUTED | LOC730100(dist=57042),MIR4431(dist=237563) | A | C | 0.00035845 | 3 | 30.4641286 | 6.62605292 | 4.27E-06 | 4.50E-06 |
| rs558373680 | 2 | 52697272 | IMPUTED | LOC730100(dist=62217),MIR4431(dist=232388) | G | T | 0.00035845 | 3 | 30.4641286 | 6.62605292 | 4.27E-06 | 4.50E-06 |
| rs145480617 | 2 | 26707907 | IMPUTED | OTOF | G | A | 0.00473794 | 40 | 8.68752713 | 1.88958906 | 4.27E-06 | 4.50E-06 |
| rs532102415 | 2 | 53190323 | IMPUTED | MIR4431(dist=260570),ASB3(dist=706794) | T | C | 0.00036511 | 3 | 30.4430034 | 6.62464137 | 4.32E-06 | 4.55E-06 |
| rs569942221 | 2 | 53191303 | IMPUTED | MIR4431(dist=261550),ASB3(dist=705814) | C | T | 0.00036511 | 3 | 30.4430034 | 6.62464137 | 4.32E-06 | 4.55E-06 |
| rs556549829 | 2 | 53193143 | IMPUTED | MIR4431(dist=263390),ASB3(dist=703974) | A | C | 0.00036511 | 3 | 30.4430034 | 6.62464137 | 4.32E-06 | 4.55E-06 |
| rs567607749 | 3 | 151202307 | IMPUTED | IGSF10(dist=25810),MIR5186(dist=81357) | T | C | 0.00058949 | 5 | -24.278583 | 5.28406735 | 4.33E-06 | 4.56E-06 |
| rs551354564 | 2 | 52689780 | IMPUTED | LOC730100(dist=54725),MIR4431(dist=239880) | A | T | 0.00035809 | 3 | 30.4397671 | 6.62623531 | 4.35E-06 | 4.58E-06 |
| rs539473468 | 2 | 172183467 | IMPUTED | METTL8 | T | C | 0.00039244 | 3 | 30.8579847 | 6.717395 | 4.35E-06 | 4.59E-06 |
| rs1006366033 | 9 | 74292533 | IMPUTED | TRPM3(dist=556019),TMEM2(dist=5749) | G | A | 0.00036237 | 3 | 30.4341595 | 6.6259248 | 4.37E-06 | 4.60E-06 |
| rs578084149 | 2 | 53184688 | IMPUTED | MIR4431(dist=254935),ASB3(dist=712429) | T | A | 0.00036368 | 3 | 30.4268661 | 6.62525423 | 4.38E-06 | 4.61E-06 |
| rs113523800 | 19 | 428527 | IMPUTED | SHC2 | G | A | 0.00036059 | 3 | 31.2731041 | 6.81005985 | 4.39E-06 | 4.62E-06 |
| rs536554775 | 19 | 8120238 | IMPUTED | CCL25 | G | C | 0.00021013 | 2 | -38.8663 | 8.46362698 | 4.39E-06 | 4.62E-06 |
| rs151263175 | 17 | 30061792 | IMPUTED | MIR365B(dist=159252),COPRS(dist=117092) | G | A | 0.0003714 | 3 | 30.4242454 | 6.62564238 | 4.39E-06 | 4.63E-06 |
| rs545597295 | 2 | 53184895 | IMPUTED | MIR4431(dist=255142),ASB3(dist=712222) | C | T | 0.00036332 | 3 | 30.4226768 | 6.62539078 | 4.39E-06 | 4.63E-06 |
| rs115302759 | 6 | 91606574 | IMPUTED | MAP3K7(dist=309554),MIR4643(dist=624804) | T | C | 0.0011885 | 10 | 15.4559054 | 3.36614142 | 4.40E-06 | 4.63E-06 |
| rs183211857 | 5 | 180640700 | IMPUTED | TRIM7(dist=8407),MIR4638(dist=8866) | T | C | 0.00036011 | 3 | 30.4236937 | 6.62618993 | 4.40E-06 | 4.64E-06 |
| rs562402350 | 2 | 52655614 | IMPUTED | LOC730100(dist=20559),MIR4431(dist=274046) | G | A | 0.00036594 | 3 | 30.4178372 | 6.62535869 | 4.41E-06 | 4.64E-06 |
| rs544182806 | 11 | 58770212 | IMPUTED | LOC283194 | A | G | 0.00014072 | 1 | -44.718842 | 9.74117761 | 4.42E-06 | 4.65E-06 |
| rs373074099 | 18 | 72153912 | IMPUTED | FAM69C(dist=29409),CNDP2(dist=9588) | G | A | 0.00127989 | 11 | 15.6305003 | 3.40570129 | 4.44E-06 | 4.68E-06 |
| rs779721380 | 11 | 5683159 | IMPUTED | TRIM6-TRIM34(dist=17534),TRIM5(dist=1266) | G | A | 0.00038864 | 3 | 33.1404517 | 7.22108595 | 4.45E-06 | 4.68E-06 |
| rs113851346 | 17 | 30035361 | IMPUTED | MIR365B(dist=132821),COPRS(dist=143523) | G | A | 0.00036332 | 3 | 30.4099148 | 6.6262451 | 4.45E-06 | 4.68E-06 |
| rs768176549 | 6 | 151784878 | IMPUTED | ARMT1 | C | G | 0.00041205 | 3 | 30.2240762 | 6.58617185 | 4.45E-06 | 4.69E-06 |
| rs191931350 | 6 | 91618301 | IMPUTED | MAP3K7(dist=321281),MIR4643(dist=613077) | C | T | 0.0011885 | 10 | 15.4164899 | 3.35982228 | 4.46E-06 | 4.70E-06 |

| | | | | | | | | | | | | |
|---|---|---|---|---|---|---|---|---|---|---|---|---|
| rs6920882 | 6 | 91618401 | GENOTYPED | MAP3K7(dist=321381),MIR4643(dist=612977) | T | C | 0.0011885 | 10 | 15.4164899 | 3.35982228 | 4.46E-06 | 4.70E-06 |
| rs369292656 | 2 | 52727688 | IMPUTED | LOC730100(dist=92633),MIR4431(dist=201972) | G | A | 0.00039125 | 3 | 30.3026987 | 6.6055844 | 4.49E-06 | 4.72E-06 |
| rs185741126 | 5 | 180611687 | IMPUTED | OR2V2(dist=28797),LINC01962(dist=6359) | C | T | 0.00036083 | 3 | 30.3963438 | 6.62614108 | 4.49E-06 | 4.73E-06 |
| rs531114386 | 2 | 52609783 | IMPUTED | LINC01867,LOC730100 | T | G | 0.0003594 | 3 | 30.3959923 | 6.62664088 | 4.50E-06 | 4.74E-06 |
| rs559962898 | 2 | 52606500 | IMPUTED | LINC01867,LOC730100 | G | A | 0.00036356 | 3 | 30.3940991 | 6.62659182 | 4.50E-06 | 4.74E-06 |
| rs113182702 | 2 | 16577350 | IMPUTED | GACAT3(dist=351539),FAM49A(dist=153380) | A | G | 0.00025481 | 2 | -32.268864 | 7.03550682 | 4.51E-06 | 4.74E-06 |
| rs563891857 | 2 | 52636044 | IMPUTED | LOC730100(dist=989) | T | C | 0.00038377 | 3 | 30.375517 | 6.6228615 | 4.51E-06 | 4.75E-06 |
| rs565163634 | 9 | 122593493 | IMPUTED | BRINP1(dist=461754),LINC01613(dist=103845) | T | C | 0.00081899 | 7 | 20.5331856 | 4.47695348 | 4.51E-06 | 4.75E-06 |
| rs113793355 | 19 | 483664 | IMPUTED | ODF3L2(dist=8681),MADCAM1(dist=12826) | G | A | 0.00035298 | 3 | 30.919049 | 6.74170613 | 4.51E-06 | 4.75E-06 |
| rs574982708 | 6 | 45921190 | IMPUTED | CLIC5 | G | A | 0.00014999 | 1 | -56.363037 | 12.2900527 | 4.52E-06 | 4.76E-06 |
| rs114294515 | 5 | 180619981 | IMPUTED | TRIM7(dist=943) | T | C | 0.00036083 | 3 | 30.384878 | 6.62620782 | 4.53E-06 | 4.77E-06 |
| rs116838278 | 5 | 180633941 | IMPUTED | TRIM7(dist=1648),MIR4638(dist=15625) | T | G | 0.0003613 | 3 | 30.3790713 | 6.62620044 | 4.55E-06 | 4.79E-06 |
| rs536174889 | 2 | 53182125 | IMPUTED | MIR4431(dist=252372),ASB3(dist=714992) | A | G | 0.00035976 | 3 | 30.3773808 | 6.62638935 | 4.56E-06 | 4.80E-06 |
| rs144023056 | 12 | 63197659 | IMPUTED | PPM1H | C | T | 0.00035904 | 3 | 30.3766923 | 6.6264064 | 4.56E-06 | 4.80E-06 |
| rs74464242 | 1 | 187976734 | IMPUTED | LINC01037(dist=530380),NONE(dist=NONE) | C | T | 0.00021393 | 2 | -37.201794 | 8.11536296 | 4.56E-06 | 4.80E-06 |
| rs534063655 | 2 | 52525631 | IMPUTED | LOC730100 | G | A | 0.00038614 | 3 | 30.3694855 | 6.62524755 | 4.56E-06 | 4.80E-06 |
| rs77173096 | 1 | 97893807 | IMPUTED | DPYD | G | A | 0.00033159 | 3 | 33.2469645 | 7.25359442 | 4.57E-06 | 4.81E-06 |
| rs138545977 | 1 | 187979976 | IMPUTED | LINC01037(dist=533622),NONE(dist=NONE) | C | T | 0.0002144 | 2 | -37.195431 | 8.1153905 | 4.58E-06 | 4.82E-06 |
| rs537003747 | 2 | 55990459 | IMPUTED | PNPT1(dist=69414),EFEMP1(dist=102638) | A | G | 0.00035797 | 3 | 31.3605799 | 6.84285109 | 4.58E-06 | 4.83E-06 |
| rs138279271 | 16 | 61055027 | IMPUTED | LOC729159(dist=661330),MIR4426(dist=34584) | G | A | 0.00340623 | 29 | -8.8538829 | 1.93210511 | 4.59E-06 | 4.84E-06 |
| rs370077527 | 2 | 53107179 | IMPUTED | MIR4431(dist=177426),ASB3(dist=789938) | T | C | 0.00035738 | 3 | 30.3664276 | 6.62677359 | 4.60E-06 | 4.84E-06 |
| rs9309230 | 2 | 52749068 | IMPUTED | LOC730100(dist=114013),MIR4431(dist=180592) | T | C | 0.00035881 | 3 | 30.3636701 | 6.62666809 | 4.60E-06 | 4.85E-06 |
| rs75390181 | 2 | 52956699 | IMPUTED | MIR4431(dist=26946),ASB3(dist=940418) | G | C | 0.00035762 | 3 | 30.3612696 | 6.626642 | 4.61E-06 | 4.85E-06 |
| rs542361546 | 2 | 52622737 | IMPUTED | LOC730100 | A | G | 0.00035869 | 3 | 30.361632 | 6.62678732 | 4.61E-06 | 4.86E-06 |
| rs372194531 | 2 | 52718516 | IMPUTED | LOC730100(dist=83461),MIR4431(dist=211144) | G | C | 0.00035786 | 3 | 30.3610307 | 6.62670306 | 4.61E-06 | 4.86E-06 |
| rs565615053 | 2 | 53181004 | IMPUTED | MIR4431(dist=251251),ASB3(dist=716113) | C | G | 0.00035833 | 3 | 30.357533 | 6.62660185 | 4.62E-06 | 4.87E-06 |
| rs17119918 | 11 | 116585613 | IMPUTED | LOC101929011(dist=56644),BUD13(dist=33273) | G | A | 0.0003638 | 3 | 30.341258 | 6.62311748 | 4.63E-06 | 4.87E-06 |
| rs548028748 | 2 | 52720097 | IMPUTED | LOC730100(dist=85042),MIR4431(dist=209563) | G | A | 0.0003575 | 3 | 30.3559562 | 6.6266542 | 4.63E-06 | 4.87E-06 |
| rs570153897 | 2 | 52610437 | IMPUTED | LINC01867,LOC730100 | C | T | 0.00035845 | 3 | 30.3560879 | 6.62678112 | 4.63E-06 | 4.88E-06 |
| rs369180864 | 2 | 52715235 | IMPUTED | LOC730100(dist=80180),MIR4431(dist=214425) | C | G | 0.00035786 | 3 | 30.3551454 | 6.62671106 | 4.63E-06 | 4.88E-06 |
| rs532715151 | 2 | 52720079 | IMPUTED | LOC730100(dist=85024),MIR4431(dist=209581) | T | G | 0.00035714 | 3 | 30.354095 | 6.62667377 | 4.64E-06 | 4.88E-06 |
| rs369287560 | 2 | 53023383 | IMPUTED | MIR4431(dist=93630),ASB3(dist=873734) | G | A | 0.00035928 | 3 | 30.3523325 | 6.62669034 | 4.64E-06 | 4.89E-06 |
| rs960532183 | 9 | 119508058 | IMPUTED | ASTN2 | C | T | 0.0003594 | 3 | 30.3517362 | 6.62668329 | 4.64E-06 | 4.89E-06 |
| rs369972488 | 2 | 52719190 | IMPUTED | LOC730100(dist=84135),MIR4431(dist=210470) | A | G | 0.00035702 | 3 | 30.3495988 | 6.62668846 | 4.65E-06 | 4.90E-06 |
| rs7257230 | 19 | 454351 | IMPUTED | SHC2 | C | A | 0.00036748 | 3 | 30.3385369 | 6.62428193 | 4.65E-06 | 4.90E-06 |
| rs528976078 | 2 | 52640735 | IMPUTED | LOC730100(dist=5680),MIR4431(dist=288925) | T | G | 0.00035845 | 3 | 30.3490075 | 6.62664491 | 4.65E-06 | 4.90E-06 |
| rs111661085 | 9 | 88730110 | IMPUTED | GOLM1(dist=14994),LOC101927623(dist=12345) | G | A | 0.00098835 | 8 | 19.1314239 | 4.17733725 | 4.65E-06 | 4.90E-06 |
| rs546783102 | 2 | 52871208 | IMPUTED | LOC730100(dist=236153),MIR4431(dist=58452) | C | T | 0.00035809 | 3 | 30.3484305 | 6.62660881 | 4.65E-06 | 4.90E-06 |
| rs543762175 | 2 | 52604220 | IMPUTED | LINC01867,LOC730100 | T | G | 0.00035833 | 3 | 30.3492248 | 6.62679021 | 4.65E-06 | 4.90E-06 |
| rs565285485 | 2 | 52604228 | IMPUTED | LINC01867,LOC730100 | A | G | 0.00035833 | 3 | 30.3492248 | 6.62679021 | 4.65E-06 | 4.90E-06 |
| rs545828979 | 2 | 52617216 | IMPUTED | LOC730100 | G | C | 0.00035833 | 3 | 30.3492248 | 6.62679021 | 4.65E-06 | 4.90E-06 |
| rs531226054 | 2 | 52623488 | IMPUTED | LOC730100 | T | A | 0.00035833 | 3 | 30.3492248 | 6.62679021 | 4.65E-06 | 4.90E-06 |
| rs551597492 | 2 | 52655879 | IMPUTED | LOC730100(dist=20824),MIR4431(dist=273781) | C | G | 0.00035904 | 3 | 30.3476019 | 6.62650235 | 4.66E-06 | 4.90E-06 |
| rs373811446 | 2 | 52716581 | IMPUTED | LOC730100(dist=81526),MIR4431(dist=213079) | A | G | 0.00035714 | 3 | 30.3482375 | 6.62668478 | 4.66E-06 | 4.90E-06 |
| rs367623001 | 2 | 52717034 | IMPUTED | LOC730100(dist=81979),MIR4431(dist=212626) | A | G | 0.00035809 | 3 | 30.3473117 | 6.62654838 | 4.66E-06 | 4.90E-06 |
| rs1526679 | 2 | 52644317 | IMPUTED | LOC730100(dist=9262),MIR4431(dist=285343) | A | G | 0.00035845 | 3 | 30.3464967 | 6.62637395 | 4.66E-06 | 4.90E-06 |
| rs377200444 | 2 | 52956505 | IMPUTED | MIR4431(dist=26752),ASB3(dist=940612) | T | C | 0.00035738 | 3 | 30.3470745 | 6.62663116 | 4.66E-06 | 4.90E-06 |
| rs369146550 | 2 | 53010458 | IMPUTED | MIR4431(dist=80705),ASB3(dist=886659) | G | A | 0.00035988 | 3 | 30.3459672 | 6.62661838 | 4.66E-06 | 4.91E-06 |
| rs2356041 | 2 | 52598458 | IMPUTED | LINC01867,LOC730100 | T | C | 0.00035797 | 3 | 30.3457822 | 6.62679028 | 4.67E-06 | 4.91E-06 |
| rs7256741 | 19 | 453898 | IMPUTED | SHC2 | G | A | 0.00037283 | 3 | 30.3315526 | 6.62377141 | 4.67E-06 | 4.91E-06 |
| rs539458152 | 11 | 116520256 | IMPUTED | LOC101929011 | C | T | 0.0003575 | 3 | 30.375172 | 6.63331343 | 4.67E-06 | 4.91E-06 |
| rs371212315 | 2 | 52956080 | IMPUTED | MIR4431(dist=26327),ASB3(dist=941037) | T | G | 0.00035702 | 3 | 30.344431 | 6.62667472 | 4.67E-06 | 4.91E-06 |

| | | | | | | | | | | | | |
|---|---|---|---|---|---|---|---|---|---|---|---|---|
| rs1358186 | 2 | 52642930 | IMPUTED | LOC730100(dist=7875),MIR4431(dist=286730) | C | T | 0.00035786 | 3 | 30.3435244 | 6.62667205 | 4.67E-06 | 4.92E-06 |
| rs533062649 | 2 | 52655676 | IMPUTED | LOC730100(dist=20621),MIR4431(dist=273984) | A | G | 0.00035797 | 3 | 30.34191 | 6.62664663 | 4.68E-06 | 4.92E-06 |
| rs571326552 | 2 | 53180537 | IMPUTED | MIR4431(dist=250784),ASB3(dist=716580) | G | A | 0.00035726 | 3 | 30.3420002 | 6.62669109 | 4.68E-06 | 4.92E-06 |
| rs575452662 | 2 | 52616725 | IMPUTED | LINC01867 | G | T | 0.00037699 | 3 | 30.3360657 | 6.62553783 | 4.68E-06 | 4.93E-06 |
| rs574775108 | 2 | 52645035 | IMPUTED | LOC730100(dist=9980),MIR4431(dist=284625) | C | A | 0.00035762 | 3 | 30.3407461 | 6.62661986 | 4.68E-06 | 4.93E-06 |
| rs369033119 | 2 | 52723678 | IMPUTED | LOC730100(dist=88623),MIR4431(dist=205982) | C | T | 0.00035679 | 3 | 30.3405234 | 6.62670883 | 4.68E-06 | 4.93E-06 |
| rs368825583 | 2 | 52955395 | IMPUTED | MIR4431(dist=25642),ASB3(dist=941722) | T | C | 0.00035691 | 3 | 30.3398512 | 6.62667976 | 4.68E-06 | 4.93E-06 |
| rs569134995 | 2 | 52955412 | IMPUTED | MIR4431(dist=25659),ASB3(dist=941705) | A | G | 0.00035691 | 3 | 30.3398512 | 6.62667976 | 4.68E-06 | 4.93E-06 |
| rs564794266 | 2 | 52688608 | IMPUTED | LOC730100(dist=53553),MIR4431(dist=241052) | T | G | 0.00035667 | 3 | 30.3398426 | 6.62669792 | 4.68E-06 | 4.93E-06 |
| rs374768224 | 2 | 52507602 | IMPUTED | LOC730100 | C | T | 0.00035833 | 3 | 30.3492169 | 6.62880352 | 4.69E-06 | 4.93E-06 |
| rs373312188 | 2 | 52916201 | IMPUTED | LOC730100(dist=281146),MIR4431(dist=13459) | A | G | 0.00035691 | 3 | 30.3387498 | 6.62669782 | 4.69E-06 | 4.93E-06 |
| rs367979351 | 2 | 52735275 | IMPUTED | LOC730100(dist=100220),MIR4431(dist=194385) | G | A | 0.00035691 | 3 | 30.338693 | 6.62670358 | 4.69E-06 | 4.93E-06 |
| rs186240407 | 2 | 53136950 | IMPUTED | MIR4431(dist=207197),ASB3(dist=760167) | C | T | 0.00036665 | 3 | 30.3321996 | 6.62547443 | 4.69E-06 | 4.94E-06 |
| rs376654178 | 2 | 52955934 | IMPUTED | MIR4431(dist=26181),ASB3(dist=941183) | T | C | 0.00035845 | 3 | 30.3371351 | 6.62667701 | 4.69E-06 | 4.94E-06 |
| rs115761220 | 2 | 52544377 | IMPUTED | LOC730100 | T | A | 0.00035667 | 3 | 30.3372686 | 6.62671523 | 4.69E-06 | 4.94E-06 |
| rs115791435 | 2 | 52544380 | IMPUTED | LOC730100 | C | T | 0.00035667 | 3 | 30.3372686 | 6.62671523 | 4.69E-06 | 4.94E-06 |
| rs115553765 | 2 | 52545698 | IMPUTED | LOC730100 | C | A | 0.00035667 | 3 | 30.3372686 | 6.62671523 | 4.69E-06 | 4.94E-06 |
| rs371862794 | 2 | 52908691 | IMPUTED | LOC730100(dist=273636),MIR4431(dist=20969) | G | A | 0.00035702 | 3 | 30.3368534 | 6.62668143 | 4.69E-06 | 4.94E-06 |
| rs149851742 | 2 | 53008138 | IMPUTED | MIR4431(dist=78385),ASB3(dist=888979) | C | T | 0.00035691 | 3 | 30.3362328 | 6.62668865 | 4.70E-06 | 4.94E-06 |
| rs4386361 | 2 | 53008780 | IMPUTED | MIR4431(dist=79027),ASB3(dist=888337) | A | G | 0.00035691 | 3 | 30.3362328 | 6.62668865 | 4.70E-06 | 4.94E-06 |
| rs375787925 | 2 | 52895590 | IMPUTED | LOC730100(dist=260535),MIR4431(dist=34070) | T | C | 0.00035667 | 3 | 30.3359649 | 6.62671277 | 4.70E-06 | 4.94E-06 |
| rs371346212 | 2 | 52897986 | IMPUTED | LOC730100(dist=262931),MIR4431(dist=31674) | C | T | 0.00035667 | 3 | 30.3359649 | 6.62671277 | 4.70E-06 | 4.94E-06 |
| rs576662858 | 2 | 52714108 | IMPUTED | LOC730100(dist=79053),MIR4431(dist=215552) | A | G | 0.00035667 | 3 | 30.3359442 | 6.62671451 | 4.70E-06 | 4.94E-06 |
| rs6736060 | 2 | 52993749 | IMPUTED | MIR4431(dist=63996),ASB3(dist=903368) | A | G | 0.00035845 | 3 | 30.3351032 | 6.62656449 | 4.70E-06 | 4.95E-06 |
| rs144116923 | 2 | 52996116 | IMPUTED | MIR4431(dist=66363),ASB3(dist=901001) | A | G | 0.00035809 | 3 | 30.3344039 | 6.62656349 | 4.70E-06 | 4.95E-06 |
| rs539771008 | 2 | 53146868 | IMPUTED | MIR4431(dist=217115),ASB3(dist=750249) | A | G | 0.00035679 | 3 | 30.3344745 | 6.62671169 | 4.70E-06 | 4.95E-06 |
| rs1843038 | 2 | 52959633 | IMPUTED | MIR4431(dist=29880),ASB3(dist=937484) | C | A | 0.00035857 | 3 | 30.3341157 | 6.62666888 | 4.70E-06 | 4.95E-06 |
| rs555469022 | 2 | 52557221 | IMPUTED | LOC730100 | C | A | 0.00035679 | 3 | 30.3341993 | 6.62669196 | 4.70E-06 | 4.95E-06 |
| rs377759156 | 2 | 52954092 | IMPUTED | MIR4431(dist=24339),ASB3(dist=943025) | T | G | 0.00035667 | 3 | 30.3342029 | 6.62670771 | 4.70E-06 | 4.95E-06 |
| rs142898785 | 2 | 52999581 | IMPUTED | MIR4431(dist=69828),ASB3(dist=897536) | C | T | 0.00035679 | 3 | 30.3341223 | 6.62670224 | 4.70E-06 | 4.95E-06 |
| rs371650770 | 2 | 53037950 | IMPUTED | MIR4431(dist=108197),ASB3(dist=859167) | T | C | 0.00035667 | 3 | 30.3341145 | 6.62670479 | 4.70E-06 | 4.95E-06 |
| rs377442641 | 2 | 53038287 | IMPUTED | MIR4431(dist=108534),ASB3(dist=858830) | A | C | 0.00035667 | 3 | 30.3341145 | 6.62670479 | 4.70E-06 | 4.95E-06 |
| rs370510072 | 2 | 53039220 | IMPUTED | MIR4431(dist=109467),ASB3(dist=857897) | T | G | 0.00035667 | 3 | 30.3341145 | 6.62670479 | 4.70E-06 | 4.95E-06 |
| rs373725461 | 2 | 53039313 | IMPUTED | MIR4431(dist=109560),ASB3(dist=857804) | C | T | 0.00035667 | 3 | 30.3341145 | 6.62670479 | 4.70E-06 | 4.95E-06 |
| rs576529791 | 2 | 53046781 | IMPUTED | MIR4431(dist=117028),ASB3(dist=850336) | T | C | 0.00035667 | 3 | 30.3341145 | 6.62670479 | 4.70E-06 | 4.95E-06 |
| rs572170177 | 2 | 52804757 | IMPUTED | LOC730100(dist=169702),MIR4431(dist=124903) | A | G | 0.00035667 | 3 | 30.3339567 | 6.62670993 | 4.71E-06 | 4.95E-06 |
| rs561186991 | 2 | 52806051 | IMPUTED | LOC730100(dist=170996),MIR4431(dist=123609) | C | T | 0.00035667 | 3 | 30.3339567 | 6.62670993 | 4.71E-06 | 4.95E-06 |
| rs531508153 | 2 | 52807272 | IMPUTED | LOC730100(dist=172217),MIR4431(dist=122388) | G | A | 0.00035667 | 3 | 30.3339567 | 6.62670993 | 4.71E-06 | 4.95E-06 |
| rs375240447 | 19 | 471395 | IMPUTED | ODF3L2 | G | A | 0.00035738 | 3 | 30.3336305 | 6.62675184 | 4.71E-06 | 4.95E-06 |
| rs190537399 | 12 | 78933270 | IMPUTED | LINC02424(dist=179744),SYT1(dist=324503) | T | A | 0.00022807 | 2 | -37.693042 | 8.23455355 | 4.71E-06 | 4.95E-06 |
| rs146381520 | 12 | 60080996 | IMPUTED | SLC16A7 | G | C | 0.00035881 | 3 | 30.3323637 | 6.62652443 | 4.71E-06 | 4.95E-06 |
| rs11894007 | 2 | 52743925 | IMPUTED | LOC730100(dist=108870),MIR4431(dist=185735) | A | G | 0.00035726 | 3 | 30.3329009 | 6.6267223 | 4.71E-06 | 4.96E-06 |
| rs143416013 | 12 | 60099743 | IMPUTED | SLC16A7 | T | C | 0.00035952 | 3 | 30.330913 | 6.62640024 | 4.71E-06 | 4.96E-06 |
| rs139335791 | 12 | 60100424 | IMPUTED | SLC16A7 | T | A | 0.00035952 | 3 | 30.330913 | 6.62640024 | 4.71E-06 | 4.96E-06 |
| rs539714680 | 12 | 60101339 | IMPUTED | SLC16A7 | A | G | 0.00035952 | 3 | 30.330913 | 6.62640024 | 4.71E-06 | 4.96E-06 |
| rs375110574 | 2 | 53112450 | IMPUTED | MIR4431(dist=182697),ASB3(dist=784667) | A | C | 0.00035667 | 3 | 30.3319784 | 6.62671767 | 4.71E-06 | 4.96E-06 |
| rs374302389 | 2 | 53085983 | IMPUTED | MIR4431(dist=156230),ASB3(dist=811134) | C | T | 0.00035809 | 3 | 30.3315286 | 6.62669404 | 4.71E-06 | 4.96E-06 |
| rs115998730 | 12 | 60109624 | IMPUTED | SLC16A7 | T | C | 0.00035976 | 3 | 30.329787 | 6.6263431 | 4.71E-06 | 4.96E-06 |
| rs147462597 | 12 | 60121592 | IMPUTED | SLC16A7 | G | T | 0.00035976 | 3 | 30.329787 | 6.6263431 | 4.71E-06 | 4.96E-06 |
| rs187674492 | 12 | 60122836 | IMPUTED | SLC16A7 | T | G | 0.00035976 | 3 | 30.329787 | 6.6263431 | 4.71E-06 | 4.96E-06 |
| rs146527019 | 12 | 60159496 | IMPUTED | SLC16A7 | A | G | 0.00035976 | 3 | 30.329787 | 6.6263431 | 4.71E-06 | 4.96E-06 |
| rs73489637 | 19 | 452105 | GENOTYPED | SHC2 | T | C | 0.00035655 | 3 | 30.3313373 | 6.62671719 | 4.71E-06 | 4.96E-06 |

| | | | | | | | | | | | | |
|---|---|---|---|---|---|---|---|---|---|---|---|---|
| rs141882698 | 11 | 116601338 | IMPUTED | LOC101929011(dist=72369),BUD13(dist=17548) | A | C | 0.00035655 | 3 | 30.3313373 | 6.62671719 | 4.71E-06 | 4.96E-06 |
| rs544617045 | 2 | 52541999 | IMPUTED | LOC730100 | G | A | 0.00035655 | 3 | 30.3313373 | 6.62671719 | 4.71E-06 | 4.96E-06 |
| rs115334491 | 2 | 52545366 | IMPUTED | LOC730100 | A | G | 0.00035655 | 3 | 30.3313373 | 6.62671719 | 4.71E-06 | 4.96E-06 |
| rs527392879 | 2 | 52547080 | IMPUTED | LOC730100 | C | T | 0.00035655 | 3 | 30.3313373 | 6.62671719 | 4.71E-06 | 4.96E-06 |
| rs372232719 | 2 | 52549212 | IMPUTED | LOC730100 | T | C | 0.00035655 | 3 | 30.3313373 | 6.62671719 | 4.71E-06 | 4.96E-06 |
| rs566577469 | 2 | 52549491 | IMPUTED | LOC730100 | G | T | 0.00035655 | 3 | 30.3313373 | 6.62671719 | 4.71E-06 | 4.96E-06 |
| rs555529050 | 2 | 52551773 | IMPUTED | LOC730100 | G | A | 0.00035655 | 3 | 30.3313373 | 6.62671719 | 4.71E-06 | 4.96E-06 |
| rs377373388 | 2 | 52553502 | IMPUTED | LOC730100 | C | T | 0.00035655 | 3 | 30.3313373 | 6.62671719 | 4.71E-06 | 4.96E-06 |
| rs544639429 | 2 | 52553534 | IMPUTED | LOC730100 | A | T | 0.00035655 | 3 | 30.3313373 | 6.62671719 | 4.71E-06 | 4.96E-06 |
| rs368726778 | 2 | 52553734 | IMPUTED | LOC730100 | T | C | 0.00035655 | 3 | 30.3313373 | 6.62671719 | 4.71E-06 | 4.96E-06 |
| rs377108751 | 2 | 52553792 | IMPUTED | LOC730100 | C | A | 0.00035655 | 3 | 30.3313373 | 6.62671719 | 4.71E-06 | 4.96E-06 |
| rs370510488 | 2 | 52553823 | IMPUTED | LOC730100 | G | A | 0.00035655 | 3 | 30.3313373 | 6.62671719 | 4.71E-06 | 4.96E-06 |
| rs376826721 | 2 | 52553993 | IMPUTED | LOC730100 | G | A | 0.00035655 | 3 | 30.3313373 | 6.62671719 | 4.71E-06 | 4.96E-06 |
| rs368715703 | 2 | 52554493 | IMPUTED | LOC730100 | G | A | 0.00035655 | 3 | 30.3313373 | 6.62671719 | 4.71E-06 | 4.96E-06 |
| rs571419511 | 2 | 52554607 | IMPUTED | LOC730100 | A | T | 0.00035655 | 3 | 30.3313373 | 6.62671719 | 4.71E-06 | 4.96E-06 |
| rs368523060 | 2 | 52559306 | IMPUTED | LOC730100 | A | G | 0.00035655 | 3 | 30.3313373 | 6.62671719 | 4.71E-06 | 4.96E-06 |
| rs571728366 | 2 | 52581588 | IMPUTED | LOC730100 | C | A | 0.00035655 | 3 | 30.3313373 | 6.62671719 | 4.71E-06 | 4.96E-06 |
| rs539209253 | 2 | 52582267 | IMPUTED | LOC730100 | A | G | 0.00035655 | 3 | 30.3313373 | 6.62671719 | 4.71E-06 | 4.96E-06 |
| rs540786989 | 2 | 52587032 | IMPUTED | LOC730100 | G | A | 0.00035655 | 3 | 30.3313373 | 6.62671719 | 4.71E-06 | 4.96E-06 |
| rs375126523 | 2 | 52587641 | IMPUTED | LOC730100 | G | A | 0.00035655 | 3 | 30.3313373 | 6.62671719 | 4.71E-06 | 4.96E-06 |
| rs561112087 | 2 | 52587714 | IMPUTED | LOC730100 | T | A | 0.00035655 | 3 | 30.3313373 | 6.62671719 | 4.71E-06 | 4.96E-06 |
| rs116707515 | 2 | 52679675 | GENOTYPED | LOC730100(dist=44620),MIR4431(dist=249985) | T | G | 0.00035655 | 3 | 30.3313373 | 6.62671719 | 4.71E-06 | 4.96E-06 |
| rs531499393 | 2 | 52680402 | IMPUTED | LOC730100(dist=45347),MIR4431(dist=249258) | A | G | 0.00035655 | 3 | 30.3313373 | 6.62671719 | 4.71E-06 | 4.96E-06 |
| rs538141757 | 2 | 52685905 | IMPUTED | LOC730100(dist=50850),MIR4431(dist=243755) | A | G | 0.00035655 | 3 | 30.3313373 | 6.62671719 | 4.71E-06 | 4.96E-06 |
| rs556861356 | 2 | 52686390 | IMPUTED | LOC730100(dist=51335),MIR4431(dist=243270) | T | A | 0.00035655 | 3 | 30.3313373 | 6.62671719 | 4.71E-06 | 4.96E-06 |
| rs572161315 | 2 | 52687214 | IMPUTED | LOC730100(dist=52159),MIR4431(dist=242446) | G | C | 0.00035655 | 3 | 30.3313373 | 6.62671719 | 4.71E-06 | 4.96E-06 |
| rs4380279 | 2 | 52688148 | IMPUTED | LOC730100(dist=53093),MIR4431(dist=241512) | C | T | 0.00035655 | 3 | 30.3313373 | 6.62671719 | 4.71E-06 | 4.96E-06 |
| rs546769066 | 2 | 52735604 | IMPUTED | LOC730100(dist=100549),MIR4431(dist=194056) | T | C | 0.00035655 | 3 | 30.3313373 | 6.62671719 | 4.71E-06 | 4.96E-06 |
| rs10187972 | 2 | 52736860 | GENOTYPED | LOC730100(dist=101805),MIR4431(dist=192800) | C | T | 0.00035655 | 3 | 30.3313373 | 6.62671719 | 4.71E-06 | 4.96E-06 |
| rs10209599 | 2 | 52739102 | GENOTYPED | LOC730100(dist=104047),MIR4431(dist=190558) | T | C | 0.00035655 | 3 | 30.3313373 | 6.62671719 | 4.71E-06 | 4.96E-06 |
| rs563784859 | 2 | 52739868 | IMPUTED | LOC730100(dist=104813),MIR4431(dist=189792) | C | T | 0.00035655 | 3 | 30.3313373 | 6.62671719 | 4.71E-06 | 4.96E-06 |
| rs546411610 | 2 | 52740102 | IMPUTED | LOC730100(dist=105047),MIR4431(dist=189558) | T | C | 0.00035655 | 3 | 30.3313373 | 6.62671719 | 4.71E-06 | 4.96E-06 |
| rs115477931 | 2 | 52740286 | IMPUTED | LOC730100(dist=105231),MIR4431(dist=189374) | G | A | 0.00035655 | 3 | 30.3313373 | 6.62671719 | 4.71E-06 | 4.96E-06 |
| rs75988214 | 2 | 52740556 | IMPUTED | LOC730100(dist=105501),MIR4431(dist=189104) | G | A | 0.00035655 | 3 | 30.3313373 | 6.62671719 | 4.71E-06 | 4.96E-06 |
| rs550751645 | 2 | 52742082 | IMPUTED | LOC730100(dist=107027),MIR4431(dist=187578) | C | T | 0.00035655 | 3 | 30.3313373 | 6.62671719 | 4.71E-06 | 4.96E-06 |
| rs569129441 | 2 | 52742087 | IMPUTED | LOC730100(dist=107032),MIR4431(dist=187573) | C | T | 0.00035655 | 3 | 30.3313373 | 6.62671719 | 4.71E-06 | 4.96E-06 |
| rs539792866 | 2 | 52742434 | IMPUTED | LOC730100(dist=107379),MIR4431(dist=187226) | A | G | 0.00035655 | 3 | 30.3313373 | 6.62671719 | 4.71E-06 | 4.96E-06 |
| rs570893929 | 2 | 52743488 | IMPUTED | LOC730100(dist=108433),MIR4431(dist=186172) | G | C | 0.00035655 | 3 | 30.3313373 | 6.62671719 | 4.71E-06 | 4.96E-06 |
| rs546114948 | 2 | 52748536 | IMPUTED | LOC730100(dist=113481),MIR4431(dist=181124) | C | T | 0.00035655 | 3 | 30.3313373 | 6.62671719 | 4.71E-06 | 4.96E-06 |
| rs527569542 | 2 | 52788118 | IMPUTED | LOC730100(dist=153063),MIR4431(dist=141542) | A | G | 0.00035655 | 3 | 30.3313373 | 6.62671719 | 4.71E-06 | 4.96E-06 |
| rs529221520 | 2 | 52790307 | IMPUTED | LOC730100(dist=155252),MIR4431(dist=139353) | T | C | 0.00035655 | 3 | 30.3313373 | 6.62671719 | 4.71E-06 | 4.96E-06 |
| rs550562911 | 2 | 52791266 | IMPUTED | LOC730100(dist=156211),MIR4431(dist=138394) | C | T | 0.00035655 | 3 | 30.3313373 | 6.62671719 | 4.71E-06 | 4.96E-06 |
| rs568967913 | 2 | 52791270 | IMPUTED | LOC730100(dist=156215),MIR4431(dist=138390) | C | A | 0.00035655 | 3 | 30.3313373 | 6.62671719 | 4.71E-06 | 4.96E-06 |
| rs537801577 | 2 | 52796544 | IMPUTED | LOC730100(dist=161489),MIR4431(dist=133116) | C | T | 0.00035655 | 3 | 30.3313373 | 6.62671719 | 4.71E-06 | 4.96E-06 |
| rs551819697 | 2 | 52813175 | IMPUTED | LOC730100(dist=178120),MIR4431(dist=116485) | T | A | 0.00035655 | 3 | 30.3313373 | 6.62671719 | 4.71E-06 | 4.96E-06 |
| rs538587569 | 2 | 52816899 | IMPUTED | LOC730100(dist=181844),MIR4431(dist=112761) | C | T | 0.00035655 | 3 | 30.3313373 | 6.62671719 | 4.71E-06 | 4.96E-06 |
| rs572048758 | 2 | 52821697 | IMPUTED | LOC730100(dist=186642),MIR4431(dist=107963) | A | C | 0.00035655 | 3 | 30.3313373 | 6.62671719 | 4.71E-06 | 4.96E-06 |
| rs554504698 | 2 | 52824896 | IMPUTED | LOC730100(dist=189841),MIR4431(dist=104764) | G | A | 0.00035655 | 3 | 30.3313373 | 6.62671719 | 4.71E-06 | 4.96E-06 |
| rs545079054 | 2 | 52827210 | IMPUTED | LOC730100(dist=192155),MIR4431(dist=102450) | T | C | 0.00035655 | 3 | 30.3313373 | 6.62671719 | 4.71E-06 | 4.96E-06 |
| rs532027353 | 2 | 52828580 | IMPUTED | LOC730100(dist=193525),MIR4431(dist=101080) | C | T | 0.00035655 | 3 | 30.3313373 | 6.62671719 | 4.71E-06 | 4.96E-06 |
| rs530399323 | 2 | 52883641 | IMPUTED | LOC730100(dist=248586),MIR4431(dist=46019) | G | C | 0.00035655 | 3 | 30.3313373 | 6.62671719 | 4.71E-06 | 4.96E-06 |
| rs113715155 | 2 | 52887694 | IMPUTED | LOC730100(dist=252639),MIR4431(dist=41966) | C | A | 0.00035655 | 3 | 30.3313373 | 6.62671719 | 4.71E-06 | 4.96E-06 |

| | | | | | | | | | | | | |
|---|---|---|---|---|---|---|---|---|---|---|---|---|
| rs372713390 | 2 | 52891535 | IMPUTED | LOC730100(dist=256480),MIR4431(dist=38125) | A | C | 0.00035655 | 3 | 30.3313373 | 6.62671719 | 4.71E-06 | 4.96E-06 |
| rs373211730 | 2 | 52902479 | IMPUTED | LOC730100(dist=267424),MIR4431(dist=27181) | A | G | 0.00035655 | 3 | 30.3313373 | 6.62671719 | 4.71E-06 | 4.96E-06 |
| rs375452509 | 2 | 52906030 | IMPUTED | LOC730100(dist=270975),MIR4431(dist=23630) | A | C | 0.00035655 | 3 | 30.3313373 | 6.62671719 | 4.71E-06 | 4.96E-06 |
| rs368001174 | 2 | 52907612 | IMPUTED | LOC730100(dist=272557),MIR4431(dist=22048) | G | A | 0.00035655 | 3 | 30.3313373 | 6.62671719 | 4.71E-06 | 4.96E-06 |
| rs374199178 | 2 | 52943891 | IMPUTED | MIR4431(dist=14138),ASB3(dist=953226) | T | A | 0.00035655 | 3 | 30.3313373 | 6.62671719 | 4.71E-06 | 4.96E-06 |
| rs376783662 | 2 | 52945859 | IMPUTED | MIR4431(dist=16106),ASB3(dist=951258) | A | C | 0.00035655 | 3 | 30.3313373 | 6.62671719 | 4.71E-06 | 4.96E-06 |
| rs367548755 | 2 | 52945977 | IMPUTED | MIR4431(dist=16224),ASB3(dist=951140) | A | G | 0.00035655 | 3 | 30.3313373 | 6.62671719 | 4.71E-06 | 4.96E-06 |
| rs567053198 | 2 | 52946641 | IMPUTED | MIR4431(dist=16888),ASB3(dist=950476) | T | A | 0.00035655 | 3 | 30.3313373 | 6.62671719 | 4.71E-06 | 4.96E-06 |
| rs376851429 | 2 | 52946746 | IMPUTED | MIR4431(dist=16993),ASB3(dist=950371) | T | C | 0.00035655 | 3 | 30.3313373 | 6.62671719 | 4.71E-06 | 4.96E-06 |
| rs369745827 | 2 | 52949008 | IMPUTED | MIR4431(dist=19255),ASB3(dist=948109) | G | C | 0.00035655 | 3 | 30.3313373 | 6.62671719 | 4.71E-06 | 4.96E-06 |
| rs373229057 | 2 | 52949073 | IMPUTED | MIR4431(dist=19320),ASB3(dist=948044) | A | T | 0.00035655 | 3 | 30.3313373 | 6.62671719 | 4.71E-06 | 4.96E-06 |
| rs556310119 | 2 | 52949752 | IMPUTED | MIR4431(dist=19999),ASB3(dist=947365) | A | G | 0.00035655 | 3 | 30.3313373 | 6.62671719 | 4.71E-06 | 4.96E-06 |
| rs377608108 | 2 | 52952455 | IMPUTED | MIR4431(dist=22702),ASB3(dist=944662) | A | G | 0.00035655 | 3 | 30.3313373 | 6.62671719 | 4.71E-06 | 4.96E-06 |
| rs371103615 | 2 | 52952570 | IMPUTED | MIR4431(dist=22817),ASB3(dist=944547) | G | T | 0.00035655 | 3 | 30.3313373 | 6.62671719 | 4.71E-06 | 4.96E-06 |
| rs371111972 | 2 | 52970008 | IMPUTED | MIR4431(dist=40255),ASB3(dist=927109) | T | C | 0.00035655 | 3 | 30.3313373 | 6.62671719 | 4.71E-06 | 4.96E-06 |
| rs369173152 | 2 | 52974817 | IMPUTED | MIR4431(dist=45064),ASB3(dist=922300) | G | A | 0.00035655 | 3 | 30.3313373 | 6.62671719 | 4.71E-06 | 4.96E-06 |
| rs374048736 | 2 | 52975352 | IMPUTED | MIR4431(dist=45599),ASB3(dist=921765) | T | C | 0.00035655 | 3 | 30.3313373 | 6.62671719 | 4.71E-06 | 4.96E-06 |
| rs376844753 | 2 | 52980403 | IMPUTED | MIR4431(dist=50650),ASB3(dist=916714) | C | T | 0.00035655 | 3 | 30.3313373 | 6.62671719 | 4.71E-06 | 4.96E-06 |
| rs372848560 | 2 | 52980866 | IMPUTED | MIR4431(dist=51113),ASB3(dist=916251) | A | C | 0.00035655 | 3 | 30.3313373 | 6.62671719 | 4.71E-06 | 4.96E-06 |
| rs185725044 | 2 | 52986949 | IMPUTED | MIR4431(dist=57196),ASB3(dist=910168) | C | A | 0.00035655 | 3 | 30.3313373 | 6.62671719 | 4.71E-06 | 4.96E-06 |
| rs373907362 | 2 | 52987346 | IMPUTED | MIR4431(dist=57593),ASB3(dist=909771) | C | T | 0.00035655 | 3 | 30.3313373 | 6.62671719 | 4.71E-06 | 4.96E-06 |
| rs370340869 | 2 | 52988086 | IMPUTED | MIR4431(dist=58333),ASB3(dist=909031) | A | G | 0.00035655 | 3 | 30.3313373 | 6.62671719 | 4.71E-06 | 4.96E-06 |
| rs377613717 | 2 | 52989589 | IMPUTED | MIR4431(dist=59836),ASB3(dist=907528) | T | C | 0.00035655 | 3 | 30.3313373 | 6.62671719 | 4.71E-06 | 4.96E-06 |
| rs371746508 | 2 | 52993493 | IMPUTED | MIR4431(dist=63740),ASB3(dist=903624) | G | A | 0.00035655 | 3 | 30.3313373 | 6.62671719 | 4.71E-06 | 4.96E-06 |
| rs4263147 | 2 | 52996436 | IMPUTED | MIR4431(dist=66683),ASB3(dist=900681) | G | A | 0.00035655 | 3 | 30.3313373 | 6.62671719 | 4.71E-06 | 4.96E-06 |
| rs374957640 | 2 | 52997217 | IMPUTED | MIR4431(dist=67464),ASB3(dist=899900) | A | G | 0.00035655 | 3 | 30.3313373 | 6.62671719 | 4.71E-06 | 4.96E-06 |
| rs7584468 | 2 | 52998684 | IMPUTED | MIR4431(dist=68931),ASB3(dist=898433) | A | G | 0.00035655 | 3 | 30.3313373 | 6.62671719 | 4.71E-06 | 4.96E-06 |
| rs370552692 | 2 | 52999439 | IMPUTED | MIR4431(dist=69686),ASB3(dist=897678) | T | G | 0.00035655 | 3 | 30.3313373 | 6.62671719 | 4.71E-06 | 4.96E-06 |
| rs578038220 | 2 | 53000477 | IMPUTED | MIR4431(dist=70724),ASB3(dist=896640) | G | A | 0.00035655 | 3 | 30.3313373 | 6.62671719 | 4.71E-06 | 4.96E-06 |
| rs375008586 | 2 | 53003027 | IMPUTED | MIR4431(dist=73274),ASB3(dist=894090) | T | C | 0.00035655 | 3 | 30.3313373 | 6.62671719 | 4.71E-06 | 4.96E-06 |
| rs376418644 | 2 | 53009954 | IMPUTED | MIR4431(dist=80201),ASB3(dist=887163) | G | C | 0.00035655 | 3 | 30.3313373 | 6.62671719 | 4.71E-06 | 4.96E-06 |
| rs532066277 | 2 | 53012669 | IMPUTED | MIR4431(dist=82916),ASB3(dist=884448) | G | T | 0.00035655 | 3 | 30.3313373 | 6.62671719 | 4.71E-06 | 4.96E-06 |
| rs535999858 | 2 | 53012670 | IMPUTED | MIR4431(dist=82917),ASB3(dist=884447) | C | T | 0.00035655 | 3 | 30.3313373 | 6.62671719 | 4.71E-06 | 4.96E-06 |
| rs372024582 | 2 | 53015305 | IMPUTED | MIR4431(dist=85552),ASB3(dist=881812) | T | C | 0.00035655 | 3 | 30.3313373 | 6.62671719 | 4.71E-06 | 4.96E-06 |
| rs375216538 | 2 | 53015379 | IMPUTED | MIR4431(dist=85626),ASB3(dist=881738) | A | C | 0.00035655 | 3 | 30.3313373 | 6.62671719 | 4.71E-06 | 4.96E-06 |
| rs376159637 | 2 | 53016909 | IMPUTED | MIR4431(dist=87156),ASB3(dist=880208) | A | G | 0.00035655 | 3 | 30.3313373 | 6.62671719 | 4.71E-06 | 4.96E-06 |
| rs374125682 | 2 | 53018752 | IMPUTED | MIR4431(dist=88999),ASB3(dist=878365) | C | G | 0.00035655 | 3 | 30.3313373 | 6.62671719 | 4.71E-06 | 4.96E-06 |
| rs375720110 | 2 | 53018789 | IMPUTED | MIR4431(dist=89036),ASB3(dist=878328) | C | T | 0.00035655 | 3 | 30.3313373 | 6.62671719 | 4.71E-06 | 4.96E-06 |
| rs146387452 | 2 | 53019922 | IMPUTED | MIR4431(dist=90169),ASB3(dist=877195) | C | T | 0.00035655 | 3 | 30.3313373 | 6.62671719 | 4.71E-06 | 4.96E-06 |
| rs371168552 | 2 | 53022057 | IMPUTED | MIR4431(dist=92304),ASB3(dist=875060) | C | T | 0.00035655 | 3 | 30.3313373 | 6.62671719 | 4.71E-06 | 4.96E-06 |
| rs376614829 | 2 | 53022790 | IMPUTED | MIR4431(dist=93037),ASB3(dist=874327) | G | A | 0.00035655 | 3 | 30.3313373 | 6.62671719 | 4.71E-06 | 4.96E-06 |
| rs375027288 | 2 | 53023367 | IMPUTED | MIR4431(dist=93614),ASB3(dist=873750) | T | C | 0.00035655 | 3 | 30.3313373 | 6.62671719 | 4.71E-06 | 4.96E-06 |
| rs372448654 | 2 | 53023769 | IMPUTED | MIR4431(dist=94016),ASB3(dist=873348) | C | T | 0.00035655 | 3 | 30.3313373 | 6.62671719 | 4.71E-06 | 4.96E-06 |
| rs368385681 | 2 | 53024808 | IMPUTED | MIR4431(dist=95055),ASB3(dist=872309) | T | G | 0.00035655 | 3 | 30.3313373 | 6.62671719 | 4.71E-06 | 4.96E-06 |
| rs374720584 | 2 | 53025662 | IMPUTED | MIR4431(dist=95909),ASB3(dist=871455) | A | G | 0.00035655 | 3 | 30.3313373 | 6.62671719 | 4.71E-06 | 4.96E-06 |
| rs368579989 | 2 | 53027201 | IMPUTED | MIR4431(dist=97448),ASB3(dist=869916) | T | C | 0.00035655 | 3 | 30.3313373 | 6.62671719 | 4.71E-06 | 4.96E-06 |
| rs376974781 | 2 | 53045609 | IMPUTED | MIR4431(dist=115856),ASB3(dist=851508) | A | G | 0.00035655 | 3 | 30.3313373 | 6.62671719 | 4.71E-06 | 4.96E-06 |
| rs1451461 | 2 | 53045987 | IMPUTED | MIR4431(dist=116234),ASB3(dist=851130) | T | C | 0.00035655 | 3 | 30.3313373 | 6.62671719 | 4.71E-06 | 4.96E-06 |
| rs377305018 | 2 | 53049078 | IMPUTED | MIR4431(dist=119325),ASB3(dist=848039) | G | A | 0.00035655 | 3 | 30.3313373 | 6.62671719 | 4.71E-06 | 4.96E-06 |
| rs372780258 | 2 | 53051083 | IMPUTED | MIR4431(dist=121330),ASB3(dist=846034) | G | C | 0.00035655 | 3 | 30.3313373 | 6.62671719 | 4.71E-06 | 4.96E-06 |
| rs374393210 | 2 | 53052556 | IMPUTED | MIR4431(dist=122803),ASB3(dist=844561) | C | A | 0.00035655 | 3 | 30.3313373 | 6.62671719 | 4.71E-06 | 4.96E-06 |
| rs572221454 | 2 | 53053088 | IMPUTED | MIR4431(dist=123335),ASB3(dist=844029) | A | G | 0.00035655 | 3 | 30.3313373 | 6.62671719 | 4.71E-06 | 4.96E-06 |

| | | | | | | | | | | | | |
|---|---|---|---|---|---|---|---|---|---|---|---|---|
| rs374596773 | 2 | 53054512 | IMPUTED | MIR4431(dist=124759),ASB3(dist=842605) | T | A | 0.00035655 | 3 | 30.3313373 | 6.62671719 | 4.71E-06 | 4.96E-06 |
| rs370658242 | 2 | 53057573 | IMPUTED | MIR4431(dist=127820),ASB3(dist=839544) | T | C | 0.00035655 | 3 | 30.3313373 | 6.62671719 | 4.71E-06 | 4.96E-06 |
| rs368423498 | 2 | 53067509 | IMPUTED | MIR4431(dist=137756),ASB3(dist=829608) | T | C | 0.00035655 | 3 | 30.3313373 | 6.62671719 | 4.71E-06 | 4.96E-06 |
| rs370673490 | 2 | 53069385 | IMPUTED | MIR4431(dist=139632),ASB3(dist=827732) | A | T | 0.00035655 | 3 | 30.3313373 | 6.62671719 | 4.71E-06 | 4.96E-06 |
| rs372673144 | 2 | 53070194 | IMPUTED | MIR4431(dist=140441),ASB3(dist=826923) | C | T | 0.00035655 | 3 | 30.3313373 | 6.62671719 | 4.71E-06 | 4.96E-06 |
| rs368886465 | 2 | 53070352 | IMPUTED | MIR4431(dist=140599),ASB3(dist=826765) | A | G | 0.00035655 | 3 | 30.3313373 | 6.62671719 | 4.71E-06 | 4.96E-06 |
| rs368684152 | 2 | 53074254 | IMPUTED | MIR4431(dist=144501),ASB3(dist=822863) | A | G | 0.00035655 | 3 | 30.3313373 | 6.62671719 | 4.71E-06 | 4.96E-06 |
| rs367793883 | 2 | 53074634 | IMPUTED | MIR4431(dist=144881),ASB3(dist=822483) | A | C | 0.00035655 | 3 | 30.3313373 | 6.62671719 | 4.71E-06 | 4.96E-06 |
| rs2122834 | 2 | 53076508 | IMPUTED | MIR4431(dist=146755),ASB3(dist=820609) | G | C | 0.00035655 | 3 | 30.3313373 | 6.62671719 | 4.71E-06 | 4.96E-06 |
| rs374828848 | 2 | 53078963 | IMPUTED | MIR4431(dist=149210),ASB3(dist=818154) | G | A | 0.00035655 | 3 | 30.3313373 | 6.62671719 | 4.71E-06 | 4.96E-06 |
| rs372283813 | 2 | 53079813 | IMPUTED | MIR4431(dist=150060),ASB3(dist=817304) | C | T | 0.00035655 | 3 | 30.3313373 | 6.62671719 | 4.71E-06 | 4.96E-06 |
| rs370070279 | 2 | 53083856 | IMPUTED | MIR4431(dist=154103),ASB3(dist=813261) | A | G | 0.00035655 | 3 | 30.3313373 | 6.62671719 | 4.71E-06 | 4.96E-06 |
| rs187031480 | 2 | 53084831 | IMPUTED | MIR4431(dist=155078),ASB3(dist=812286) | C | G | 0.00035655 | 3 | 30.3313373 | 6.62671719 | 4.71E-06 | 4.96E-06 |
| rs374699037 | 2 | 53085539 | IMPUTED | MIR4431(dist=155786),ASB3(dist=811578) | A | T | 0.00035655 | 3 | 30.3313373 | 6.62671719 | 4.71E-06 | 4.96E-06 |
| rs371759740 | 2 | 53086199 | IMPUTED | MIR4431(dist=156446),ASB3(dist=810918) | G | A | 0.00035655 | 3 | 30.3313373 | 6.62671719 | 4.71E-06 | 4.96E-06 |
| rs367818531 | 2 | 53086329 | IMPUTED | MIR4431(dist=156576),ASB3(dist=810788) | T | A | 0.00035655 | 3 | 30.3313373 | 6.62671719 | 4.71E-06 | 4.96E-06 |
| rs184160873 | 2 | 53090179 | IMPUTED | MIR4431(dist=160426),ASB3(dist=806938) | C | T | 0.00035655 | 3 | 30.3313373 | 6.62671719 | 4.71E-06 | 4.96E-06 |
| rs138517075 | 2 | 53090960 | IMPUTED | MIR4431(dist=161207),ASB3(dist=806157) | T | G | 0.00035655 | 3 | 30.3313373 | 6.62671719 | 4.71E-06 | 4.96E-06 |
| rs376352658 | 2 | 53092151 | IMPUTED | MIR4431(dist=162398),ASB3(dist=804966) | G | C | 0.00035655 | 3 | 30.3313373 | 6.62671719 | 4.71E-06 | 4.96E-06 |
| rs375953088 | 2 | 53093169 | IMPUTED | MIR4431(dist=163416),ASB3(dist=803948) | T | G | 0.00035655 | 3 | 30.3313373 | 6.62671719 | 4.71E-06 | 4.96E-06 |
| rs368040583 | 2 | 53093425 | IMPUTED | MIR4431(dist=163672),ASB3(dist=803692) | C | T | 0.00035655 | 3 | 30.3313373 | 6.62671719 | 4.71E-06 | 4.96E-06 |
| rs190919296 | 2 | 53093463 | IMPUTED | MIR4431(dist=163710),ASB3(dist=803654) | A | G | 0.00035655 | 3 | 30.3313373 | 6.62671719 | 4.71E-06 | 4.96E-06 |
| rs182625218 | 2 | 53093767 | IMPUTED | MIR4431(dist=164014),ASB3(dist=803350) | C | T | 0.00035655 | 3 | 30.3313373 | 6.62671719 | 4.71E-06 | 4.96E-06 |
| rs374272665 | 2 | 53095847 | IMPUTED | MIR4431(dist=166094),ASB3(dist=801270) | A | G | 0.00035655 | 3 | 30.3313373 | 6.62671719 | 4.71E-06 | 4.96E-06 |
| rs371176997 | 2 | 53096275 | IMPUTED | MIR4431(dist=166522),ASB3(dist=800842) | C | A | 0.00035655 | 3 | 30.3313373 | 6.62671719 | 4.71E-06 | 4.96E-06 |
| rs375647791 | 2 | 53097666 | IMPUTED | MIR4431(dist=167913),ASB3(dist=799451) | T | A | 0.00035655 | 3 | 30.3313373 | 6.62671719 | 4.71E-06 | 4.96E-06 |
| rs367901292 | 2 | 53097825 | IMPUTED | MIR4431(dist=168072),ASB3(dist=799292) | G | C | 0.00035655 | 3 | 30.3313373 | 6.62671719 | 4.71E-06 | 4.96E-06 |
| rs140739207 | 2 | 53098046 | IMPUTED | MIR4431(dist=168293),ASB3(dist=799071) | C | T | 0.00035655 | 3 | 30.3313373 | 6.62671719 | 4.71E-06 | 4.96E-06 |
| rs1376566 | 2 | 53098542 | IMPUTED | MIR4431(dist=168789),ASB3(dist=798575) | A | C | 0.00035655 | 3 | 30.3313373 | 6.62671719 | 4.71E-06 | 4.96E-06 |
| rs552594509 | 2 | 53099558 | IMPUTED | MIR4431(dist=169805),ASB3(dist=797559) | A | T | 0.00035655 | 3 | 30.3313373 | 6.62671719 | 4.71E-06 | 4.96E-06 |
| rs374268300 | 2 | 53100764 | IMPUTED | MIR4431(dist=171011),ASB3(dist=796353) | T | C | 0.00035655 | 3 | 30.3313373 | 6.62671719 | 4.71E-06 | 4.96E-06 |
| rs377035611 | 2 | 53101538 | IMPUTED | MIR4431(dist=171785),ASB3(dist=795579) | T | C | 0.00035655 | 3 | 30.3313373 | 6.62671719 | 4.71E-06 | 4.96E-06 |
| rs369835291 | 2 | 53102987 | IMPUTED | MIR4431(dist=173234),ASB3(dist=794130) | G | C | 0.00035655 | 3 | 30.3313373 | 6.62671719 | 4.71E-06 | 4.96E-06 |
| rs1868906 | 2 | 53105047 | IMPUTED | MIR4431(dist=175294),ASB3(dist=792070) | T | C | 0.00035655 | 3 | 30.3313373 | 6.62671719 | 4.71E-06 | 4.96E-06 |
| rs368428877 | 2 | 53105628 | IMPUTED | MIR4431(dist=175875),ASB3(dist=791489) | G | A | 0.00035655 | 3 | 30.3313373 | 6.62671719 | 4.71E-06 | 4.96E-06 |
| rs376861736 | 2 | 53107240 | IMPUTED | MIR4431(dist=177487),ASB3(dist=789877) | C | G | 0.00035655 | 3 | 30.3313373 | 6.62671719 | 4.71E-06 | 4.96E-06 |
| rs369494773 | 2 | 53107659 | IMPUTED | MIR4431(dist=177906),ASB3(dist=789458) | C | G | 0.00035655 | 3 | 30.3313373 | 6.62671719 | 4.71E-06 | 4.96E-06 |
| rs377436118 | 2 | 53107705 | IMPUTED | MIR4431(dist=177952),ASB3(dist=789412) | C | T | 0.00035655 | 3 | 30.3313373 | 6.62671719 | 4.71E-06 | 4.96E-06 |
| rs1864533 | 2 | 53109379 | IMPUTED | MIR4431(dist=179626),ASB3(dist=787738) | A | T | 0.00035655 | 3 | 30.3313373 | 6.62671719 | 4.71E-06 | 4.96E-06 |
| rs369471378 | 2 | 53112597 | IMPUTED | MIR4431(dist=182844),ASB3(dist=784520) | G | A | 0.00035655 | 3 | 30.3313373 | 6.62671719 | 4.71E-06 | 4.96E-06 |
| rs370668232 | 2 | 53114029 | IMPUTED | MIR4431(dist=184276),ASB3(dist=783088) | A | G | 0.00035655 | 3 | 30.3313373 | 6.62671719 | 4.71E-06 | 4.96E-06 |
| rs374881775 | 2 | 53119483 | IMPUTED | MIR4431(dist=189730),ASB3(dist=777634) | C | T | 0.00035655 | 3 | 30.3313373 | 6.62671719 | 4.71E-06 | 4.96E-06 |
| rs557719687 | 2 | 53147235 | IMPUTED | MIR4431(dist=217482),ASB3(dist=749882) | C | T | 0.00035655 | 3 | 30.3313373 | 6.62671719 | 4.71E-06 | 4.96E-06 |
| rs374175094 | 12 | 60058530 | IMPUTED | SLC16A7 | C | T | 0.00035655 | 3 | 30.3313373 | 6.62671719 | 4.71E-06 | 4.96E-06 |
| rs150258437 | 12 | 60087460 | IMPUTED | SLC16A7 | T | C | 0.00035655 | 3 | 30.3313373 | 6.62671719 | 4.71E-06 | 4.96E-06 |
| rs138452335 | 12 | 60091008 | IMPUTED | SLC16A7 | G | A | 0.00035655 | 3 | 30.3313373 | 6.62671719 | 4.71E-06 | 4.96E-06 |
| rs151027047 | 12 | 60108470 | IMPUTED | SLC16A7 | C | T | 0.00035655 | 3 | 30.3313373 | 6.62671719 | 4.71E-06 | 4.96E-06 |
| rs11173127 | 12 | 60133321 | IMPUTED | SLC16A7 | G | A | 0.00035655 | 3 | 30.3313373 | 6.62671719 | 4.71E-06 | 4.96E-06 |
| rs767930336 | 12 | 60140721 | IMPUTED | SLC16A7 | C | T | 0.00035655 | 3 | 30.3313373 | 6.62671719 | 4.71E-06 | 4.96E-06 |
| rs12303722 | 12 | 60144427 | IMPUTED | SLC16A7 | G | A | 0.00035655 | 3 | 30.3313373 | 6.62671719 | 4.71E-06 | 4.96E-06 |
| rs189827842 | 12 | 60145004 | IMPUTED | SLC16A7 | A | G | 0.00035655 | 3 | 30.3313373 | 6.62671719 | 4.71E-06 | 4.96E-06 |
| rs74096452 | 12 | 60164585 | IMPUTED | SLC16A7 | A | G | 0.00035655 | 3 | 30.3313373 | 6.62671719 | 4.71E-06 | 4.96E-06 |

| | | | | | | | | | | | | |
|---|---|---|---|---|---|---|---|---|---|---|---|---|
| rs74096453 | 12 | 60164673 | IMPUTED | SLC16A7 | C | T | 0.00035655 | 3 | 30.3313373 | 6.62671719 | 4.71E-06 | 4.96E-06 |
| rs13378061 | 12 | 60169806 | IMPUTED | SLC16A7 | T | A | 0.00035655 | 3 | 30.3313373 | 6.62671719 | 4.71E-06 | 4.96E-06 |
| rs7316320 | 12 | 60171528 | IMPUTED | SLC16A7 | T | C | 0.00035655 | 3 | 30.3313373 | 6.62671719 | 4.71E-06 | 4.96E-06 |
| rs11173140 | 12 | 60173052 | IMPUTED | SLC16A7 | A | T | 0.00035655 | 3 | 30.3313373 | 6.62671719 | 4.71E-06 | 4.96E-06 |
| rs11173141 | 12 | 60173806 | GENOTYPED | SLC16A7(NM_001270623:c.*3460>0,NM_001270622:c.*34 | A | G | 0.00035655 | 3 | 30.3313373 | 6.62671719 | 4.71E-06 | 4.96E-06 |
| rs11173142 | 12 | 60173878 | GENOTYPED | SLC16A7(NM_001270623:c.*4180>0,NM_001270622:c.*41 | A | C | 0.00035655 | 3 | 30.3313373 | 6.62671719 | 4.71E-06 | 4.96E-06 |
| rs149148185 | 12 | 60174087 | IMPUTED | SLC16A7(NM_001270623:c.*6270>0,NM_001270622:c.*62 | A | T | 0.00035655 | 3 | 30.3313373 | 6.62671719 | 4.71E-06 | 4.96E-06 |
| rs11173144 | 12 | 60176349 | IMPUTED | SLC16A7(NM_001270623:c.*28890>0,NM_001270622:c.*2 | A | T | 0.00035655 | 3 | 30.3313373 | 6.62671719 | 4.71E-06 | 4.96E-06 |
| rs11173145 | 12 | 60176474 | IMPUTED | SLC16A7(NM_001270623:c.*30140>0,NM_001270622:c.*3 | C | T | 0.00035655 | 3 | 30.3313373 | 6.62671719 | 4.71E-06 | 4.96E-06 |
| rs11173146 | 12 | 60176576 | IMPUTED | SLC16A7(NM_001270623:c.*31160>0,NM_001270622:c.*3 | C | A | 0.00035655 | 3 | 30.3313373 | 6.62671719 | 4.71E-06 | 4.96E-06 |
| rs12316816 | 12 | 60177483 | IMPUTED | SLC16A7(NM_001270623:c.*40230>0,NM_001270622:c.*4 | C | A | 0.00035655 | 3 | 30.3313373 | 6.62671719 | 4.71E-06 | 4.96E-06 |
| rs12303301 | 12 | 60177494 | IMPUTED | SLC16A7(NM_001270623:c.*40340>0,NM_001270622:c.*4 | A | G | 0.00035655 | 3 | 30.3313373 | 6.62671719 | 4.71E-06 | 4.96E-06 |
| rs7307407 | 12 | 60178234 | IMPUTED | SLC16A7(NM_001270623:c.*47740>0,NM_001270622:c.*4 | G | C | 0.00035655 | 3 | 30.3313373 | 6.62671719 | 4.71E-06 | 4.96E-06 |
| rs7307832 | 12 | 60178589 | IMPUTED | SLC16A7(NM_001270623:c.*51290>0,NM_001270622:c.*5 | G | C | 0.00035655 | 3 | 30.3313373 | 6.62671719 | 4.71E-06 | 4.96E-06 |
| rs7310780 | 12 | 60178801 | IMPUTED | SLC16A7(NM_001270623:c.*53410>0,NM_001270622:c.*5 | A | G | 0.00035655 | 3 | 30.3313373 | 6.62671719 | 4.71E-06 | 4.96E-06 |
| rs7311237 | 12 | 60178910 | IMPUTED | SLC16A7(NM_001270623:c.*54500>0,NM_001270622:c.*5 | G | C | 0.00035655 | 3 | 30.3313373 | 6.62671719 | 4.71E-06 | 4.96E-06 |
| rs28701946 | 12 | 60179814 | IMPUTED | SLC16A7(NM_001270623:c.*63540>0,NM_001270622:c.*6 | G | T | 0.00035655 | 3 | 30.3313373 | 6.62671719 | 4.71E-06 | 4.96E-06 |
| rs28534781 | 12 | 60179847 | IMPUTED | SLC16A7(NM_001270623:c.*63870>0,NM_001270622:c.*6 | G | A | 0.00035655 | 3 | 30.3313373 | 6.62671719 | 4.71E-06 | 4.96E-06 |
| rs12304208 | 12 | 60180842 | IMPUTED | SLC16A7(NM_001270623:c.*73820>0,NM_001270622:c.*7 | T | C | 0.00035655 | 3 | 30.3313373 | 6.62671719 | 4.71E-06 | 4.96E-06 |
| rs12310755 | 12 | 60181280 | IMPUTED | SLC16A7(NM_001270623:c.*78200>0,NM_001270622:c.*7 | A | G | 0.00035655 | 3 | 30.3313373 | 6.62671719 | 4.71E-06 | 4.96E-06 |
| rs12304515 | 12 | 60181337 | IMPUTED | SLC16A7(NM_001270623:c.*78770>0,NM_001270622:c.*7 | T | A | 0.00035655 | 3 | 30.3313373 | 6.62671719 | 4.71E-06 | 4.96E-06 |
| rs12306387 | 12 | 60182244 | IMPUTED | SLC16A7(NM_001270623:c.*87840>0,NM_001270622:c.*8 | T | C | 0.00035655 | 3 | 30.3313373 | 6.62671719 | 4.71E-06 | 4.96E-06 |
| rs74095498 | 12 | 60182681 | IMPUTED | SLC16A7(NM_001270623:c.*92210>0,NM_001270622:c.*9 | A | C | 0.00035655 | 3 | 30.3313373 | 6.62671719 | 4.71E-06 | 4.96E-06 |
| rs74095500 | 12 | 60182765 | IMPUTED | SLC16A7(NM_001270623:c.*93050>0,NM_001270622:c.*9 | T | C | 0.00035655 | 3 | 30.3313373 | 6.62671719 | 4.71E-06 | 4.96E-06 |
| rs74095501 | 12 | 60182906 | IMPUTED | SLC16A7(NM_001270623:c.*94460>0,NM_001270622:c.*9 | A | G | 0.00035655 | 3 | 30.3313373 | 6.62671719 | 4.71E-06 | 4.96E-06 |
| rs56121641 | 12 | 60183692 | IMPUTED | SLC16A7(dist=57) | A | G | 0.00035655 | 3 | 30.3313373 | 6.62671719 | 4.71E-06 | 4.96E-06 |
| rs10161078 | 12 | 60185335 | IMPUTED | SLC16A7(dist=1700),FAM19A2(dist=1916694) | C | T | 0.00035655 | 3 | 30.3313373 | 6.62671719 | 4.71E-06 | 4.96E-06 |
| rs10160899 | 12 | 60186828 | IMPUTED | SLC16A7(dist=3193),FAM19A2(dist=1915201) | C | T | 0.00035655 | 3 | 30.3313373 | 6.62671719 | 4.71E-06 | 4.96E-06 |
| rs10161101 | 12 | 60187174 | IMPUTED | SLC16A7(dist=3539),FAM19A2(dist=1914855) | T | G | 0.00035655 | 3 | 30.3313373 | 6.62671719 | 4.71E-06 | 4.96E-06 |
| rs12300901 | 12 | 60188479 | IMPUTED | SLC16A7(dist=4844),FAM19A2(dist=1913550) | G | C | 0.00035655 | 3 | 30.3313373 | 6.62671719 | 4.71E-06 | 4.96E-06 |
| rs11173154 | 12 | 60189332 | IMPUTED | SLC16A7(dist=5697),FAM19A2(dist=1912697) | G | A | 0.00035655 | 3 | 30.3313373 | 6.62671719 | 4.71E-06 | 4.96E-06 |
| rs11504057 | 12 | 60189610 | IMPUTED | SLC16A7(dist=5975),FAM19A2(dist=1912419) | C | T | 0.00035655 | 3 | 30.3313373 | 6.62671719 | 4.71E-06 | 4.96E-06 |
| rs78279772 | 12 | 60192365 | IMPUTED | SLC16A7(dist=8730),FAM19A2(dist=1909664) | A | G | 0.00035655 | 3 | 30.3313373 | 6.62671719 | 4.71E-06 | 4.96E-06 |
| rs12298827 | 12 | 60194678 | IMPUTED | SLC16A7(dist=11043),FAM19A2(dist=1907351) | A | G | 0.00035655 | 3 | 30.3313373 | 6.62671719 | 4.71E-06 | 4.96E-06 |
| rs12296304 | 12 | 60196574 | IMPUTED | SLC16A7(dist=12939),FAM19A2(dist=1905455) | T | C | 0.00035655 | 3 | 30.3313373 | 6.62671719 | 4.71E-06 | 4.96E-06 |
| rs12318590 | 12 | 60199283 | IMPUTED | SLC16A7(dist=15648),FAM19A2(dist=1902746) | T | C | 0.00035655 | 3 | 30.3313373 | 6.62671719 | 4.71E-06 | 4.96E-06 |
| rs12318639 | 12 | 60199337 | IMPUTED | SLC16A7(dist=15702),FAM19A2(dist=1902692) | T | C | 0.00035655 | 3 | 30.3313373 | 6.62671719 | 4.71E-06 | 4.96E-06 |
| rs12301821 | 12 | 60201300 | IMPUTED | SLC16A7(dist=17665),FAM19A2(dist=1900729) | A | G | 0.00035655 | 3 | 30.3313373 | 6.62671719 | 4.71E-06 | 4.96E-06 |
| rs10437931 | 12 | 60201882 | IMPUTED | SLC16A7(dist=18247),FAM19A2(dist=1900147) | G | A | 0.00035655 | 3 | 30.3313373 | 6.62671719 | 4.71E-06 | 4.96E-06 |
| rs11173161 | 12 | 60205072 | GENOTYPED | SLC16A7(dist=21437),FAM19A2(dist=1896957) | A | G | 0.00035655 | 3 | 30.3313373 | 6.62671719 | 4.71E-06 | 4.96E-06 |
| rs115609179 | 12 | 60222966 | IMPUTED | SLC16A7(dist=39331),FAM19A2(dist=1879063) | C | G | 0.00035655 | 3 | 30.3313373 | 6.62671719 | 4.71E-06 | 4.96E-06 |
| rs74095777 | 12 | 60227712 | IMPUTED | SLC16A7(dist=44077),FAM19A2(dist=1874317) | G | C | 0.00035655 | 3 | 30.3313373 | 6.62671719 | 4.71E-06 | 4.96E-06 |
| rs74095781 | 12 | 60229512 | IMPUTED | SLC16A7(dist=45877),FAM19A2(dist=1872517) | C | T | 0.00035655 | 3 | 30.3313373 | 6.62671719 | 4.71E-06 | 4.96E-06 |
| rs74095794 | 12 | 60237301 | GENOTYPED | SLC16A7(dist=53666),FAM19A2(dist=1864728) | T | C | 0.00035655 | 3 | 30.3313373 | 6.62671719 | 4.71E-06 | 4.96E-06 |
| rs865864711 | 12 | 60645607 | IMPUTED | SLC16A7(dist=461972),FAM19A2(dist=1456422) | T | C | 0.00035655 | 3 | 30.3313373 | 6.62671719 | 4.71E-06 | 4.96E-06 |
| rs748926027 | 9 | 120716095 | IMPUTED | TLR4(dist=236326),BRINP1(dist=1212813) | C | A | 0.00035512 | 3 | 30.4531502 | 6.65333056 | 4.71E-06 | 4.96E-06 |
| rs116802691 | 2 | 52985351 | IMPUTED | MIR4431(dist=55598),ASB3(dist=911766) | A | G | 0.00035667 | 3 | 30.3310373 | 6.62671057 | 4.71E-06 | 4.96E-06 |
| rs375636911 | 2 | 52912902 | IMPUTED | LOC730100(dist=277847),MIR4431(dist=16758) | G | T | 0.00035714 | 3 | 30.3309445 | 6.62669753 | 4.72E-06 | 4.96E-06 |
| rs373775879 | 2 | 52732897 | IMPUTED | LOC730100(dist=97842),MIR4431(dist=196763) | C | A | 0.00035667 | 3 | 30.3304595 | 6.62671695 | 4.72E-06 | 4.96E-06 |
| rs376126118 | 2 | 52733140 | IMPUTED | LOC730100(dist=98085),MIR4431(dist=196520) | C | T | 0.00035667 | 3 | 30.3304595 | 6.62671695 | 4.72E-06 | 4.96E-06 |
| rs544170814 | 2 | 52733150 | IMPUTED | LOC730100(dist=98095),MIR4431(dist=196510) | C | G | 0.00035667 | 3 | 30.3304595 | 6.62671695 | 4.72E-06 | 4.96E-06 |
| rs374849837 | 2 | 52733210 | IMPUTED | LOC730100(dist=98155),MIR4431(dist=196450) | A | G | 0.00035667 | 3 | 30.3304595 | 6.62671695 | 4.72E-06 | 4.96E-06 |

| | | | | | | | | | | | | |
|---|---|---|---|---|---|---|---|---|---|---|---|---|
| rs185117219 | 2 | 52733514 | IMPUTED | LOC730100(dist=98459),MIR4431(dist=196146) | A | G | 0.00035667 | 3 | 30.3304595 | 6.62671695 | 4.72E-06 | 4.96E-06 |
| rs191491025 | 2 | 52733708 | IMPUTED | LOC730100(dist=98653),MIR4431(dist=195952) | C | T | 0.00035667 | 3 | 30.3304595 | 6.62671695 | 4.72E-06 | 4.96E-06 |
| rs183840040 | 2 | 52733748 | IMPUTED | LOC730100(dist=98693),MIR4431(dist=195912) | A | G | 0.00035667 | 3 | 30.3304595 | 6.62671695 | 4.72E-06 | 4.96E-06 |
| rs376687316 | 2 | 52733882 | IMPUTED | LOC730100(dist=98827),MIR4431(dist=195778) | C | T | 0.00035667 | 3 | 30.3304595 | 6.62671695 | 4.72E-06 | 4.96E-06 |
| rs182110122 | 2 | 52733968 | IMPUTED | LOC730100(dist=98913),MIR4431(dist=195692) | G | A | 0.00035667 | 3 | 30.3304595 | 6.62671695 | 4.72E-06 | 4.96E-06 |
| rs193221137 | 2 | 52734017 | IMPUTED | LOC730100(dist=98962),MIR4431(dist=195643) | T | C | 0.00035667 | 3 | 30.3304595 | 6.62671695 | 4.72E-06 | 4.96E-06 |
| rs377145324 | 2 | 52734248 | IMPUTED | LOC730100(dist=99193),MIR4431(dist=195412) | C | T | 0.00035667 | 3 | 30.3304595 | 6.62671695 | 4.72E-06 | 4.96E-06 |
| rs190751367 | 2 | 52734316 | IMPUTED | LOC730100(dist=99261),MIR4431(dist=195344) | G | A | 0.00035667 | 3 | 30.3304595 | 6.62671695 | 4.72E-06 | 4.96E-06 |
| rs530397536 | 2 | 52734395 | IMPUTED | LOC730100(dist=99340),MIR4431(dist=195265) | A | C | 0.00035667 | 3 | 30.3304595 | 6.62671695 | 4.72E-06 | 4.96E-06 |
| rs374373550 | 2 | 52735159 | IMPUTED | LOC730100(dist=100104),MIR4431(dist=194501) | C | T | 0.00035667 | 3 | 30.3304595 | 6.62671695 | 4.72E-06 | 4.96E-06 |
| rs552186163 | 2 | 55744929 | IMPUTED | CCDC88A(dist=97872),CFAP36(dist=1802) | T | C | 0.00035584 | 3 | 30.4442775 | 6.6515885 | 4.72E-06 | 4.96E-06 |
| rs554319096 | 2 | 52835517 | IMPUTED | LOC730100(dist=200462),MIR4431(dist=94143) | T | A | 0.00035691 | 3 | 30.3300558 | 6.62669591 | 4.72E-06 | 4.97E-06 |
| rs181035023 | 12 | 60083836 | IMPUTED | SLC16A7 | A | C | 0.00035893 | 3 | 30.3290844 | 6.62650103 | 4.72E-06 | 4.97E-06 |
| rs376638129 | 2 | 53034115 | IMPUTED | MIR4431(dist=104362),ASB3(dist=863002) | G | A | 0.00035667 | 3 | 30.330035 | 6.62672999 | 4.72E-06 | 4.97E-06 |
| rs12312017 | 12 | 60194008 | IMPUTED | SLC16A7(dist=10373),FAM19A2(dist=1908021) | G | A | 0.00035679 | 3 | 30.3298188 | 6.62674555 | 4.72E-06 | 4.97E-06 |
| rs376043750 | 2 | 52729403 | IMPUTED | LOC730100(dist=94348),MIR4431(dist=200257) | T | C | 0.00035679 | 3 | 30.3295545 | 6.62671375 | 4.72E-06 | 4.97E-06 |
| rs370547329 | 2 | 52729443 | IMPUTED | LOC730100(dist=94388),MIR4431(dist=200217) | C | T | 0.00035679 | 3 | 30.3295545 | 6.62671375 | 4.72E-06 | 4.97E-06 |
| rs371923479 | 2 | 52730307 | IMPUTED | LOC730100(dist=95252),MIR4431(dist=199353) | A | C | 0.00035679 | 3 | 30.3295545 | 6.62671375 | 4.72E-06 | 4.97E-06 |
| rs375202743 | 2 | 52730349 | IMPUTED | LOC730100(dist=95294),MIR4431(dist=199311) | C | T | 0.00035679 | 3 | 30.3295545 | 6.62671375 | 4.72E-06 | 4.97E-06 |
| rs375581355 | 2 | 52730992 | IMPUTED | LOC730100(dist=95937),MIR4431(dist=198668) | A | G | 0.00035679 | 3 | 30.3295545 | 6.62671375 | 4.72E-06 | 4.97E-06 |
| rs189626675 | 2 | 52731052 | IMPUTED | LOC730100(dist=95997),MIR4431(dist=198608) | G | T | 0.00035679 | 3 | 30.3295545 | 6.62671375 | 4.72E-06 | 4.97E-06 |
| rs375248108 | 2 | 52731202 | IMPUTED | LOC730100(dist=96147),MIR4431(dist=198458) | G | C | 0.00035679 | 3 | 30.3295545 | 6.62671375 | 4.72E-06 | 4.97E-06 |
| rs368013119 | 2 | 52731376 | IMPUTED | LOC730100(dist=96321),MIR4431(dist=198284) | C | T | 0.00035679 | 3 | 30.3295545 | 6.62671375 | 4.72E-06 | 4.97E-06 |
| rs376675214 | 2 | 52731939 | IMPUTED | LOC730100(dist=96884),MIR4431(dist=197721) | C | T | 0.00035679 | 3 | 30.3295545 | 6.62671375 | 4.72E-06 | 4.97E-06 |
| rs192842229 | 2 | 52732817 | IMPUTED | LOC730100(dist=97762),MIR4431(dist=196843) | G | A | 0.00035679 | 3 | 30.3295545 | 6.62671375 | 4.72E-06 | 4.97E-06 |
| rs148369445 | 19 | 450882 | IMPUTED | SHC2 | G | A | 0.00036701 | 3 | 30.321857 | 6.62509816 | 4.72E-06 | 4.97E-06 |
| rs538136531 | 2 | 52547581 | IMPUTED | LOC730100 | C | T | 0.00035691 | 3 | 30.3292213 | 6.6267188 | 4.72E-06 | 4.97E-06 |
| rs369074393 | 2 | 52972000 | IMPUTED | MIR4431(dist=42247),ASB3(dist=925117) | A | T | 0.00035667 | 3 | 30.3289107 | 6.62671019 | 4.72E-06 | 4.97E-06 |
| rs372007188 | 2 | 52974427 | IMPUTED | MIR4431(dist=44674),ASB3(dist=922690) | C | A | 0.00035667 | 3 | 30.3289107 | 6.62671019 | 4.72E-06 | 4.97E-06 |
| rs981157 | 2 | 52975415 | IMPUTED | MIR4431(dist=45662),ASB3(dist=921702) | C | T | 0.00035726 | 3 | 30.3288894 | 6.6267104 | 4.72E-06 | 4.97E-06 |
| rs531515532 | 2 | 55556616 | IMPUTED | CCDC88A | C | G | 0.00035881 | 3 | 30.3913641 | 6.64039351 | 4.72E-06 | 4.97E-06 |
| rs565522352 | 2 | 52855488 | IMPUTED | LOC730100(dist=220433),MIR4431(dist=74172) | T | C | 0.00035667 | 3 | 30.3281631 | 6.62671308 | 4.72E-06 | 4.97E-06 |
| rs547807201 | 2 | 52856336 | IMPUTED | LOC730100(dist=221281),MIR4431(dist=73324) | G | C | 0.00035667 | 3 | 30.3281631 | 6.62671308 | 4.72E-06 | 4.97E-06 |
| rs547794581 | 2 | 52857107 | IMPUTED | LOC730100(dist=222052),MIR4431(dist=72553) | G | A | 0.00035667 | 3 | 30.3281631 | 6.62671308 | 4.72E-06 | 4.97E-06 |
| rs576606058 | 2 | 52866519 | IMPUTED | LOC730100(dist=231464),MIR4431(dist=63141) | T | C | 0.00035667 | 3 | 30.3281631 | 6.62671308 | 4.72E-06 | 4.97E-06 |
| rs543958949 | 2 | 52866882 | IMPUTED | LOC730100(dist=231827),MIR4431(dist=62778) | A | C | 0.00035667 | 3 | 30.3281631 | 6.62671308 | 4.72E-06 | 4.97E-06 |
| rs559090140 | 2 | 52866914 | IMPUTED | LOC730100(dist=231859),MIR4431(dist=62746) | T | G | 0.00035667 | 3 | 30.3281631 | 6.62671308 | 4.72E-06 | 4.97E-06 |
| rs577331164 | 2 | 52867026 | IMPUTED | LOC730100(dist=231971),MIR4431(dist=62634) | T | C | 0.00035667 | 3 | 30.3281631 | 6.62671308 | 4.72E-06 | 4.97E-06 |
| rs528226082 | 2 | 52870621 | IMPUTED | LOC730100(dist=235566),MIR4431(dist=59039) | T | C | 0.00035667 | 3 | 30.3281631 | 6.62671308 | 4.72E-06 | 4.97E-06 |
| rs548867931 | 11 | 116625948 | IMPUTED | BUD13 | G | T | 0.00035643 | 3 | 30.3886077 | 6.63994067 | 4.73E-06 | 4.97E-06 |
| rs1568992 | 2 | 52708546 | IMPUTED | LOC730100(dist=73491),MIR4431(dist=221114) | C | T | 0.00035714 | 3 | 30.3277011 | 6.62669145 | 4.73E-06 | 4.97E-06 |
| rs371296017 | 2 | 52537462 | IMPUTED | LOC730100 | G | T | 0.00035667 | 3 | 30.3272512 | 6.6267205 | 4.73E-06 | 4.98E-06 |
| rs573025486 | 2 | 52539006 | IMPUTED | LOC730100 | G | A | 0.00035667 | 3 | 30.3272512 | 6.6267205 | 4.73E-06 | 4.98E-06 |
| rs531911265 | 2 | 52842946 | IMPUTED | LOC730100(dist=207891),MIR4431(dist=86714) | G | C | 0.00035691 | 3 | 30.3269058 | 6.6266875 | 4.73E-06 | 4.98E-06 |
| rs1483882 | 2 | 52953704 | IMPUTED | MIR4431(dist=23951),ASB3(dist=943413) | A | G | 0.00035809 | 3 | 30.3269068 | 6.62670997 | 4.73E-06 | 4.98E-06 |
| rs78935322 | 17 | 30032498 | IMPUTED | MIR365B(dist=129958),COPRS(dist=146386) | C | A | 0.00036463 | 3 | 30.3231337 | 6.6258876 | 4.73E-06 | 4.98E-06 |
| rs374368987 | 2 | 52912191 | IMPUTED | LOC730100(dist=277136),MIR4431(dist=17469) | T | C | 0.00035821 | 3 | 30.3267397 | 6.62668598 | 4.73E-06 | 4.98E-06 |
| rs116822094 | 2 | 52543145 | IMPUTED | LOC730100 | C | G | 0.00035679 | 3 | 30.3268183 | 6.62675588 | 4.73E-06 | 4.98E-06 |
| rs139929104 | 2 | 52996963 | IMPUTED | MIR4431(dist=67210),ASB3(dist=900154) | G | C | 0.00035774 | 3 | 30.3256868 | 6.62660632 | 4.73E-06 | 4.98E-06 |
| rs563451204 | 12 | 63484318 | IMPUTED | PPM1H(dist=155653),AVPR1A(dist=52221) | T | C | 0.00035702 | 3 | 30.325181 | 6.62670998 | 4.73E-06 | 4.98E-06 |
| rs79084599 | 2 | 52997421 | IMPUTED | MIR4431(dist=67668),ASB3(dist=899696) | A | T | 0.00035774 | 3 | 30.3245351 | 6.62660762 | 4.74E-06 | 4.98E-06 |
| rs4392298 | 2 | 53123847 | IMPUTED | MIR4431(dist=194094),ASB3(dist=773270) | T | C | 0.00035702 | 3 | 30.3239883 | 6.62667725 | 4.74E-06 | 4.99E-06 |

| | | | | | | | | | | | | |
|---|---|---|---|---|---|---|---|---|---|---|---|---|
| rs556888371 | 2 | 52616642 | IMPUTED | LINC01867 | G | T | 0.00035904 | 3 | 30.324523 | 6.62685677 | 4.74E-06 | 4.99E-06 |
| rs551132210 | 2 | 53160711 | IMPUTED | MIR4431(dist=230958),ASB3(dist=736406) | T | C | 0.00035691 | 3 | 30.3236357 | 6.62672067 | 4.74E-06 | 4.99E-06 |
| rs767463635 | 8 | 138721525 | IMPUTED | LOC101927915(dist=295694),FAM135B(dist=420741) | A | G | 0.00024899 | 2 | -33.334346 | 7.28471873 | 4.74E-06 | 4.99E-06 |
| rs116726745 | 17 | 30045836 | IMPUTED | MIR365B(dist=143296),COPRS(dist=133048) | G | A | 0.00036736 | 3 | 30.3123589 | 6.62434761 | 4.74E-06 | 4.99E-06 |
| rs549215614 | 2 | 52534543 | IMPUTED | LOC730100 | T | C | 0.00035679 | 3 | 30.3231388 | 6.62672095 | 4.74E-06 | 4.99E-06 |
| rs567527525 | 2 | 52534646 | IMPUTED | LOC730100 | A | G | 0.00035679 | 3 | 30.3231388 | 6.62672095 | 4.74E-06 | 4.99E-06 |
| rs538141205 | 2 | 52535009 | IMPUTED | LOC730100 | T | C | 0.00035679 | 3 | 30.3231388 | 6.62672095 | 4.74E-06 | 4.99E-06 |
| rs368894322 | 2 | 52536103 | IMPUTED | LOC730100 | G | A | 0.00035679 | 3 | 30.3231388 | 6.62672095 | 4.74E-06 | 4.99E-06 |
| rs369224228 | 2 | 52536378 | IMPUTED | LOC730100 | C | T | 0.00035679 | 3 | 30.3231388 | 6.62672095 | 4.74E-06 | 4.99E-06 |
| rs927161683 | 9 | 120693777 | IMPUTED | TLR4(dist=214008),BRINP1(dist=1235131) | G | A | 0.00035667 | 3 | 30.3323089 | 6.62879266 | 4.74E-06 | 4.99E-06 |
| rs550086640 | 2 | 52680796 | IMPUTED | LOC730100(dist=45741),MIR4431(dist=248864) | T | A | 0.00035667 | 3 | 30.3222921 | 6.62673382 | 4.75E-06 | 4.99E-06 |
| rs117405005 | 18 | 178216 | IMPUTED | USP14 | C | T | 0.0003556 | 3 | 30.566758 | 6.68024427 | 4.75E-06 | 4.99E-06 |
| rs112488623 | 8 | 82440157 | IMPUTED | FABP12 | G | A | 0.00033088 | 3 | -29.610887 | 6.47141325 | 4.75E-06 | 5.00E-06 |
| rs1843030 | 2 | 52841168 | IMPUTED | LOC730100(dist=206113),MIR4431(dist=88492) | G | A | 0.00035679 | 3 | 30.3205413 | 6.62669134 | 4.75E-06 | 5.00E-06 |
| rs542789750 | 2 | 52842539 | IMPUTED | LOC730100(dist=207484),MIR4431(dist=87121) | T | C | 0.00035679 | 3 | 30.3205413 | 6.62669134 | 4.75E-06 | 5.00E-06 |
| rs533790736 | 2 | 53148225 | IMPUTED | MIR4431(dist=218472),ASB3(dist=748892) | G | A | 0.00035774 | 3 | 30.3189499 | 6.62658751 | 4.75E-06 | 5.00E-06 |
| rs755646528 | 9 | 120549489 | IMPUTED | TLR4(dist=69720),BRINP1(dist=1379419) | G | A | 0.00035714 | 3 | 30.4100732 | 6.64651459 | 4.75E-06 | 5.00E-06 |
| rs532640589 | 2 | 52605778 | IMPUTED | LINC01867,LOC730100 | T | C | 0.00035916 | 3 | 30.3196386 | 6.62685295 | 4.76E-06 | 5.00E-06 |
| rs570590317 | 2 | 52859091 | IMPUTED | LOC730100(dist=224036),MIR4431(dist=70569) | C | T | 0.00035714 | 3 | 30.3188083 | 6.62670652 | 4.76E-06 | 5.01E-06 |
| rs572045690 | 2 | 53168467 | IMPUTED | MIR4431(dist=238714),ASB3(dist=728650) | C | G | 0.00035702 | 3 | 30.318053 | 6.6267111 | 4.76E-06 | 5.01E-06 |
| rs140699806 | 2 | 52840408 | IMPUTED | LOC730100(dist=205353),MIR4431(dist=89252) | A | G | 0.00035726 | 3 | 30.3176101 | 6.62667334 | 4.76E-06 | 5.01E-06 |
| rs1949927 | 2 | 52703501 | IMPUTED | LOC730100(dist=68446),MIR4431(dist=226159) | C | T | 0.00035881 | 3 | 30.3170411 | 6.62656132 | 4.76E-06 | 5.01E-06 |
| rs565861418 | 2 | 53157289 | IMPUTED | MIR4431(dist=227536),ASB3(dist=739828) | T | C | 0.00035833 | 3 | 30.3156966 | 6.62652295 | 4.76E-06 | 5.01E-06 |
| rs183149432 | 14 | 30019949 | IMPUTED | LINC01551(dist=755949),PRKD1(dist=25737) | G | A | 0.00026087 | 2 | -33.213623 | 7.26006135 | 4.77E-06 | 5.01E-06 |
| rs545785774 | 2 | 52884260 | IMPUTED | LOC730100(dist=249205),MIR4431(dist=45400) | G | A | 0.00035691 | 3 | 30.315726 | 6.62674474 | 4.77E-06 | 5.02E-06 |
| rs569417827 | 2 | 52849837 | IMPUTED | LOC730100(dist=214782),MIR4431(dist=79823) | C | T | 0.00035691 | 3 | 30.3151028 | 6.62667401 | 4.77E-06 | 5.02E-06 |
| rs529411347 | 2 | 53159441 | IMPUTED | MIR4431(dist=229688),ASB3(dist=737676) | T | C | 0.00035786 | 3 | 30.3150371 | 6.62667795 | 4.77E-06 | 5.02E-06 |
| rs563775596 | 2 | 52518812 | IMPUTED | LOC730100 | A | C | 0.00035702 | 3 | 30.314835 | 6.62671326 | 4.77E-06 | 5.02E-06 |
| rs1922208 | 2 | 52519636 | IMPUTED | LOC730100 | C | A | 0.00035702 | 3 | 30.314835 | 6.62671326 | 4.77E-06 | 5.02E-06 |
| rs546338448 | 2 | 52519976 | IMPUTED | LOC730100 | G | C | 0.00035702 | 3 | 30.314835 | 6.62671326 | 4.77E-06 | 5.02E-06 |
| rs771876650 | 9 | 120670555 | IMPUTED | TLR4(dist=190786),BRINP1(dist=1258353) | G | A | 0.00035702 | 3 | 30.3142055 | 6.62668176 | 4.77E-06 | 5.02E-06 |
| rs546159418 | 2 | 53139536 | IMPUTED | MIR4431(dist=209783),ASB3(dist=757581) | C | T | 0.00035988 | 3 | 30.310387 | 6.6259367 | 4.77E-06 | 5.02E-06 |
| rs370296178 | 2 | 52954734 | IMPUTED | MIR4431(dist=24981),ASB3(dist=942383) | G | A | 0.0003575 | 3 | 30.3132331 | 6.62664725 | 4.77E-06 | 5.02E-06 |
| rs376179130 | 2 | 52920989 | IMPUTED | LOC730100(dist=285934),MIR4431(dist=8671) | T | G | 0.00035762 | 3 | 30.3131567 | 6.62665672 | 4.78E-06 | 5.02E-06 |
| rs537456120 | 2 | 52875961 | IMPUTED | LOC730100(dist=240906),MIR4431(dist=53699) | A | T | 0.00035667 | 3 | 30.3129406 | 6.6267255 | 4.78E-06 | 5.03E-06 |
| rs1922190 | 2 | 52512210 | IMPUTED | LOC730100 | A | G | 0.00035691 | 3 | 30.3126883 | 6.62676158 | 4.78E-06 | 5.03E-06 |
| rs533762077 | 2 | 53164653 | IMPUTED | MIR4431(dist=234900),ASB3(dist=732464) | G | A | 0.00035726 | 3 | 30.3112445 | 6.62668995 | 4.78E-06 | 5.03E-06 |
| rs145796806 | 11 | 116650184 | IMPUTED | ZPR1 | C | T | 0.00035619 | 3 | 30.4020957 | 6.64662059 | 4.78E-06 | 5.03E-06 |
| rs533586809 | 2 | 53163001 | IMPUTED | MIR4431(dist=233248),ASB3(dist=734116) | A | G | 0.00035762 | 3 | 30.3106328 | 6.62666918 | 4.78E-06 | 5.03E-06 |
| rs373904199 | 2 | 53099935 | IMPUTED | MIR4431(dist=170182),ASB3(dist=797182) | T | C | 0.00036011 | 3 | 30.3094996 | 6.62656042 | 4.79E-06 | 5.04E-06 |
| rs557335377 | 2 | 52662014 | IMPUTED | LOC730100(dist=26959),MIR4431(dist=267646) | G | A | 0.00035833 | 3 | 30.3100775 | 6.62674415 | 4.79E-06 | 5.04E-06 |
| rs535792665 | 2 | 52873266 | IMPUTED | LOC730100(dist=238211),MIR4431(dist=56394) | T | A | 0.00035679 | 3 | 30.3097665 | 6.6267214 | 4.79E-06 | 5.04E-06 |
| rs550930665 | 2 | 53143403 | IMPUTED | MIR4431(dist=213650),ASB3(dist=753714) | G | T | 0.00035928 | 3 | 30.3067739 | 6.62607557 | 4.79E-06 | 5.04E-06 |
| rs535265668 | 2 | 52741408 | IMPUTED | LOC730100(dist=106353),MIR4431(dist=188252) | C | T | 0.00035714 | 3 | 30.3096683 | 6.62675648 | 4.79E-06 | 5.04E-06 |
| rs574902418 | 2 | 52737341 | IMPUTED | LOC730100(dist=102286),MIR4431(dist=192319) | A | G | 0.0003575 | 3 | 30.3090182 | 6.62673568 | 4.79E-06 | 5.04E-06 |
| rs574920246 | 2 | 52514121 | IMPUTED | LOC730100 | C | A | 0.00035702 | 3 | 30.3085495 | 6.62675916 | 4.79E-06 | 5.04E-06 |
| rs111702906 | 19 | 554425 | IMPUTED | GZMM(dist=4505),BSG(dist=16852) | C | T | 0.00037295 | 3 | 30.2973162 | 6.62441703 | 4.79E-06 | 5.04E-06 |
| rs544087633 | 2 | 53137882 | IMPUTED | MIR4431(dist=208129),ASB3(dist=759235) | A | G | 0.00035988 | 3 | 30.3037864 | 6.6258769 | 4.80E-06 | 5.05E-06 |
| rs562793161 | 2 | 53138286 | IMPUTED | MIR4431(dist=208533),ASB3(dist=758831) | T | C | 0.00035988 | 3 | 30.3037864 | 6.6258769 | 4.80E-06 | 5.05E-06 |
| rs533123190 | 2 | 53138320 | IMPUTED | MIR4431(dist=208567),ASB3(dist=758797) | A | G | 0.00035988 | 3 | 30.3037864 | 6.6258769 | 4.80E-06 | 5.05E-06 |
| rs111977451 | 17 | 39116513 | IMPUTED | KRT39 | G | A | 0.00379677 | 32 | -8.8056827 | 1.92535915 | 4.80E-06 | 5.05E-06 |
| rs56109124 | 2 | 52771895 | IMPUTED | LOC730100(dist=136840),MIR4431(dist=157765) | T | A | 0.00036736 | 3 | 30.3011681 | 6.62539625 | 4.80E-06 | 5.05E-06 |

| | | | | | | | | | | | | |
|---|---|---|---|---|---|---|---|---|---|---|---|---|
| rs573835048 | 2 | 53137673 | IMPUTED | MIR4431(dist=207920),ASB3(dist=759444) | C | A | 0.00035976 | 3 | 30.3031719 | 6.62587929 | 4.80E-06 | 5.05E-06 |
| rs535448254 | 2 | 52616242 | IMPUTED | LINC01867,LOC730100 | T | C | 0.00035845 | 3 | 30.3074326 | 6.62688356 | 4.80E-06 | 5.05E-06 |
| rs557194704 | 2 | 52600815 | IMPUTED | LINC01867,LOC730100 | T | C | 0.00035893 | 3 | 30.3072315 | 6.62689516 | 4.80E-06 | 5.05E-06 |
| rs374221587 | 2 | 52729166 | IMPUTED | LOC730100(dist=94111),MIR4431(dist=200494) | C | T | 0.00035821 | 3 | 30.3063751 | 6.62675177 | 4.80E-06 | 5.05E-06 |
| rs372377537 | 2 | 52704541 | IMPUTED | LOC730100(dist=69486),MIR4431(dist=225119) | T | A | 0.00036011 | 3 | 30.3028916 | 6.62601093 | 4.80E-06 | 5.05E-06 |
| rs533135336 | 2 | 52766733 | IMPUTED | LOC730100(dist=131678),MIR4431(dist=162927) | C | T | 0.00035916 | 3 | 30.305284 | 6.62669509 | 4.80E-06 | 5.05E-06 |
| rs560147099 | 2 | 52768261 | IMPUTED | LOC730100(dist=133206),MIR4431(dist=161399) | G | C | 0.00035916 | 3 | 30.305284 | 6.62669509 | 4.80E-06 | 5.05E-06 |
| rs568469982 | 2 | 52770892 | IMPUTED | LOC730100(dist=135837),MIR4431(dist=158768) | A | G | 0.00035916 | 3 | 30.305284 | 6.62669509 | 4.80E-06 | 5.05E-06 |
| rs535560757 | 2 | 52771898 | IMPUTED | LOC730100(dist=136843),MIR4431(dist=157762) | T | G | 0.00035916 | 3 | 30.305284 | 6.62669509 | 4.80E-06 | 5.05E-06 |
| rs539595685 | 2 | 52775275 | IMPUTED | LOC730100(dist=140220),MIR4431(dist=154385) | T | A | 0.00035916 | 3 | 30.305284 | 6.62669509 | 4.80E-06 | 5.05E-06 |
| rs573446993 | 2 | 52775400 | IMPUTED | LOC730100(dist=140345),MIR4431(dist=154260) | G | T | 0.00035916 | 3 | 30.305284 | 6.62669509 | 4.80E-06 | 5.05E-06 |
| rs555055715 | 2 | 52778733 | IMPUTED | LOC730100(dist=143678),MIR4431(dist=150927) | C | T | 0.00035916 | 3 | 30.305284 | 6.62669509 | 4.80E-06 | 5.05E-06 |
| rs573548698 | 2 | 52778763 | IMPUTED | LOC730100(dist=143708),MIR4431(dist=150897) | C | T | 0.00035916 | 3 | 30.305284 | 6.62669509 | 4.80E-06 | 5.05E-06 |
| rs137932806 | 19 | 536014 | IMPUTED | CDC34 | C | T | 0.000372 | 3 | 30.874768 | 6.75123669 | 4.80E-06 | 5.05E-06 |
| rs556358565 | 2 | 53134151 | IMPUTED | MIR4431(dist=204398),ASB3(dist=762966) | G | T | 0.00035976 | 3 | 30.3010639 | 6.62580745 | 4.80E-06 | 5.05E-06 |
| rs555516619 | 2 | 53135616 | IMPUTED | MIR4431(dist=205863),ASB3(dist=761501) | T | C | 0.00035976 | 3 | 30.3010639 | 6.62580745 | 4.80E-06 | 5.05E-06 |
| rs61369041 | 2 | 52757732 | IMPUTED | LOC730100(dist=122677),MIR4431(dist=171928) | C | T | 0.00036772 | 3 | 30.2989357 | 6.62538646 | 4.80E-06 | 5.06E-06 |
| rs570097903 | 2 | 52874483 | IMPUTED | LOC730100(dist=239428),MIR4431(dist=55177) | A | G | 0.00035786 | 3 | 30.3045342 | 6.62664667 | 4.80E-06 | 5.06E-06 |
| rs533251316 | 2 | 52752898 | IMPUTED | LOC730100(dist=117843),MIR4431(dist=176762) | G | A | 0.00035738 | 3 | 30.3044362 | 6.62674403 | 4.81E-06 | 5.06E-06 |
| rs566795496 | 2 | 52754974 | IMPUTED | LOC730100(dist=119919),MIR4431(dist=174686) | A | G | 0.00035738 | 3 | 30.3044362 | 6.62674403 | 4.81E-06 | 5.06E-06 |
| rs530144269 | 2 | 52621533 | IMPUTED | LOC730100 | T | C | 0.00035869 | 3 | 30.3040121 | 6.62689261 | 4.81E-06 | 5.06E-06 |
| rs552629121 | 2 | 52623746 | IMPUTED | LOC730100 | T | C | 0.00035869 | 3 | 30.3040121 | 6.62689261 | 4.81E-06 | 5.06E-06 |
| rs377508110 | 2 | 53062231 | IMPUTED | MIR4431(dist=132478),ASB3(dist=834886) | A | G | 0.00035714 | 3 | 30.3026794 | 6.62663955 | 4.81E-06 | 5.06E-06 |
| rs536813480 | 2 | 52850328 | IMPUTED | LOC730100(dist=215273),MIR4431(dist=79332) | G | T | 0.0003575 | 3 | 30.3024716 | 6.62667539 | 4.81E-06 | 5.06E-06 |
| rs538042079 | 2 | 53149162 | IMPUTED | MIR4431(dist=219409),ASB3(dist=747955) | C | A | 0.00035904 | 3 | 30.3001282 | 6.62628215 | 4.81E-06 | 5.07E-06 |
| rs531732261 | 2 | 53151413 | IMPUTED | MIR4431(dist=221660),ASB3(dist=745704) | C | T | 0.00035904 | 3 | 30.3001282 | 6.62628215 | 4.81E-06 | 5.07E-06 |
| rs558373841 | 2 | 52850981 | IMPUTED | LOC730100(dist=215926),MIR4431(dist=78679) | A | C | 0.00035762 | 3 | 30.3014298 | 6.626661 | 4.82E-06 | 5.07E-06 |
| rs577065460 | 2 | 52851096 | IMPUTED | LOC730100(dist=216041),MIR4431(dist=78564) | G | A | 0.00035762 | 3 | 30.3014298 | 6.626661 | 4.82E-06 | 5.07E-06 |
| rs374185367 | 2 | 53062136 | IMPUTED | MIR4431(dist=132383),ASB3(dist=834981) | G | A | 0.00038888 | 3 | 30.066415 | 6.5754043 | 4.82E-06 | 5.07E-06 |
| rs559036395 | 2 | 52736744 | IMPUTED | LOC730100(dist=101689),MIR4431(dist=192916) | G | T | 0.00035786 | 3 | 30.3002025 | 6.62669408 | 4.82E-06 | 5.07E-06 |
| rs577415351 | 2 | 52736773 | IMPUTED | LOC730100(dist=101718),MIR4431(dist=192887) | T | C | 0.00035786 | 3 | 30.3002025 | 6.62669408 | 4.82E-06 | 5.07E-06 |
| rs150859756 | 2 | 52815352 | IMPUTED | LOC730100(dist=180297),MIR4431(dist=114308) | C | T | 0.00036047 | 3 | 30.2911133 | 6.6247647 | 4.82E-06 | 5.07E-06 |
| rs535204752 | 2 | 52629081 | IMPUTED | LOC730100 | T | C | 0.00035833 | 3 | 30.3005694 | 6.62689266 | 4.82E-06 | 5.07E-06 |
| rs759324230 | 5 | 171210528 | IMPUTED | FGF18(dist=325898),SMIM23(dist=2348) | C | G | 0.00039256 | 3 | 30.2682764 | 6.61990436 | 4.82E-06 | 5.08E-06 |
| rs7421904 | 2 | 52762547 | IMPUTED | LOC730100(dist=127492),MIR4431(dist=167113) | G | A | 0.0003575 | 3 | 30.2982671 | 6.62674644 | 4.83E-06 | 5.08E-06 |
| rs569029246 | 2 | 52775249 | IMPUTED | LOC730100(dist=140194),MIR4431(dist=154411) | C | A | 0.0003575 | 3 | 30.2982671 | 6.62674644 | 4.83E-06 | 5.08E-06 |
| rs374615507 | 2 | 53120357 | IMPUTED | MIR4431(dist=190604),ASB3(dist=776760) | C | T | 0.00035774 | 3 | 30.297376 | 6.62659892 | 4.83E-06 | 5.08E-06 |
| rs73489636 | 19 | 452062 | IMPUTED | SHC2 | C | T | 0.00036261 | 3 | 30.2893857 | 6.62487925 | 4.83E-06 | 5.08E-06 |
| rs74363876 | 2 | 52517336 | IMPUTED | LOC730100 | G | A | 0.00035714 | 3 | 30.2980725 | 6.62679694 | 4.83E-06 | 5.08E-06 |
| rs115676637 | 19 | 450068 | IMPUTED | SHC2 | G | A | 0.00036166 | 3 | 30.2911729 | 6.62532627 | 4.83E-06 | 5.08E-06 |
| rs116669807 | 19 | 450165 | IMPUTED | SHC2 | G | A | 0.00036166 | 3 | 30.2911729 | 6.62532627 | 4.83E-06 | 5.08E-06 |
| rs575628829 | 2 | 52736374 | IMPUTED | LOC730100(dist=101319),MIR4431(dist=193286) | T | G | 0.00035797 | 3 | 30.2972099 | 6.62667433 | 4.83E-06 | 5.08E-06 |
| rs537393677 | 2 | 52736378 | IMPUTED | LOC730100(dist=101323),MIR4431(dist=193282) | G | A | 0.00035797 | 3 | 30.2972099 | 6.62667433 | 4.83E-06 | 5.08E-06 |
| rs17123009 | 12 | 60175789 | IMPUTED | SLC16A7(NM_001270623:c.*23290>0,NM_001270622:c.*2 | G | A | 0.00035952 | 3 | 30.2935782 | 6.62602425 | 4.83E-06 | 5.09E-06 |
| rs367906151 | 2 | 53086176 | IMPUTED | MIR4431(dist=156423),ASB3(dist=810941) | C | T | 0.00036071 | 3 | 30.2961288 | 6.6266092 | 4.83E-06 | 5.09E-06 |
| rs148648064 | 19 | 471828 | IMPUTED | ODF3L2 | C | T | 0.00036629 | 3 | 30.2789202 | 6.62284565 | 4.83E-06 | 5.09E-06 |
| rs796326000 | 19 | 451079 | IMPUTED | SHC2 | C | T | 0.00036213 | 3 | 30.2886988 | 6.62510896 | 4.84E-06 | 5.09E-06 |
| rs370767240 | 2 | 52568339 | IMPUTED | LOC730100 | A | C | 0.00035774 | 3 | 30.2957935 | 6.62680565 | 4.84E-06 | 5.09E-06 |
| rs111824506 | 19 | 451666 | IMPUTED | SHC2 | C | T | 0.00036225 | 3 | 30.2870453 | 6.62500997 | 4.84E-06 | 5.09E-06 |
| rs111572517 | 19 | 451746 | IMPUTED | SHC2 | C | T | 0.00036225 | 3 | 30.2870453 | 6.62500997 | 4.84E-06 | 5.09E-06 |
| rs569104524 | 2 | 52752394 | IMPUTED | LOC730100(dist=117339),MIR4431(dist=177266) | G | A | 0.00035904 | 3 | 30.2947422 | 6.62669987 | 4.84E-06 | 5.09E-06 |
| rs551669043 | 2 | 52753497 | IMPUTED | LOC730100(dist=118442),MIR4431(dist=176163) | G | C | 0.00035904 | 3 | 30.2947422 | 6.62669987 | 4.84E-06 | 5.09E-06 |

| | | | | | | | | | | | | |
|---|---|---|---|---|---|---|---|---|---|---|---|---|
| rs568453194 | 2 | 52757915 | IMPUTED | LOC730100(dist=122860),MIR4431(dist=171745) | T | C | 0.00035904 | 3 | 30.2947422 | 6.62669987 | 4.84E-06 | 5.09E-06 |
| rs535990627 | 2 | 52758087 | IMPUTED | LOC730100(dist=123032),MIR4431(dist=171573) | T | C | 0.00035904 | 3 | 30.2947422 | 6.62669987 | 4.84E-06 | 5.09E-06 |
| rs575466167 | 2 | 52759253 | IMPUTED | LOC730100(dist=124198),MIR4431(dist=170407) | G | C | 0.00035904 | 3 | 30.2947422 | 6.62669987 | 4.84E-06 | 5.09E-06 |
| rs112676561 | 9 | 88752947 | IMPUTED | LOC101927623 | G | A | 0.00098336 | 8 | 19.0932209 | 4.17655536 | 4.84E-06 | 5.09E-06 |
| rs372931574 | 2 | 52565855 | IMPUTED | LOC730100 | A | C | 0.00035774 | 3 | 30.2942978 | 6.62682453 | 4.84E-06 | 5.10E-06 |
| rs13407120 | 2 | 52736614 | IMPUTED | LOC730100(dist=101559),MIR4431(dist=193046) | C | T | 0.00035797 | 3 | 30.2929162 | 6.62669787 | 4.85E-06 | 5.10E-06 |
| rs117725199 | 18 | 289856 | IMPUTED | THOC1(dist=21797),COLEC12(dist=29499) | C | A | 0.00035821 | 3 | 30.2923151 | 6.62664387 | 4.85E-06 | 5.10E-06 |
| rs558053953 | 2 | 52630672 | IMPUTED | LOC730100 | T | C | 0.00036071 | 3 | 30.2930988 | 6.62683061 | 4.85E-06 | 5.10E-06 |
| rs552519443 | 2 | 52503465 | IMPUTED | LOC730100 | A | G | 0.00035049 | 3 | 30.8219408 | 6.74266906 | 4.85E-06 | 5.10E-06 |
| rs545730630 | 2 | 52560097 | IMPUTED | LOC730100 | T | C | 0.00035738 | 3 | 30.2922064 | 6.6267968 | 4.85E-06 | 5.10E-06 |
| rs553795061 | 2 | 52560104 | IMPUTED | LOC730100 | G | A | 0.00035738 | 3 | 30.2922064 | 6.6267968 | 4.85E-06 | 5.10E-06 |
| rs547319884 | 2 | 52565681 | IMPUTED | LOC730100 | C | T | 0.00035762 | 3 | 30.2921286 | 6.62683514 | 4.85E-06 | 5.10E-06 |
| rs369867526 | 2 | 52567276 | IMPUTED | LOC730100 | G | C | 0.00035762 | 3 | 30.2921286 | 6.62683514 | 4.85E-06 | 5.10E-06 |
| rs541382616 | 2 | 54958264 | IMPUTED | EML6 | A | G | 0.00036178 | 3 | 30.2895892 | 6.62628045 | 4.85E-06 | 5.10E-06 |
| rs189535388 | 2 | 52662612 | IMPUTED | LOC730100(dist=27557),MIR4431(dist=267048) | C | T | 0.00035833 | 3 | 30.2917675 | 6.6267679 | 4.85E-06 | 5.10E-06 |
| rs371284192 | 2 | 52562748 | IMPUTED | LOC730100 | G | T | 0.00035726 | 3 | 30.291513 | 6.62681137 | 4.85E-06 | 5.11E-06 |
| rs137868799 | 2 | 55621333 | IMPUTED | CCDC88A | C | T | 0.0003594 | 3 | 30.3824546 | 6.64692585 | 4.86E-06 | 5.11E-06 |
| rs994068702 | 11 | 120312581 | IMPUTED | ARHGEF12 | A | G | 0.00035821 | 3 | 30.289314 | 6.62670245 | 4.86E-06 | 5.11E-06 |
| rs542724820 | 2 | 52560729 | IMPUTED | LOC730100 | C | T | 0.00035714 | 3 | 30.2893438 | 6.62682199 | 4.86E-06 | 5.11E-06 |
| rs367851740 | 2 | 52561853 | IMPUTED | LOC730100 | G | T | 0.00035714 | 3 | 30.2893438 | 6.62682199 | 4.86E-06 | 5.11E-06 |
| rs565131686 | 2 | 52563049 | IMPUTED | LOC730100 | T | C | 0.00035714 | 3 | 30.2893438 | 6.62682199 | 4.86E-06 | 5.11E-06 |
| rs531702345 | 2 | 52578740 | IMPUTED | LOC730100 | A | C | 0.00035714 | 3 | 30.2893438 | 6.62682199 | 4.86E-06 | 5.11E-06 |
| rs574057725 | 2 | 55484418 | IMPUTED | MTIF2 | C | G | 0.00036023 | 3 | 30.2871147 | 6.62658726 | 4.86E-06 | 5.12E-06 |
| rs558937324 | 2 | 55358340 | IMPUTED | RTN4 | T | C | 0.00036213 | 3 | 30.2860137 | 6.62653698 | 4.87E-06 | 5.12E-06 |
| rs368720644 | 2 | 52567528 | IMPUTED | LOC730100 | C | G | 0.00035762 | 3 | 30.2871203 | 6.62682754 | 4.87E-06 | 5.12E-06 |
| rs368707584 | 2 | 52569011 | IMPUTED | LOC730100 | G | C | 0.00035762 | 3 | 30.2871203 | 6.62682754 | 4.87E-06 | 5.12E-06 |
| rs577741617 | 2 | 52678367 | IMPUTED | LOC730100(dist=43312),MIR4431(dist=251293) | G | A | 0.00035714 | 3 | 30.285848 | 6.62677176 | 4.87E-06 | 5.13E-06 |
| rs149614586 | 2 | 52559435 | IMPUTED | LOC730100 | G | C | 0.00035738 | 3 | 30.2855103 | 6.62682904 | 4.87E-06 | 5.13E-06 |
| rs554536411 | 2 | 52569404 | IMPUTED | LOC730100 | C | T | 0.00035774 | 3 | 30.2850668 | 6.62683022 | 4.88E-06 | 5.13E-06 |
| rs375501748 | 2 | 52569709 | IMPUTED | LOC730100 | G | A | 0.00035774 | 3 | 30.2850668 | 6.62683022 | 4.88E-06 | 5.13E-06 |
| rs373708219 | 2 | 52573401 | IMPUTED | LOC730100 | G | A | 0.00035774 | 3 | 30.2850668 | 6.62683022 | 4.88E-06 | 5.13E-06 |
| rs113739919 | 19 | 450931 | IMPUTED | SHC2 | A | G | 0.00036261 | 3 | 30.2762822 | 6.62514081 | 4.88E-06 | 5.13E-06 |
| rs79053136 | 11 | 36076320 | IMPUTED | LDLRAD3 | G | C | 0.05600226 | 471 | -2.3232562 | 0.50838385 | 4.88E-06 | 5.13E-06 |
| rs148674242 | 2 | 52569030 | IMPUTED | LOC730100 | G | A | 0.00035786 | 3 | 30.2837004 | 6.62682832 | 4.88E-06 | 5.13E-06 |
| rs559807846 | 19 | 460403 | IMPUTED | SHC2 | G | A | 0.00036368 | 3 | 30.2684529 | 6.62398264 | 4.89E-06 | 5.14E-06 |
| rs566648755 | 2 | 52534240 | IMPUTED | LOC730100 | C | T | 0.00035738 | 3 | 30.2811452 | 6.62682574 | 4.89E-06 | 5.14E-06 |
| rs549035362 | 2 | 52657316 | IMPUTED | LOC730100(dist=22261),MIR4431(dist=272344) | A | G | 0.00036071 | 3 | 30.2789673 | 6.62641899 | 4.89E-06 | 5.14E-06 |
| rs372237158 | 2 | 52569628 | IMPUTED | LOC730100 | C | T | 0.00035762 | 3 | 30.2789871 | 6.6268365 | 4.90E-06 | 5.15E-06 |
| rs369944585 | 2 | 52569929 | IMPUTED | LOC730100 | T | A | 0.00035762 | 3 | 30.2789871 | 6.6268365 | 4.90E-06 | 5.15E-06 |
| rs8104724 | 19 | 462352 | IMPUTED | ODF3L2(dist=994) | G | C | 0.00036439 | 3 | 30.264838 | 6.62400079 | 4.90E-06 | 5.16E-06 |
| rs550501118 | 2 | 52532072 | IMPUTED | LOC730100 | T | C | 0.0003575 | 3 | 30.2770065 | 6.62682332 | 4.90E-06 | 5.16E-06 |
| rs533037143 | 2 | 52533449 | IMPUTED | LOC730100 | C | G | 0.0003575 | 3 | 30.2770065 | 6.62682332 | 4.90E-06 | 5.16E-06 |
| rs369943916 | 2 | 52533626 | IMPUTED | LOC730100 | G | A | 0.0003575 | 3 | 30.2770065 | 6.62682332 | 4.90E-06 | 5.16E-06 |
| rs572181905 | 2 | 52679204 | IMPUTED | LOC730100(dist=44149),MIR4431(dist=250456) | A | C | 0.00035726 | 3 | 30.2766714 | 6.62677408 | 4.90E-06 | 5.16E-06 |
| rs111276455 | 19 | 450507 | IMPUTED | SHC2 | C | T | 0.00036285 | 3 | 30.2694247 | 6.62525064 | 4.91E-06 | 5.16E-06 |
| rs75408787 | 19 | 450626 | IMPUTED | SHC2 | C | T | 0.00036285 | 3 | 30.2694247 | 6.62525064 | 4.91E-06 | 5.16E-06 |
| rs74459457 | 19 | 450627 | IMPUTED | SHC2 | A | G | 0.00036285 | 3 | 30.2694247 | 6.62525064 | 4.91E-06 | 5.16E-06 |
| rs73489617 | 19 | 450715 | IMPUTED | SHC2 | C | T | 0.00036285 | 3 | 30.2694247 | 6.62525064 | 4.91E-06 | 5.16E-06 |
| rs111367969 | 19 | 450814 | IMPUTED | SHC2 | G | A | 0.00036285 | 3 | 30.2694247 | 6.62525064 | 4.91E-06 | 5.16E-06 |
| rs561016172 | 1 | 91085967 | IMPUTED | ZNF326(dist=584878),SNORD3G(dist=37340) | C | T | 0.00025707 | 2 | -33.2033 | 7.26758108 | 4.91E-06 | 5.16E-06 |
| rs568891853 | 2 | 52522576 | IMPUTED | LOC730100 | A | T | 0.00035762 | 3 | 30.2728415 | 6.62681804 | 4.92E-06 | 5.17E-06 |
| rs566590054 | 2 | 52525433 | IMPUTED | LOC730100 | A | T | 0.00035762 | 3 | 30.2728415 | 6.62681804 | 4.92E-06 | 5.17E-06 |

| | | | | | | | | | | | | |
|---|---|---|---|---|---|---|---|---|---|---|---|---|
| rs574004397 | 2 | 52526286 | IMPUTED | LOC730100 | C | T | 0.00035762 | 3 | 30.2728415 | 6.62681804 | 4.92E-06 | 5.17E-06 |
| rs557087923 | 2 | 52527447 | IMPUTED | LOC730100 | T | G | 0.00035762 | 3 | 30.2728415 | 6.62681804 | 4.92E-06 | 5.17E-06 |
| rs367619843 | 2 | 52528524 | IMPUTED | LOC730100 | C | T | 0.00035762 | 3 | 30.2728415 | 6.62681804 | 4.92E-06 | 5.17E-06 |
| rs562008490 | 2 | 52529152 | IMPUTED | LOC730100 | T | G | 0.00035762 | 3 | 30.2728415 | 6.62681804 | 4.92E-06 | 5.17E-06 |
| rs73489613 | 19 | 449982 | IMPUTED | SHC2 | C | A | 0.00036285 | 3 | 30.2660003 | 6.62535787 | 4.92E-06 | 5.17E-06 |
| rs374706023 | 2 | 52952647 | IMPUTED | MIR4431(dist=22894),ASB3(dist=944470) | C | T | 0.00035893 | 3 | 30.2709 | 6.62670732 | 4.92E-06 | 5.18E-06 |
| rs147188679 | 17 | 30023108 | IMPUTED | MIR365B(dist=120568),COPRS(dist=155776) | T | C | 0.00044141 | 4 | 28.8872855 | 6.32400702 | 4.93E-06 | 5.18E-06 |
| rs76490838 | 19 | 450238 | IMPUTED | SHC2 | T | G | 0.0003632 | 3 | 30.2630871 | 6.6253517 | 4.93E-06 | 5.19E-06 |
| rs77744218 | 19 | 450302 | IMPUTED | SHC2 | A | G | 0.00036332 | 3 | 30.2614927 | 6.62525907 | 4.93E-06 | 5.19E-06 |
| rs531639262 | 2 | 52670686 | IMPUTED | LOC730100(dist=35631),MIR4431(dist=258974) | G | A | 0.00035738 | 3 | 30.2674686 | 6.62677355 | 4.94E-06 | 5.19E-06 |
| rs544928443 | 2 | 52675896 | IMPUTED | LOC730100(dist=40841),MIR4431(dist=253764) | C | T | 0.00036011 | 3 | 30.2663443 | 6.62653914 | 4.94E-06 | 5.19E-06 |
| rs112434157 | 19 | 450231 | IMPUTED | SHC2 | A | G | 0.0003632 | 3 | 30.2573391 | 6.6253905 | 4.95E-06 | 5.21E-06 |
| rs560255773 | 2 | 52679092 | IMPUTED | LOC730100(dist=44037),MIR4431(dist=250568) | A | G | 0.0003575 | 3 | 30.2582396 | 6.62677016 | 4.97E-06 | 5.23E-06 |
| rs964215441 | 1 | 108294081 | IMPUTED | VAV3 | G | A | 0.00035298 | 3 | 30.9362152 | 6.77531789 | 4.97E-06 | 5.23E-06 |
| rs2043080 | 2 | 3726163 | GENOTYPED | ALLC | T | C | 0.47976194 | 4037 | 1.08171528 | 0.23690851 | 4.97E-06 | 5.23E-06 |
| rs17841752 | 19 | 536527 | IMPUTED | CDC34 | C | T | 0.0003714 | 3 | 30.820263 | 6.75126736 | 4.99E-06 | 5.25E-06 |
| rs549162189 | 9 | 117659401 | IMPUTED | TNFSF8 | G | A | 0.00043 | 4 | 30.1302321 | 6.60031514 | 5.00E-06 | 5.25E-06 |
| rs73489629 | 19 | 451908 | IMPUTED | SHC2 | A | G | 0.00036344 | 3 | 30.2404027 | 6.62497147 | 5.00E-06 | 5.26E-06 |
| rs73489619 | 19 | 451465 | IMPUTED | SHC2 | G | A | 0.00036356 | 3 | 30.2386213 | 6.62500089 | 5.01E-06 | 5.27E-06 |
| rs73489621 | 19 | 451583 | IMPUTED | SHC2 | C | G | 0.00036356 | 3 | 30.2386213 | 6.62500089 | 5.01E-06 | 5.27E-06 |
| rs17042677 | 2 | 52507535 | IMPUTED | LOC730100 | A | T | 0.00035786 | 3 | 30.2651262 | 6.63108905 | 5.02E-06 | 5.28E-06 |
| rs372168808 | 2 | 52704357 | IMPUTED | LOC730100(dist=69302),MIR4431(dist=225303) | T | A | 0.00036522 | 3 | 30.2229712 | 6.62240449 | 5.02E-06 | 5.28E-06 |
| rs113708455 | 19 | 451089 | IMPUTED | SHC2 | T | C | 0.00036392 | 3 | 30.2344702 | 6.6251228 | 5.03E-06 | 5.29E-06 |
| rs112335970 | 19 | 451158 | IMPUTED | SHC2 | T | C | 0.0003638 | 3 | 30.2338296 | 6.62511959 | 5.03E-06 | 5.29E-06 |
| rs115064953 | 19 | 451261 | IMPUTED | SHC2 | C | T | 0.0003638 | 3 | 30.2338296 | 6.62511959 | 5.03E-06 | 5.29E-06 |
| rs540445501 | 2 | 52666399 | IMPUTED | LOC730100(dist=31344),MIR4431(dist=263261) | T | A | 0.00035833 | 3 | 30.2361812 | 6.62675766 | 5.05E-06 | 5.31E-06 |
| rs112680260 | 19 | 450067 | IMPUTED | SHC2 | T | C | 0.0003638 | 3 | 30.2278599 | 6.62540936 | 5.06E-06 | 5.32E-06 |
| rs79621546 | 19 | 452976 | IMPUTED | SHC2 | G | A | 0.00037057 | 3 | 30.2228457 | 6.62450253 | 5.06E-06 | 5.32E-06 |
| rs112068208 | 19 | 451007 | IMPUTED | SHC2 | G | A | 0.00036392 | 3 | 30.2250514 | 6.62510975 | 5.06E-06 | 5.32E-06 |
| rs145228918 | 21 | 31124413 | IMPUTED | GRIK1-AS1 | G | A | 0.00040064 | 3 | 30.1367771 | 6.60587807 | 5.06E-06 | 5.33E-06 |
| rs368783350 | 2 | 52928992 | IMPUTED | MIR4431(dist=668) | T | C | 0.00036154 | 3 | 30.2171422 | 6.62425942 | 5.08E-06 | 5.34E-06 |
| rs367596224 | 11 | 116707929 | IMPUTED | APOA1-AS | G | A | 0.0003632 | 3 | 30.6624223 | 6.72193215 | 5.08E-06 | 5.34E-06 |
| rs549967454 | 1 | 240778266 | IMPUTED | MIR1273E | G | A | 0.00037366 | 3 | 30.2147875 | 6.62476154 | 5.09E-06 | 5.36E-06 |
| rs192239501 | 15 | 64466050 | IMPUTED | CSNK1G1 | T | C | 0.00820787 | 69 | -6.2133839 | 1.36238516 | 5.10E-06 | 5.36E-06 |
| rs76821818 | 19 | 439566 | IMPUTED | SHC2 | C | T | 0.00036522 | 3 | 30.2831704 | 6.64051933 | 5.11E-06 | 5.37E-06 |
| rs373143962 | 2 | 52960571 | IMPUTED | MIR4431(dist=30818),ASB3(dist=936546) | T | C | 0.00036178 | 3 | 30.2168928 | 6.62658818 | 5.12E-06 | 5.38E-06 |
| rs368598615 | 2 | 52930117 | IMPUTED | MIR4431(dist=364) | G | A | 0.00036202 | 3 | 30.2038656 | 6.62376252 | 5.12E-06 | 5.38E-06 |
| rs543812413 | 2 | 52668274 | IMPUTED | LOC730100(dist=33219),MIR4431(dist=261386) | A | G | 0.00035845 | 3 | 30.2170621 | 6.62671988 | 5.12E-06 | 5.38E-06 |
| rs546540293 | 17 | 54482918 | IMPUTED | ANKFN1 | T | A | 0.00142608 | 12 | 16.2366447 | 3.56083589 | 5.12E-06 | 5.38E-06 |
| rs557798528 | 2 | 52672521 | IMPUTED | LOC730100(dist=37466),MIR4431(dist=257139) | A | C | 0.00035809 | 3 | 30.2117027 | 6.62671025 | 5.14E-06 | 5.40E-06 |
| rs566637930 | 2 | 52672633 | IMPUTED | LOC730100(dist=37578),MIR4431(dist=257027) | G | A | 0.00035809 | 3 | 30.2117027 | 6.62671025 | 5.14E-06 | 5.40E-06 |
| rs145513604 | 1 | 186629504 | IMPUTED | LOC102724919(dist=190081),PTGS2(dist=11440) | T | C | 0.00084359 | 7 | -19.738398 | 4.32963694 | 5.14E-06 | 5.41E-06 |
| rs149566183 | 17 | 30045735 | IMPUTED | MIR365B(dist=143195),COPRS(dist=133149) | G | A | 0.00037271 | 3 | 30.1964999 | 6.62430915 | 5.15E-06 | 5.42E-06 |
| rs150202361 | 2 | 52704153 | IMPUTED | LOC730100(dist=69098),MIR4431(dist=225507) | A | G | 0.0003676 | 3 | 30.1706471 | 6.61905977 | 5.16E-06 | 5.43E-06 |
| rs550698104 | 2 | 52670937 | IMPUTED | LOC730100(dist=35882),MIR4431(dist=258723) | T | C | 0.00035821 | 3 | 30.2023171 | 6.62668969 | 5.17E-06 | 5.44E-06 |
| rs750101917 | 18 | 39248855 | IMPUTED | KC6(dist=148294),PIK3C3(dist=286308) | G | A | 0.00025184 | 2 | -33.171313 | 7.278477 | 5.18E-06 | 5.44E-06 |
| rs542073652 | 1 | 104856346 | IMPUTED | LOC100129138(dist=236653),LINC01676(dist=1275970) | T | G | 0.00043725 | 4 | 29.7776132 | 6.53389161 | 5.18E-06 | 5.45E-06 |
| rs193213051 | 2 | 53000443 | IMPUTED | MIR4431(dist=70690),ASB3(dist=896674) | G | A | 0.00036285 | 3 | 30.1939328 | 6.62560909 | 5.19E-06 | 5.45E-06 |
| rs146369142 | 5 | 180621561 | IMPUTED | TRIM7(NM_203293:c.*6050>0,NM_203295:c.*6050>0,NM_ | C | T | 0.00038388 | 3 | 30.1494873 | 6.61676035 | 5.20E-06 | 5.47E-06 |
| rs568953477 | 2 | 52671022 | IMPUTED | LOC730100(dist=35967),MIR4431(dist=258638) | G | A | 0.00035845 | 3 | 30.1933572 | 6.62667433 | 5.21E-06 | 5.47E-06 |
| rs79252187 | 11 | 36074245 | IMPUTED | LDLRAD3 | C | T | 0.05562408 | 468 | -2.3256672 | 0.51051639 | 5.23E-06 | 5.49E-06 |
| rs1227101 | 2 | 148525896 | IMPUTED | PABPC1P2(dist=1177338),ACVR2A(dist=76190) | T | C | 0.99999941 | | 16674.6132 | 3660.40865 | 5.23E-06 | |

| | | | | | | | | | | | | |
|---|---|---|---|---|---|---|---|---|---|---|---|---|
| rs986773683 | 6 | 43120291 | IMPUTED | PTK7 | C | T | 0.00024162 | 2 | -33.273942 | 7.30433932 | 5.23E-06 | 5.50E-06 |
| rs749650930 | 12 | 102111320 | IMPUTED | CHPT1 | C | T | 0.00012646 | 1 | 47.1269193 | 10.3458063 | 5.23E-06 | 5.50E-06 |
| rs550436085 | 8 | 136090272 | IMPUTED | LOC101927845(dist=196130),LINC01591(dist=156102) | C | T | 0.00024115 | 2 | -33.254378 | 7.30074068 | 5.24E-06 | 5.51E-06 |
| rs527958282 | 6 | 101572582 | IMPUTED | ASCC3(dist=243334),GRIK2(dist=274279) | A | T | 0.00023806 | 2 | -33.118117 | 7.27165739 | 5.25E-06 | 5.52E-06 |
| rs757096891 | 18 | 39379864 | IMPUTED | KC6(dist=279303),PIK3C3(dist=155299) | T | C | 0.00023461 | 2 | -33.819617 | 7.42764018 | 5.28E-06 | 5.55E-06 |
| rs749433443 | 6 | 170870119 | IMPUTED | TBP | G | A | 0.00026004 | 2 | -33.898265 | 7.44505741 | 5.29E-06 | 5.56E-06 |
| rs766229253 | 6 | 55521755 | IMPUTED | HMGCLL1(dist=77743),BMP5(dist=96696) | T | C | 0.00023699 | 2 | -33.221908 | 7.29722763 | 5.30E-06 | 5.57E-06 |
| rs73277212 | 8 | 82451471 | IMPUTED | FABP12(dist=7846),IMPA1P1(dist=64648) | T | A | 0.0002875 | 2 | -31.995011 | 7.0279443 | 5.30E-06 | 5.57E-06 |
| rs7251996 | 19 | 482288 | IMPUTED | ODF3L2(dist=7305),MADCAM1(dist=14202) | T | C | 0.00032648 | 3 | 32.8144809 | 7.20902477 | 5.32E-06 | 5.59E-06 |
| rs530214521 | 2 | 52499703 | IMPUTED | LOC730100 | G | T | 0.00034454 | 3 | 31.2619396 | 6.86828784 | 5.32E-06 | 5.60E-06 |
| rs756788725 | 1 | 75202091 | IMPUTED | TYW3 | G | A | 0.00024566 | 2 | -33.126694 | 7.27855743 | 5.33E-06 | 5.61E-06 |
| rs190859994 | 1 | 215169142 | IMPUTED | CENPF(dist=331228),KCNK2(dist=9743) | T | C | 0.00063917 | 5 | -22.05965 | 4.8469839 | 5.33E-06 | 5.61E-06 |
| rs565654994 | 3 | 2176571 | IMPUTED | CNTN4-AS2 | C | A | 0.00036974 | 3 | 30.2018575 | 6.63604018 | 5.33E-06 | 5.61E-06 |
| rs374739658 | 2 | 52933597 | IMPUTED | MIR4431(dist=3844),ASB3(dist=963520) | A | G | 0.00036475 | 3 | 30.1280739 | 6.62021133 | 5.34E-06 | 5.61E-06 |
| rs370658532 | 2 | 53108117 | IMPUTED | MIR4431(dist=178364),ASB3(dist=789000) | G | A | 0.00036237 | 3 | 30.1422274 | 6.62354728 | 5.35E-06 | 5.62E-06 |
| rs114112804 | 4 | 184756015 | IMPUTED | NONE(dist=NONE),NONE(dist=NONE) | G | A | 0.00082517 | 7 | -19.573704 | 4.30165239 | 5.36E-06 | 5.63E-06 |
| rs559783246 | 2 | 52499447 | IMPUTED | LOC730100 | G | A | 0.00034407 | 3 | 31.2811584 | 6.87477939 | 5.36E-06 | 5.64E-06 |
| rs373411394 | 2 | 52934689 | IMPUTED | MIR4431(dist=4936),ASB3(dist=962428) | A | C | 0.00036511 | 3 | 30.1161433 | 6.6196211 | 5.38E-06 | 5.65E-06 |
| rs899857383 | 17 | 30843917 | IMPUTED | MYO1D | G | A | 0.00037117 | 3 | 30.2913737 | 6.65923841 | 5.40E-06 | 5.67E-06 |
| rs185758937 | 2 | 59958337 | IMPUTED | LINC01793(dist=451802),MIR4432HG(dist=628014) | C | G | 0.00055384 | 5 | 25.0762981 | 5.51417164 | 5.43E-06 | 5.70E-06 |
| rs535362015 | 12 | 97739345 | IMPUTED | NEDD1(dist=391876),RMST(dist=119454) | T | C | 0.00035488 | 3 | 30.9330756 | 6.80207793 | 5.43E-06 | 5.70E-06 |
| rs7972335 | 12 | 60144966 | IMPUTED | SLC16A7 | C | T | 0.0004029 | 3 | 30.0029672 | 6.59790692 | 5.43E-06 | 5.71E-06 |
| rs181380052 | 2 | 59958310 | IMPUTED | LINC01793(dist=451775),MIR4432HG(dist=628041) | A | G | 0.00055384 | 5 | 25.0740957 | 5.51444614 | 5.44E-06 | 5.72E-06 |
| rs113579383 | 9 | 90137915 | IMPUTED | DAPK1 | G | A | 0.00037782 | 3 | 30.158936 | 6.63324687 | 5.45E-06 | 5.73E-06 |
| rs183898472 | 2 | 96840725 | IMPUTED | DUSP2(dist=29546),STARD7(dist=9878) | C | T | 0.00188733 | 16 | 12.3824137 | 2.72349862 | 5.45E-06 | 5.73E-06 |
| rs564375608 | 2 | 52498780 | IMPUTED | LOC730100 | T | A | 0.00034252 | 3 | 31.4107697 | 6.90885018 | 5.46E-06 | 5.73E-06 |
| rs55840820 | 13 | 80450978 | IMPUTED | LINC00382 | T | C | 0.00843487 | 71 | 6.14249146 | 1.35124799 | 5.47E-06 | 5.75E-06 |
| rs114934434 | 2 | 137252609 | IMPUTED | CXCR4(dist=376884),THSD7B(dist=270523) | T | C | 0.00024222 | 2 | -33.383997 | 7.34500027 | 5.49E-06 | 5.77E-06 |
| rs948160859 | 18 | 71588667 | IMPUTED | LOC100505817(dist=571543),FBXO15(dist=151921) | G | A | 0.00053007 | 4 | 24.9194605 | 5.48293131 | 5.50E-06 | 5.78E-06 |
| rs545765809 | 2 | 52498530 | IMPUTED | LOC730100 | T | C | 0.00034181 | 3 | 31.4520187 | 6.92030766 | 5.50E-06 | 5.78E-06 |
| rs540848381 | 2 | 52571977 | IMPUTED | LOC730100 | A | T | 0.00036641 | 3 | 30.1072549 | 6.6245601 | 5.50E-06 | 5.78E-06 |
| rs114306868 | 2 | 16534049 | GENOTYPED | GACAT3(dist=308238),FAM49A(dist=196681) | C | A | 0.00023794 | 2 | -31.970494 | 7.03654546 | 5.53E-06 | 5.81E-06 |
| rs6791206 | 3 | 61236328 | IMPUTED | FHIT | A | G | 0.27760245 | 2336 | -1.1798026 | 0.25970177 | 5.55E-06 | 5.83E-06 |
| rs757283490 | 4 | 182401877 | IMPUTED | LINC00290(dist=321575),TEMN3-AS1(dist=339281) | T | C | 0.00031519 | 3 | 33.9499731 | 7.47385738 | 5.56E-06 | 5.84E-06 |
| rs778133878 | 16 | 74059073 | IMPUTED | LINC01568(dist=603778),LOC101928035(dist=167218) | C | A | 0.00041835 | 4 | -26.853251 | 5.9116694 | 5.56E-06 | 5.84E-06 |
| rs28693834 | 12 | 60179829 | IMPUTED | SLC16A7(NM_001270623:c.*63690>0,NM_001270622:c.*6 | G | A | 0.00037034 | 3 | 30.0912285 | 6.62450539 | 5.56E-06 | 5.84E-06 |
| rs531720237 | 2 | 52703591 | IMPUTED | LOC730100(dist=68536),MIR4431(dist=226069) | C | T | 0.00037295 | 3 | 30.0138036 | 6.60790028 | 5.57E-06 | 5.85E-06 |
| rs78660840 | 2 | 16583501 | GENOTYPED | GACAT3(dist=357690),FAM49A(dist=147229) | C | T | 0.0002377 | 2 | -31.957888 | 7.03654483 | 5.58E-06 | 5.86E-06 |
| rs115823749 | 2 | 16532294 | IMPUTED | GACAT3(dist=306483),FAM49A(dist=198436) | T | A | 0.00023806 | 2 | -31.956881 | 7.03653579 | 5.58E-06 | 5.87E-06 |
| rs775820042 | 2 | 19678882 | IMPUTED | OSR1(dist=120510),LINC00954(dist=389733) | C | A | 0.00024269 | 2 | -33.015234 | 7.27031481 | 5.60E-06 | 5.88E-06 |
| rs556737465 | 2 | 52497944 | IMPUTED | LOC730100 | G | A | 0.00033991 | 3 | 31.5759089 | 6.95402087 | 5.61E-06 | 5.89E-06 |
| rs542666267 | 9 | 37663948 | IMPUTED | FRMPD1 | C | T | 0.00034169 | 3 | -30.611504 | 6.74179569 | 5.61E-06 | 5.89E-06 |
| rs114248084 | 19 | 450056 | IMPUTED | SHC2 | G | A | 0.00037188 | 3 | 30.0431756 | 6.61770902 | 5.63E-06 | 5.92E-06 |
| rs375877370 | 1 | 75211488 | IMPUTED | TYW3 | A | T | 0.00025137 | 2 | -33.036808 | 7.2779091 | 5.64E-06 | 5.93E-06 |
| rs566408803 | 1 | 80702723 | IMPUTED | ADGRL4(dist=1230228),LINC01781(dist=298717) | G | T | 0.00024305 | 2 | -35.31559 | 7.7801451 | 5.65E-06 | 5.93E-06 |
| rs749746138 | 9 | 120852171 | IMPUTED | TLR4(dist=372402),BRINP1(dist=1076737) | T | A | 0.00036891 | 3 | 30.249422 | 6.66417705 | 5.65E-06 | 5.94E-06 |
| rs372492838 | 2 | 52710357 | IMPUTED | LOC730100(dist=75302),MIR4431(dist=219303) | A | G | 0.00041276 | 3 | 29.7891194 | 6.56397775 | 5.67E-06 | 5.96E-06 |
| rs144759232 | 12 | 40714394 | IMPUTED | LRRK2 | G | A | 0.00026254 | 2 | -33.816424 | 7.45235554 | 5.69E-06 | 5.97E-06 |
| rs114605485 | 2 | 16521419 | IMPUTED | GACAT3(dist=295608),FAM49A(dist=209311) | A | T | 0.00023853 | 2 | -31.923436 | 7.03654235 | 5.71E-06 | 6.00E-06 |
| rs574154145 | 2 | 52496962 | IMPUTED | LOC730100 | A | C | 0.00033777 | 3 | 31.7166142 | 6.99303182 | 5.75E-06 | 6.04E-06 |
| rs1052639176 | 7 | 83658598 | IMPUTED | SEMA3A | C | G | 0.00030402 | 3 | -33.143383 | 7.30799294 | 5.75E-06 | 6.04E-06 |
| rs374460855 | 2 | 53108073 | IMPUTED | MIR4431(dist=178320),ASB3(dist=789044) | G | C | 0.00038139 | 3 | 30.0263082 | 6.62211144 | 5.78E-06 | 6.07E-06 |

| | | | | | | | | | | | | |
|---|---|---|---|---|---|---|---|---|---|---|---|---|
| rs549411482 | 16 | 65031164 | IMPUTED | CDH11 | C | T | 0.00025434 | 2 | -32.912155 | 7.25922356 | 5.79E-06 | 6.08E-06 |
| rs916860201 | 6 | 53400634 | IMPUTED | GCLC | T | C | 0.00050856 | 4 | 25.2356824 | 5.5666119 | 5.80E-06 | 6.10E-06 |
| rs141588875 | 9 | 112054565 | IMPUTED | EPB41L4B | C | T | 0.00924935 | 78 | -5.587597 | 1.23291595 | 5.84E-06 | 6.14E-06 |
| rs114938410 | 4 | 164722318 | IMPUTED | 1-Mar | G | T | 3.09E-06 | | 4105.26668 | 905.88683 | 5.85E-06 | |
| rs62460338 | 7 | 68602418 | IMPUTED | LOC102723427(dist=1104741),LOC100507468(dist=458706 | A | T | 0.09684407 | 815 | -1.8842021 | 0.4157816 | 5.85E-06 | 6.14E-06 |
| rs181788788 | 2 | 97464013 | IMPUTED | CNNM4 | C | A | 0.00440492 | 37 | 8.59679547 | 1.89718258 | 5.86E-06 | 6.16E-06 |
| rs547801833 | 11 | 117313688 | IMPUTED | DSCAML1 | G | C | 0.00079962 | 7 | 18.8278949 | 4.15660832 | 5.91E-06 | 6.21E-06 |
| rs115300716 | 2 | 63729942 | IMPUTED | WDPCP | A | T | 0.00335417 | 28 | -10.229793 | 2.25860013 | 5.92E-06 | 6.22E-06 |
| rs73489616 | 19 | 450644 | IMPUTED | SHC2 | G | A | 0.00037188 | 3 | 29.9896419 | 6.62264676 | 5.94E-06 | 6.24E-06 |
| rs17042683 | 2 | 52510537 | IMPUTED | LOC730100 | A | G | 0.00036106 | 3 | 30.0059974 | 6.62705996 | 5.96E-06 | 6.26E-06 |
| rs184772703 | 2 | 55797387 | IMPUTED | PPP4R3B | A | G | 0.0003827 | 3 | 30.1437956 | 6.65778559 | 5.97E-06 | 6.26E-06 |
| rs550145157 | 3 | 1874208 | IMPUTED | CNTN6(dist=428916),CNTN4(dist=266279) | G | T | 0.00037842 | 3 | 30.0200298 | 6.63135028 | 5.98E-06 | 6.28E-06 |
| rs374633957 | 2 | 53042776 | IMPUTED | MIR4431(dist=113023),ASB3(dist=854341) | T | C | 0.00036736 | 3 | 29.9488347 | 6.61681525 | 6.01E-06 | 6.31E-06 |
| rs532274002 | 11 | 103845616 | IMPUTED | PDGFD | T | C | 0.00162218 | 14 | 12.4658838 | 2.75430438 | 6.01E-06 | 6.31E-06 |
| rs988902039 | 4 | 184718514 | IMPUTED | TRAPPC11(dist=83767),NONE(dist=NONE) | C | T | 0.00061683 | 5 | -21.759863 | 4.80939199 | 6.06E-06 | 6.36E-06 |
| rs74090138 | 14 | 105432498 | IMPUTED | AHNAK2 | G | T | 0.00036843 | 3 | 27.2527553 | 6.02451555 | 6.08E-06 | 6.38E-06 |
| rs76744615 | 2 | 53010207 | IMPUTED | MIR4431(dist=80454),ASB3(dist=886910) | G | A | 0.00038781 | 3 | 29.9456812 | 6.61988673 | 6.08E-06 | 6.38E-06 |
| rs74090139 | 14 | 105432549 | IMPUTED | AHNAK2 | C | T | 0.0003682 | 3 | 27.2515161 | 6.0249143 | 6.09E-06 | 6.40E-06 |
| rs181094857 | 18 | 2758416 | IMPUTED | SMCHD1 | C | T | 0.00070383 | 6 | 19.9967316 | 4.42107816 | 6.10E-06 | 6.40E-06 |
| rs941006524 | 1 | 196040073 | IMPUTED | LINC01724 | G | A | 0.00081959 | 7 | 20.4683024 | 4.52550452 | 6.10E-06 | 6.40E-06 |
| rs534584559 | 2 | 52494745 | IMPUTED | LOC730100 | A | C | 0.00033409 | 3 | 31.9984405 | 7.07539266 | 6.11E-06 | 6.42E-06 |
| rs12425587 | 12 | 86003568 | IMPUTED | ALX1(dist=308007),RASSF9(dist=194763) | A | C | 0.00045769 | 4 | 27.886765 | 6.16673498 | 6.12E-06 | 6.43E-06 |
| rs561942096 | 6 | 41992649 | IMPUTED | CCND3 | A | T | 0.0002421 | 2 | -32.873527 | 7.2714585 | 6.16E-06 | 6.46E-06 |
| rs189929690 | 18 | 2788624 | IMPUTED | SMCHD1 | A | G | 0.00070062 | 6 | 20.0702371 | 4.44092871 | 6.20E-06 | 6.51E-06 |
| rs73820011 | 3 | 24636476 | IMPUTED | MIR4792(dist=73550),RARB(dist=234338) | C | T | 0.00166401 | 14 | 12.7318305 | 2.81856114 | 6.27E-06 | 6.58E-06 |
| rs563650515 | 7 | 20113634 | IMPUTED | LOC101927668 | G | A | 0.00045888 | 4 | 25.7922794 | 5.70996372 | 6.27E-06 | 6.58E-06 |
| rs376620740 | 2 | 52989466 | IMPUTED | MIR4431(dist=59713),ASB3(dist=907651) | A | G | 0.00043689 | 4 | 28.2582756 | 6.25680691 | 6.29E-06 | 6.60E-06 |
| rs188219563 | 8 | 99163374 | IMPUTED | POP1 | C | T | 0.00022879 | 2 | -34.956823 | 7.74277162 | 6.34E-06 | 6.65E-06 |
| rs28501597 | 18 | 73234731 | IMPUTED | SMIM21(dist=95073),LINC01898(dist=173307) | A | G | 0.07499917 | 631 | -1.9878269 | 0.44055391 | 6.42E-06 | 6.74E-06 |
| rs781835964 | 1 | 147138901 | IMPUTED | ACP6,NBPF19 | G | A | 0.00039993 | 3 | -28.510524 | 6.31932224 | 6.43E-06 | 6.75E-06 |
| rs6884393 | 5 | 171272310 | IMPUTED | SMIM23(dist=54218),FBXW11(dist=16246) | A | C | 0.00029724 | 3 | -31.285007 | 6.9352846 | 6.45E-06 | 6.77E-06 |
| rs537133298 | 2 | 54555797 | IMPUTED | ACYP2(dist=23362),C2orf73(dist=2274) | T | C | 0.00037307 | 3 | 29.9257504 | 6.63436914 | 6.46E-06 | 6.78E-06 |
| rs61759787 | 14 | 105251426 | IMPUTED | AKT1 | C | T | 0.00037366 | 3 | 27.2328114 | 6.03748798 | 6.46E-06 | 6.78E-06 |
| rs111915241 | 6 | 167000370 | IMPUTED | RPS6KA2 | C | G | 0.00119907 | 10 | 16.8575807 | 3.7373948 | 6.47E-06 | 6.79E-06 |
| rs142065631 | 14 | 105249989 | IMPUTED | AKT1 | G | A | 0.00037152 | 3 | 27.2618072 | 6.04509964 | 6.49E-06 | 6.81E-06 |
| rs182347375 | 5 | 16439388 | IMPUTED | LINC02150 | G | A | 0.00026812 | 2 | -33.874245 | 7.51213082 | 6.51E-06 | 6.83E-06 |
| rs57829652 | 6 | 21272089 | IMPUTED | CDKAL1(dist=39455),LINC00581(dist=214203) | A | T | 0.00027193 | 2 | -32.163438 | 7.13326793 | 6.52E-06 | 6.84E-06 |
| rs34072234 | 13 | 30091775 | IMPUTED | SLC7A1 | G | A | 0.00117459 | 10 | -15.729179 | 3.48876973 | 6.53E-06 | 6.85E-06 |
| rs537333512 | 2 | 52493569 | IMPUTED | LOC730100 | G | T | 0.0003304 | 3 | 32.2027205 | 7.1433683 | 6.54E-06 | 6.86E-06 |
| rs114106530 | 5 | 171273376 | IMPUTED | SMIM23(dist=55284),FBXW11(dist=15180) | G | A | 0.000297 | 2 | -31.264527 | 6.93525672 | 6.54E-06 | 6.86E-06 |
| rs533086661 | 2 | 54593020 | IMPUTED | C2orf73(dist=4306),SPTBN1(dist=90434) | T | C | 0.00037414 | 3 | 29.896945 | 6.63281533 | 6.56E-06 | 6.88E-06 |
| rs751978714 | 8 | 42508224 | IMPUTED | SMIM19(dist=100084),CHRNB3(dist=44295) | C | T | 0.00026741 | 2 | 31.5723375 | 7.0069797 | 6.61E-06 | 6.94E-06 |
| rs906391415 | 5 | 173413458 | IMPUTED | CPEB4(dist=25464),C5orf47(dist=2704) | G | A | 0.00037236 | 3 | 29.764338 | 6.60580008 | 6.61E-06 | 6.94E-06 |
| rs140040390 | 13 | 54348781 | IMPUTED | LINC01065(dist=622746),LINC00558(dist=40773) | C | T | 0.0008077 | 7 | -19.660027 | 4.36421419 | 6.64E-06 | 6.97E-06 |
| rs10912126 | 1 | 187636974 | IMPUTED | LINC01037(dist=190620),NONE(dist=NONE) | G | A | 0.00026254 | 2 | -32.741201 | 7.26901643 | 6.66E-06 | 6.99E-06 |
| rs533516434 | 21 | 38066225 | IMPUTED | CLDN14(dist=117358),SIM2(dist=5196) | G | A | 0.00037081 | 3 | 29.83304 | 6.62347671 | 6.66E-06 | 6.99E-06 |
| rs146689853 | 1 | 187632912 | IMPUTED | LINC01037(dist=186558),NONE(dist=NONE) | T | C | 0.00026242 | 2 | -32.739196 | 7.2690199 | 6.67E-06 | 7.00E-06 |
| rs542688940 | 12 | 59661267 | IMPUTED | LRIG3(dist=346948),SLC16A7(dist=328554) | C | A | 0.000401 | 3 | 30.4017304 | 6.75040361 | 6.68E-06 | 7.01E-06 |
| rs112298944 | 4 | 110742801 | IMPUTED | GAR1 | G | A | 0.00041764 | 4 | 28.291726 | 6.28221121 | 6.69E-06 | 7.01E-06 |
| rs111378532 | 13 | 93204033 | IMPUTED | GPC5 | A | G | 0.05790231 | 487 | -2.2742418 | 0.50505329 | 6.70E-06 | 7.03E-06 |
| rs757504941 | 18 | 38713901 | IMPUTED | LINC01477(dist=1034704),KC6(dist=346335) | A | G | 0.00020977 | 2 | -38.28546 | 8.50304667 | 6.71E-06 | 7.04E-06 |
| rs112543146 | 2 | 52517531 | IMPUTED | LOC730100 | T | G | 0.00036618 | 3 | 29.8274883 | 6.62550114 | 6.73E-06 | 7.06E-06 |

| | | | | | | | | | | | | |
|---|---|---|---|---|---|---|---|---|---|---|---|---|
| rs780529937 | 8 | 80843646 | IMPUTED | MRPS28 | T | C | 0.00027704 | 2 | -33.651893 | 7.47585191 | 6.75E-06 | 7.08E-06 |
| rs758723783 | 9 | 35953589 | IMPUTED | SPAAR(dist=41972),OR2S2(dist=3516) | T | C | 0.00023758 | 2 | -34.569391 | 7.68015813 | 6.76E-06 | 7.09E-06 |
| rs73336159 | 5 | 176137439 | IMPUTED | TSPAN17(dist=51380),LINC01574(dist=32767) | C | T | 0.0005246 | 4 | 24.3279946 | 5.40539579 | 6.77E-06 | 7.10E-06 |
| rs145866128 | 19 | 450803 | IMPUTED | SHC2 | G | A | 0.00046803 | 4 | 29.3848737 | 6.52979819 | 6.79E-06 | 7.12E-06 |
| rs150095180 | 12 | 87851874 | IMPUTED | LOC105369879(dist=125776),MKRN9P(dist=324789) | G | A | 0.00744331 | 63 | -6.8279447 | 1.51753712 | 6.82E-06 | 7.15E-06 |
| rs116991706 | 9 | 5618386 | IMPUTED | PDCD1LG2(dist=47104),RIC1(dist=10733) | A | G | 0.0057995 | 49 | 6.86101194 | 1.5252007 | 6.85E-06 | 7.18E-06 |
| rs115494049 | 8 | 82649057 | IMPUTED | CHMP4C | G | C | 0.00023092 | 2 | -34.019227 | 7.56320853 | 6.86E-06 | 7.19E-06 |
| rs118088040 | 10 | 112508087 | GENOTYPED | RBM20 | C | T | 0.02253696 | 190 | -3.5338673 | 0.78568375 | 6.87E-06 | 7.20E-06 |
| rs73916585 | 2 | 16514796 | IMPUTED | GACAT3(dist=288985),FAM49A(dist=215934) | T | A | 0.00035809 | 3 | -25.925019 | 5.7648655 | 6.89E-06 | 7.22E-06 |
| rs183914121 | 3 | 117437693 | IMPUTED | LINC02024(dist=27921),LOC105374060(dist=789565) | G | A | 0.00460483 | 39 | -7.7461274 | 1.72290892 | 6.93E-06 | 7.26E-06 |
| rs13377557 | 11 | 36077902 | GENOTYPED | LDLRAD3 | T | G | 0.05574507 | 469 | -2.2926291 | 0.50994308 | 6.93E-06 | 7.27E-06 |
| rs562098716 | 17 | 43278078 | IMPUTED | LOC339192 | C | T | 0.00350856 | 30 | 9.17360705 | 2.04096225 | 6.97E-06 | 7.30E-06 |
| rs80180234 | 2 | 16520328 | IMPUTED | GACAT3(dist=294517),FAM49A(dist=210402) | A | G | 0.00035857 | 3 | -25.916289 | 5.76674143 | 6.99E-06 | 7.33E-06 |
| rs1039113713 | 12 | 97055973 | IMPUTED | CFAP54 | A | C | 0.00038602 | 3 | 29.8105429 | 6.6334185 | 6.99E-06 | 7.33E-06 |
| rs144694633 | 8 | 82569926 | IMPUTED | IMPA1(NM_001144878:c.*16600>0,NM_005536:c.*16600> | G | A | 0.00023164 | 2 | -33.883274 | 7.54101751 | 7.02E-06 | 7.36E-06 |
| rs73217694 | 2 | 16519533 | IMPUTED | GACAT3(dist=293722),FAM49A(dist=211197) | A | G | 0.00035869 | 3 | -25.908458 | 5.76675639 | 7.03E-06 | 7.37E-06 |
| rs571970220 | 5 | 66483273 | IMPUTED | CD180 | C | T | 0.00024531 | 2 | -33.013631 | 7.35175813 | 7.10E-06 | 7.45E-06 |
| rs1015156934 | 6 | 36055015 | IMPUTED | MAPK14 | C | T | 0.00013002 | 1 | -46.740064 | 10.4089993 | 7.11E-06 | 7.45E-06 |
| rs574282721 | 17 | 54544365 | IMPUTED | ANKFN1 | C | A | 0.0013184 | 11 | 16.7470382 | 3.73046295 | 7.15E-06 | 7.49E-06 |
| rs373182914 | 2 | 52556877 | IMPUTED | LOC730100 | C | T | 0.0005183 | 4 | 25.6584439 | 5.71782844 | 7.21E-06 | 7.56E-06 |
| rs141930236 | 8 | 82239070 | IMPUTED | FABP5(dist=42058),PMP2(dist=113493) | G | A | 0.00023342 | 2 | -34.042432 | 7.58795641 | 7.24E-06 | 7.59E-06 |
| rs540661404 | 5 | 58204331 | IMPUTED | RAB3C(dist=49109),PDE4D(dist=60535) | T | C | 0.00035512 | 3 | 27.2557279 | 6.07556792 | 7.25E-06 | 7.60E-06 |
| rs574682246 | 12 | 63410783 | IMPUTED | PPM1H(dist=82118),AVPR1A(dist=125756) | G | A | 0.00036843 | 3 | 29.6775817 | 6.61552712 | 7.26E-06 | 7.61E-06 |
| rs777593453 | 18 | 46245088 | IMPUTED | CTIF | G | A | 0.00028393 | 2 | -32.350115 | 7.21405909 | 7.31E-06 | 7.67E-06 |
| rs144646239 | 18 | 54126954 | IMPUTED | LINC01539(dist=322187),TXNL1(dist=143099) | C | T | 0.00031008 | 3 | -33.358883 | 7.44007487 | 7.34E-06 | 7.69E-06 |
| rs189653044 | 8 | 82321312 | IMPUTED | FABP5(dist=124300),PMP2(dist=31251) | T | C | 0.00023734 | 2 | -33.438378 | 7.45975429 | 7.38E-06 | 7.73E-06 |
| rs960923972 | 15 | 89964468 | IMPUTED | MIR9-3HG(dist=22750),RHCG(dist=50172) | G | T | 0.00023984 | 2 | -35.430607 | 7.90901633 | 7.47E-06 | 7.83E-06 |
| rs141153583 | 8 | 82440058 | IMPUTED | FABP12 | G | T | 0.00023568 | 2 | -33.323027 | 7.44138545 | 7.53E-06 | 7.89E-06 |
| rs10166649 | 2 | 16520757 | IMPUTED | GACAT3(dist=294946),FAM49A(dist=209973) | T | C | 0.00037984 | 3 | -25.790952 | 5.76059173 | 7.57E-06 | 7.93E-06 |
| rs532500304 | 13 | 54361368 | IMPUTED | LINC01065(dist=635333),LINC00558(dist=28186) | T | C | 0.00080592 | 7 | -19.522871 | 4.36127299 | 7.59E-06 | 7.95E-06 |
| rs775026608 | 1 | 104104807 | IMPUTED | AMY2B | T | C | 0.00028405 | 2 | -32.850636 | 7.34019124 | 7.63E-06 | 7.99E-06 |
| rs76723802 | 3 | 24482538 | IMPUTED | THRB | T | C | 0.00087913 | 7 | 17.9113686 | 4.00240932 | 7.64E-06 | 8.00E-06 |
| rs190566429 | 8 | 82330221 | IMPUTED | FABP5(dist=133209),PMP2(dist=22342) | T | C | 0.00023437 | 2 | -33.36345 | 7.45626514 | 7.66E-06 | 8.02E-06 |
| rs73217696 | 2 | 16521354 | IMPUTED | GACAT3(dist=295543),FAM49A(dist=209376) | T | A | 0.00037996 | 3 | -25.774142 | 5.76060711 | 7.67E-06 | 8.04E-06 |
| rs75888469 | 19 | 449822 | IMPUTED | SHC2 | G | A | 0.00043261 | 4 | 29.4511926 | 6.58287295 | 7.68E-06 | 8.05E-06 |
| rs550387016 | 6 | 72846297 | IMPUTED | RIMS1 | C | T | 0.00024768 | 2 | -32.933161 | 7.36196525 | 7.70E-06 | 8.06E-06 |
| rs146838899 | 15 | 83749289 | IMPUTED | MIR4515(dist=13122),TM6SF1(dist=26949) | C | T | 0.00023104 | 2 | -33.972431 | 7.59463552 | 7.71E-06 | 8.07E-06 |
| rs551235699 | 2 | 45748144 | IMPUTED | SRBD1 | C | T | 0.00047896 | 4 | -24.277934 | 5.427481 | 7.71E-06 | 8.07E-06 |
| rs330083 | 8 | 9152654 | GENOTYPED | LOC101929128(dist=92288),LOC157273(dist=29907) | G | A | 0.94746589 | 442 | -2.3688943 | 0.52967739 | 7.74E-06 | 8.11E-06 |
| rs547943183 | 1 | 83271460 | IMPUTED | ADGRL2(dist=811844),LINC01362(dist=97406) | C | A | 0.00028999 | 2 | -31.867775 | 7.12657121 | 7.76E-06 | 8.13E-06 |
| rs145507590 | 16 | 84512201 | IMPUTED | TLDC1(NM_020947:c.*13180>0) | A | G | 0.00397338 | 33 | -8.1587668 | 1.82465177 | 7.77E-06 | 8.14E-06 |
| rs190253462 | 8 | 82874581 | IMPUTED | SNX16(dist=120060),LOC101927141(dist=949758) | G | A | 0.00024317 | 2 | -32.874947 | 7.35311264 | 7.79E-06 | 8.16E-06 |
| rs779966965 | 9 | 120744155 | IMPUTED | TLR4(dist=264386),BRINP1(dist=1184753) | A | G | 0.00037521 | 3 | 29.575494 | 6.61513017 | 7.79E-06 | 8.16E-06 |
| rs903168264 | 1 | 91913712 | IMPUTED | HFM1(dist=43286),CDC7(dist=52692) | G | C | 0.00037687 | 3 | 29.8552835 | 6.67781514 | 7.79E-06 | 8.16E-06 |
| rs73198481 | 8 | 15819676 | IMPUTED | TUSC3(dist=195518),MSR1(dist=145711) | A | T | 0.18310827 | 1541 | -1.3687093 | 0.3061995 | 7.82E-06 | 8.19E-06 |
| rs551518632 | 3 | 83063032 | IMPUTED | LINC02008(dist=550206),LINC00971(dist=1624524) | G | A | 0.00070858 | 6 | -20.062778 | 4.48884822 | 7.84E-06 | 8.21E-06 |
| rs150903802 | 8 | 82879135 | IMPUTED | SNX16(dist=124614),LOC101927141(dist=945204) | G | A | 0.00024222 | 2 | -32.854531 | 7.35318222 | 7.89E-06 | 8.27E-06 |
| rs369314523 | 2 | 52713930 | IMPUTED | LOC730100(dist=78875),MIR4431(dist=215730) | T | C | 0.00037913 | 3 | 29.4241496 | 6.58617424 | 7.91E-06 | 8.29E-06 |
| rs59841133 | 2 | 16498770 | IMPUTED | GACAT3(dist=272959),FAM49A(dist=231960) | C | T | 0.0002251 | 2 | -33.584574 | 7.51835321 | 7.93E-06 | 8.31E-06 |
| rs73275496 | 8 | 82443303 | IMPUTED | FABP12 | G | T | 0.0002396 | 2 | -32.842838 | 7.35312804 | 7.95E-06 | 8.33E-06 |
| rs115320746 | 1 | 162817025 | IMPUTED | HSD17B7(dist=34417),CCDC190(dist=7062) | G | A | 0.00024911 | 2 | -32.805235 | 7.34529141 | 7.96E-06 | 8.34E-06 |
| rs181970914 | 3 | 32430717 | IMPUTED | CMTM8(dist=18900),CMTM7(dist=2446) | G | T | 0.00448419 | 38 | 8.43089994 | 1.88784375 | 7.97E-06 | 8.35E-06 |

| | | | | | | | | | | | | |
|---|---|---|---|---|---|---|---|---|---|---|---|---|
| rs368114493 | 4 | 7940268 | IMPUTED | AFAP1 | G | C | 0.00090219 | 8 | 18.5408614 | 4.1517877 | 7.98E-06 | 8.36E-06 |
| rs563754974 | 7 | 20123324 | IMPUTED | LOC101927668 | A | G | 0.00040005 | 3 | 26.6771894 | 5.97531511 | 8.02E-06 | 8.40E-06 |
| rs774049907 | 4 | 63613948 | IMPUTED | ADGRL3-AS1(dist=586465),TECRL(dist=1530229) | C | A | 0.00047278 | 4 | 28.1258015 | 6.30006207 | 8.03E-06 | 8.41E-06 |
| rs922706564 | 12 | 22090721 | IMPUTED | ABCC9(dist=1093),CMAS(dist=108387) | C | A | 0.00049727 | 4 | 22.3075321 | 4.9972043 | 8.04E-06 | 8.42E-06 |
| rs534331977 | 12 | 67431608 | IMPUTED | GRIP1(dist=358683),LOC102724421(dist=39798) | T | A | 0.00048122 | 4 | 28.832642 | 6.45955101 | 8.06E-06 | 8.44E-06 |
| rs135440 | 22 | 44654739 | IMPUTED | KIAA1644 | C | G | 0.88486249 | 969 | -1.6934686 | 0.3794396 | 8.08E-06 | 8.46E-06 |
| rs372910642 | 1 | 66723172 | IMPUTED | PDE4B | C | T | 0.00028084 | 2 | -32.81245 | 7.35399689 | 8.13E-06 | 8.51E-06 |
| rs73217697 | 2 | 16521414 | IMPUTED | GACAT3(dist=295603),FAM49A(dist=209316) | T | C | 0.00038697 | 3 | -25.685064 | 5.75825724 | 8.17E-06 | 8.56E-06 |
| rs138513080 | 7 | 20119934 | IMPUTED | LOC101927668 | A | G | 0.00040397 | 3 | 26.4958832 | 5.94242503 | 8.24E-06 | 8.63E-06 |
| rs373619954 | 1 | 105045980 | IMPUTED | LOC100129138(dist=426287),LINC01676(dist=1086336) | A | G | 0.00024008 | 2 | -32.773821 | 7.3537693 | 8.32E-06 | 8.71E-06 |
| rs79990010 | 1 | 163489142 | IMPUTED | LOC100422212(dist=96161),PBX1(dist=1039455) | G | T | 0.00024126 | 2 | -32.912835 | 7.38649969 | 8.36E-06 | 8.75E-06 |
| rs9775241 | 9 | 140189845 | IMPUTED | TOR4A(dist=12752),NRARP(dist=4238) | T | G | 0.08617875 | 725 | 1.85734993 | 0.41689355 | 8.38E-06 | 8.77E-06 |
| rs558963078 | 6 | 83231861 | IMPUTED | TPBG(dist=154728),UBE3D(dist=370256) | G | T | 0.0004836 | 4 | 23.6510681 | 5.30905336 | 8.39E-06 | 8.79E-06 |
| rs144843801 | 4 | 110746431 | IMPUTED | GAR1(dist=538) | G | C | 0.0003991 | 3 | 28.7264161 | 6.44898761 | 8.41E-06 | 8.81E-06 |
| rs145351234 | 8 | 98900292 | IMPUTED | MATN2 | G | A | 0.00023889 | 2 | -32.755329 | 7.35375134 | 8.42E-06 | 8.81E-06 |
| rs554463 | 13 | 21525833 | IMPUTED | LINC00367(dist=2266),LATS2(dist=21343) | A | G | 0.99510221 | 41 | 7.48841758 | 1.68119737 | 8.42E-06 | 8.81E-06 |
| rs143839811 | 8 | 82780246 | IMPUTED | SNX16(dist=25725),LOC101927141(dist=1044093) | T | C | 0.00023734 | 2 | -32.801537 | 7.36477426 | 8.43E-06 | 8.83E-06 |
| rs73029638 | 1 | 163534200 | IMPUTED | LOC100422212(dist=141219),PBX1(dist=994397) | T | C | 0.00023913 | 2 | -32.745259 | 7.35356678 | 8.47E-06 | 8.86E-06 |
| rs75958227 | 11 | 116816591 | IMPUTED | SIK3 | G | A | 0.00039351 | 3 | 29.4077738 | 6.60407796 | 8.47E-06 | 8.86E-06 |
| rs532124872 | 6 | 72125965 | IMPUTED | LINC00472 | T | A | 0.00023841 | 2 | -32.744015 | 7.35382125 | 8.48E-06 | 8.88E-06 |
| rs561941741 | 1 | 96569567 | IMPUTED | LOC102723661(dist=81131),LINC01787(dist=150058) | T | C | 0.00050606 | 4 | 24.6389671 | 5.53361323 | 8.48E-06 | 8.88E-06 |
| rs7323729 | 13 | 21525247 | IMPUTED | LINC00367(dist=1680),LATS2(dist=21929) | C | T | 0.99509627 | 41 | 7.51016511 | 1.68674847 | 8.49E-06 | 8.89E-06 |
| rs185366171 | 7 | 29175181 | IMPUTED | CPVL | C | T | 0.00018541 | 2 | 38.2224585 | 8.58464283 | 8.49E-06 | 8.89E-06 |
| rs114632222 | 8 | 82745388 | IMPUTED | SNX16 | A | G | 0.00023758 | 2 | -32.756751 | 7.35748589 | 8.50E-06 | 8.90E-06 |
| rs141094782 | 8 | 99018996 | IMPUTED | MATN2 | T | G | 0.00023758 | 2 | -32.756751 | 7.35748589 | 8.50E-06 | 8.90E-06 |
| rs73277205 | 8 | 82447770 | IMPUTED | FABP12(dist=4145),IMPA1P1(dist=68349) | C | T | 0.00023782 | 2 | -32.740085 | 7.35385347 | 8.50E-06 | 8.90E-06 |
| rs150556028 | 8 | 82705793 | IMPUTED | CHMP4C(dist=34045),SNX16(dist=6025) | G | T | 0.00023782 | 2 | -32.740085 | 7.35385347 | 8.50E-06 | 8.90E-06 |
| rs193187075 | 8 | 99021713 | IMPUTED | MATN2 | A | G | 0.0002377 | 2 | -32.755501 | 7.35749161 | 8.51E-06 | 8.90E-06 |
| rs189119988 | 8 | 99056751 | IMPUTED | RPL30 | T | G | 0.00023782 | 2 | -32.754617 | 7.35748099 | 8.51E-06 | 8.91E-06 |
| rs115100734 | 1 | 162794207 | GENOTYPED | HSD17B7(dist=11599),CCDC190(dist=29880) | A | G | 0.00023782 | 2 | -32.736231 | 7.35383389 | 8.52E-06 | 8.92E-06 |
| rs115309715 | 1 | 162712336 | IMPUTED | DDR2 | A | G | 0.00023782 | 2 | -32.734638 | 7.35383062 | 8.53E-06 | 8.93E-06 |
| rs11484775 | 1 | 162719087 | IMPUTED | DDR2 | G | C | 0.00023782 | 2 | -32.734638 | 7.35383062 | 8.53E-06 | 8.93E-06 |
| rs73277204 | 8 | 82447200 | GENOTYPED | FABP12(dist=3575),IMPA1P1(dist=68919) | G | A | 0.0002377 | 2 | -32.734369 | 7.35384362 | 8.53E-06 | 8.93E-06 |
| rs188764513 | 8 | 82720924 | IMPUTED | SNX16 | G | A | 0.0002377 | 2 | -32.734369 | 7.35384362 | 8.53E-06 | 8.93E-06 |
| rs115571718 | 8 | 82783619 | IMPUTED | SNX16(dist=29098),LOC101927141(dist=1040720) | T | C | 0.0002377 | 2 | -32.734369 | 7.35384362 | 8.53E-06 | 8.93E-06 |
| rs7015194 | 8 | 82800296 | GENOTYPED | SNX16(dist=45775),LOC101927141(dist=1024043) | C | T | 0.0002377 | 2 | -32.734369 | 7.35384362 | 8.53E-06 | 8.93E-06 |
| rs148381942 | 8 | 82805168 | IMPUTED | SNX16(dist=50647),LOC101927141(dist=1019171) | T | C | 0.0002377 | 2 | -32.734369 | 7.35384362 | 8.53E-06 | 8.93E-06 |
| rs142735588 | 8 | 82807786 | IMPUTED | SNX16(dist=53265),LOC101927141(dist=1016553) | G | C | 0.0002377 | 2 | -32.734369 | 7.35384362 | 8.53E-06 | 8.93E-06 |
| rs143666707 | 8 | 98883291 | IMPUTED | MATN2 | A | C | 0.0002377 | 2 | -32.734369 | 7.35384362 | 8.53E-06 | 8.93E-06 |
| rs143416200 | 8 | 98889339 | IMPUTED | MATN2 | A | G | 0.0002377 | 2 | -32.734369 | 7.35384362 | 8.53E-06 | 8.93E-06 |
| rs183597429 | 8 | 98898180 | IMPUTED | MATN2 | C | G | 0.0002377 | 2 | -32.734369 | 7.35384362 | 8.53E-06 | 8.93E-06 |
| rs73278205 | 8 | 98948761 | IMPUTED | MATN2 | C | A | 0.0002377 | 2 | -32.734369 | 7.35384362 | 8.53E-06 | 8.93E-06 |
| rs148205164 | 8 | 98977355 | IMPUTED | MATN2 | C | A | 0.0002377 | 2 | -32.734369 | 7.35384362 | 8.53E-06 | 8.93E-06 |
| rs184388695 | 8 | 98982974 | IMPUTED | MATN2 | G | C | 0.0002377 | 2 | -32.734369 | 7.35384362 | 8.53E-06 | 8.93E-06 |
| rs187568831 | 8 | 98986701 | IMPUTED | MATN2 | G | A | 0.0002377 | 2 | -32.734369 | 7.35384362 | 8.53E-06 | 8.93E-06 |
| rs16974569 | 15 | 85399645 | GENOTYPED | ALPK3 | C | T | 0.0002377 | 2 | -32.734369 | 7.35384362 | 8.53E-06 | 8.93E-06 |
| rs57149748 | 1 | 162694835 | GENOTYPED | DDR2 | T | C | 0.0002377 | 2 | -32.734369 | 7.35384362 | 8.53E-06 | 8.93E-06 |
| rs143601614 | 1 | 162723518 | IMPUTED | DDR2 | C | G | 0.0002377 | 2 | -32.734369 | 7.35384362 | 8.53E-06 | 8.93E-06 |
| rs6699299 | 1 | 162735213 | GENOTYPED | DDR2 | C | T | 0.0002377 | 2 | -32.734369 | 7.35384362 | 8.53E-06 | 8.93E-06 |
| rs75108476 | 1 | 162813814 | IMPUTED | HSD17B7(dist=31206),CCDC190(dist=10273) | A | G | 0.0002377 | 2 | -32.734369 | 7.35384362 | 8.53E-06 | 8.93E-06 |
| rs75065453 | 1 | 162853093 | IMPUTED | CCDC190(dist=14488),RGS4(dist=185303) | C | T | 0.0002377 | 2 | -32.734369 | 7.35384362 | 8.53E-06 | 8.93E-06 |
| rs116697211 | 1 | 163471359 | GENOTYPED | LOC100422212(dist=78378),PBX1(dist=1057238) | G | A | 0.0002377 | 2 | -32.734369 | 7.35384362 | 8.53E-06 | 8.93E-06 |

| | | | | | | | | | | | | |
|---|---|---|---|---|---|---|---|---|---|---|---|---|
| rs73023495 | 1 | 163477913 | IMPUTED | LOC100422212(dist=84932),PBX1(dist=1050684) | G | A | 0.0002377 | 2 | -32.734369 | 7.35384362 | 8.53E-06 | 8.93E-06 |
| rs79836011 | 1 | 163477992 | GENOTYPED | LOC100422212(dist=85011),PBX1(dist=1050605) | G | A | 0.0002377 | 2 | -32.734369 | 7.35384362 | 8.53E-06 | 8.93E-06 |
| rs7538628 | 1 | 163478826 | GENOTYPED | LOC100422212(dist=85845),PBX1(dist=1049771) | A | G | 0.0002377 | 2 | -32.734369 | 7.35384362 | 8.53E-06 | 8.93E-06 |
| rs12565204 | 1 | 163479889 | GENOTYPED | LOC100422212(dist=86908),PBX1(dist=1048708) | T | C | 0.0002377 | 2 | -32.734369 | 7.35384362 | 8.53E-06 | 8.93E-06 |
| rs113488456 | 1 | 163481828 | IMPUTED | LOC100422212(dist=88847),PBX1(dist=1046769) | G | A | 0.0002377 | 2 | -32.734369 | 7.35384362 | 8.53E-06 | 8.93E-06 |
| rs145379856 | 1 | 163491096 | IMPUTED | LOC100422212(dist=98115),PBX1(dist=1037501) | G | A | 0.0002377 | 2 | -32.734369 | 7.35384362 | 8.53E-06 | 8.93E-06 |
| rs76142447 | 1 | 163528507 | GENOTYPED | LOC100422212(dist=135526),PBX1(dist=1000090) | A | G | 0.0002377 | 2 | -32.734369 | 7.35384362 | 8.53E-06 | 8.93E-06 |
| rs12086244 | 1 | 163542696 | IMPUTED | LOC100422212(dist=149715),PBX1(dist=985901) | G | A | 0.0002377 | 2 | -32.734369 | 7.35384362 | 8.53E-06 | 8.93E-06 |
| rs10917803 | 1 | 163545323 | IMPUTED | LOC100422212(dist=152342),PBX1(dist=983274) | C | T | 0.0002377 | 2 | -32.734369 | 7.35384362 | 8.53E-06 | 8.93E-06 |
| rs115967054 | 1 | 163846640 | GENOTYPED | LOC100422212(dist=453659),PBX1(dist=681957) | G | A | 0.0002377 | 2 | -32.734369 | 7.35384362 | 8.53E-06 | 8.93E-06 |
| rs1432428 | 1 | 187993896 | GENOTYPED | LINC01037(dist=547542),NONE(dist=NONE) | C | T | 0.0002377 | 2 | -32.734369 | 7.35384362 | 8.53E-06 | 8.93E-06 |
| rs78818485 | 1 | 162687331 | IMPUTED | DDR2 | C | G | 0.00023877 | 2 | -32.732332 | 7.35377085 | 8.54E-06 | 8.94E-06 |
| rs146989532 | 1 | 163840741 | IMPUTED | LOC100422212(dist=447760),PBX1(dist=687856) | G | A | 0.00023806 | 2 | -32.732012 | 7.3538498 | 8.55E-06 | 8.95E-06 |
| rs76606125 | 1 | 163841505 | IMPUTED | LOC100422212(dist=448524),PBX1(dist=687092) | C | T | 0.00023806 | 2 | -32.732012 | 7.3538498 | 8.55E-06 | 8.95E-06 |
| rs6659194 | 1 | 162685364 | IMPUTED | DDR2 | G | A | 0.00023889 | 2 | -32.72907 | 7.35375484 | 8.56E-06 | 8.96E-06 |
| rs140441781 | 8 | 82826977 | IMPUTED | SNX16(dist=72456),LOC101927141(dist=997362) | C | T | 0.00023806 | 2 | -32.726226 | 7.35382363 | 8.58E-06 | 8.98E-06 |
| rs192033741 | 8 | 82827375 | IMPUTED | SNX16(dist=72854),LOC101927141(dist=996964) | C | T | 0.00023806 | 2 | -32.726226 | 7.35382363 | 8.58E-06 | 8.98E-06 |
| rs115098909 | 1 | 162714006 | IMPUTED | DDR2 | G | A | 0.00024055 | 2 | -32.724767 | 7.35380221 | 8.59E-06 | 8.99E-06 |
| rs9506584 | 13 | 21525511 | IMPUTED | LINC00367(dist=1944),LATS2(dist=21665) | G | T | 0.99510887 | 41 | 7.48942407 | 1.68310399 | 8.60E-06 | 9.00E-06 |
| rs558451449 | 1 | 104900815 | IMPUTED | LOC100129138(dist=281122),LINC01676(dist=1231501) | T | C | 0.00024067 | 2 | -32.722598 | 7.3538345 | 8.60E-06 | 9.00E-06 |
| rs115086013 | 8 | 82774081 | IMPUTED | SNX16(dist=19560),LOC101927141(dist=1050258) | G | A | 0.00024495 | 2 | -32.720233 | 7.35361979 | 8.61E-06 | 9.01E-06 |
| rs142466649 | 1 | 162674578 | IMPUTED | DDR2 | C | T | 0.00023806 | 2 | -32.718864 | 7.35388454 | 8.62E-06 | 9.02E-06 |
| rs150243641 | 7 | 20118216 | IMPUTED | LOC101927668 | G | A | 0.00025089 | 2 | -32.852377 | 7.38546424 | 8.66E-06 | 9.06E-06 |
| rs150484952 | 1 | 163490907 | IMPUTED | LOC100422212(dist=97926),PBX1(dist=1037690) | A | G | 0.00023782 | 2 | -32.792364 | 7.37240218 | 8.67E-06 | 9.07E-06 |
| rs9316030 | 13 | 21525398 | IMPUTED | LINC00367(dist=1831),LATS2(dist=21778) | C | T | 0.99511303 | 41 | 7.49458548 | 1.68518055 | 8.69E-06 | 9.10E-06 |
| rs16855210 | 1 | 163479854 | IMPUTED | LOC100422212(dist=86873),PBX1(dist=1048743) | G | T | 0.0002371 | 2 | -32.78593 | 7.37242101 | 8.70E-06 | 9.11E-06 |
| rs16855237 | 1 | 163480472 | IMPUTED | LOC100422212(dist=87491),PBX1(dist=1048125) | C | A | 0.0002371 | 2 | -32.78593 | 7.37242101 | 8.70E-06 | 9.11E-06 |
| rs7526481 | 1 | 163484242 | IMPUTED | LOC100422212(dist=91261),PBX1(dist=1044355) | C | T | 0.0002371 | 2 | -32.78593 | 7.37242101 | 8.70E-06 | 9.11E-06 |
| rs146563539 | 1 | 163486488 | IMPUTED | LOC100422212(dist=93507),PBX1(dist=1042109) | C | G | 0.0002371 | 2 | -32.78593 | 7.37242101 | 8.70E-06 | 9.11E-06 |
| rs111461063 | 1 | 163490575 | IMPUTED | LOC100422212(dist=97594),PBX1(dist=1038022) | G | A | 0.0002371 | 2 | -32.78593 | 7.37242101 | 8.70E-06 | 9.11E-06 |
| rs553414377 | 12 | 40663237 | IMPUTED | LRRK2 | T | C | 0.00023734 | 2 | -33.531364 | 7.54025672 | 8.71E-06 | 9.11E-06 |
| rs115110312 | 10 | 90801365 | IMPUTED | FAS(dist=24547),MIR4679-2(dist=21727) | T | C | 0.00022142 | 2 | -33.919513 | 7.62811289 | 8.72E-06 | 9.13E-06 |
| rs73825624 | 3 | 24400340 | IMPUTED | THRB | T | C | 0.00381994 | 32 | 8.48251873 | 1.90835033 | 8.79E-06 | 9.20E-06 |
| rs746863731 | 6 | 54713225 | IMPUTED | FAM83B | C | A | 0.00051117 | 4 | 24.6241463 | 5.54290362 | 8.89E-06 | 9.31E-06 |
| rs117966882 | 15 | 82396031 | IMPUTED | LINC01583(dist=5997),EFL1(dist=26530) | G | A | 0.00023829 | 2 | -34.04605 | 7.66442652 | 8.91E-06 | 9.32E-06 |
| rs754401661 | 5 | 6297339 | IMPUTED | ICE1(dist=806992),LINC02145(dist=13215) | C | T | 0.00018148 | 2 | -41.095679 | 9.25376401 | 8.96E-06 | 9.37E-06 |
| rs17057178 | 18 | 73233917 | GENOTYPED | SMIM21(dist=94259),LINC01898(dist=174121) | C | T | 0.08236095 | 693 | -1.8718627 | 0.42155405 | 8.98E-06 | 9.39E-06 |
| rs74330829 | 10 | 114568863 | IMPUTED | VTI1A | G | C | 0.00012634 | 1 | 46.3979386 | 10.4502788 | 9.00E-06 | 9.42E-06 |
| rs142267675 | 8 | 99053275 | IMPUTED | RPL30(dist=663) | C | T | 0.00024958 | 2 | -32.656556 | 7.35644183 | 9.03E-06 | 9.45E-06 |
| rs530243109 | 3 | 169314534 | IMPUTED | MECOM | C | T | 0.00024329 | 2 | -32.644083 | 7.35380025 | 9.03E-06 | 9.45E-06 |
| rs531930435 | 6 | 83208846 | IMPUTED | TPBG(dist=131713),UBE3D(dist=393271) | G | C | 0.00046268 | 4 | 23.5921471 | 5.31590606 | 9.08E-06 | 9.50E-06 |
| rs544069805 | 6 | 16307271 | IMPUTED | ATXN1 | G | A | 0.00042108 | 4 | 29.3740418 | 6.61910326 | 9.09E-06 | 9.51E-06 |
| rs114241402 | 1 | 163493815 | IMPUTED | LOC100422212(dist=100834),PBX1(dist=1034782) | G | A | 0.00023592 | 2 | -32.914781 | 7.41709639 | 9.09E-06 | 9.51E-06 |
| rs572615835 | 2 | 54790377 | IMPUTED | SPTBN1 | C | G | 0.00040064 | 3 | 29.1028845 | 6.55835877 | 9.10E-06 | 9.52E-06 |
| rs773629474 | 3 | 169291717 | IMPUTED | MECOM | T | C | 0.00080509 | 7 | -19.627994 | 4.42343721 | 9.11E-06 | 9.53E-06 |
| rs78092409 | 10 | 114567138 | IMPUTED | VTI1A | A | C | 0.00012515 | 1 | 46.4257756 | 10.4627688 | 9.11E-06 | 9.53E-06 |
| rs67883680 | 18 | 73235450 | IMPUTED | SMIM21(dist=95792),LINC01898(dist=172588) | T | C | 0.0752667 | 633 | -1.9535908 | 0.44034532 | 9.14E-06 | 9.56E-06 |
| rs537514827 | 6 | 83312747 | IMPUTED | TPBG(dist=235614),UBE3D(dist=289370) | A | G | 0.00046268 | 4 | 23.5857767 | 5.31672247 | 9.16E-06 | 9.58E-06 |
| rs775525523 | 16 | 54258059 | IMPUTED | FTO(dist=109680),LINC02169(dist=21398) | T | A | 0.0002995 | 3 | -31.919993 | 7.19553446 | 9.16E-06 | 9.58E-06 |
| rs145330758 | 8 | 98907261 | IMPUTED | MATN2 | C | T | 0.00024673 | 2 | -32.605501 | 7.35103059 | 9.19E-06 | 9.61E-06 |
| rs567409192 | 2 | 52670336 | IMPUTED | LOC730100(dist=35281),MIR4431(dist=259324) | C | A | 0.00038103 | 3 | 29.183307 | 6.57975196 | 9.19E-06 | 9.62E-06 |
| rs528472306 | 6 | 83242587 | IMPUTED | TPBG(dist=165454),UBE3D(dist=359530) | A | T | 0.00045876 | 4 | 23.7870388 | 5.36322166 | 9.20E-06 | 9.62E-06 |

| | | | | | | | | | | | | |
|---|---|---|---|---|---|---|---|---|---|---|---|---|
| rs73328073 | 5 | 171274975 | IMPUTED | SMIM23(dist=56883),FBXW11(dist=13581) | C | T | 0.00044985 | 4 | -25.696611 | 5.79428507 | 9.21E-06 | 9.64E-06 |
| rs554265290 | 1 | 41219161 | IMPUTED | NFYC | G | A | 0.00025256 | 2 | -34.040299 | 7.67585142 | 9.22E-06 | 9.64E-06 |
| rs947620457 | 12 | 127133829 | IMPUTED | LINC02347(dist=176498),LOC100996671(dist=3663) | G | A | 0.00111909 | 9 | 15.7242332 | 3.54609307 | 9.24E-06 | 9.66E-06 |
| rs553929719 | 2 | 54661230 | IMPUTED | C2orf73(dist=72516),SPTBN1(dist=22224) | G | A | 0.00039838 | 3 | 29.1041103 | 6.56375633 | 9.25E-06 | 9.67E-06 |
| rs115689294 | 17 | 30047103 | IMPUTED | MIR365B(dist=144563),COPRS(dist=131781) | A | C | 0.00041229 | 3 | 28.9442349 | 6.52798565 | 9.26E-06 | 9.68E-06 |
| rs557232295 | 18 | 66914932 | IMPUTED | CCDC102B(dist=192506),DOK6(dist=153352) | G | T | 0.00177454 | 15 | -13.59696 | 3.06705342 | 9.28E-06 | 9.71E-06 |
| rs138992017 | 15 | 65075140 | IMPUTED | RBPMS2(dist=7354),PIF1(dist=32689) | T | G | 0.00903173 | 76 | -5.790683 | 1.30623482 | 9.29E-06 | 9.71E-06 |
| rs560366370 | 2 | 52669698 | IMPUTED | LOC730100(dist=34643),MIR4431(dist=259962) | T | C | 0.00038151 | 3 | 29.1663502 | 6.57971418 | 9.30E-06 | 9.73E-06 |
| rs118150476 | 10 | 92050053 | IMPUTED | LINC01375(dist=332923),LOC101926942(dist=112225) | G | C | 0.00649548 | 55 | 7.07415768 | 1.59589886 | 9.31E-06 | 9.73E-06 |
| rs79343744 | 2 | 151551293 | IMPUTED | LOC101929282(dist=59422),RBM43(dist=553435) | T | C | 0.00042227 | 4 | -26.217223 | 5.91520114 | 9.33E-06 | 9.76E-06 |
| rs772998989 | 3 | 156502603 | IMPUTED | LINC00886 | G | A | 0.00037568 | 3 | 28.3823993 | 6.40416027 | 9.34E-06 | 9.77E-06 |
| rs7520505 | 1 | 162669855 | IMPUTED | DDR2 | T | C | 0.00024222 | 2 | -32.580055 | 7.35258778 | 9.38E-06 | 9.80E-06 |
| rs78145580 | 10 | 114560843 | IMPUTED | VTI1A | A | G | 0.00012348 | 1 | 46.2956457 | 10.4482661 | 9.38E-06 | 9.81E-06 |
| rs116880183 | 18 | 57503119 | IMPUTED | CCBE1(dist=138475),PMAIP1(dist=64073) | T | C | 0.01592073 | 134 | -4.5119371 | 1.01893875 | 9.51E-06 | 9.94E-06 |
| rs117528598 | 14 | 21202538 | IMPUTED | RNASE4(dist=33777),EDDM3A(dist=11561) | A | T | 0.00074257 | 6 | 19.2787867 | 4.35376642 | 9.51E-06 | 9.94E-06 |
| rs184467205 | 8 | 76988559 | IMPUTED | HNF4G(dist=509482),LINC01111(dist=330330) | G | T | 0.0015025 | 13 | -14.362121 | 3.24344572 | 9.51E-06 | 9.94E-06 |
| rs72885535 | 2 | 52497133 | IMPUTED | LOC730100 | A | G | 0.00036772 | 3 | 30.7523699 | 6.94544485 | 9.52E-06 | 9.96E-06 |
| rs561792757 | 12 | 97348275 | IMPUTED | NEDD1(dist=806) | G | C | 0.0003972 | 3 | 29.2598733 | 6.60945476 | 9.56E-06 | 9.99E-06 |
| rs62460339 | 7 | 68614093 | IMPUTED | LOC102723427(dist=1116416),LOC100507468(dist=447031 | G | A | 0.09810364 | 825 | -1.8006291 | 0.40679546 | 9.58E-06 | 1.00E-05 |
| rs10399864 | 1 | 163473431 | IMPUTED | LOC100422212(dist=80450),PBX1(dist=1055166) | A | T | 0.00024031 | 2 | -32.623189 | 7.37189984 | 9.63E-06 | 1.01E-05 |
| rs892691836 | 6 | 69678427 | IMPUTED | ADGRB3 | A | C | 0.00028797 | 2 | 30.1104144 | 6.8050576 | 9.66E-06 | 1.01E-05 |
| rs573486780 | 1 | 172964004 | IMPUTED | FASLG(dist=327992),TNFSF18(dist=46356) | A | C | 0.00035714 | 3 | 26.4979676 | 5.98897093 | 9.67E-06 | 1.01E-05 |
| rs542193620 | 12 | 92394209 | IMPUTED | LINC01619 | T | C | 0.00049156 | 4 | -22.700544 | 5.13080809 | 9.67E-06 | 1.01E-05 |
| rs745981658 | 1 | 212213715 | IMPUTED | DTL | G | T | 0.00013525 | 1 | 44.696166 | 10.1076838 | 9.78E-06 | 1.02E-05 |
| rs765061610 | 1 | 90492457 | IMPUTED | ZNF326 | T | C | 0.00033254 | 3 | -28.505844 | 6.44708834 | 9.80E-06 | 1.02E-05 |
| rs902592460 | 12 | 127093914 | IMPUTED | LINC02347(dist=136583),LOC100996671(dist=43578) | G | A | 0.00123746 | 10 | 15.2795149 | 3.45682541 | 9.87E-06 | 1.03E-05 |
| rs75684812 | 1 | 163525360 | IMPUTED | LOC100422212(dist=132379),PBX1(dist=1003237) | G | A | 0.00021429 | 2 | -36.67865 | 8.29975664 | 9.90E-06 | 1.04E-05 |
| rs762567682 | 19 | 22540017 | IMPUTED | ZNF729(dist=40039),ZNF98(dist=33882) | G | A | 0.00049334 | 4 | -23.835378 | 5.39454264 | 9.94E-06 | 1.04E-05 |
| rs59090179 | 3 | 24063857 | IMPUTED | NR1D2(dist=41748),LINC00691(dist=77608) | A | G | 0.00059449 | 5 | 20.2950509 | 4.59329105 | 9.94E-06 | 1.04E-05 |
| rs143716771 | 16 | 60949631 | IMPUTED | LOC729159(dist=555934),MIR4426(dist=139980) | A | G | 0.00341467 | 29 | -8.5653442 | 1.93861719 | 9.95E-06 | 1.04E-05 |
| rs112165405 | 1 | 164617327 | IMPUTED | PBX1 | C | T | 0.00026789 | 2 | -32.366972 | 7.32663507 | 9.98E-06 | 1.04E-05 |
| rs777334205 | 2 | 33844386 | IMPUTED | FAM98A(dist=19957),LINC01317(dist=87567) | A | G | 0.0002289 | 2 | 34.8294972 | 7.88481552 | 9.99E-06 | 1.04E-05 |
