## Supplemental_table2 for "Rare Genetic Variants Correlate with Better Processing Speed"

**Supplement Table2. SNPs with Wald p-values < $1\times10^{-6}$ from the longitudianl GWAS in LLFS.**

Chr, Chromosome; Position, position of SNP on GRCh37 reference panel; Ref and Alt, reference allele and alternative allele; EAF, effect allele frequency; Wald.p, p-values from Wald test; Score.p, p-values from Score test

| SNP | Chr | Position | Gene | Ref | Alt | EAF | Effect | SE | Wald.p | Score.p |
|---|---|---|---|---|---|---|---|---|---|---|
| rs58169119 | 8 | 82350918 | FABP5(dist=153906),PMP2(dist=1645) | G | A | 0.00047076 | -27.882322 | 4.85932218 | 9.59E-09 | 1.71E-08 |
| rs6754826 | 2 | 174325120 | CDCA7(dist=91402),SP3(dist=446067) | T | C | 0.99945436 | -26.285385 | 4.72180036 | 2.59E-08 | 1.46E-07 |
| rs140387800 | 3 | 26853308 | LRRC3B(dist=101043),NEK10(dist=299086) | G | C | 0.00027419 | 40.4182023 | 7.27669515 | 2.78E-08 | 1.14E-07 |
| rs9821587 | 3 | 24712189 | MIR4792(dist=149263),RARB(dist=158625) | G | T | 0.00056204 | 26.5487019 | 4.81440899 | 3.50E-08 | 3.98E-08 |
| rs7623455 | 3 | 24713169 | MIR4792(dist=150243),RARB(dist=157645) | G | A | 0.0005624 | 26.5430775 | 4.81341333 | 3.50E-08 | 3.98E-08 |
| rs9821776 | 3 | 24712332 | MIR4792(dist=149406),RARB(dist=158482) | G | C | 0.0005624 | 26.5408788 | 4.81340824 | 3.51E-08 | 3.98E-08 |
| rs116193953 | 1 | 163629475 | LOC100422212(dist=236494),PBX1(dist=899122) | A | C | 0.00055764 | -28.087092 | 5.10813629 | 3.83E-08 | 7.80E-08 |
| rs78188599 | 8 | 9141514 | LOC101929128(dist=81148),LOC157273(dist=41047) | G | A | 0.02427098 | 3.88226317 | 0.71314821 | 5.21E-08 | 3.98E-08 |
| rs148022846 | 3 | 25100109 | RARB | A | T | 0.00048205 | 28.1894779 | 5.2211336 | 6.70E-08 | 6.01E-08 |
| rs146691363 | 2 | 160767097 | LY75-CD302(dist=5830),PLA2R1(dist=30163) | G | C | 0.00072463 | -24.868492 | 4.60687146 | 6.73E-08 | 4.04E-08 |
| rs183090477 | 2 | 36794491 | FEZ2 | T | G | 0.00073247 | 22.4290637 | 4.15696037 | 6.83E-08 | 3.97E-08 |
| rs75963215 | 3 | 24423893 | THRB | A | C | 0.00095139 | 19.3024106 | 3.58146373 | 7.06E-08 | 4.09E-08 |
| rs59914825 | 3 | 24424415 | THRB | T | C | 0.00095127 | 19.3023012 | 3.58147024 | 7.07E-08 | 4.09E-08 |
| rs4266131 | 3 | 24425760 | THRB | A | G | 0.99902817 | -19.250597 | 3.57343155 | 7.16E-08 | 4.09E-08 |
| rs78704059 | 3 | 25170804 | RARB | A | G | 0.00047611 | 27.9688845 | 5.21957624 | 8.39E-08 | 6.01E-08 |
| rs189337466 | 3 | 25228798 | RARB | T | A | 0.00047504 | 27.9975045 | 5.22499929 | 8.40E-08 | 6.01E-08 |
| rs181742183 | 15 | 85216399 | SEC11A | G | A | 0.00039315 | -29.145748 | 5.44034448 | 8.45E-08 | 1.68E-07 |
| rs77306558 | 3 | 25223736 | RARB | A | T | 0.00047659 | 27.9611791 | 5.2209107 | 8.53E-08 | 6.01E-08 |
| rs59296535 | 3 | 25206678 | RARB | C | T | 0.00047659 | 27.9525645 | 5.21948055 | 8.54E-08 | 6.01E-08 |
| rs74467766 | 3 | 25206522 | RARB | A | C | 0.00047682 | 27.9509397 | 5.21947547 | 8.55E-08 | 6.01E-08 |
| rs79046847 | 3 | 25217506 | RARB | A | G | 0.00047682 | 27.9492166 | 5.21949638 | 8.57E-08 | 6.01E-08 |
| rs80164536 | 3 | 25212666 | RARB | C | G | 0.00047694 | 27.9474743 | 5.21948383 | 8.58E-08 | 6.01E-08 |
| rs77025115 | 3 | 25192336 | RARB | T | G | 0.00047778 | 27.9461239 | 5.21945115 | 8.59E-08 | 6.01E-08 |
| rs80011850 | 3 | 25194665 | RARB | A | G | 0.00047742 | 27.943132 | 5.21946544 | 8.62E-08 | 6.01E-08 |
| rs80223246 | 3 | 25206289 | RARB | C | T | 0.0004773 | 27.9406643 | 5.21948668 | 8.64E-08 | 6.01E-08 |
| rs554910598 | 20 | 25474435 | NINL | G | A | 0.00034894 | -30.082026 | 5.63262095 | 9.26E-08 | 1.52E-07 |
| rs719408 | 8 | 9151138 | LOC101929128(dist=90772),LOC157273(dist=31423) | G | T | 0.02462111 | 3.77882494 | 0.70798057 | 9.43E-08 | 8.83E-08 |
| rs77189115 | 8 | 9143416 | LOC101929128(dist=83050),LOC157273(dist=39145) | C | T | 0.02435346 | 3.80374009 | 0.7132034 | 9.64E-08 | 1.14E-07 |
| rs111975479 | 8 | 9148463 | LOC101929128(dist=88097),LOC157273(dist=34098) | G | C | 0.02459698 | 3.7728038 | 0.70767245 | 9.75E-08 | 8.83E-08 |
| rs111795201 | 8 | 9147367 | LOC101929128(dist=87001),LOC157273(dist=35194) | G | A | 0.02460007 | 3.7714529 | 0.70760866 | 9.83E-08 | 8.83E-08 |
| rs75170749 | 8 | 9149731 | LOC101929128(dist=89365),LOC157273(dist=32830) | C | T | 0.02460304 | 3.77022869 | 0.70753156 | 9.89E-08 | 8.83E-08 |
| rs76745642 | 8 | 9151712 | LOC101929128(dist=91346),LOC157273(dist=30849) | C | T | 0.02458272 | 3.77661669 | 0.70974904 | 1.03E-07 | 8.83E-08 |
| rs79907700 | 2 | 151544901 | LOC101929282(dist=53030),RBM43(dist=559827) | C | T | 0.00046541 | -27.77933 | 5.22966898 | 1.09E-07 | 1.89E-07 |
| rs77977386 | 2 | 151550092 | LOC101929282(dist=58221),RBM43(dist=554636) | G | A | 0.00046637 | -27.740419 | 5.22278375 | 1.09E-07 | 1.89E-07 |
| rs74397915 | 2 | 151554303 | LOC101929282(dist=62432),RBM43(dist=550425) | A | G | 0.00046637 | -27.740419 | 5.22278375 | 1.09E-07 | 1.89E-07 |
| rs74761575 | 2 | 151541269 | LOC101929282(dist=49398),RBM43(dist=563459) | A | T | 0.00046589 | -27.766644 | 5.23019719 | 1.10E-07 | 1.89E-07 |
| rs113378776 | 2 | 151540798 | LOC101929282(dist=48927),RBM43(dist=563930) | T | C | 0.00046007 | -28.098585 | 5.29622666 | 1.12E-07 | 1.89E-07 |
| rs112905701 | 2 | 151546989 | LOC101929282(dist=55118),RBM43(dist=557739) | C | T | 0.00044046 | -29.031452 | 5.50963367 | 1.37E-07 | 1.89E-07 |
| rs75464434 | 2 | 151548853 | LOC101929282(dist=56982),RBM43(dist=555875) | C | A | 0.00043986 | -29.055398 | 5.51603711 | 1.38E-07 | 1.89E-07 |
| rs75060076 | 3 | 25235889 | RARB | C | T | 0.00045698 | 28.5849445 | 5.42816151 | 1.39E-07 | 6.01E-08 |
| rs114219304 | 2 | 151583525 | LOC101929282(dist=91654),RBM43(dist=521203) | G | A | 0.00043725 | -29.101785 | 5.53926502 | 1.49E-07 | 1.89E-07 |
| rs77473871 | 3 | 25279859 | RARB | T | C | 0.00049465 | 27.3747542 | 5.21584911 | 1.53E-07 | 6.01E-08 |
| rs75702282 | 8 | 9150378 | LOC101929128(dist=90012),LOC157273(dist=32183) | T | C | 0.02483955 | 3.68528259 | 0.70365654 | 1.63E-07 | 1.32E-07 |
| rs77444611 | 8 | 9150131 | LOC101929128(dist=89765),LOC157273(dist=32430) | C | T | 0.0248392 | 3.685337 | 0.70366889 | 1.63E-07 | 1.32E-07 |
| rs141237895 | 1 | 215308848 | KCNK2 | A | C | 0.00019277 | -42.932068 | 8.21069644 | 1.71E-07 | 1.25E-06 |
| rs74596618 | 2 | 151544939 | LOC101929282(dist=53068),RBM43(dist=559789) | C | T | 0.00043404 | -29.280624 | 5.60994634 | 1.79E-07 | 1.89E-07 |
| rs76789477 | 3 | 25378703 | RARB | C | G | 0.00050297 | 26.976478 | 5.1842628 | 1.96E-07 | 6.01E-08 |
| rs4972570 | 2 | 174325744 | CDCA7(dist=92026),SP3(dist=445443) | A | T | 0.99952341 | -25.121463 | 4.83794948 | 2.07E-07 | 1.46E-07 |
| rs4972402 | 2 | 174327221 | CDCA7(dist=93503),SP3(dist=443966) | A | G | 0.99950844 | -25.089382 | 4.83809214 | 2.15E-07 | 1.46E-07 |

| | | | | | | | | | | |
|---|---|---|---|---|---|---|---|---|---|---|
| rs1405229 | 2 | 174324895 | CDCA7(dist=91177),SP3(dist=446292) | A | G | 0.99951854 | -25.078153 | 4.83621997 | 2.15E-07 | 1.46E-07 |
| rs75407235 | 1 | 188102370 | LINC01037(dist=656016),BRINP3(dist=1964427) | A | T | 0.00022058 | -39.557629 | 7.63155194 | 2.18E-07 | 1.25E-06 |
| rs113543017 | 2 | 151564112 | LOC101929282(dist=72241),RBM43(dist=540616) | G | A | 0.00045496 | -28.453328 | 5.51711552 | 2.51E-07 | 1.89E-07 |
| rs142018669 | 12 | 53411481 | EIF4B | G | A | 0.00023283 | -38.280117 | 7.43227765 | 2.60E-07 | 1.25E-06 |
| rs577268629 | 5 | 150322147 | ZNF300P1 | A | T | 0.00053625 | 25.9592908 | 5.05932703 | 2.88E-07 | 3.85E-07 |
| rs749433443 | 6 | 170870119 | TBP | G | A | 0.00026004 | -36.094884 | 7.03914936 | 2.93E-07 | 1.25E-06 |
| rs111661085 | 9 | 88730110 | GOLM1(dist=14994),LOC101927623(dist=12345) | G | A | 0.00098835 | 20.1741464 | 3.93650893 | 2.98E-07 | 2.04E-07 |
| rs74464242 | 1 | 187976734 | LINC01037(dist=530380),NONE(dist=NONE) | C | T | 0.00021393 | -38.867231 | 7.59237893 | 3.07E-07 | 1.25E-06 |
| rs138545977 | 1 | 187979976 | LINC01037(dist=533622),NONE(dist=NONE) | C | T | 0.0002144 | -38.860562 | 7.59240603 | 3.08E-07 | 1.25E-06 |
| rs112676561 | 9 | 88752947 | LOC101927623 | G | A | 0.00098336 | 20.1421493 | 3.93593123 | 3.10E-07 | 2.04E-07 |
| rs190537399 | 12 | 78933270 | LINC02424(dist=179744),SYT1(dist=324503) | T | A | 0.00022807 | -39.389688 | 7.69845965 | 3.11E-07 | 1.25E-06 |
| rs782769776 | 1 | 147130330 | ACP6,NBPF19 | A | G | 0.00048657 | -27.180253 | 5.31414836 | 3.14E-07 | 4.07E-07 |
| rs778464311 | 6 | 95898881 | TSG1(dist=1412682),MANEA-AS1(dist=109091) | C | A | 0.00076159 | 20.4860283 | 4.00743214 | 3.19E-07 | 9.75E-08 |
| rs76432074 | 2 | 151565341 | LOC101929282(dist=73470),RBM43(dist=539387) | A | G | 0.00044925 | -28.644709 | 5.61321054 | 3.34E-07 | 1.89E-07 |
| rs74649461 | 2 | 151564122 | LOC101929282(dist=72251),RBM43(dist=540606) | G | C | 0.00044937 | -28.635599 | 5.61316738 | 3.37E-07 | 1.89E-07 |
| rs60248650 | 1 | 162885760 | CCDC190(dist=47155),RGS4(dist=152636) | G | A | 0.00033409 | -29.175136 | 5.72360534 | 3.44E-07 | 7.42E-07 |
| rs531471021 | 18 | 29650959 | RNF125(NM_017831:c.*26120>0) | G | A | 0.00048158 | -27.264443 | 5.36548314 | 3.75E-07 | 4.41E-07 |
| rs101774090 | 6 | 92831492 | CASC6(dist=431346),EPHA7(dist=1118248) | A | G | 0.00071892 | 20.6714868 | 4.07477352 | 3.92E-07 | 3.59E-07 |
| rs113920675 | 8 | 82440603 | FABP12 | C | T | 0.00030307 | -32.478891 | 6.40935441 | 4.03E-07 | 4.43E-06 |
| rs144759232 | 12 | 40714394 | LRRK2 | G | A | 0.00026254 | -35.621787 | 7.0432116 | 4.25E-07 | 1.25E-06 |
| rs102498301 | 6 | 92781358 | CASC6(dist=381212),EPHA7(dist=1168382) | A | G | 0.00070692 | 20.9369734 | 4.14470879 | 4.38E-07 | 3.59E-07 |
| rs61764931 | 1 | 4986475 | AJAP1(dist=142624),MIR4417(dist=637656) | A | G | 0.01469907 | -4.8717448 | 0.96547285 | 4.51E-07 | 2.89E-06 |
| rs780529937 | 8 | 80843646 | MRPS28 | T | C | 0.00027704 | -35.704375 | 7.08225385 | 4.62E-07 | 1.25E-06 |
| rs188219563 | 8 | 99163374 | POP1 | C | T | 0.00022879 | -36.661408 | 7.28024334 | 4.76E-07 | 1.25E-06 |
| rs566408803 | 1 | 80702723 | ADGRL4(dist=1230228),LINC01781(dist=298717) | G | T | 0.00024305 | -37.074196 | 7.37028298 | 4.90E-07 | 1.25E-06 |
| rs573486780 | 1 | 172964004 | FASLG(dist=327992),TNFSF18(dist=46356) | A | C | 0.00035714 | 28.1941718 | 5.60745613 | 4.96E-07 | 8.06E-07 |
| rs902845237 | 6 | 93232345 | CASC6(dist=832199),EPHA7(dist=717395) | G | T | 0.0007251 | 20.0127892 | 3.98501896 | 5.11E-07 | 3.59E-07 |
| rs553961701 | 6 | 93833544 | CASC6(dist=1433398),EPHA7(dist=116196) | C | T | 0.00060233 | 22.0981021 | 4.40132468 | 5.15E-07 | 3.50E-07 |
| rs104791120 | 6 | 94594504 | TSG1(dist=108305),MANEA-AS1(dist=1413468) | C | A | 0.00059246 | 22.1589295 | 4.41514678 | 5.20E-07 | 3.50E-07 |
| rs182347375 | 5 | 16439388 | LINC02150 | G | A | 0.00026812 | -35.677117 | 7.11165694 | 5.26E-07 | 1.25E-06 |
| rs931770839 | 6 | 92645910 | CASC6(dist=245764),EPHA7(dist=1303830) | A | G | 0.0007421 | 19.9810987 | 3.98447593 | 5.31E-07 | 3.59E-07 |
| rs115494049 | 8 | 82649057 | CHMP4C | G | C | 0.00023092 | -35.729232 | 7.13203839 | 5.45E-07 | 1.25E-06 |
| rs73277212 | 8 | 82451471 | FABP12(dist=7846),IMPA1P1(dist=64648) | T | A | 0.0002875 | -33.451016 | 6.68157599 | 5.54E-07 | 1.25E-06 |
| rs758723783 | 9 | 35953589 | SPAAR(dist=41972),OR2S2(dist=3516) | T | C | 0.00023758 | -36.289703 | 7.24867357 | 5.55E-07 | 1.25E-06 |
| rs144694633 | 8 | 82569926 | IMPA1(NM_001144878:c.*16600>0,NM_005536:c.*16600 | G | A | 0.00023164 | -35.595379 | 7.11386162 | 5.63E-07 | 1.25E-06 |
| rs775026608 | 1 | 104104807 | AMY2B | T | C | 0.00028405 | -34.728545 | 6.94709202 | 5.76E-07 | 1.25E-06 |
| rs141930236 | 8 | 82239070 | FABP5(dist=42058),PMP2(dist=113493) | G | A | 0.00023342 | -35.755719 | 7.15890855 | 5.90E-07 | 1.25E-06 |
| rs189653044 | 8 | 82321312 | FABP5(dist=124300),PMP2(dist=31251) | T | C | 0.00023734 | -35.141306 | 7.04707514 | 6.14E-07 | 1.25E-06 |
| rs105073917 | 2 | 156651436 | KCNJ3(dist=936572),LINC01876(dist=225611) | A | G | 0.00165023 | 15.3340855 | 3.07613399 | 6.20E-07 | 3.94E-06 |
| rs571970220 | 5 | 66483273 | CD180 | C | T | 0.00024531 | -34.679633 | 6.95755647 | 6.21E-07 | 1.25E-06 |
| rs141153583 | 8 | 82440058 | FABP12 | G | T | 0.00023568 | -35.041503 | 7.03190572 | 6.25E-07 | 1.25E-06 |
| rs10912126 | 1 | 187636974 | LINC01037(dist=190620),NONE(dist=NONE) | G | A | 0.00026254 | -34.322035 | 6.88777209 | 6.26E-07 | 1.25E-06 |
| rs146689853 | 1 | 187632912 | LINC01037(dist=186558),NONE(dist=NONE) | T | C | 0.00026242 | -34.320537 | 6.88777246 | 6.27E-07 | 1.25E-06 |
| rs35899516 | 13 | 107450854 | LINC00443(dist=126326),FAM155A(dist=370025) | T | G | 0.00054528 | 23.961982 | 4.80962693 | 6.29E-07 | 1.58E-06 |
| rs190566429 | 8 | 82330221 | FABP5(dist=133209),PMP2(dist=22342) | T | C | 0.00023437 | -35.082364 | 7.04421602 | 6.35E-07 | 1.25E-06 |
| rs146838899 | 15 | 83749289 | MIR4515(dist=13122),TM6SF1(dist=26949) | C | T | 0.00023104 | -35.720976 | 7.17380403 | 6.38E-07 | 1.25E-06 |
| rs112488623 | 8 | 82440157 | FABP12 | G | A | 0.00033088 | -30.180154 | 6.06169042 | 6.40E-07 | 4.43E-06 |
| rs188948126 | 20 | 25316590 | ABHD12 | T | C | 0.00049334 | -23.534679 | 4.72747804 | 6.42E-07 | 1.29E-06 |
| rs550387016 | 6 | 72846297 | RIMS1 | C | T | 0.00024768 | -34.663307 | 6.96762774 | 6.53E-07 | 1.25E-06 |
| rs150903802 | 8 | 82879135 | SNX16(dist=124614),LOC101927141(dist=945204) | G | A | 0.00024222 | -34.599423 | 6.95890111 | 6.63E-07 | 1.25E-06 |
| rs190253462 | 8 | 82874581 | SNX16(dist=120060),LOC101927141(dist=949758) | G | A | 0.00024317 | -34.598869 | 6.95884164 | 6.63E-07 | 1.25E-06 |
| rs181020921 | 20 | 25373429 | ABHD12(dist=1811),GINS1(dist=14890) | A | G | 0.00049394 | -23.446848 | 4.71882251 | 6.74E-07 | 1.29E-06 |

| | | | | | | | | | | |
|---|---|---|---|---|---|---|---|---|---|---|
| rs73275496 | 8 | 82443303 | FABP12 | G | T | 0.0002396 | -34.564166 | 6.958828 | 6.80E-07 | 1.25E-06 |
| rs542666267 | 9 | 37663948 | FRMPD1 | C | T | 0.00034169 | -31.960674 | 6.43508612 | 6.81E-07 | 5.90E-06 |
| rs563992891 | 6 | 40903799 | LOC101929555 | C | T | 0.00099643 | -18.11546 | 3.65136766 | 7.00E-07 | 4.81E-07 |
| rs115320746 | 1 | 162817025 | HSD17B7(dist=34417),CCDC190(dist=7062) | G | A | 0.00024911 | -34.48523 | 6.95238596 | 7.04E-07 | 1.25E-06 |
| rs762567682 | 19 | 22540017 | ZNF729(dist=40039),ZNF98(dist=33882) | G | A | 0.00049334 | -24.774469 | 4.99616603 | 7.10E-07 | 1.58E-06 |
| rs373619954 | 1 | 105045980 | LOC100129138(dist=426287),LINC01676(dist=1086336) | A | G | 0.00024008 | -34.508041 | 6.95949031 | 7.11E-07 | 1.25E-06 |
| rs56991769 | 12 | 83104383 | TMTC2 | G | T | 0.00014547 | -43.47708 | 8.76930899 | 7.13E-07 | 4.05E-06 |
| rs143839811 | 8 | 82780246 | SNX16(dist=25725),LOC101927141(dist=1044093) | T | C | 0.00023734 | -34.525639 | 6.96863779 | 7.25E-07 | 1.25E-06 |
| rs553414377 | 12 | 40663237 | LRRK2 | T | C | 0.00023734 | -35.333114 | 7.13227831 | 7.27E-07 | 1.25E-06 |
| rs145351234 | 8 | 98900292 | MATN2 | G | A | 0.00023889 | -34.473787 | 6.95950592 | 7.29E-07 | 1.25E-06 |
| rs532124872 | 6 | 72125965 | LINC00472 | T | A | 0.00023841 | -34.468853 | 6.9595528 | 7.32E-07 | 1.25E-06 |
| rs189119988 | 8 | 99056751 | RPL30 | T | G | 0.00023782 | -34.483598 | 6.96259133 | 7.32E-07 | 1.25E-06 |
| rs79990010 | 1 | 163489142 | LOC100422212(dist=96161),PBX1(dist=1039455) | G | T | 0.00024126 | -34.626786 | 6.9917704 | 7.33E-07 | 1.25E-06 |
| rs114632222 | 8 | 82745388 | SNX16 | A | G | 0.00023758 | -34.48117 | 6.96259959 | 7.33E-07 | 1.25E-06 |
| rs141094782 | 8 | 99018996 | MATN2 | T | G | 0.00023758 | -34.48117 | 6.96259959 | 7.33E-07 | 1.25E-06 |
| rs193187075 | 8 | 99021713 | MATN2 | A | G | 0.0002377 | -34.47942 | 6.96260561 | 7.34E-07 | 1.25E-06 |
| rs960923972 | 15 | 89964468 | MIR9-3HG(dist=22750),RHCG(dist=50172) | G | T | 0.00023984 | -37.061083 | 7.4839667 | 7.34E-07 | 1.25E-06 |
| rs73277205 | 8 | 82447770 | FABP12(dist=4145),IMPA1P1(dist=68349) | C | T | 0.00023782 | -34.463532 | 6.95958888 | 7.35E-07 | 1.25E-06 |
| rs150556028 | 8 | 82705793 | CHMP4C(dist=34045),SNX16(dist=6025) | G | T | 0.00023782 | -34.463532 | 6.95958888 | 7.35E-07 | 1.25E-06 |
| rs115100734 | 1 | 162794207 | HSD17B7(dist=11599),CCDC190(dist=29880) | A | G | 0.00023782 | -34.461692 | 6.95956882 | 7.36E-07 | 1.25E-06 |
| rs115086013 | 8 | 82774081 | SNX16(dist=19560),LOC101927141(dist=1050258) | G | A | 0.00024495 | -34.457762 | 6.9593402 | 7.37E-07 | 1.25E-06 |
| rs57149748 | 1 | 162694835 | DDR2 | T | C | 0.0002377 | -34.458941 | 6.95958111 | 7.37E-07 | 1.25E-06 |
| rs143601614 | 1 | 162723518 | DDR2 | C | G | 0.0002377 | -34.458941 | 6.95958111 | 7.37E-07 | 1.25E-06 |
| rs6699299 | 1 | 162735213 | DDR2 | C | T | 0.0002377 | -34.458941 | 6.95958111 | 7.37E-07 | 1.25E-06 |
| rs75108476 | 1 | 162813814 | HSD17B7(dist=31206),CCDC190(dist=10273) | A | G | 0.0002377 | -34.458941 | 6.95958111 | 7.37E-07 | 1.25E-06 |
| rs75065453 | 1 | 162853093 | CCDC190(dist=14488),RGS4(dist=185303) | C | T | 0.0002377 | -34.458941 | 6.95958111 | 7.37E-07 | 1.25E-06 |
| rs116697211 | 1 | 163471359 | LOC100422212(dist=78378),PBX1(dist=1057238) | G | A | 0.0002377 | -34.458941 | 6.95958111 | 7.37E-07 | 1.25E-06 |
| rs73023495 | 1 | 163477913 | LOC100422212(dist=84932),PBX1(dist=1050684) | G | A | 0.0002377 | -34.458941 | 6.95958111 | 7.37E-07 | 1.25E-06 |
| rs79836011 | 1 | 163477992 | LOC100422212(dist=85011),PBX1(dist=1050605) | G | A | 0.0002377 | -34.458941 | 6.95958111 | 7.37E-07 | 1.25E-06 |
| rs7538628 | 1 | 163478826 | LOC100422212(dist=85845),PBX1(dist=1049771) | A | G | 0.0002377 | -34.458941 | 6.95958111 | 7.37E-07 | 1.25E-06 |
| rs12565204 | 1 | 163479889 | LOC100422212(dist=86908),PBX1(dist=1048708) | T | C | 0.0002377 | -34.458941 | 6.95958111 | 7.37E-07 | 1.25E-06 |
| rs113488456 | 1 | 163481828 | LOC100422212(dist=88847),PBX1(dist=1046769) | G | A | 0.0002377 | -34.458941 | 6.95958111 | 7.37E-07 | 1.25E-06 |
| rs145379856 | 1 | 163491096 | LOC100422212(dist=98115),PBX1(dist=1037501) | G | A | 0.0002377 | -34.458941 | 6.95958111 | 7.37E-07 | 1.25E-06 |
| rs76142447 | 1 | 163528507 | LOC100422212(dist=135526),PBX1(dist=1000090) | A | G | 0.0002377 | -34.458941 | 6.95958111 | 7.37E-07 | 1.25E-06 |
| rs12086244 | 1 | 163542696 | LOC100422212(dist=149715),PBX1(dist=985901) | G | A | 0.0002377 | -34.458941 | 6.95958111 | 7.37E-07 | 1.25E-06 |
| rs10917803 | 1 | 163545323 | LOC100422212(dist=152342),PBX1(dist=983274) | C | T | 0.0002377 | -34.458941 | 6.95958111 | 7.37E-07 | 1.25E-06 |
| rs115967054 | 1 | 163846640 | LOC100422212(dist=453659),PBX1(dist=681957) | G | A | 0.0002377 | -34.458941 | 6.95958111 | 7.37E-07 | 1.25E-06 |
| rs73277204 | 8 | 82447200 | FABP12(dist=3575),IMPA1P1(dist=68919) | G | A | 0.0002377 | -34.458941 | 6.95958111 | 7.37E-07 | 1.25E-06 |
| rs1432428 | 1 | 187993896 | LINC01037(dist=547542),NONE(dist=NONE) | C | T | 0.0002377 | -34.458941 | 6.95958111 | 7.37E-07 | 1.25E-06 |
| rs188764513 | 8 | 82720924 | SNX16 | G | A | 0.0002377 | -34.458941 | 6.95958111 | 7.37E-07 | 1.25E-06 |
| rs115571718 | 8 | 82783619 | SNX16(dist=29098),LOC101927141(dist=1040720) | T | C | 0.0002377 | -34.458941 | 6.95958111 | 7.37E-07 | 1.25E-06 |
| rs7015194 | 8 | 82800296 | SNX16(dist=45775),LOC101927141(dist=1024043) | C | T | 0.0002377 | -34.458941 | 6.95958111 | 7.37E-07 | 1.25E-06 |
| rs148381942 | 8 | 82805168 | SNX16(dist=50647),LOC101927141(dist=1019171) | T | C | 0.0002377 | -34.458941 | 6.95958111 | 7.37E-07 | 1.25E-06 |
| rs142735588 | 8 | 82807786 | SNX16(dist=53265),LOC101927141(dist=1016553) | G | C | 0.0002377 | -34.458941 | 6.95958111 | 7.37E-07 | 1.25E-06 |
| rs143666707 | 8 | 98883291 | MATN2 | A | C | 0.0002377 | -34.458941 | 6.95958111 | 7.37E-07 | 1.25E-06 |
| rs143416200 | 8 | 98889339 | MATN2 | A | G | 0.0002377 | -34.458941 | 6.95958111 | 7.37E-07 | 1.25E-06 |
| rs183597429 | 8 | 98898180 | MATN2 | C | G | 0.0002377 | -34.458941 | 6.95958111 | 7.37E-07 | 1.25E-06 |
| rs73278205 | 8 | 98948761 | MATN2 | C | A | 0.0002377 | -34.458941 | 6.95958111 | 7.37E-07 | 1.25E-06 |
| rs148205164 | 8 | 98977355 | MATN2 | C | A | 0.0002377 | -34.458941 | 6.95958111 | 7.37E-07 | 1.25E-06 |
| rs184388695 | 8 | 98982974 | MATN2 | G | C | 0.0002377 | -34.458941 | 6.95958111 | 7.37E-07 | 1.25E-06 |
| rs187568831 | 8 | 98986701 | MATN2 | G | A | 0.0002377 | -34.458941 | 6.95958111 | 7.37E-07 | 1.25E-06 |
| rs16974569 | 15 | 85399645 | ALPK3 | C | T | 0.0002377 | -34.458941 | 6.95958111 | 7.37E-07 | 1.25E-06 |

| | | | | | | | | | | |
|---|---|---|---|---|---|---|---|---|---|---|
| rs146989532 | 1 | 163840741 | LOC100422212(dist=447760),PBX1(dist=687856) | G | A | 0.00023806 | -34.458827 | 6.95958485 | 7.37E-07 | 1.25E-06 |
| rs76606125 | 1 | 163841505 | LOC100422212(dist=448524),PBX1(dist=687092) | C | T | 0.00023806 | -34.458827 | 6.95958485 | 7.37E-07 | 1.25E-06 |
| rs115309715 | 1 | 162712336 | DDR2 | A | G | 0.00023782 | -34.45866 | 6.95957108 | 7.37E-07 | 1.25E-06 |
| rs11484775 | 1 | 162719087 | DDR2 | G | C | 0.00023782 | -34.45866 | 6.95957108 | 7.37E-07 | 1.25E-06 |
| rs78818485 | 1 | 162687331 | DDR2 | C | G | 0.00023877 | -34.456448 | 6.95951769 | 7.38E-07 | 1.25E-06 |
| rs112165405 | 1 | 164617327 | PBX1 | C | T | 0.00026789 | -34.314432 | 6.93120796 | 7.39E-07 | 1.25E-06 |
| rs117966882 | 15 | 82396031 | LINC01583(dist=5997),EFL1(dist=26530) | G | A | 0.00023829 | -35.793198 | 7.23013742 | 7.40E-07 | 1.25E-06 |
| rs558451449 | 1 | 104900815 | LOC100129138(dist=281122),LINC01676(dist=1231501) | T | C | 0.00024067 | -34.45308 | 6.95956654 | 7.40E-07 | 1.25E-06 |
| rs6659194 | 1 | 162685364 | DDR2 | G | A | 0.00023889 | -34.452772 | 6.95950493 | 7.40E-07 | 1.25E-06 |
| rs115098909 | 1 | 162714006 | DDR2 | G | A | 0.00024055 | -34.449397 | 6.95954564 | 7.42E-07 | 1.25E-06 |
| rs140441781 | 8 | 82826977 | SNX16(dist=72456),LOC101927141(dist=997362) | C | T | 0.00023806 | -34.449346 | 6.95956887 | 7.42E-07 | 1.25E-06 |
| rs192033741 | 8 | 82827375 | SNX16(dist=72854),LOC101927141(dist=996964) | C | T | 0.00023806 | -34.449346 | 6.95956887 | 7.42E-07 | 1.25E-06 |
| rs73029638 | 1 | 163534200 | LOC100422212(dist=141219),PBX1(dist=994397) | T | C | 0.00023913 | -34.443715 | 6.95933878 | 7.45E-07 | 1.25E-06 |
| rs142466649 | 1 | 162674578 | DDR2 | C | T | 0.00023806 | -34.444409 | 6.95962147 | 7.45E-07 | 1.25E-06 |
| rs150484952 | 1 | 163490907 | LOC100422212(dist=97926),PBX1(dist=1037690) | A | G | 0.00023782 | -34.526548 | 6.97929268 | 7.54E-07 | 1.25E-06 |
| rs16855210 | 1 | 163479854 | LOC100422212(dist=86873),PBX1(dist=1048743) | G | T | 0.0002371 | -34.519886 | 6.97931482 | 7.57E-07 | 1.25E-06 |
| rs16855237 | 1 | 163480472 | LOC100422212(dist=87491),PBX1(dist=1048125) | C | A | 0.0002371 | -34.519886 | 6.97931482 | 7.57E-07 | 1.25E-06 |
| rs7526481 | 1 | 163484242 | LOC100422212(dist=91261),PBX1(dist=1044355) | C | T | 0.0002371 | -34.519886 | 6.97931482 | 7.57E-07 | 1.25E-06 |
| rs146563539 | 1 | 163486488 | LOC100422212(dist=93507),PBX1(dist=1042109) | C | G | 0.0002371 | -34.519886 | 6.97931482 | 7.57E-07 | 1.25E-06 |
| rs111461063 | 1 | 163490575 | LOC100422212(dist=97594),PBX1(dist=1038022) | G | A | 0.0002371 | -34.519886 | 6.97931482 | 7.57E-07 | 1.25E-06 |
| rs535373912 | 20 | 29991638 | DEFB119(dist=13186),DEFB121(dist=1010) | G | A | 0.00047255 | -23.555713 | 4.76556589 | 7.70E-07 | 1.29E-06 |
| rs761025544 | 18 | 72134950 | FAM69C(dist=10447),CNDP2(dist=28550) | C | T | 0.001285 | 16.2387518 | 3.28542535 | 7.71E-07 | 2.74E-06 |
| rs530243109 | 3 | 169314534 | MECOM | C | T | 0.00024329 | -34.391016 | 6.95951122 | 7.75E-07 | 1.25E-06 |
| rs140760764 | 2 | 103757580 | LINC01935(dist=156693),LOC100287010(dist=1237728) | C | G | 0.00038377 | 30.9994702 | 6.27491636 | 7.80E-07 | 2.41E-06 |
| rs529668978 | 20 | 29526419 | LINC01597(dist=5206),LINC01598(dist=32069) | G | T | 0.00047064 | -23.510674 | 4.76114437 | 7.89E-07 | 1.29E-06 |
| rs557067262 | 2 | 14741334 | LINC00276(dist=200252),FAM84A(dist=31473) | T | C | 0.00028322 | -33.498581 | 6.78441354 | 7.91E-07 | 1.25E-06 |
| rs986373288 | 3 | 87267145 | LINC00506(dist=60926),MIR4795(dist=8194) | G | C | 0.00022522 | -38.713627 | 7.84219289 | 7.95E-07 | 2.06E-07 |
| rs142267675 | 8 | 99053275 | RPL30(dist=663) | C | T | 0.00024958 | -34.355808 | 6.96158735 | 8.01E-07 | 1.25E-06 |
| rs114241402 | 1 | 163493815 | LOC100422212(dist=100834),PBX1(dist=1034782) | G | A | 0.00023592 | -34.670116 | 7.02684973 | 8.06E-07 | 1.25E-06 |
| rs529459930 | 6 | 151843034 | CCDC170 | C | T | 0.00044367 | 29.4049427 | 5.95993062 | 8.07E-07 | 2.41E-06 |
| rs541726732 | 20 | 29962442 | DEFB118(dist=737) | G | T | 0.00048835 | -23.325307 | 4.72905951 | 8.13E-07 | 1.29E-06 |
| rs554265290 | 1 | 41219161 | NFYC | G | A | 0.00025256 | -35.828015 | 7.26408455 | 8.13E-07 | 1.25E-06 |
| rs7520505 | 1 | 162669855 | DDR2 | T | C | 0.00024222 | -34.311934 | 6.95856928 | 8.19E-07 | 1.25E-06 |
| rs144763189 | 2 | 103366466 | TMEM182 | A | G | 0.00050452 | 25.7904689 | 5.23562306 | 8.39E-07 | 8.40E-07 |
| rs141219189 | 2 | 103429638 | TMEM182 | T | G | 0.00049418 | 25.8306242 | 5.24387815 | 8.40E-07 | 8.40E-07 |
| rs150243641 | 7 | 20118216 | LOC101927668 | G | A | 0.00025089 | -34.429728 | 6.99012447 | 8.42E-07 | 1.25E-06 |
| rs564609140 | 19 | 22785642 | GOLGA2P9 | G | A | 0.00051212 | -24.580583 | 4.9935833 | 8.55E-07 | 1.58E-06 |
| rs10399864 | 1 | 163473431 | LOC100422212(dist=80450),PBX1(dist=1055166) | A | T | 0.00024031 | -34.332355 | 6.97883769 | 8.68E-07 | 1.25E-06 |
| rs145330758 | 8 | 98907261 | MATN2 | C | T | 0.00024673 | -34.22206 | 6.956911 | 8.69E-07 | 1.25E-06 |
| rs371781211 | 8 | 73888194 | KCNB2(dist=37610),TERF1(dist=32903) | C | A | 0.00025125 | -34.218823 | 6.95739115 | 8.73E-07 | 1.25E-06 |
| rs76625767 | 2 | 103393185 | TMEM182 | G | A | 0.0004975 | 25.7569808 | 5.2399385 | 8.86E-07 | 8.40E-07 |
| rs139275277 | 20 | 13137998 | SPTLC3 | C | T | 0.00038686 | 29.7837011 | 6.06108395 | 8.93E-07 | 2.41E-06 |
| rs565662642 | 6 | 69766487 | ADGRB3 | T | C | 0.00055443 | 24.2462996 | 4.93494994 | 8.96E-07 | 7.72E-07 |
| rs539822723 | 9 | 81426863 | PSAT1(dist=481854),LOC101927450(dist=323475) | C | T | 0.000244 | -34.948018 | 7.11432204 | 9.00E-07 | 1.25E-06 |
| rs373434487 | 6 | 69675348 | ADGRB3 | G | T | 0.00057095 | 24.1738575 | 4.92120672 | 9.01E-07 | 7.72E-07 |
| rs188505066 | 16 | 60432138 | LOC729159(dist=38441),MIR4426(dist=657473) | C | G | 0.0003909 | -28.05164 | 5.71096493 | 9.02E-07 | 1.98E-06 |
| rs75684812 | 1 | 163525360 | LOC100422212(dist=132379),PBX1(dist=1003237) | G | A | 0.00021429 | -38.612898 | 7.86905874 | 9.25E-07 | 1.25E-06 |
| rs770048470 | 6 | 77378157 | IMPG1(dist=595762),HTR1B(dist=792408) | G | T | 0.00024768 | -34.132208 | 6.95875143 | 9.35E-07 | 1.25E-06 |
| rs552210261 | 12 | 92706705 | LOC101928617(dist=127145),CLLU1OS(dist=107165) | C | T | 0.00025814 | -35.980309 | 7.34154955 | 9.54E-07 | 1.25E-06 |
| rs988902039 | 4 | 184718514 | TRAPPC11(dist=83767),NONE(dist=NONE) | C | T | 0.00061683 | -22.217371 | 4.53423714 | 9.59E-07 | 1.29E-06 |
| rs185185799 | 9 | 74187011 | TRPM3(dist=450497),TMEM2(dist=111271) | A | T | 0.0018922 | 13.2146748 | 2.70019572 | 9.88E-07 | 3.08E-06 |
| rs117326470 | 2 | 103722596 | LINC01935(dist=121709),LOC100287010(dist=1272712) | A | G | 0.00047195 | 26.3500912 | 5.38491327 | 9.92E-07 | 8.40E-07 |

| | | | | | | | | | | |
|---|---|---|---|---|---|---|---|---|---|---|
| rs570203963 | 12 | 66271102 | HMGA2,LOC100129940 | T | C | 0.00027466 | -34.351772 | 7.0204864 | 9.93E-07 | 1.25E-06 |
| rs9301165 | 13 | 107437444 | LINC00443(dist=112916),FAM155A(dist=383435) | T | C | 0.00049762 | 24.4630262 | 5.00291173 | 1.01E-06 | 1.18E-06 |
| rs538595234 | 14 | 101390810 | MEG8 | A | C | 0.00121702 | 16.5891323 | 3.39291197 | 1.01E-06 | 4.49E-07 |
| rs1934529 | 1 | 163553145 | LOC100422212(dist=160164),PBX1(dist=975452) | T | C | 0.00025208 | -33.877842 | 6.93741323 | 1.04E-06 | 1.25E-06 |
| rs112182732 | 1 | 163649757 | LOC100422212(dist=256776),PBX1(dist=878840) | T | C | 0.00023259 | -35.535886 | 7.2854105 | 1.07E-06 | 1.25E-06 |
| rs551533844 | 8 | 74105341 | SBSPON(dist=99834),C8orf89(dist=48318) | C | T | 0.00026753 | -33.914524 | 6.95533243 | 1.08E-06 | 1.25E-06 |
| rs373074099 | 18 | 72153912 | FAM69C(dist=29409),CNDP2(dist=9588) | G | A | 0.00127989 | 16.0312003 | 3.28889028 | 1.09E-06 | 2.74E-06 |
| rs535347456 | 8 | 32294257 | NRG1 | G | C | 0.00092786 | 19.8221326 | 4.06684814 | 1.09E-06 | 1.07E-06 |
| rs370514761 | 16 | 81922041 | PLCG2 | A | G | 0.00077953 | 21.3645502 | 4.38340206 | 1.09E-06 | 5.50E-07 |
| rs776395852 | 5 | 95526964 | LOC101929710 | G | A | 0.00026801 | -34.323581 | 7.04252488 | 1.09E-06 | 1.25E-06 |
| rs114723842 | 1 | 163498600 | LOC100422212(dist=105619),PBX1(dist=1029997) | T | C | 0.00022974 | -35.366281 | 7.26893853 | 1.14E-06 | 1.25E-06 |
| rs13183308 | 5 | 163996922 | LOC102546299(dist=26933),NONE(dist=NONE) | C | G | 0.20474745 | 1.34540704 | 0.2765424 | 1.14E-06 | 7.75E-07 |
| rs541787240 | 22 | 32378798 | YWHAH(dist=25208),SLC5A1(dist=60221) | C | T | 0.0004773 | -24.451356 | 5.02851692 | 1.16E-06 | 2.70E-06 |
| rs59494028 | 13 | 107436325 | LINC00443(dist=111797),FAM155A(dist=384554) | C | T | 0.0004773 | 24.3358562 | 5.01114243 | 1.20E-06 | 1.18E-06 |
| rs7544144 | 1 | 163059868 | RGS4(dist=13276),RGS5(dist=52221) | A | G | 0.00011932 | -43.87206 | 9.03674745 | 1.20E-06 | 4.05E-06 |
| rs772781056 | 5 | 52021339 | LINC02118(dist=681596),PELO(dist=62435) | G | T | 0.00025386 | -33.761228 | 6.95506994 | 1.21E-06 | 1.25E-06 |
| rs192971782 | 2 | 96901964 | STARD7-AS1 | A | C | 0.00179629 | 13.0119163 | 2.68095232 | 1.21E-06 | 2.79E-06 |
| rs148137226 | 1 | 163489113 | LOC100422212(dist=96132),PBX1(dist=1039484) | A | G | 0.00025351 | -33.87147 | 6.98862874 | 1.26E-06 | 1.25E-06 |
| rs114112804 | 4 | 184756015 | NONE(dist=NONE),NONE(dist=NONE) | G | A | 0.00082517 | -19.843876 | 4.09535039 | 1.26E-06 | 7.83E-07 |
| rs542970754 | 6 | 71450425 | SMAP1 | A | G | 0.00027086 | -33.60189 | 6.93635364 | 1.27E-06 | 1.25E-06 |
| rs765061610 | 1 | 90492457 | ZNF326 | T | C | 0.00033254 | -29.909125 | 6.17556991 | 1.28E-06 | 3.57E-06 |
| rs897672626 | 6 | 69633940 | ADGRB3 | A | T | 0.00048586 | 24.3390472 | 5.0276045 | 1.29E-06 | 7.72E-07 |
| rs554968707 | 21 | 19259570 | CHODL-AS1(dist=1645),CHODL(dist=30087) | G | A | 0.00012598 | -42.160235 | 8.71508141 | 1.31E-06 | 4.05E-06 |
| rs142156809 | 21 | 31517942 | GRIK1(dist=205572),CLDN17(dist=20299) | A | C | 0.00039018 | 29.0458869 | 6.0070971 | 1.33E-06 | 2.41E-06 |
| rs753990222 | 1 | 193815600 | LINC01031(dist=480517),NONE(dist=NONE) | A | G | 0.00126337 | 16.0339954 | 3.31786019 | 1.35E-06 | 1.08E-06 |
| rs544558253 | 20 | 13123525 | SPTLC3 | G | A | 0.00037628 | 29.3189322 | 6.06740022 | 1.35E-06 | 2.41E-06 |
| | 1 | 100032890 | LINC01708(dist=79530),PALMD(dist=78541) | G | A | 0.00025386 | -33.982385 | 7.04092559 | 1.39E-06 | 1.25E-06 |
| rs190780367 | 2 | 69935425 | ANXA4 | G | T | 0.00011861 | -43.959781 | 9.11923394 | 1.43E-06 | 4.05E-06 |
| rs566658574 | 12 | 88176259 | MKRN9P(dist=404) | T | C | 0.00027537 | -33.470249 | 6.94990996 | 1.47E-06 | 1.25E-06 |
| rs115257980 | 12 | 4489564 | FGF23(dist=670) | G | A | 0.00012313 | -41.988135 | 8.72126861 | 1.48E-06 | 4.05E-06 |
| rs330083 | 8 | 9152654 | LOC101929128(dist=92288),LOC157273(dist=29907) | G | A | 0.94746589 | -2.423227 | 0.50339097 | 1.48E-06 | 2.17E-06 |
| rs75708071 | 1 | 163040464 | RGS4 | T | C | 0.00012099 | -41.790493 | 8.68283059 | 1.49E-06 | 4.05E-06 |
| rs145480617 | 2 | 26707907 | OTOF | G | A | 0.00473794 | 8.61747391 | 1.79072672 | 1.49E-06 | 2.61E-07 |
| rs543850521 | 19 | 495019 | ODF3L2(dist=20036),MADCAM1(dist=1471) | G | A | 0.00035393 | 30.7615115 | 6.39353466 | 1.50E-06 | 2.41E-06 |
| rs141858010 | 2 | 152193397 | LOC101929319(dist=945) | C | A | 9.92E-05 | -50.827765 | 10.5758011 | 1.54E-06 | 4.05E-06 |
| rs147621649 | 11 | 43162092 | LOC100507205(dist=886852),HNRNPKP3(dist=120962) | C | G | 0.00045198 | 25.8656321 | 5.38196741 | 1.54E-06 | 4.23E-06 |
| rs532779237 | 12 | 63527786 | PPM1H(dist=199121),AVPR1A(dist=8753) | T | C | 0.00039696 | 28.7727162 | 6.00404643 | 1.65E-06 | 2.41E-06 |
| rs142516872 | 14 | 55235501 | SAMD4A | G | A | 0.00024899 | -38.656101 | 8.06752405 | 1.65E-06 | 9.95E-07 |
| rs149781560 | 8 | 83141316 | SNX16(dist=386795),LOC101927141(dist=683023) | C | T | 0.00026896 | -33.160826 | 6.9222573 | 1.66E-06 | 1.25E-06 |
| rs149603239 | 1 | 104863243 | LOC100129138(dist=243550),LINC01676(dist=1269073) | G | A | 0.00036582 | 29.3649495 | 6.13806177 | 1.72E-06 | 2.41E-06 |
| rs544725047 | 2 | 96828129 | DUSP2(dist=16950),STARD7(dist=22474) | C | T | 0.00138864 | 14.9559866 | 3.12674494 | 1.72E-06 | 2.25E-07 |
| rs996561540 | 15 | 34314013 | AVEN,CHRM5 | G | A | 0.00011909 | -42.464538 | 8.87975838 | 1.73E-06 | 4.05E-06 |
| rs147491961 | 1 | 99946174 | LINC01708 | G | T | 0.00011517 | -43.029045 | 9.00143483 | 1.75E-06 | 4.05E-06 |
| rs146824245 | 1 | 99928302 | PLPPR4(dist=153164),LINC01708(dist=9674) | A | C | 0.00011481 | -42.997553 | 9.00149628 | 1.78E-06 | 4.05E-06 |
| rs549491075 | 2 | 52691266 | LOC730100(dist=56211),MIR4431(dist=238394) | G | A | 0.00036713 | 28.908326 | 6.05280122 | 1.79E-06 | 2.41E-06 |
| rs527764081 | 2 | 52691238 | LOC730100(dist=56183),MIR4431(dist=238422) | G | A | 0.00036701 | 28.9024153 | 6.05301656 | 1.80E-06 | 2.41E-06 |
| rs948160859 | 18 | 71588667 | LOC100505817(dist=571543),FBXO15(dist=151921) | G | A | 0.00053007 | 24.5627831 | 5.14548356 | 1.81E-06 | 1.75E-06 |
| rs1918085 | 2 | 52700429 | LOC730100(dist=65374),MIR4431(dist=229231) | A | G | 0.00036808 | 28.8743673 | 6.05362583 | 1.84E-06 | 2.41E-06 |
| rs12086493 | 1 | 163032285 | CCDC190(dist=193680),RGS4(dist=6111) | A | G | 0.0001217 | -41.65634 | 8.74082707 | 1.88E-06 | 4.05E-06 |
| rs148414063 | 9 | 124091618 | GSN | C | G | 0.00033551 | 28.5053331 | 5.98382007 | 1.90E-06 | 2.51E-06 |
| rs115070743 | 1 | 163502305 | LOC100422212(dist=109324),PBX1(dist=1026292) | G | A | 0.00023199 | -35.719554 | 7.49833831 | 1.90E-06 | 1.25E-06 |
| rs183898472 | 2 | 96840725 | DUSP2(dist=29546),STARD7(dist=9878) | C | T | 0.00188733 | 12.2154174 | 2.56801439 | 1.97E-06 | 2.79E-06 |

| | | | | | | | | | | |
|---|---|---|---|---|---|---|---|---|---|---|
| rs545083715 | 18 | 72851707 | ZNF407(dist=74079),ZADH2(dist=55358) | C | T | 0.00041669 | 28.3391224 | 5.96378421 | 2.02E-06 | 2.41E-06 |
| rs101445405 | 12 | 87220823 | MGAT4C | A | G | 0.00041336 | 29.3959673 | 6.1862781 | 2.02E-06 | 2.41E-06 |
| rs190509573 | 1 | 104890979 | LOC100129138(dist=271286),LINC01676(dist=1241337) | A | T | 0.00035857 | 29.3982357 | 6.19033126 | 2.04E-06 | 2.41E-06 |
| rs188748319 | 11 | 43155320 | LOC100507205(dist=880080),HNRNPKP3(dist=127734) | G | A | 0.00047516 | 25.2886756 | 5.32580729 | 2.05E-06 | 4.23E-06 |
| rs534827699 | 1 | 104801983 | LOC100129138(dist=182290),LINC01676(dist=1330333) | C | A | 0.00037129 | 28.8578382 | 6.08452501 | 2.11E-06 | 2.41E-06 |
| rs941006524 | 1 | 196040073 | LINC01724 | G | A | 0.00081959 | 20.2028284 | 4.26229006 | 2.14E-06 | 5.48E-06 |
| rs114883353 | 17 | 30026956 | MIR365B(dist=124416),COPRS(dist=151928) | G | A | 0.00040231 | 28.4017099 | 5.9967004 | 2.18E-06 | 2.41E-06 |
| rs58050068 | 1 | 99801255 | PLPPR4(dist=26117),LINC01708(dist=136721) | A | G | 0.00011314 | -45.539645 | 9.6199479 | 2.20E-06 | 4.05E-06 |
| rs116727144 | 21 | 19431222 | CHODL | G | A | 0.00014203 | -41.046668 | 8.67215906 | 2.21E-06 | 4.05E-06 |
| rs114438921 | 19 | 549382 | GZMM | G | T | 0.00039256 | 28.605808 | 6.04630814 | 2.23E-06 | 2.41E-06 |
| rs565163634 | 9 | 122593493 | BRINP1(dist=461754),LINC01613(dist=103845) | T | C | 0.00081899 | 19.6370012 | 4.1509705 | 2.24E-06 | 2.71E-06 |
| rs116760832 | 1 | 99860364 | PLPPR4(dist=85226),LINC01708(dist=77612) | T | C | 0.00012135 | -41.693153 | 8.81544954 | 2.25E-06 | 4.05E-06 |
| rs146565413 | 17 | 30027922 | MIR365B(dist=125382),COPRS(dist=150962) | A | G | 0.00038864 | 28.3945079 | 6.013334 | 2.34E-06 | 2.41E-06 |
| rs105235661 | 11 | 120408875 | GRIK4 | C | T | 0.00038127 | 28.6465054 | 6.06947519 | 2.36E-06 | 2.41E-06 |
| rs73275479 | 8 | 82438256 | FABP12 | T | G | 0.00035655 | -26.053877 | 5.52191059 | 2.38E-06 | 4.43E-06 |
| rs530523771 | 11 | 106342138 | LOC101928535(dist=206506),GUCY1A2(dist=202600) | G | T | 0.00387675 | 8.3558437 | 1.77136258 | 2.39E-06 | 4.25E-06 |
| rs73336159 | 5 | 176137439 | TSPAN17(dist=51380),LINC01574(dist=32767) | C | T | 0.0005246 | 23.3514285 | 4.95058561 | 2.39E-06 | 3.94E-06 |
| rs537086621 | 6 | 87537334 | SNHG5(dist=1148883),HTR1E(dist=109690) | A | C | 0.00093867 | 18.3742885 | 3.89753553 | 2.43E-06 | 4.27E-06 |
| rs113316432 | 8 | 82434557 | FABP4(dist=39084),FABP12(dist=2659) | C | G | 0.00035845 | -26.019234 | 5.5218607 | 2.45E-06 | 4.43E-06 |
| rs527468128 | 2 | 153245962 | FMNL2 | A | G | 0.00033682 | 28.9692581 | 6.15147553 | 2.49E-06 | 3.13E-06 |
| rs111977451 | 17 | 39116513 | KRT39 | G | A | 0.00379677 | -8.6522416 | 1.83783192 | 2.50E-06 | 4.67E-07 |
| rs780871194 | 14 | 103486835 | CDC42BPB | C | T | 0.00037556 | 28.4989896 | 6.05350351 | 2.50E-06 | 2.41E-06 |
| rs363072 | 4 | 3142528 | HTT | A | T | 0.14154956 | 1.48439626 | 0.3153699 | 2.52E-06 | 2.64E-06 |
| rs552877239 | 17 | 1290529 | YWHAE | G | A | 0.00161861 | 13.3221752 | 2.83068757 | 2.52E-06 | 2.35E-06 |
| rs560162233 | 2 | 52690443 | LOC730100(dist=55388),MIR4431(dist=239217) | A | G | 0.00036059 | 28.5168083 | 6.0617597 | 2.55E-06 | 2.41E-06 |
| rs190859994 | 1 | 215169142 | CENPF(dist=331228),KCNK2(dist=9743) | T | C | 0.00063917 | -21.788662 | 4.631626 | 2.55E-06 | 8.71E-07 |
| rs115700753 | 17 | 30029860 | MIR365B(dist=127320),COPRS(dist=149024) | T | C | 0.00038091 | 28.4313319 | 6.04940232 | 2.60E-06 | 2.41E-06 |
| rs74095773 | 12 | 60226569 | SLC16A7(dist=42934),FAM19A2(dist=1875460) | C | T | 0.00039244 | 28.4683205 | 6.05809666 | 2.61E-06 | 2.41E-06 |
| rs139487541 | 2 | 52526697 | LOC730100 | C | T | 0.00036095 | 28.4839973 | 6.06209242 | 2.62E-06 | 2.41E-06 |
| rs560100734 | 2 | 52700082 | LOC730100(dist=65027),MIR4431(dist=229578) | A | G | 0.0003594 | 28.4777436 | 6.06210674 | 2.63E-06 | 2.41E-06 |
| rs539473468 | 2 | 172183467 | METTL8 | T | C | 0.00039244 | 28.8632269 | 6.14555529 | 2.65E-06 | 2.41E-06 |
| rs181788788 | 2 | 97464013 | CNNM4 | C | A | 0.00440492 | 8.466316 | 1.80267113 | 2.65E-06 | 8.18E-06 |
| rs184293531 | 2 | 59037301 | LINC01122 | T | C | 0.00036237 | 28.4900975 | 6.06684188 | 2.65E-06 | 2.41E-06 |
| rs768176549 | 6 | 151784878 | ARMT1 | C | G | 0.00041205 | 28.3003897 | 6.03088104 | 2.70E-06 | 2.41E-06 |
| rs549162189 | 9 | 117659401 | TNFSF8 | G | A | 0.00043 | 28.3516646 | 6.04194096 | 2.70E-06 | 2.41E-06 |
| rs113523800 | 19 | 428527 | SHC2 | G | A | 0.00036059 | 29.2324769 | 6.23019649 | 2.70E-06 | 2.41E-06 |
| rs11154688 | 6 | 132912878 | TAAR5(dist=2001),TAAR3P(dist=16486) | G | A | 0.31896042 | 1.13178563 | 0.24121599 | 2.71E-06 | 3.45E-06 |
| rs552544727 | 19 | 460642 | SHC2 | C | A | 0.00040991 | 28.3765078 | 6.04790684 | 2.71E-06 | 2.41E-06 |
| rs532102415 | 2 | 53190323 | MIR4431(dist=260570),ASB3(dist=706794) | T | C | 0.00036511 | 28.4134733 | 6.06140554 | 2.76E-06 | 2.41E-06 |
| rs569942221 | 2 | 53191303 | MIR4431(dist=261550),ASB3(dist=705814) | C | T | 0.00036511 | 28.4134733 | 6.06140554 | 2.76E-06 | 2.41E-06 |
| rs556549829 | 2 | 53193143 | MIR4431(dist=263390),ASB3(dist=703974) | A | C | 0.00036511 | 28.4134733 | 6.06140554 | 2.76E-06 | 2.41E-06 |
| rs567670762 | 2 | 52692097 | LOC730100(dist=57042),MIR4431(dist=237563) | A | C | 0.00035845 | 28.4183222 | 6.06259036 | 2.77E-06 | 2.41E-06 |
| rs558373680 | 2 | 52697272 | LOC730100(dist=62217),MIR4431(dist=232388) | G | T | 0.00035845 | 28.4183222 | 6.06259036 | 2.77E-06 | 2.41E-06 |
| rs561772098 | 2 | 53168754 | MIR4431(dist=239001),ASB3(dist=728363) | G | A | 0.00036118 | 28.4128972 | 6.06213807 | 2.77E-06 | 2.41E-06 |
| rs766959296 | 17 | 16996395 | MPRIP | G | A | 0.00037224 | 28.399233 | 6.05978828 | 2.78E-06 | 2.41E-06 |
| rs113793355 | 19 | 483664 | ODF3L2(dist=8681),MADCAM1(dist=12826) | G | A | 0.00035298 | 28.9094581 | 6.16886881 | 2.78E-06 | 2.41E-06 |
| rs369292656 | 2 | 52727688 | LOC730100(dist=92633),MIR4431(dist=201972) | G | A | 0.00039125 | 28.3350972 | 6.0468007 | 2.79E-06 | 2.41E-06 |
| rs181380052 | 2 | 59958310 | LINC01793(dist=451775),MIR4432HG(dist=628041) | A | G | 0.00055384 | 23.5652636 | 5.02914115 | 2.79E-06 | 2.75E-06 |
| rs185758937 | 2 | 59958337 | LINC01793(dist=451802),MIR4432HG(dist=628014) | C | G | 0.00055384 | 23.5624977 | 5.02892977 | 2.79E-06 | 2.75E-06 |
| rs77173096 | 1 | 97893807 | DPYD | G | A | 0.00033159 | 31.0809784 | 6.63362298 | 2.79E-06 | 2.41E-06 |
| rs578084149 | 2 | 53184688 | MIR4431(dist=254935),ASB3(dist=712429) | T | A | 0.00036368 | 28.3957548 | 6.06189886 | 2.81E-06 | 2.41E-06 |
| rs551354564 | 2 | 52689780 | LOC730100(dist=54725),MIR4431(dist=239880) | A | T | 0.00035809 | 28.395682 | 6.0627342 | 2.82E-06 | 2.41E-06 |

| | | | | | | | | | | |
|---|---|---|---|---|---|---|---|---|---|---|
| rs545597295 | 2 | 53184895 | MIR4431(dist=255142),ASB3(dist=712222) | C | T | 0.00036332 | 28.3912017 | 6.06200923 | 2.82E-06 | 2.41E-06 |
| rs100636603 | 9 | 74292533 | TRPM3(dist=556019),TMEM2(dist=5749) | G | A | 0.00036237 | 28.3885686 | 6.06243128 | 2.83E-06 | 2.41E-06 |
| rs77319082 | 1 | 163653543 | LOC100422212(dist=260562),PBX1(dist=875054) | G | A | 0.00025719 | -33.860561 | 7.23118997 | 2.83E-06 | 1.25E-06 |
| rs562402350 | 2 | 52655614 | LOC730100(dist=20559),MIR4431(dist=274046) | G | A | 0.00036594 | 28.3780455 | 6.06199554 | 2.85E-06 | 2.41E-06 |
| rs7850758 | 9 | 140188225 | TOR4A(dist=11132),NRARP(dist=5858) | A | G | 0.07715023 | 1.92178742 | 0.41054849 | 2.85E-06 | 1.52E-06 |
| rs151263175 | 17 | 30061792 | MIR365B(dist=159252),COPRS(dist=117092) | G | A | 0.0003714 | 28.3722071 | 6.06240982 | 2.87E-06 | 2.41E-06 |
| rs183211857 | 5 | 180640700 | TRIM7(dist=8407),MIR4638(dist=8866) | T | C | 0.00036011 | 28.3709746 | 6.06255292 | 2.87E-06 | 2.41E-06 |
| rs12197997 | 6 | 132919550 | TAAR5(dist=8673),TAAR3P(dist=9814) | A | G | 0.31863644 | 1.12785575 | 0.24102657 | 2.88E-06 | 3.66E-06 |
| rs17119918 | 11 | 116585613 | LOC101929011(dist=56644),BUD13(dist=33273) | G | A | 0.0003638 | 28.3525801 | 6.05969123 | 2.88E-06 | 2.41E-06 |
| rs7257230 | 19 | 454351 | SHC2 | C | A | 0.00036748 | 28.3601014 | 6.06131976 | 2.88E-06 | 2.41E-06 |
| rs113851346 | 17 | 30035361 | MIR365B(dist=132821),COPRS(dist=143523) | G | A | 0.00036332 | 28.3631113 | 6.06275019 | 2.89E-06 | 2.41E-06 |
| rs7256741 | 19 | 453898 | SHC2 | G | A | 0.00037283 | 28.3514072 | 6.06069003 | 2.90E-06 | 2.41E-06 |
| rs531114386 | 2 | 52609783 | LINC01867,LOC730100 | T | G | 0.0003594 | 28.3492702 | 6.06311435 | 2.93E-06 | 2.41E-06 |
| rs559962898 | 2 | 52606500 | LINC01867,LOC730100 | G | A | 0.00036356 | 28.3481719 | 6.06305118 | 2.93E-06 | 2.41E-06 |
| rs185741126 | 5 | 180611687 | OR2V2(dist=28797),LINC01962(dist=6359) | C | T | 0.00036083 | 28.3431498 | 6.062525 | 2.94E-06 | 2.41E-06 |
| rs536174889 | 2 | 53182125 | MIR4431(dist=252372),ASB3(dist=714992) | A | G | 0.00035976 | 28.3429583 | 6.06282784 | 2.94E-06 | 2.41E-06 |
| rs534063655 | 2 | 52525631 | LOC730100 | G | A | 0.00038614 | 28.3381435 | 6.06210819 | 2.94E-06 | 2.41E-06 |
| rs186240407 | 2 | 53136950 | MIR4431(dist=207197),ASB3(dist=760167) | C | T | 0.00036665 | 28.3354378 | 6.0621346 | 2.95E-06 | 2.41E-06 |
| rs563891857 | 2 | 52636044 | LOC730100(dist=989) | T | C | 0.00038377 | 28.3258048 | 6.06027271 | 2.95E-06 | 2.41E-06 |
| rs144023056 | 12 | 63197659 | PPM1H | C | T | 0.00035904 | 28.3376353 | 6.06283817 | 2.95E-06 | 2.41E-06 |
| rs534689729 | 1 | 108007520 | NTNG1 | G | A | 0.00078928 | 20.6627015 | 4.42101326 | 2.96E-06 | 8.57E-06 |
| rs114294515 | 5 | 180619981 | TRIM7(dist=943) | T | C | 0.00036083 | 28.33299 | 6.062616 | 2.96E-06 | 2.41E-06 |
| rs148369445 | 19 | 450882 | SHC2 | G | A | 0.00036701 | 28.3291129 | 6.06179273 | 2.96E-06 | 2.41E-06 |
| rs116838278 | 5 | 180633941 | TRIM7(dist=1648),MIR4638(dist=15625) | T | G | 0.0003613 | 28.3322001 | 6.06260517 | 2.96E-06 | 2.41E-06 |
| rs569238584 | 11 | 133008828 | OPCML | C | T | 0.00040207 | 28.1423882 | 6.02290337 | 2.97E-06 | 2.41E-06 |
| rs9309230 | 2 | 52749068 | LOC730100(dist=114013),MIR4431(dist=180592) | T | C | 0.00035881 | 28.3294443 | 6.06304062 | 2.98E-06 | 2.41E-06 |
| rs148648064 | 19 | 471828 | ODF3L2 | C | T | 0.00036629 | 28.3159937 | 6.06016254 | 2.98E-06 | 2.41E-06 |
| rs370077527 | 2 | 53107179 | MIR4431(dist=177426),ASB3(dist=789938) | T | C | 0.00035738 | 28.3286097 | 6.06317798 | 2.98E-06 | 2.41E-06 |
| rs372194531 | 2 | 52718516 | LOC730100(dist=83461),MIR4431(dist=211144) | G | C | 0.00035786 | 28.3244386 | 6.06312002 | 2.99E-06 | 2.41E-06 |
| rs565615053 | 2 | 53181004 | MIR4431(dist=251251),ASB3(dist=716113) | C | G | 0.00035833 | 28.3222833 | 6.06301002 | 2.99E-06 | 2.41E-06 |
| rs1526679 | 2 | 52644317 | LOC730100(dist=9262),MIR4431(dist=285343) | A | G | 0.00035845 | 28.3197332 | 6.06289174 | 3.00E-06 | 2.41E-06 |
| rs548028748 | 2 | 52720097 | LOC730100(dist=85042),MIR4431(dist=209563) | G | A | 0.0003575 | 28.3203322 | 6.06310871 | 3.00E-06 | 2.41E-06 |
| rs532715151 | 2 | 52720079 | LOC730100(dist=85024),MIR4431(dist=209581) | T | G | 0.00035714 | 28.3181464 | 6.06312073 | 3.00E-06 | 2.41E-06 |
| rs369287560 | 2 | 53023383 | MIR4431(dist=93630),ASB3(dist=873734) | G | A | 0.00035928 | 28.3173974 | 6.06310884 | 3.01E-06 | 2.41E-06 |
| rs75390181 | 2 | 52956699 | MIR4431(dist=26946),ASB3(dist=940418) | G | C | 0.00035762 | 28.3169602 | 6.06305001 | 3.01E-06 | 2.41E-06 |
| rs73489636 | 19 | 452062 | SHC2 | C | T | 0.00036261 | 28.3093117 | 6.06169078 | 3.01E-06 | 2.41E-06 |
| rs369180864 | 2 | 52715235 | LOC730100(dist=80180),MIR4431(dist=214425) | C | G | 0.00035786 | 28.3159295 | 6.06312256 | 3.01E-06 | 2.41E-06 |
| rs528976078 | 2 | 52640735 | LOC730100(dist=5680),MIR4431(dist=288925) | T | G | 0.00035845 | 28.3155665 | 6.06311204 | 3.01E-06 | 2.41E-06 |
| rs542361546 | 2 | 52622737 | LOC730100 | A | G | 0.00035869 | 28.3154966 | 6.06322323 | 3.01E-06 | 2.41E-06 |
| rs369972488 | 2 | 52719190 | LOC730100(dist=84135),MIR4431(dist=210470) | A | G | 0.00035702 | 28.313637 | 6.06312641 | 3.01E-06 | 2.41E-06 |
| rs7251996 | 19 | 482288 | ODF3L2(dist=7305),MADCAM1(dist=14202) | T | C | 0.00032648 | 30.761612 | 6.58735036 | 3.01E-06 | 2.41E-06 |
| rs947140319 | 15 | 40521210 | BUB1B-PAK6 | C | T | 0.00033563 | 28.902322 | 6.18938908 | 3.02E-06 | 2.58E-06 |
| rs559807846 | 19 | 460403 | SHC2 | G | A | 0.00036368 | 28.3021228 | 6.06099812 | 3.02E-06 | 2.41E-06 |
| rs111824506 | 19 | 451666 | SHC2 | C | T | 0.00036225 | 28.3054768 | 6.06179566 | 3.02E-06 | 2.41E-06 |
| rs111572517 | 19 | 451746 | SHC2 | C | T | 0.00036225 | 28.3054768 | 6.06179566 | 3.02E-06 | 2.41E-06 |
| rs796326000 | 19 | 451079 | SHC2 | C | T | 0.00036213 | 28.3056163 | 6.06187252 | 3.02E-06 | 2.41E-06 |
| rs570153897 | 2 | 52610437 | LINC01867,LOC730100 | C | T | 0.00035845 | 28.3119368 | 6.06322685 | 3.02E-06 | 2.41E-06 |
| rs546783102 | 2 | 52871208 | LOC730100(dist=236153),MIR4431(dist=58452) | C | T | 0.00035809 | 28.3108731 | 6.06306094 | 3.02E-06 | 2.41E-06 |
| rs1358186 | 2 | 52642930 | LOC730100(dist=7875),MIR4431(dist=286730) | C | T | 0.00035786 | 28.3104265 | 6.06311632 | 3.02E-06 | 2.41E-06 |
| rs539458152 | 11 | 116520256 | LOC101929011 | C | T | 0.0003575 | 28.3384225 | 6.06914989 | 3.02E-06 | 2.41E-06 |
| rs574775108 | 2 | 52645035 | LOC730100(dist=9980),MIR4431(dist=284625) | C | A | 0.00035762 | 28.3097457 | 6.06306532 | 3.02E-06 | 2.41E-06 |
| rs373811446 | 2 | 52716581 | LOC730100(dist=81526),MIR4431(dist=213079) | A | G | 0.00035714 | 28.3096636 | 6.0631261 | 3.02E-06 | 2.41E-06 |

| | | | | | | | | | | |
|---|---|---|---|---|---|---|---|---|---|---|
| rs367623001 | 2 | 52717034 | LOC730100(dist=81979),MIR4431(dist=212626) | A | G | 0.00035809 | 28.3089014 | 6.06299678 | 3.02E-06 | 2.41E-06 |
| rs115676637 | 19 | 450068 | SHC2 | G | A | 0.00036166 | 28.3041217 | 6.06203686 | 3.03E-06 | 2.41E-06 |
| rs116669807 | 19 | 450165 | SHC2 | G | A | 0.00036166 | 28.3041217 | 6.06203686 | 3.03E-06 | 2.41E-06 |
| rs537003747 | 2 | 55990459 | PNPT1(dist=69414),EFEMP1(dist=102638) | A | G | 0.00035797 | 29.2358314 | 6.26159506 | 3.03E-06 | 2.41E-06 |
| rs369146550 | 2 | 53010458 | MIR4431(dist=80705),ASB3(dist=886659) | G | A | 0.00035988 | 28.3081845 | 6.06302099 | 3.03E-06 | 2.41E-06 |
| rs575452662 | 2 | 52616725 | LINC01867 | G | T | 0.00037699 | 28.3033413 | 6.06224241 | 3.03E-06 | 2.41E-06 |
| rs551597492 | 2 | 52655879 | LOC730100(dist=20824),MIR4431(dist=273781) | C | G | 0.00035904 | 28.3064211 | 6.06299881 | 3.03E-06 | 2.41E-06 |
| rs377200444 | 2 | 52956505 | MIR4431(dist=26752),ASB3(dist=940612) | T | C | 0.00035738 | 28.3064878 | 6.06303593 | 3.03E-06 | 2.41E-06 |
| rs571326552 | 2 | 53180537 | MIR4431(dist=250784),ASB3(dist=716580) | G | A | 0.00035726 | 28.3062618 | 6.06309217 | 3.03E-06 | 2.41E-06 |
| rs960532183 | 9 | 119508058 | ASTN2 | C | T | 0.0003594 | 28.3059101 | 6.06308851 | 3.03E-06 | 2.41E-06 |
| rs371212315 | 2 | 52956080 | MIR4431(dist=26327),ASB3(dist=941037) | T | G | 0.00035702 | 28.3052991 | 6.0630758 | 3.03E-06 | 2.41E-06 |
| rs543762175 | 2 | 52604220 | LINC01867,LOC730100 | T | G | 0.00035833 | 28.3057045 | 6.06323422 | 3.04E-06 | 2.41E-06 |
| rs565285485 | 2 | 52604228 | LINC01867,LOC730100 | A | G | 0.00035833 | 28.3057045 | 6.06323422 | 3.04E-06 | 2.41E-06 |
| rs545828979 | 2 | 52617216 | LOC730100 | G | C | 0.00035833 | 28.3057045 | 6.06323422 | 3.04E-06 | 2.41E-06 |
| rs531226054 | 2 | 52623488 | LOC730100 | T | A | 0.00035833 | 28.3057045 | 6.06323422 | 3.04E-06 | 2.41E-06 |
| rs533062649 | 2 | 52655676 | LOC730100(dist=20621),MIR4431(dist=273984) | A | G | 0.00035797 | 28.3049851 | 6.06309416 | 3.04E-06 | 2.41E-06 |
| rs369033119 | 2 | 52723678 | LOC730100(dist=88623),MIR4431(dist=205982) | C | T | 0.00035679 | 28.3045391 | 6.06312932 | 3.04E-06 | 2.41E-06 |
| rs368825583 | 2 | 52955395 | MIR4431(dist=25642),ASB3(dist=941722) | T | C | 0.00035691 | 28.3035845 | 6.06307861 | 3.04E-06 | 2.41E-06 |
| rs569134995 | 2 | 52955412 | MIR4431(dist=25659),ASB3(dist=941705) | A | G | 0.00035691 | 28.3035845 | 6.06307861 | 3.04E-06 | 2.41E-06 |
| rs2356041 | 2 | 52598458 | LINC01867,LOC730100 | T | C | 0.00035797 | 28.3037521 | 6.06322448 | 3.04E-06 | 2.41E-06 |
| rs564794266 | 2 | 52688608 | LOC730100(dist=53553),MIR4431(dist=241052) | T | G | 0.00035667 | 28.3031834 | 6.06310487 | 3.04E-06 | 2.41E-06 |
| rs8104724 | 19 | 462352 | ODF3L2(dist=994) | G | C | 0.00036439 | 28.2931648 | 6.0610064 | 3.04E-06 | 2.41E-06 |
| rs78935322 | 17 | 30032498 | MIR365B(dist=129958),COPRS(dist=146386) | C | A | 0.00036463 | 28.298501 | 6.06232831 | 3.04E-06 | 2.41E-06 |
| rs367979351 | 2 | 52735275 | LOC730100(dist=100220),MIR4431(dist=194385) | G | A | 0.00035691 | 28.302138 | 6.06311649 | 3.04E-06 | 2.41E-06 |
| rs374768224 | 2 | 52507602 | LOC730100 | C | T | 0.00035833 | 28.310982 | 6.0650133 | 3.04E-06 | 2.41E-06 |
| rs376654178 | 2 | 52955934 | MIR4431(dist=26181),ASB3(dist=941183) | T | C | 0.00035845 | 28.3012281 | 6.06307159 | 3.04E-06 | 2.41E-06 |
| rs371862794 | 2 | 52908691 | LOC730100(dist=273636),MIR4431(dist=20969) | G | A | 0.00035702 | 28.3013398 | 6.06311181 | 3.04E-06 | 2.41E-06 |
| rs116726745 | 17 | 30045836 | MIR365B(dist=143296),COPRS(dist=133048) | G | A | 0.00036736 | 28.2906758 | 6.06086157 | 3.04E-06 | 2.41E-06 |
| rs115761220 | 2 | 52544377 | LOC730100 | T | A | 0.00035667 | 28.3011757 | 6.06311932 | 3.05E-06 | 2.41E-06 |
| rs115791435 | 2 | 52544380 | LOC730100 | C | T | 0.00035667 | 28.3011757 | 6.06311932 | 3.05E-06 | 2.41E-06 |
| rs115553765 | 2 | 52545698 | LOC730100 | C | A | 0.00035667 | 28.3011757 | 6.06311932 | 3.05E-06 | 2.41E-06 |
| rs373312188 | 2 | 52916201 | LOC730100(dist=281146),MIR4431(dist=13459) | A | G | 0.00035691 | 28.3010552 | 6.06311385 | 3.05E-06 | 2.41E-06 |
| rs1843038 | 2 | 52959633 | MIR4431(dist=29880),ASB3(dist=937484) | C | A | 0.00035857 | 28.2997373 | 6.06306467 | 3.05E-06 | 2.41E-06 |
| rs149851742 | 2 | 53008138 | MIR4431(dist=78385),ASB3(dist=888979) | C | T | 0.00035691 | 28.2998192 | 6.06309504 | 3.05E-06 | 2.41E-06 |
| rs4386361 | 2 | 53008780 | MIR4431(dist=79027),ASB3(dist=888337) | A | G | 0.00035691 | 28.2998192 | 6.06309504 | 3.05E-06 | 2.41E-06 |
| rs576662858 | 2 | 52714108 | LOC730100(dist=79053),MIR4431(dist=215552) | A | G | 0.00035667 | 28.2999507 | 6.06312655 | 3.05E-06 | 2.41E-06 |
| rs539771008 | 2 | 53146868 | MIR4431(dist=217115),ASB3(dist=750249) | A | G | 0.00035679 | 28.2989084 | 6.0631233 | 3.05E-06 | 2.41E-06 |
| rs142898785 | 2 | 52999581 | MIR4431(dist=69828),ASB3(dist=897536) | C | T | 0.00035679 | 28.2983588 | 6.06311864 | 3.05E-06 | 2.41E-06 |
| rs11894007 | 2 | 52743925 | LOC730100(dist=108870),MIR4431(dist=185735) | A | G | 0.00035726 | 28.2981401 | 6.06311312 | 3.05E-06 | 2.41E-06 |
| rs377759156 | 2 | 52954092 | MIR4431(dist=24339),ASB3(dist=943025) | T | G | 0.00035667 | 28.2981068 | 6.06310914 | 3.05E-06 | 2.41E-06 |
| rs375787925 | 2 | 52895590 | LOC730100(dist=260535),MIR4431(dist=34070) | T | C | 0.00035667 | 28.2980323 | 6.06311619 | 3.05E-06 | 2.41E-06 |
| rs371346212 | 2 | 52897986 | LOC730100(dist=262931),MIR4431(dist=31674) | C | T | 0.00035667 | 28.2980323 | 6.06311619 | 3.05E-06 | 2.41E-06 |
| rs113739919 | 19 | 450931 | SHC2 | A | G | 0.00036261 | 28.2917578 | 6.061892 | 3.05E-06 | 2.41E-06 |
| rs375636911 | 2 | 52912902 | LOC730100(dist=277847),MIR4431(dist=16758) | G | T | 0.00035714 | 28.2970861 | 6.06311793 | 3.06E-06 | 2.41E-06 |
| rs371650770 | 2 | 53037950 | MIR4431(dist=108197),ASB3(dist=859167) | T | C | 0.00035667 | 28.2967937 | 6.06311064 | 3.06E-06 | 2.41E-06 |
| rs377442641 | 2 | 53038287 | MIR4431(dist=108534),ASB3(dist=858830) | A | C | 0.00035667 | 28.2967937 | 6.06311064 | 3.06E-06 | 2.41E-06 |
| rs370510072 | 2 | 53039220 | MIR4431(dist=109467),ASB3(dist=857897) | T | G | 0.00035667 | 28.2967937 | 6.06311064 | 3.06E-06 | 2.41E-06 |
| rs373725461 | 2 | 53039313 | MIR4431(dist=109560),ASB3(dist=857804) | C | T | 0.00035667 | 28.2967937 | 6.06311064 | 3.06E-06 | 2.41E-06 |
| rs576529791 | 2 | 53046781 | MIR4431(dist=117028),ASB3(dist=850336) | T | C | 0.00035667 | 28.2967937 | 6.06311064 | 3.06E-06 | 2.41E-06 |
| rs374302389 | 2 | 53085983 | MIR4431(dist=156230),ASB3(dist=811134) | C | T | 0.00035809 | 28.2967224 | 6.06313206 | 3.06E-06 | 2.41E-06 |
| rs375110574 | 2 | 53112450 | MIR4431(dist=182697),ASB3(dist=784667) | A | C | 0.00035667 | 28.296116 | 6.06311681 | 3.06E-06 | 2.41E-06 |
| rs554319096 | 2 | 52835517 | LOC730100(dist=200462),MIR4431(dist=94143) | T | A | 0.00035691 | 28.2954723 | 6.06309909 | 3.06E-06 | 2.41E-06 |

| | | | | | | | | | | |
|---|---|---|---|---|---|---|---|---|---|---|
| rs116802691 | 2 | 52985351 | MIR4431(dist=55598),ASB3(dist=911766) | A | G | 0.00035667 | 28.2955383 | 6.06311397 | 3.06E-06 | 2.41E-06 |
| rs375240447 | 19 | 471395 | ODF3L2 | G | A | 0.00035738 | 28.2956326 | 6.06317108 | 3.06E-06 | 2.41E-06 |
| rs748926027 | 9 | 120716095 | TLR4(dist=236326),BRINP1(dist=1212813) | C | A | 0.00035512 | 28.408972 | 6.08747091 | 3.06E-06 | 2.41E-06 |
| rs546769066 | 2 | 52735604 | LOC730100(dist=100549),MIR4431(dist=194056) | T | C | 0.00035655 | 28.2953359 | 6.06312097 | 3.06E-06 | 2.41E-06 |
| rs10187972 | 2 | 52736860 | LOC730100(dist=101805),MIR4431(dist=192800) | C | T | 0.00035655 | 28.2953359 | 6.06312097 | 3.06E-06 | 2.41E-06 |
| rs10209599 | 2 | 52739102 | LOC730100(dist=104047),MIR4431(dist=190558) | T | C | 0.00035655 | 28.2953359 | 6.06312097 | 3.06E-06 | 2.41E-06 |
| rs563784859 | 2 | 52739868 | LOC730100(dist=104813),MIR4431(dist=189792) | C | T | 0.00035655 | 28.2953359 | 6.06312097 | 3.06E-06 | 2.41E-06 |
| rs546411610 | 2 | 52740102 | LOC730100(dist=105047),MIR4431(dist=189558) | T | C | 0.00035655 | 28.2953359 | 6.06312097 | 3.06E-06 | 2.41E-06 |
| rs115477931 | 2 | 52740286 | LOC730100(dist=105231),MIR4431(dist=189374) | G | A | 0.00035655 | 28.2953359 | 6.06312097 | 3.06E-06 | 2.41E-06 |
| rs75988214 | 2 | 52740556 | LOC730100(dist=105501),MIR4431(dist=189104) | G | A | 0.00035655 | 28.2953359 | 6.06312097 | 3.06E-06 | 2.41E-06 |
| rs550751645 | 2 | 52742082 | LOC730100(dist=107027),MIR4431(dist=187578) | C | T | 0.00035655 | 28.2953359 | 6.06312097 | 3.06E-06 | 2.41E-06 |
| rs569129441 | 2 | 52742087 | LOC730100(dist=107032),MIR4431(dist=187573) | C | T | 0.00035655 | 28.2953359 | 6.06312097 | 3.06E-06 | 2.41E-06 |
| rs539792866 | 2 | 52742434 | LOC730100(dist=107379),MIR4431(dist=187226) | A | G | 0.00035655 | 28.2953359 | 6.06312097 | 3.06E-06 | 2.41E-06 |
| rs570893929 | 2 | 52743488 | LOC730100(dist=108433),MIR4431(dist=186172) | G | C | 0.00035655 | 28.2953359 | 6.06312097 | 3.06E-06 | 2.41E-06 |
| rs546114948 | 2 | 52748536 | LOC730100(dist=113481),MIR4431(dist=181124) | C | T | 0.00035655 | 28.2953359 | 6.06312097 | 3.06E-06 | 2.41E-06 |
| rs527569542 | 2 | 52788118 | LOC730100(dist=153063),MIR4431(dist=141542) | A | G | 0.00035655 | 28.2953359 | 6.06312097 | 3.06E-06 | 2.41E-06 |
| rs529221520 | 2 | 52790307 | LOC730100(dist=155252),MIR4431(dist=139353) | T | C | 0.00035655 | 28.2953359 | 6.06312097 | 3.06E-06 | 2.41E-06 |
| rs550562911 | 2 | 52791266 | LOC730100(dist=156211),MIR4431(dist=138394) | C | T | 0.00035655 | 28.2953359 | 6.06312097 | 3.06E-06 | 2.41E-06 |
| rs568967913 | 2 | 52791270 | LOC730100(dist=156215),MIR4431(dist=138390) | C | A | 0.00035655 | 28.2953359 | 6.06312097 | 3.06E-06 | 2.41E-06 |
| rs537801577 | 2 | 52796544 | LOC730100(dist=161489),MIR4431(dist=133116) | C | T | 0.00035655 | 28.2953359 | 6.06312097 | 3.06E-06 | 2.41E-06 |
| rs551819697 | 2 | 52813175 | LOC730100(dist=178120),MIR4431(dist=116485) | T | A | 0.00035655 | 28.2953359 | 6.06312097 | 3.06E-06 | 2.41E-06 |
| rs538587569 | 2 | 52816899 | LOC730100(dist=181844),MIR4431(dist=112761) | C | T | 0.00035655 | 28.2953359 | 6.06312097 | 3.06E-06 | 2.41E-06 |
| rs572048758 | 2 | 52821697 | LOC730100(dist=186642),MIR4431(dist=107963) | A | C | 0.00035655 | 28.2953359 | 6.06312097 | 3.06E-06 | 2.41E-06 |
| rs554504698 | 2 | 52824896 | LOC730100(dist=189841),MIR4431(dist=104764) | G | A | 0.00035655 | 28.2953359 | 6.06312097 | 3.06E-06 | 2.41E-06 |
| rs545079054 | 2 | 52827210 | LOC730100(dist=192155),MIR4431(dist=102450) | T | C | 0.00035655 | 28.2953359 | 6.06312097 | 3.06E-06 | 2.41E-06 |
| rs532027353 | 2 | 52828580 | LOC730100(dist=193525),MIR4431(dist=101080) | C | T | 0.00035655 | 28.2953359 | 6.06312097 | 3.06E-06 | 2.41E-06 |
| rs530399323 | 2 | 52883641 | LOC730100(dist=248586),MIR4431(dist=46019) | G | C | 0.00035655 | 28.2953359 | 6.06312097 | 3.06E-06 | 2.41E-06 |
| rs113715155 | 2 | 52887694 | LOC730100(dist=252639),MIR4431(dist=41966) | C | A | 0.00035655 | 28.2953359 | 6.06312097 | 3.06E-06 | 2.41E-06 |
| rs372713390 | 2 | 52891535 | LOC730100(dist=256480),MIR4431(dist=38125) | A | C | 0.00035655 | 28.2953359 | 6.06312097 | 3.06E-06 | 2.41E-06 |
| rs373211730 | 2 | 52902479 | LOC730100(dist=267424),MIR4431(dist=27181) | A | G | 0.00035655 | 28.2953359 | 6.06312097 | 3.06E-06 | 2.41E-06 |
| rs375452509 | 2 | 52906030 | LOC730100(dist=270975),MIR4431(dist=23630) | A | C | 0.00035655 | 28.2953359 | 6.06312097 | 3.06E-06 | 2.41E-06 |
| rs368001174 | 2 | 52907612 | LOC730100(dist=272557),MIR4431(dist=22048) | G | A | 0.00035655 | 28.2953359 | 6.06312097 | 3.06E-06 | 2.41E-06 |
| rs374199178 | 2 | 52943891 | MIR4431(dist=14138),ASB3(dist=953226) | T | A | 0.00035655 | 28.2953359 | 6.06312097 | 3.06E-06 | 2.41E-06 |
| rs376783662 | 2 | 52945859 | MIR4431(dist=16106),ASB3(dist=951258) | A | C | 0.00035655 | 28.2953359 | 6.06312097 | 3.06E-06 | 2.41E-06 |
| rs367548755 | 2 | 52945977 | MIR4431(dist=16224),ASB3(dist=951140) | A | G | 0.00035655 | 28.2953359 | 6.06312097 | 3.06E-06 | 2.41E-06 |
| rs567053198 | 2 | 52946641 | MIR4431(dist=16888),ASB3(dist=950476) | T | A | 0.00035655 | 28.2953359 | 6.06312097 | 3.06E-06 | 2.41E-06 |
| rs376851429 | 2 | 52946746 | MIR4431(dist=16993),ASB3(dist=950371) | T | C | 0.00035655 | 28.2953359 | 6.06312097 | 3.06E-06 | 2.41E-06 |
| rs369745827 | 2 | 52949008 | MIR4431(dist=19255),ASB3(dist=948109) | G | C | 0.00035655 | 28.2953359 | 6.06312097 | 3.06E-06 | 2.41E-06 |
| rs373229057 | 2 | 52949073 | MIR4431(dist=19320),ASB3(dist=948044) | A | T | 0.00035655 | 28.2953359 | 6.06312097 | 3.06E-06 | 2.41E-06 |
| rs556310119 | 2 | 52949752 | MIR4431(dist=19999),ASB3(dist=947365) | A | G | 0.00035655 | 28.2953359 | 6.06312097 | 3.06E-06 | 2.41E-06 |
| rs377608108 | 2 | 52952455 | MIR4431(dist=22702),ASB3(dist=944662) | A | G | 0.00035655 | 28.2953359 | 6.06312097 | 3.06E-06 | 2.41E-06 |
| rs371103615 | 2 | 52952570 | MIR4431(dist=22817),ASB3(dist=944547) | G | T | 0.00035655 | 28.2953359 | 6.06312097 | 3.06E-06 | 2.41E-06 |
| rs371111972 | 2 | 52970008 | MIR4431(dist=40255),ASB3(dist=927109) | T | C | 0.00035655 | 28.2953359 | 6.06312097 | 3.06E-06 | 2.41E-06 |
| rs369173152 | 2 | 52974817 | MIR4431(dist=45064),ASB3(dist=922300) | G | A | 0.00035655 | 28.2953359 | 6.06312097 | 3.06E-06 | 2.41E-06 |
| rs374048736 | 2 | 52975352 | MIR4431(dist=45599),ASB3(dist=921765) | T | C | 0.00035655 | 28.2953359 | 6.06312097 | 3.06E-06 | 2.41E-06 |
| rs376844753 | 2 | 52980403 | MIR4431(dist=50650),ASB3(dist=916714) | C | T | 0.00035655 | 28.2953359 | 6.06312097 | 3.06E-06 | 2.41E-06 |
| rs372848560 | 2 | 52980866 | MIR4431(dist=51113),ASB3(dist=916251) | A | C | 0.00035655 | 28.2953359 | 6.06312097 | 3.06E-06 | 2.41E-06 |
| rs185725044 | 2 | 52986949 | MIR4431(dist=57196),ASB3(dist=910168) | C | A | 0.00035655 | 28.2953359 | 6.06312097 | 3.06E-06 | 2.41E-06 |
| rs373907362 | 2 | 52987346 | MIR4431(dist=57593),ASB3(dist=909771) | C | T | 0.00035655 | 28.2953359 | 6.06312097 | 3.06E-06 | 2.41E-06 |
| rs370340869 | 2 | 52988086 | MIR4431(dist=58333),ASB3(dist=909031) | A | G | 0.00035655 | 28.2953359 | 6.06312097 | 3.06E-06 | 2.41E-06 |
| rs377613717 | 2 | 52989589 | MIR4431(dist=59836),ASB3(dist=907528) | T | C | 0.00035655 | 28.2953359 | 6.06312097 | 3.06E-06 | 2.41E-06 |
| rs371746508 | 2 | 52993493 | MIR4431(dist=63740),ASB3(dist=903624) | G | A | 0.00035655 | 28.2953359 | 6.06312097 | 3.06E-06 | 2.41E-06 |

| | | | | | | | | | | |
|---|---|---|---|---|---|---|---|---|---|---|
| rs4263147 | 2 | 52996436 | MIR4431(dist=66683),ASB3(dist=900681) | G | A | 0.00035655 | 28.2953359 | 6.06312097 | 3.06E-06 | 2.41E-06 |
| rs374957640 | 2 | 52997217 | MIR4431(dist=67464),ASB3(dist=899900) | A | G | 0.00035655 | 28.2953359 | 6.06312097 | 3.06E-06 | 2.41E-06 |
| rs7584468 | 2 | 52998684 | MIR4431(dist=68931),ASB3(dist=898433) | A | G | 0.00035655 | 28.2953359 | 6.06312097 | 3.06E-06 | 2.41E-06 |
| rs370552692 | 2 | 52999439 | MIR4431(dist=69686),ASB3(dist=897678) | T | G | 0.00035655 | 28.2953359 | 6.06312097 | 3.06E-06 | 2.41E-06 |
| rs578038220 | 2 | 53000477 | MIR4431(dist=70724),ASB3(dist=896640) | G | A | 0.00035655 | 28.2953359 | 6.06312097 | 3.06E-06 | 2.41E-06 |
| rs375008586 | 2 | 53003027 | MIR4431(dist=73274),ASB3(dist=894090) | T | C | 0.00035655 | 28.2953359 | 6.06312097 | 3.06E-06 | 2.41E-06 |
| rs376418644 | 2 | 53009954 | MIR4431(dist=80201),ASB3(dist=887163) | G | C | 0.00035655 | 28.2953359 | 6.06312097 | 3.06E-06 | 2.41E-06 |
| rs532066277 | 2 | 53012669 | MIR4431(dist=82916),ASB3(dist=884448) | G | T | 0.00035655 | 28.2953359 | 6.06312097 | 3.06E-06 | 2.41E-06 |
| rs535999858 | 2 | 53012670 | MIR4431(dist=82917),ASB3(dist=884447) | C | T | 0.00035655 | 28.2953359 | 6.06312097 | 3.06E-06 | 2.41E-06 |
| rs372024582 | 2 | 53015305 | MIR4431(dist=85552),ASB3(dist=881812) | T | C | 0.00035655 | 28.2953359 | 6.06312097 | 3.06E-06 | 2.41E-06 |
| rs375216538 | 2 | 53015379 | MIR4431(dist=85626),ASB3(dist=881738) | A | C | 0.00035655 | 28.2953359 | 6.06312097 | 3.06E-06 | 2.41E-06 |
| rs376159637 | 2 | 53016909 | MIR4431(dist=87156),ASB3(dist=880208) | A | G | 0.00035655 | 28.2953359 | 6.06312097 | 3.06E-06 | 2.41E-06 |
| rs374125682 | 2 | 53018752 | MIR4431(dist=88999),ASB3(dist=878365) | C | G | 0.00035655 | 28.2953359 | 6.06312097 | 3.06E-06 | 2.41E-06 |
| rs375720110 | 2 | 53018789 | MIR4431(dist=89036),ASB3(dist=878328) | C | T | 0.00035655 | 28.2953359 | 6.06312097 | 3.06E-06 | 2.41E-06 |
| rs146387452 | 2 | 53019922 | MIR4431(dist=90169),ASB3(dist=877195) | C | T | 0.00035655 | 28.2953359 | 6.06312097 | 3.06E-06 | 2.41E-06 |
| rs371168552 | 2 | 53022057 | MIR4431(dist=92304),ASB3(dist=875060) | C | T | 0.00035655 | 28.2953359 | 6.06312097 | 3.06E-06 | 2.41E-06 |
| rs376614829 | 2 | 53022790 | MIR4431(dist=93037),ASB3(dist=874327) | G | A | 0.00035655 | 28.2953359 | 6.06312097 | 3.06E-06 | 2.41E-06 |
| rs375027288 | 2 | 53023367 | MIR4431(dist=93614),ASB3(dist=873750) | T | C | 0.00035655 | 28.2953359 | 6.06312097 | 3.06E-06 | 2.41E-06 |
| rs372448654 | 2 | 53023769 | MIR4431(dist=94016),ASB3(dist=873348) | C | T | 0.00035655 | 28.2953359 | 6.06312097 | 3.06E-06 | 2.41E-06 |
| rs368385681 | 2 | 53024808 | MIR4431(dist=95055),ASB3(dist=872309) | T | G | 0.00035655 | 28.2953359 | 6.06312097 | 3.06E-06 | 2.41E-06 |
| rs374720584 | 2 | 53025662 | MIR4431(dist=95909),ASB3(dist=871455) | A | G | 0.00035655 | 28.2953359 | 6.06312097 | 3.06E-06 | 2.41E-06 |
| rs368579989 | 2 | 53027201 | MIR4431(dist=97448),ASB3(dist=869916) | T | C | 0.00035655 | 28.2953359 | 6.06312097 | 3.06E-06 | 2.41E-06 |
| rs376974781 | 2 | 53045609 | MIR4431(dist=115856),ASB3(dist=851508) | A | G | 0.00035655 | 28.2953359 | 6.06312097 | 3.06E-06 | 2.41E-06 |
| rs1451461 | 2 | 53045987 | MIR4431(dist=116234),ASB3(dist=851130) | T | C | 0.00035655 | 28.2953359 | 6.06312097 | 3.06E-06 | 2.41E-06 |
| rs377305018 | 2 | 53049078 | MIR4431(dist=119325),ASB3(dist=848039) | G | A | 0.00035655 | 28.2953359 | 6.06312097 | 3.06E-06 | 2.41E-06 |
| rs372780258 | 2 | 53051083 | MIR4431(dist=121330),ASB3(dist=846034) | G | C | 0.00035655 | 28.2953359 | 6.06312097 | 3.06E-06 | 2.41E-06 |
| rs374393210 | 2 | 53052556 | MIR4431(dist=122803),ASB3(dist=844561) | C | A | 0.00035655 | 28.2953359 | 6.06312097 | 3.06E-06 | 2.41E-06 |
| rs572221454 | 2 | 53053088 | MIR4431(dist=123335),ASB3(dist=844029) | A | G | 0.00035655 | 28.2953359 | 6.06312097 | 3.06E-06 | 2.41E-06 |
| rs374596773 | 2 | 53054512 | MIR4431(dist=124759),ASB3(dist=842605) | T | A | 0.00035655 | 28.2953359 | 6.06312097 | 3.06E-06 | 2.41E-06 |
| rs370658242 | 2 | 53057573 | MIR4431(dist=127820),ASB3(dist=839544) | T | C | 0.00035655 | 28.2953359 | 6.06312097 | 3.06E-06 | 2.41E-06 |
| rs368423498 | 2 | 53067509 | MIR4431(dist=137756),ASB3(dist=829608) | T | C | 0.00035655 | 28.2953359 | 6.06312097 | 3.06E-06 | 2.41E-06 |
| rs370673490 | 2 | 53069385 | MIR4431(dist=139632),ASB3(dist=827732) | A | T | 0.00035655 | 28.2953359 | 6.06312097 | 3.06E-06 | 2.41E-06 |
| rs372673144 | 2 | 53070194 | MIR4431(dist=140441),ASB3(dist=826923) | C | T | 0.00035655 | 28.2953359 | 6.06312097 | 3.06E-06 | 2.41E-06 |
| rs368886465 | 2 | 53070352 | MIR4431(dist=140599),ASB3(dist=826765) | A | G | 0.00035655 | 28.2953359 | 6.06312097 | 3.06E-06 | 2.41E-06 |
| rs368684152 | 2 | 53074254 | MIR4431(dist=144501),ASB3(dist=822863) | A | G | 0.00035655 | 28.2953359 | 6.06312097 | 3.06E-06 | 2.41E-06 |
| rs367793883 | 2 | 53074634 | MIR4431(dist=144881),ASB3(dist=822483) | A | C | 0.00035655 | 28.2953359 | 6.06312097 | 3.06E-06 | 2.41E-06 |
| rs2122834 | 2 | 53076508 | MIR4431(dist=146755),ASB3(dist=820609) | G | C | 0.00035655 | 28.2953359 | 6.06312097 | 3.06E-06 | 2.41E-06 |
| rs374828848 | 2 | 53078963 | MIR4431(dist=149210),ASB3(dist=818154) | G | A | 0.00035655 | 28.2953359 | 6.06312097 | 3.06E-06 | 2.41E-06 |
| rs372283813 | 2 | 53079813 | MIR4431(dist=150060),ASB3(dist=817304) | C | T | 0.00035655 | 28.2953359 | 6.06312097 | 3.06E-06 | 2.41E-06 |
| rs370070279 | 2 | 53083856 | MIR4431(dist=154103),ASB3(dist=813261) | A | G | 0.00035655 | 28.2953359 | 6.06312097 | 3.06E-06 | 2.41E-06 |
| rs187031480 | 2 | 53084831 | MIR4431(dist=155078),ASB3(dist=812286) | C | G | 0.00035655 | 28.2953359 | 6.06312097 | 3.06E-06 | 2.41E-06 |
| rs374699037 | 2 | 53085539 | MIR4431(dist=155786),ASB3(dist=811578) | A | T | 0.00035655 | 28.2953359 | 6.06312097 | 3.06E-06 | 2.41E-06 |
| rs371759740 | 2 | 53086199 | MIR4431(dist=156446),ASB3(dist=810918) | G | A | 0.00035655 | 28.2953359 | 6.06312097 | 3.06E-06 | 2.41E-06 |
| rs367818531 | 2 | 53086329 | MIR4431(dist=156576),ASB3(dist=810788) | T | A | 0.00035655 | 28.2953359 | 6.06312097 | 3.06E-06 | 2.41E-06 |
| rs184160873 | 2 | 53090179 | MIR4431(dist=160426),ASB3(dist=806938) | C | T | 0.00035655 | 28.2953359 | 6.06312097 | 3.06E-06 | 2.41E-06 |
| rs138517075 | 2 | 53090960 | MIR4431(dist=161207),ASB3(dist=806157) | T | G | 0.00035655 | 28.2953359 | 6.06312097 | 3.06E-06 | 2.41E-06 |
| rs376352658 | 2 | 53092151 | MIR4431(dist=162398),ASB3(dist=804966) | G | C | 0.00035655 | 28.2953359 | 6.06312097 | 3.06E-06 | 2.41E-06 |
| rs375953088 | 2 | 53093169 | MIR4431(dist=163416),ASB3(dist=803948) | T | G | 0.00035655 | 28.2953359 | 6.06312097 | 3.06E-06 | 2.41E-06 |
| rs368040583 | 2 | 53093425 | MIR4431(dist=163672),ASB3(dist=803692) | C | T | 0.00035655 | 28.2953359 | 6.06312097 | 3.06E-06 | 2.41E-06 |
| rs190919296 | 2 | 53093463 | MIR4431(dist=163710),ASB3(dist=803654) | A | G | 0.00035655 | 28.2953359 | 6.06312097 | 3.06E-06 | 2.41E-06 |
| rs182625218 | 2 | 53093767 | MIR4431(dist=164014),ASB3(dist=803350) | C | T | 0.00035655 | 28.2953359 | 6.06312097 | 3.06E-06 | 2.41E-06 |
| rs374272665 | 2 | 53095847 | MIR4431(dist=166094),ASB3(dist=801270) | A | G | 0.00035655 | 28.2953359 | 6.06312097 | 3.06E-06 | 2.41E-06 |

| | | | | | | | | | | |
|---|---|---|---|---|---|---|---|---|---|---|
| rs371176997 | 2 | 53096275 | MIR4431(dist=166522),ASB3(dist=800842) | C | A | 0.00035655 | 28.2953359 | 6.06312097 | 3.06E-06 | 2.41E-06 |
| rs375647791 | 2 | 53097666 | MIR4431(dist=167913),ASB3(dist=799451) | T | A | 0.00035655 | 28.2953359 | 6.06312097 | 3.06E-06 | 2.41E-06 |
| rs367901292 | 2 | 53097825 | MIR4431(dist=168072),ASB3(dist=799292) | G | C | 0.00035655 | 28.2953359 | 6.06312097 | 3.06E-06 | 2.41E-06 |
| rs140739207 | 2 | 53098046 | MIR4431(dist=168293),ASB3(dist=799071) | C | T | 0.00035655 | 28.2953359 | 6.06312097 | 3.06E-06 | 2.41E-06 |
| rs1376566 | 2 | 53098542 | MIR4431(dist=168789),ASB3(dist=798575) | A | C | 0.00035655 | 28.2953359 | 6.06312097 | 3.06E-06 | 2.41E-06 |
| rs552594509 | 2 | 53099558 | MIR4431(dist=169805),ASB3(dist=797559) | A | T | 0.00035655 | 28.2953359 | 6.06312097 | 3.06E-06 | 2.41E-06 |
| rs374268300 | 2 | 53100764 | MIR4431(dist=171011),ASB3(dist=796353) | T | C | 0.00035655 | 28.2953359 | 6.06312097 | 3.06E-06 | 2.41E-06 |
| rs377035611 | 2 | 53101538 | MIR4431(dist=171785),ASB3(dist=795579) | T | C | 0.00035655 | 28.2953359 | 6.06312097 | 3.06E-06 | 2.41E-06 |
| rs369835291 | 2 | 53102987 | MIR4431(dist=173234),ASB3(dist=794130) | G | C | 0.00035655 | 28.2953359 | 6.06312097 | 3.06E-06 | 2.41E-06 |
| rs1868906 | 2 | 53105047 | MIR4431(dist=175294),ASB3(dist=792070) | T | C | 0.00035655 | 28.2953359 | 6.06312097 | 3.06E-06 | 2.41E-06 |
| rs368428877 | 2 | 53105628 | MIR4431(dist=175875),ASB3(dist=791489) | G | A | 0.00035655 | 28.2953359 | 6.06312097 | 3.06E-06 | 2.41E-06 |
| rs376861736 | 2 | 53107240 | MIR4431(dist=177487),ASB3(dist=789877) | C | G | 0.00035655 | 28.2953359 | 6.06312097 | 3.06E-06 | 2.41E-06 |
| rs369494773 | 2 | 53107659 | MIR4431(dist=177906),ASB3(dist=789458) | C | G | 0.00035655 | 28.2953359 | 6.06312097 | 3.06E-06 | 2.41E-06 |
| rs377436118 | 2 | 53107705 | MIR4431(dist=177952),ASB3(dist=789412) | C | T | 0.00035655 | 28.2953359 | 6.06312097 | 3.06E-06 | 2.41E-06 |
| rs1864533 | 2 | 53109379 | MIR4431(dist=179626),ASB3(dist=787738) | A | T | 0.00035655 | 28.2953359 | 6.06312097 | 3.06E-06 | 2.41E-06 |
| rs369471378 | 2 | 53112597 | MIR4431(dist=182844),ASB3(dist=784520) | G | A | 0.00035655 | 28.2953359 | 6.06312097 | 3.06E-06 | 2.41E-06 |
| rs370668232 | 2 | 53114029 | MIR4431(dist=184276),ASB3(dist=783088) | A | G | 0.00035655 | 28.2953359 | 6.06312097 | 3.06E-06 | 2.41E-06 |
| rs374881775 | 2 | 53119483 | MIR4431(dist=189730),ASB3(dist=777634) | C | T | 0.00035655 | 28.2953359 | 6.06312097 | 3.06E-06 | 2.41E-06 |
| rs557719687 | 2 | 53147235 | MIR4431(dist=217482),ASB3(dist=749882) | C | T | 0.00035655 | 28.2953359 | 6.06312097 | 3.06E-06 | 2.41E-06 |
| rs141882698 | 11 | 116601338 | LOC101929011(dist=72369),BUD13(dist=17548) | A | C | 0.00035655 | 28.2953359 | 6.06312097 | 3.06E-06 | 2.41E-06 |
| rs374175094 | 12 | 60058530 | SLC16A7 | C | T | 0.00035655 | 28.2953359 | 6.06312097 | 3.06E-06 | 2.41E-06 |
| rs150258437 | 12 | 60087460 | SLC16A7 | T | C | 0.00035655 | 28.2953359 | 6.06312097 | 3.06E-06 | 2.41E-06 |
| rs138452335 | 12 | 60091008 | SLC16A7 | G | A | 0.00035655 | 28.2953359 | 6.06312097 | 3.06E-06 | 2.41E-06 |
| rs151027047 | 12 | 60108470 | SLC16A7 | C | T | 0.00035655 | 28.2953359 | 6.06312097 | 3.06E-06 | 2.41E-06 |
| rs11173127 | 12 | 60133321 | SLC16A7 | G | A | 0.00035655 | 28.2953359 | 6.06312097 | 3.06E-06 | 2.41E-06 |
| rs767930336 | 12 | 60140721 | SLC16A7 | C | T | 0.00035655 | 28.2953359 | 6.06312097 | 3.06E-06 | 2.41E-06 |
| rs12303722 | 12 | 60144427 | SLC16A7 | G | A | 0.00035655 | 28.2953359 | 6.06312097 | 3.06E-06 | 2.41E-06 |
| rs189827842 | 12 | 60145004 | SLC16A7 | A | G | 0.00035655 | 28.2953359 | 6.06312097 | 3.06E-06 | 2.41E-06 |
| rs74096452 | 12 | 60164585 | SLC16A7 | A | G | 0.00035655 | 28.2953359 | 6.06312097 | 3.06E-06 | 2.41E-06 |
| rs74096453 | 12 | 60164673 | SLC16A7 | C | T | 0.00035655 | 28.2953359 | 6.06312097 | 3.06E-06 | 2.41E-06 |
| rs13378061 | 12 | 60169806 | SLC16A7 | T | A | 0.00035655 | 28.2953359 | 6.06312097 | 3.06E-06 | 2.41E-06 |
| rs7316320 | 12 | 60171528 | SLC16A7 | T | C | 0.00035655 | 28.2953359 | 6.06312097 | 3.06E-06 | 2.41E-06 |
| rs11173140 | 12 | 60173052 | SLC16A7 | A | T | 0.00035655 | 28.2953359 | 6.06312097 | 3.06E-06 | 2.41E-06 |
| rs11173141 | 12 | 60173806 | SLC16A7(NM_001270623:c.*3460>0,NM_001270622:c.*34 | A | G | 0.00035655 | 28.2953359 | 6.06312097 | 3.06E-06 | 2.41E-06 |
| rs11173142 | 12 | 60173878 | SLC16A7(NM_001270623:c.*4180>0,NM_001270622:c.*41 | A | C | 0.00035655 | 28.2953359 | 6.06312097 | 3.06E-06 | 2.41E-06 |
| rs149148185 | 12 | 60174087 | SLC16A7(NM_001270623:c.*6270>0,NM_001270622:c.*62 | A | T | 0.00035655 | 28.2953359 | 6.06312097 | 3.06E-06 | 2.41E-06 |
| rs11173144 | 12 | 60176349 | SLC16A7(NM_001270623:c.*28890>0,NM_001270622:c.*2 | A | T | 0.00035655 | 28.2953359 | 6.06312097 | 3.06E-06 | 2.41E-06 |
| rs11173145 | 12 | 60176474 | SLC16A7(NM_001270623:c.*30140>0,NM_001270622:c.*3 | C | T | 0.00035655 | 28.2953359 | 6.06312097 | 3.06E-06 | 2.41E-06 |
| rs11173146 | 12 | 60176576 | SLC16A7(NM_001270623:c.*31160>0,NM_001270622:c.*3 | C | A | 0.00035655 | 28.2953359 | 6.06312097 | 3.06E-06 | 2.41E-06 |
| rs12316816 | 12 | 60177483 | SLC16A7(NM_001270623:c.*40230>0,NM_001270622:c.*4 | C | A | 0.00035655 | 28.2953359 | 6.06312097 | 3.06E-06 | 2.41E-06 |
| rs12303301 | 12 | 60177494 | SLC16A7(NM_001270623:c.*40340>0,NM_001270622:c.*4 | A | G | 0.00035655 | 28.2953359 | 6.06312097 | 3.06E-06 | 2.41E-06 |
| rs7307407 | 12 | 60178234 | SLC16A7(NM_001270623:c.*47740>0,NM_001270622:c.*4 | G | C | 0.00035655 | 28.2953359 | 6.06312097 | 3.06E-06 | 2.41E-06 |
| rs7307832 | 12 | 60178589 | SLC16A7(NM_001270623:c.*51290>0,NM_001270622:c.*5 | G | C | 0.00035655 | 28.2953359 | 6.06312097 | 3.06E-06 | 2.41E-06 |
| rs7310780 | 12 | 60178801 | SLC16A7(NM_001270623:c.*53410>0,NM_001270622:c.*5 | A | G | 0.00035655 | 28.2953359 | 6.06312097 | 3.06E-06 | 2.41E-06 |
| rs7311237 | 12 | 60178910 | SLC16A7(NM_001270623:c.*54500>0,NM_001270622:c.*5 | G | C | 0.00035655 | 28.2953359 | 6.06312097 | 3.06E-06 | 2.41E-06 |
| rs28701946 | 12 | 60179814 | SLC16A7(NM_001270623:c.*63540>0,NM_001270622:c.*6 | G | T | 0.00035655 | 28.2953359 | 6.06312097 | 3.06E-06 | 2.41E-06 |
| rs28534781 | 12 | 60179847 | SLC16A7(NM_001270623:c.*63870>0,NM_001270622:c.*6 | G | A | 0.00035655 | 28.2953359 | 6.06312097 | 3.06E-06 | 2.41E-06 |
| rs12304208 | 12 | 60180842 | SLC16A7(NM_001270623:c.*73820>0,NM_001270622:c.*7 | T | C | 0.00035655 | 28.2953359 | 6.06312097 | 3.06E-06 | 2.41E-06 |
| rs12310755 | 12 | 60181280 | SLC16A7(NM_001270623:c.*78200>0,NM_001270622:c.*7 | A | G | 0.00035655 | 28.2953359 | 6.06312097 | 3.06E-06 | 2.41E-06 |
| rs12304515 | 12 | 60181337 | SLC16A7(NM_001270623:c.*78770>0,NM_001270622:c.*7 | T | A | 0.00035655 | 28.2953359 | 6.06312097 | 3.06E-06 | 2.41E-06 |
| rs12306387 | 12 | 60182244 | SLC16A7(NM_001270623:c.*87840>0,NM_001270622:c.*8 | T | C | 0.00035655 | 28.2953359 | 6.06312097 | 3.06E-06 | 2.41E-06 |
| rs74095498 | 12 | 60182681 | SLC16A7(NM_001270623:c.*92210>0,NM_001270622:c.*9 | A | C | 0.00035655 | 28.2953359 | 6.06312097 | 3.06E-06 | 2.41E-06 |

| | | | | | | | | | | |
|---|---|---|---|---|---|---|---|---|---|---|
| rs74095500 | 12 | 60182765 | SLC16A7(NM_001270623:c.*93050>0,NM_001270622:c.*9 | T | C | 0.00035655 | 28.2953359 | 6.06312097 | 3.06E-06 | 2.41E-06 |
| rs74095501 | 12 | 60182906 | SLC16A7(NM_001270623:c.*94460>0,NM_001270622:c.*9 | A | G | 0.00035655 | 28.2953359 | 6.06312097 | 3.06E-06 | 2.41E-06 |
| rs56121641 | 12 | 60183692 | SLC16A7(dist=57) | A | G | 0.00035655 | 28.2953359 | 6.06312097 | 3.06E-06 | 2.41E-06 |
| rs10161078 | 12 | 60185335 | SLC16A7(dist=1700),FAM19A2(dist=1916694) | C | T | 0.00035655 | 28.2953359 | 6.06312097 | 3.06E-06 | 2.41E-06 |
| rs10160899 | 12 | 60186828 | SLC16A7(dist=3193),FAM19A2(dist=1915201) | C | T | 0.00035655 | 28.2953359 | 6.06312097 | 3.06E-06 | 2.41E-06 |
| rs10161101 | 12 | 60187174 | SLC16A7(dist=3539),FAM19A2(dist=1914855) | T | G | 0.00035655 | 28.2953359 | 6.06312097 | 3.06E-06 | 2.41E-06 |
| rs12300901 | 12 | 60188479 | SLC16A7(dist=4844),FAM19A2(dist=1913550) | G | C | 0.00035655 | 28.2953359 | 6.06312097 | 3.06E-06 | 2.41E-06 |
| rs11173154 | 12 | 60189332 | SLC16A7(dist=5697),FAM19A2(dist=1912697) | G | A | 0.00035655 | 28.2953359 | 6.06312097 | 3.06E-06 | 2.41E-06 |
| rs11504057 | 12 | 60189610 | SLC16A7(dist=5975),FAM19A2(dist=1912419) | C | T | 0.00035655 | 28.2953359 | 6.06312097 | 3.06E-06 | 2.41E-06 |
| rs78279772 | 12 | 60192365 | SLC16A7(dist=8730),FAM19A2(dist=1909664) | A | G | 0.00035655 | 28.2953359 | 6.06312097 | 3.06E-06 | 2.41E-06 |
| rs12298827 | 12 | 60194678 | SLC16A7(dist=11043),FAM19A2(dist=1907351) | A | G | 0.00035655 | 28.2953359 | 6.06312097 | 3.06E-06 | 2.41E-06 |
| rs12296304 | 12 | 60196574 | SLC16A7(dist=12939),FAM19A2(dist=1905455) | T | C | 0.00035655 | 28.2953359 | 6.06312097 | 3.06E-06 | 2.41E-06 |
| rs12318590 | 12 | 60199283 | SLC16A7(dist=15648),FAM19A2(dist=1902746) | T | C | 0.00035655 | 28.2953359 | 6.06312097 | 3.06E-06 | 2.41E-06 |
| rs12318639 | 12 | 60199337 | SLC16A7(dist=15702),FAM19A2(dist=1902692) | T | C | 0.00035655 | 28.2953359 | 6.06312097 | 3.06E-06 | 2.41E-06 |
| rs12301821 | 12 | 60201300 | SLC16A7(dist=17665),FAM19A2(dist=1900729) | A | G | 0.00035655 | 28.2953359 | 6.06312097 | 3.06E-06 | 2.41E-06 |
| rs10437931 | 12 | 60201882 | SLC16A7(dist=18247),FAM19A2(dist=1900147) | G | A | 0.00035655 | 28.2953359 | 6.06312097 | 3.06E-06 | 2.41E-06 |
| rs11173161 | 12 | 60205072 | SLC16A7(dist=21437),FAM19A2(dist=1896957) | A | G | 0.00035655 | 28.2953359 | 6.06312097 | 3.06E-06 | 2.41E-06 |
| rs115609179 | 12 | 60222966 | SLC16A7(dist=39331),FAM19A2(dist=1879063) | C | G | 0.00035655 | 28.2953359 | 6.06312097 | 3.06E-06 | 2.41E-06 |
| rs74095777 | 12 | 60227712 | SLC16A7(dist=44077),FAM19A2(dist=1874317) | G | C | 0.00035655 | 28.2953359 | 6.06312097 | 3.06E-06 | 2.41E-06 |
| rs74095781 | 12 | 60229512 | SLC16A7(dist=45877),FAM19A2(dist=1872517) | C | T | 0.00035655 | 28.2953359 | 6.06312097 | 3.06E-06 | 2.41E-06 |
| rs74095794 | 12 | 60237301 | SLC16A7(dist=53666),FAM19A2(dist=1864728) | T | C | 0.00035655 | 28.2953359 | 6.06312097 | 3.06E-06 | 2.41E-06 |
| rs865864711 | 12 | 60645607 | SLC16A7(dist=461972),FAM19A2(dist=1456422) | T | C | 0.00035655 | 28.2953359 | 6.06312097 | 3.06E-06 | 2.41E-06 |
| rs73489637 | 19 | 452105 | SHC2 | T | C | 0.00035655 | 28.2953359 | 6.06312097 | 3.06E-06 | 2.41E-06 |
| rs544617045 | 2 | 52541999 | LOC730100 | G | A | 0.00035655 | 28.2953359 | 6.06312097 | 3.06E-06 | 2.41E-06 |
| rs115334491 | 2 | 52545366 | LOC730100 | A | G | 0.00035655 | 28.2953359 | 6.06312097 | 3.06E-06 | 2.41E-06 |
| rs527392879 | 2 | 52547080 | LOC730100 | C | T | 0.00035655 | 28.2953359 | 6.06312097 | 3.06E-06 | 2.41E-06 |
| rs372232719 | 2 | 52549212 | LOC730100 | T | C | 0.00035655 | 28.2953359 | 6.06312097 | 3.06E-06 | 2.41E-06 |
| rs566577469 | 2 | 52549491 | LOC730100 | G | T | 0.00035655 | 28.2953359 | 6.06312097 | 3.06E-06 | 2.41E-06 |
| rs555529050 | 2 | 52551773 | LOC730100 | G | A | 0.00035655 | 28.2953359 | 6.06312097 | 3.06E-06 | 2.41E-06 |
| rs377373388 | 2 | 52553502 | LOC730100 | C | T | 0.00035655 | 28.2953359 | 6.06312097 | 3.06E-06 | 2.41E-06 |
| rs544639429 | 2 | 52553534 | LOC730100 | A | T | 0.00035655 | 28.2953359 | 6.06312097 | 3.06E-06 | 2.41E-06 |
| rs368726778 | 2 | 52553734 | LOC730100 | T | C | 0.00035655 | 28.2953359 | 6.06312097 | 3.06E-06 | 2.41E-06 |
| rs377108751 | 2 | 52553792 | LOC730100 | C | A | 0.00035655 | 28.2953359 | 6.06312097 | 3.06E-06 | 2.41E-06 |
| rs370510488 | 2 | 52553823 | LOC730100 | G | A | 0.00035655 | 28.2953359 | 6.06312097 | 3.06E-06 | 2.41E-06 |
| rs376826721 | 2 | 52553993 | LOC730100 | G | A | 0.00035655 | 28.2953359 | 6.06312097 | 3.06E-06 | 2.41E-06 |
| rs368715703 | 2 | 52554493 | LOC730100 | G | A | 0.00035655 | 28.2953359 | 6.06312097 | 3.06E-06 | 2.41E-06 |
| rs571419511 | 2 | 52554607 | LOC730100 | A | T | 0.00035655 | 28.2953359 | 6.06312097 | 3.06E-06 | 2.41E-06 |
| rs368523060 | 2 | 52559306 | LOC730100 | A | G | 0.00035655 | 28.2953359 | 6.06312097 | 3.06E-06 | 2.41E-06 |
| rs571728366 | 2 | 52581588 | LOC730100 | C | A | 0.00035655 | 28.2953359 | 6.06312097 | 3.06E-06 | 2.41E-06 |
| rs539209253 | 2 | 52582267 | LOC730100 | A | G | 0.00035655 | 28.2953359 | 6.06312097 | 3.06E-06 | 2.41E-06 |
| rs540786989 | 2 | 52587032 | LOC730100 | G | A | 0.00035655 | 28.2953359 | 6.06312097 | 3.06E-06 | 2.41E-06 |
| rs375126523 | 2 | 52587641 | LOC730100 | G | A | 0.00035655 | 28.2953359 | 6.06312097 | 3.06E-06 | 2.41E-06 |
| rs561112087 | 2 | 52587714 | LOC730100 | T | A | 0.00035655 | 28.2953359 | 6.06312097 | 3.06E-06 | 2.41E-06 |
| rs116707515 | 2 | 52679675 | LOC730100(dist=44620),MIR4431(dist=249985) | T | G | 0.00035655 | 28.2953359 | 6.06312097 | 3.06E-06 | 2.41E-06 |
| rs531499393 | 2 | 52680402 | LOC730100(dist=45347),MIR4431(dist=249258) | A | G | 0.00035655 | 28.2953359 | 6.06312097 | 3.06E-06 | 2.41E-06 |
| rs538141757 | 2 | 52685905 | LOC730100(dist=50850),MIR4431(dist=243755) | A | G | 0.00035655 | 28.2953359 | 6.06312097 | 3.06E-06 | 2.41E-06 |
| rs556861356 | 2 | 52686390 | LOC730100(dist=51335),MIR4431(dist=243270) | T | A | 0.00035655 | 28.2953359 | 6.06312097 | 3.06E-06 | 2.41E-06 |
| rs572161315 | 2 | 52687214 | LOC730100(dist=52159),MIR4431(dist=242446) | G | C | 0.00035655 | 28.2953359 | 6.06312097 | 3.06E-06 | 2.41E-06 |
| rs4380279 | 2 | 52688148 | LOC730100(dist=53093),MIR4431(dist=241512) | C | T | 0.00035655 | 28.2953359 | 6.06312097 | 3.06E-06 | 2.41E-06 |
| rs531911265 | 2 | 52842946 | LOC730100(dist=207891),MIR4431(dist=86714) | G | C | 0.00035691 | 28.2949755 | 6.06309641 | 3.06E-06 | 2.41E-06 |
| rs572170177 | 2 | 52804757 | LOC730100(dist=169702),MIR4431(dist=124903) | A | G | 0.00035667 | 28.2944946 | 6.06311883 | 3.06E-06 | 2.41E-06 |
| rs561186991 | 2 | 52806051 | LOC730100(dist=170996),MIR4431(dist=123609) | C | T | 0.00035667 | 28.2944946 | 6.06311883 | 3.06E-06 | 2.41E-06 |

| | | | | | | | | | | |
|---|---|---|---|---|---|---|---|---|---|---|
| rs531508153 | 2 | 52807272 | LOC730100(dist=172217),MIR4431(dist=122388) | G | A | 0.00035667 | 28.2944946 | 6.06311883 | 3.06E-06 | 2.41E-06 |
| rs552186163 | 2 | 55744929 | CCDC88A(dist=97872),CFAP36(dist=1802) | T | C | 0.00035584 | 28.4011127 | 6.08597375 | 3.06E-06 | 2.41E-06 |
| rs538136531 | 2 | 52547581 | LOC730100 | C | T | 0.00035691 | 28.294404 | 6.06312904 | 3.06E-06 | 2.41E-06 |
| rs12312017 | 12 | 60194008 | SLC16A7(dist=10373),FAM19A2(dist=1908021) | G | A | 0.00035679 | 28.294495 | 6.06315442 | 3.06E-06 | 2.41E-06 |
| rs376638129 | 2 | 53034115 | MIR4431(dist=104362),ASB3(dist=863002) | G | A | 0.00035667 | 28.29422 | 6.06312466 | 3.06E-06 | 2.41E-06 |
| rs555469022 | 2 | 52557221 | LOC730100 | C | A | 0.00035679 | 28.2941295 | 6.06310701 | 3.06E-06 | 2.41E-06 |
| rs193221137 | 2 | 52734017 | LOC730100(dist=98962),MIR4431(dist=195643) | T | C | 0.00035667 | 28.294139 | 6.06312136 | 3.06E-06 | 2.41E-06 |
| rs377145324 | 2 | 52734248 | LOC730100(dist=99193),MIR4431(dist=195412) | C | T | 0.00035667 | 28.294139 | 6.06312136 | 3.06E-06 | 2.41E-06 |
| rs190751367 | 2 | 52734316 | LOC730100(dist=99261),MIR4431(dist=195344) | G | A | 0.00035667 | 28.294139 | 6.06312136 | 3.06E-06 | 2.41E-06 |
| rs530397536 | 2 | 52734395 | LOC730100(dist=99340),MIR4431(dist=195265) | A | C | 0.00035667 | 28.294139 | 6.06312136 | 3.06E-06 | 2.41E-06 |
| rs374373550 | 2 | 52735159 | LOC730100(dist=100104),MIR4431(dist=194501) | C | T | 0.00035667 | 28.294139 | 6.06312136 | 3.06E-06 | 2.41E-06 |
| rs373775879 | 2 | 52732897 | LOC730100(dist=97842),MIR4431(dist=196763) | C | A | 0.00035667 | 28.294139 | 6.06312136 | 3.06E-06 | 2.41E-06 |
| rs376126118 | 2 | 52733140 | LOC730100(dist=98085),MIR4431(dist=196520) | C | T | 0.00035667 | 28.294139 | 6.06312136 | 3.06E-06 | 2.41E-06 |
| rs544170814 | 2 | 52733150 | LOC730100(dist=98095),MIR4431(dist=196510) | C | G | 0.00035667 | 28.294139 | 6.06312136 | 3.06E-06 | 2.41E-06 |
| rs374849837 | 2 | 52733210 | LOC730100(dist=98155),MIR4431(dist=196450) | A | G | 0.00035667 | 28.294139 | 6.06312136 | 3.06E-06 | 2.41E-06 |
| rs185117219 | 2 | 52733514 | LOC730100(dist=98459),MIR4431(dist=196146) | A | G | 0.00035667 | 28.294139 | 6.06312136 | 3.06E-06 | 2.41E-06 |
| rs191491025 | 2 | 52733708 | LOC730100(dist=98653),MIR4431(dist=195952) | C | T | 0.00035667 | 28.294139 | 6.06312136 | 3.06E-06 | 2.41E-06 |
| rs183840040 | 2 | 52733748 | LOC730100(dist=98693),MIR4431(dist=195912) | A | G | 0.00035667 | 28.294139 | 6.06312136 | 3.06E-06 | 2.41E-06 |
| rs376687316 | 2 | 52733882 | LOC730100(dist=98827),MIR4431(dist=195778) | C | T | 0.00035667 | 28.294139 | 6.06312136 | 3.06E-06 | 2.41E-06 |
| rs182110122 | 2 | 52733968 | LOC730100(dist=98913),MIR4431(dist=195692) | G | A | 0.00035667 | 28.294139 | 6.06312136 | 3.06E-06 | 2.41E-06 |
| rs1483882 | 2 | 52953704 | MIR4431(dist=23951),ASB3(dist=943413) | A | G | 0.00035809 | 28.2940356 | 6.06310491 | 3.06E-06 | 2.41E-06 |
| rs565522352 | 2 | 52855488 | LOC730100(dist=220433),MIR4431(dist=74172) | T | C | 0.00035667 | 28.2938007 | 6.06311197 | 3.06E-06 | 2.41E-06 |
| rs547807201 | 2 | 52856336 | LOC730100(dist=221281),MIR4431(dist=73324) | G | C | 0.00035667 | 28.2938007 | 6.06311197 | 3.06E-06 | 2.41E-06 |
| rs547794581 | 2 | 52857107 | LOC730100(dist=222052),MIR4431(dist=72553) | G | A | 0.00035667 | 28.2938007 | 6.06311197 | 3.06E-06 | 2.41E-06 |
| rs576606058 | 2 | 52866519 | LOC730100(dist=231464),MIR4431(dist=63141) | T | C | 0.00035667 | 28.2938007 | 6.06311197 | 3.06E-06 | 2.41E-06 |
| rs543958949 | 2 | 52866882 | LOC730100(dist=231827),MIR4431(dist=62778) | A | C | 0.00035667 | 28.2938007 | 6.06311197 | 3.06E-06 | 2.41E-06 |
| rs559090140 | 2 | 52866914 | LOC730100(dist=231859),MIR4431(dist=62746) | T | G | 0.00035667 | 28.2938007 | 6.06311197 | 3.06E-06 | 2.41E-06 |
| rs577331164 | 2 | 52867026 | LOC730100(dist=231971),MIR4431(dist=62634) | T | C | 0.00035667 | 28.2938007 | 6.06311197 | 3.06E-06 | 2.41E-06 |
| rs528226082 | 2 | 52870621 | LOC730100(dist=235566),MIR4431(dist=59039) | T | C | 0.00035667 | 28.2938007 | 6.06311197 | 3.06E-06 | 2.41E-06 |
| rs369074393 | 2 | 52972000 | MIR4431(dist=42247),ASB3(dist=925117) | A | T | 0.00035667 | 28.2937031 | 6.06311403 | 3.06E-06 | 2.41E-06 |
| rs372007188 | 2 | 52974427 | MIR4431(dist=44674),ASB3(dist=922690) | C | A | 0.00035667 | 28.2937031 | 6.06311403 | 3.06E-06 | 2.41E-06 |
| rs6736060 | 2 | 52993749 | MIR4431(dist=63996),ASB3(dist=903368) | A | G | 0.00035845 | 28.2925281 | 6.06296714 | 3.06E-06 | 2.41E-06 |
| rs548867931 | 11 | 116625948 | BUD13 | G | T | 0.00035643 | 28.3493723 | 6.07518468 | 3.06E-06 | 2.41E-06 |
| rs376043750 | 2 | 52729403 | LOC730100(dist=94348),MIR4431(dist=200257) | T | C | 0.00035679 | 28.2929161 | 6.06311898 | 3.07E-06 | 2.41E-06 |
| rs370547329 | 2 | 52729443 | LOC730100(dist=94388),MIR4431(dist=200217) | C | T | 0.00035679 | 28.2929161 | 6.06311898 | 3.07E-06 | 2.41E-06 |
| rs371923479 | 2 | 52730307 | LOC730100(dist=95252),MIR4431(dist=199353) | A | C | 0.00035679 | 28.2929161 | 6.06311898 | 3.07E-06 | 2.41E-06 |
| rs375202743 | 2 | 52730349 | LOC730100(dist=95294),MIR4431(dist=199311) | C | T | 0.00035679 | 28.2929161 | 6.06311898 | 3.07E-06 | 2.41E-06 |
| rs375581355 | 2 | 52730992 | LOC730100(dist=95937),MIR4431(dist=198668) | A | G | 0.00035679 | 28.2929161 | 6.06311898 | 3.07E-06 | 2.41E-06 |
| rs189626675 | 2 | 52731052 | LOC730100(dist=95997),MIR4431(dist=198608) | G | T | 0.00035679 | 28.2929161 | 6.06311898 | 3.07E-06 | 2.41E-06 |
| rs375248108 | 2 | 52731202 | LOC730100(dist=96147),MIR4431(dist=198458) | G | C | 0.00035679 | 28.2929161 | 6.06311898 | 3.07E-06 | 2.41E-06 |
| rs368013119 | 2 | 52731376 | LOC730100(dist=96321),MIR4431(dist=198284) | C | T | 0.00035679 | 28.2929161 | 6.06311898 | 3.07E-06 | 2.41E-06 |
| rs376675214 | 2 | 52731939 | LOC730100(dist=96884),MIR4431(dist=197721) | C | T | 0.00035679 | 28.2929161 | 6.06311898 | 3.07E-06 | 2.41E-06 |
| rs192842229 | 2 | 52732817 | LOC730100(dist=97762),MIR4431(dist=196843) | G | A | 0.00035679 | 28.2929161 | 6.06311898 | 3.07E-06 | 2.41E-06 |
| rs374368987 | 2 | 52912191 | LOC730100(dist=277136),MIR4431(dist=17469) | T | C | 0.00035821 | 28.2927366 | 6.06313621 | 3.07E-06 | 2.41E-06 |
| rs371296017 | 2 | 52537462 | LOC730100 | G | T | 0.00035667 | 28.2922017 | 6.06312291 | 3.07E-06 | 2.41E-06 |
| rs573025486 | 2 | 52539006 | LOC730100 | G | A | 0.00035667 | 28.2922017 | 6.06312291 | 3.07E-06 | 2.41E-06 |
| rs755646528 | 9 | 120549489 | TLR4(dist=69720),BRINP1(dist=1379419) | G | A | 0.00035714 | 28.3767322 | 6.08123995 | 3.07E-06 | 2.41E-06 |
| rs533790736 | 2 | 53148225 | MIR4431(dist=218472),ASB3(dist=748892) | G | A | 0.00035774 | 28.2918518 | 6.06306947 | 3.07E-06 | 2.41E-06 |
| rs563451204 | 12 | 63484318 | PPM1H(dist=155653),AVPR1A(dist=52221) | T | C | 0.00035702 | 28.2915567 | 6.06310946 | 3.07E-06 | 2.41E-06 |
| rs116822094 | 2 | 52543145 | LOC730100 | C | G | 0.00035679 | 28.2917282 | 6.06316347 | 3.07E-06 | 2.41E-06 |
| rs927161683 | 9 | 120693777 | TLR4(dist=214008),BRINP1(dist=1235131) | G | A | 0.00035667 | 28.3000117 | 6.06499065 | 3.07E-06 | 2.41E-06 |
| rs981157 | 2 | 52975415 | MIR4431(dist=45662),ASB3(dist=921702) | C | T | 0.00035726 | 28.2906748 | 6.06311941 | 3.07E-06 | 2.41E-06 |

| | | | | | | | | | | |
|---|---|---|---|---|---|---|---|---|---|---|
| rs144116923 | 2 | 52996116 | MIR4431(dist=66363),ASB3(dist=901001) | A | G | 0.00035809 | 28.2896541 | 6.06297619 | 3.07E-06 | 2.41E-06 |
| rs4392298 | 2 | 53123847 | MIR4431(dist=194094),ASB3(dist=773270) | T | C | 0.00035702 | 28.2898707 | 6.06308976 | 3.07E-06 | 2.41E-06 |
| rs551132210 | 2 | 53160711 | MIR4431(dist=230958),ASB3(dist=736406) | T | C | 0.00035691 | 28.2900315 | 6.06313122 | 3.07E-06 | 2.41E-06 |
| rs111276455 | 19 | 450507 | SHC2 | C | T | 0.00036285 | 28.2844317 | 6.06196925 | 3.07E-06 | 2.41E-06 |
| rs75408787 | 19 | 450626 | SHC2 | C | T | 0.00036285 | 28.2844317 | 6.06196925 | 3.07E-06 | 2.41E-06 |
| rs74459457 | 19 | 450627 | SHC2 | A | G | 0.00036285 | 28.2844317 | 6.06196925 | 3.07E-06 | 2.41E-06 |
| rs73489617 | 19 | 450715 | SHC2 | C | T | 0.00036285 | 28.2844317 | 6.06196925 | 3.07E-06 | 2.41E-06 |
| rs111367969 | 19 | 450814 | SHC2 | G | A | 0.00036285 | 28.2844317 | 6.06196925 | 3.07E-06 | 2.41E-06 |
| rs565861418 | 2 | 53157289 | MIR4431(dist=227536),ASB3(dist=739828) | T | C | 0.00035833 | 28.2888609 | 6.06303078 | 3.07E-06 | 2.41E-06 |
| rs549215614 | 2 | 52534543 | LOC730100 | T | C | 0.00035679 | 28.289047 | 6.06312265 | 3.07E-06 | 2.41E-06 |
| rs567527525 | 2 | 52534646 | LOC730100 | A | G | 0.00035679 | 28.289047 | 6.06312265 | 3.07E-06 | 2.41E-06 |
| rs538141205 | 2 | 52535009 | LOC730100 | T | C | 0.00035679 | 28.289047 | 6.06312265 | 3.07E-06 | 2.41E-06 |
| rs368894322 | 2 | 52536103 | LOC730100 | G | A | 0.00035679 | 28.289047 | 6.06312265 | 3.07E-06 | 2.41E-06 |
| rs369224228 | 2 | 52536378 | LOC730100 | C | T | 0.00035679 | 28.289047 | 6.06312265 | 3.07E-06 | 2.41E-06 |
| rs546159418 | 2 | 53139536 | MIR4431(dist=209783),ASB3(dist=757581) | C | T | 0.00035988 | 28.285932 | 6.06250418 | 3.08E-06 | 2.41E-06 |
| rs552519443 | 2 | 52503465 | LOC730100 | A | G | 0.00035049 | 28.7760539 | 6.1676541 | 3.08E-06 | 2.41E-06 |
| rs1568992 | 2 | 52708546 | LOC730100(dist=73491),MIR4431(dist=221114) | C | T | 0.00035714 | 28.2878253 | 6.06311332 | 3.08E-06 | 2.41E-06 |
| rs150859756 | 2 | 52815352 | LOC730100(dist=180297),MIR4431(dist=114308) | C | T | 0.00036047 | 28.2779221 | 6.06110052 | 3.08E-06 | 2.41E-06 |
| rs556888371 | 2 | 52616642 | LINC01867 | G | T | 0.00035904 | 28.2879432 | 6.06328002 | 3.08E-06 | 2.41E-06 |
| rs1843030 | 2 | 52841168 | LOC730100(dist=206113),MIR4431(dist=88492) | G | A | 0.00035679 | 28.2870889 | 6.06310077 | 3.08E-06 | 2.41E-06 |
| rs542789750 | 2 | 52842539 | LOC730100(dist=207484),MIR4431(dist=87121) | T | C | 0.00035679 | 28.2870889 | 6.06310077 | 3.08E-06 | 2.41E-06 |
| rs550086640 | 2 | 52680796 | LOC730100(dist=45741),MIR4431(dist=248864) | T | A | 0.00035667 | 28.2867607 | 6.06314113 | 3.08E-06 | 2.41E-06 |
| rs570590317 | 2 | 52859091 | LOC730100(dist=224036),MIR4431(dist=70569) | C | T | 0.00035714 | 28.2865369 | 6.06311096 | 3.08E-06 | 2.41E-06 |
| rs771876650 | 9 | 120670555 | TLR4(dist=190786),BRINP1(dist=1258353) | G | A | 0.00035702 | 28.2861307 | 6.06308046 | 3.08E-06 | 2.41E-06 |
| rs140699806 | 2 | 52840408 | LOC730100(dist=205353),MIR4431(dist=89252) | A | G | 0.00035726 | 28.2860268 | 6.06309292 | 3.08E-06 | 2.41E-06 |
| rs550930665 | 2 | 53143403 | MIR4431(dist=213650),ASB3(dist=753714) | G | T | 0.00035928 | 28.2828789 | 6.0626231 | 3.08E-06 | 2.41E-06 |
| rs572045690 | 2 | 53168467 | MIR4431(dist=238714),ASB3(dist=728650) | C | G | 0.00035702 | 28.2851447 | 6.06312194 | 3.08E-06 | 2.41E-06 |
| rs544087633 | 2 | 53137882 | MIR4431(dist=208129),ASB3(dist=759235) | A | G | 0.00035988 | 28.2819826 | 6.06246108 | 3.08E-06 | 2.41E-06 |
| rs562793161 | 2 | 53138286 | MIR4431(dist=208533),ASB3(dist=758831) | T | C | 0.00035988 | 28.2819826 | 6.06246108 | 3.08E-06 | 2.41E-06 |
| rs533123190 | 2 | 53138320 | MIR4431(dist=208567),ASB3(dist=758797) | A | G | 0.00035988 | 28.2819826 | 6.06246108 | 3.08E-06 | 2.41E-06 |
| rs376179130 | 2 | 52920989 | LOC730100(dist=285934),MIR4431(dist=8671) | T | G | 0.00035762 | 28.2843043 | 6.06305561 | 3.09E-06 | 2.41E-06 |
| rs532640589 | 2 | 52605778 | LINC01867,LOC730100 | T | C | 0.00035916 | 28.2851062 | 6.06327466 | 3.09E-06 | 2.41E-06 |
| rs573835048 | 2 | 53137673 | MIR4431(dist=207920),ASB3(dist=759444) | C | A | 0.00035976 | 28.2812233 | 6.06246744 | 3.09E-06 | 2.41E-06 |
| rs529411347 | 2 | 53159441 | MIR4431(dist=229688),ASB3(dist=737676) | T | C | 0.00035786 | 28.2841498 | 6.0631203 | 3.09E-06 | 2.41E-06 |
| rs139929104 | 2 | 52996963 | MIR4431(dist=67210),ASB3(dist=900154) | G | C | 0.00035774 | 28.2831039 | 6.06301726 | 3.09E-06 | 2.41E-06 |
| rs569417827 | 2 | 52849837 | LOC730100(dist=214782),MIR4431(dist=79823) | C | T | 0.00035691 | 28.2829336 | 6.06308727 | 3.09E-06 | 2.41E-06 |
| rs556358565 | 2 | 53134151 | MIR4431(dist=204398),ASB3(dist=762966) | G | T | 0.00035976 | 28.27959 | 6.06241669 | 3.09E-06 | 2.41E-06 |
| rs555516619 | 2 | 53135616 | MIR4431(dist=205863),ASB3(dist=761501) | T | C | 0.00035976 | 28.27959 | 6.06241669 | 3.09E-06 | 2.41E-06 |
| rs73489613 | 19 | 449982 | SHC2 | C | A | 0.00036285 | 28.2778705 | 6.06206267 | 3.09E-06 | 2.41E-06 |
| rs563775596 | 2 | 52518812 | LOC730100 | A | C | 0.00035702 | 28.2826757 | 6.06311554 | 3.09E-06 | 2.41E-06 |
| rs1922208 | 2 | 52519636 | LOC730100 | C | A | 0.00035702 | 28.2826757 | 6.06311554 | 3.09E-06 | 2.41E-06 |
| rs546338448 | 2 | 52519976 | LOC730100 | G | C | 0.00035702 | 28.2826757 | 6.06311554 | 3.09E-06 | 2.41E-06 |
| rs117405005 | 18 | 178216 | USP14 | C | T | 0.0003556 | 28.5098616 | 6.11211485 | 3.09E-06 | 2.41E-06 |
| rs79084599 | 2 | 52997421 | MIR4431(dist=67668),ASB3(dist=899696) | A | T | 0.00035774 | 28.2807567 | 6.0630112 | 3.09E-06 | 2.41E-06 |
| rs146381520 | 12 | 60080996 | SLC16A7 | G | C | 0.00035881 | 28.2801188 | 6.0629846 | 3.10E-06 | 2.41E-06 |
| rs77744218 | 19 | 450302 | SHC2 | A | G | 0.00036332 | 28.2753933 | 6.06197621 | 3.10E-06 | 2.41E-06 |
| rs531515532 | 2 | 55556616 | CCDC88A | C | G | 0.00035881 | 28.3394806 | 6.07572591 | 3.10E-06 | 2.41E-06 |
| rs145796806 | 11 | 116650184 | ZPR1 | C | T | 0.00035619 | 28.3656422 | 6.08134455 | 3.10E-06 | 2.41E-06 |
| rs145866128 | 19 | 450803 | SHC2 | G | A | 0.00046803 | 27.8771467 | 5.97661597 | 3.10E-06 | 2.41E-06 |
| rs76490838 | 19 | 450238 | SHC2 | T | G | 0.0003632 | 28.2754864 | 6.0620482 | 3.10E-06 | 2.41E-06 |
| rs537456120 | 2 | 52875961 | LOC730100(dist=240906),MIR4431(dist=53699) | A | T | 0.00035667 | 28.2802315 | 6.06311311 | 3.10E-06 | 2.41E-06 |
| rs533762077 | 2 | 53164653 | MIR4431(dist=234900),ASB3(dist=732464) | G | A | 0.00035726 | 28.27992 | 6.06310862 | 3.10E-06 | 2.41E-06 |

| | | | | | | | | | | |
|---|---|---|---|---|---|---|---|---|---|---|
| rs545785774 | 2 | 52884260 | LOC730100(dist=249205),MIR4431(dist=45400) | G | A | 0.00035691 | 28.2799467 | 6.06315336 | 3.10E-06 | 2.41E-06 |
| rs189412822 | 2 | 59920678 | LINC01793(dist=414143),MIR4432HG(dist=665673) | A | G | 0.00042025 | 28.0610244 | 6.01624777 | 3.10E-06 | 2.41E-06 |
| rs117717189 | 2 | 39154242 | ARHGEF33 | T | C | 0.00073199 | 18.8777955 | 4.04739517 | 3.10E-06 | 4.05E-06 |
| rs1922190 | 2 | 52512210 | LOC730100 | A | G | 0.00035691 | 28.2795993 | 6.06316678 | 3.10E-06 | 2.41E-06 |
| rs535265668 | 2 | 52741408 | LOC730100(dist=106353),MIR4431(dist=188252) | C | T | 0.00035714 | 28.279413 | 6.06313429 | 3.10E-06 | 2.41E-06 |
| rs574902418 | 2 | 52737341 | LOC730100(dist=102286),MIR4431(dist=192319) | A | G | 0.0003575 | 28.279487 | 6.0631562 | 3.10E-06 | 2.41E-06 |
| rs535792665 | 2 | 52873266 | LOC730100(dist=238211),MIR4431(dist=56394) | T | A | 0.00035679 | 28.2786964 | 6.06310411 | 3.10E-06 | 2.41E-06 |
| rs1949927 | 2 | 52703501 | LOC730100(dist=68446),MIR4431(dist=226159) | C | T | 0.00035881 | 28.2776615 | 6.06306155 | 3.10E-06 | 2.41E-06 |
| rs572615835 | 2 | 54790377 | SPTBN1 | C | G | 0.00040064 | 27.9824079 | 5.99976166 | 3.10E-06 | 2.41E-06 |
| rs533586809 | 2 | 53163001 | MIR4431(dist=233248),ASB3(dist=734116) | A | G | 0.00035762 | 28.2777546 | 6.063094 | 3.10E-06 | 2.41E-06 |
| rs541382616 | 2 | 54958264 | EML6 | A | G | 0.00036178 | 28.2760249 | 6.06277334 | 3.10E-06 | 2.41E-06 |
| rs536813480 | 2 | 52850328 | LOC730100(dist=215273),MIR4431(dist=79332) | G | T | 0.0003575 | 28.2773997 | 6.06307477 | 3.10E-06 | 2.41E-06 |
| rs79621546 | 19 | 452976 | SHC2 | G | A | 0.00037057 | 28.2702362 | 6.06155298 | 3.10E-06 | 2.41E-06 |
| rs377508110 | 2 | 53062231 | MIR4431(dist=132478),ASB3(dist=834886) | A | G | 0.00035714 | 28.277132 | 6.06305979 | 3.10E-06 | 2.41E-06 |
| rs538042079 | 2 | 53149162 | MIR4431(dist=219409),ASB3(dist=747955) | C | A | 0.00035904 | 28.2762814 | 6.06288639 | 3.10E-06 | 2.41E-06 |
| rs531732261 | 2 | 53151413 | MIR4431(dist=221660),ASB3(dist=745704) | C | T | 0.00035904 | 28.2762814 | 6.06288639 | 3.10E-06 | 2.41E-06 |
| rs370296178 | 2 | 52954734 | MIR4431(dist=24981),ASB3(dist=942383) | G | A | 0.0003575 | 28.2769798 | 6.06306438 | 3.10E-06 | 2.41E-06 |
| rs558373841 | 2 | 52850981 | LOC730100(dist=215926),MIR4431(dist=78679) | A | C | 0.00035762 | 28.2768718 | 6.06306164 | 3.10E-06 | 2.41E-06 |
| rs577065460 | 2 | 52851096 | LOC730100(dist=216041),MIR4431(dist=78564) | G | A | 0.00035762 | 28.2768718 | 6.06306164 | 3.10E-06 | 2.41E-06 |
| rs374221587 | 2 | 52729166 | LOC730100(dist=94111),MIR4431(dist=200494) | C | T | 0.00035821 | 28.2764573 | 6.06312934 | 3.11E-06 | 2.41E-06 |
| rs574920246 | 2 | 52514121 | LOC730100 | C | A | 0.00035702 | 28.2764239 | 6.06316432 | 3.11E-06 | 2.41E-06 |
| rs181035023 | 12 | 60083836 | SLC16A7 | A | C | 0.00035893 | 28.2754525 | 6.06296699 | 3.11E-06 | 2.41E-06 |
| rs112434157 | 19 | 450231 | SHC2 | A | G | 0.0003632 | 28.270309 | 6.06207015 | 3.11E-06 | 2.41E-06 |
| rs367906151 | 2 | 53086176 | MIR4431(dist=156423),ASB3(dist=810941) | C | T | 0.00036071 | 28.2740881 | 6.06308785 | 3.11E-06 | 2.41E-06 |
| rs73489629 | 19 | 451908 | SHC2 | A | G | 0.00036344 | 28.2679198 | 6.06177646 | 3.11E-06 | 2.41E-06 |
| rs570097903 | 2 | 52874483 | LOC730100(dist=239428),MIR4431(dist=55177) | A | G | 0.00035786 | 28.273778 | 6.06311406 | 3.11E-06 | 2.41E-06 |
| rs373904199 | 2 | 53099935 | MIR4431(dist=170182),ASB3(dist=797182) | T | C | 0.00036011 | 28.2728561 | 6.06294256 | 3.11E-06 | 2.41E-06 |
| rs559036395 | 2 | 52736744 | LOC730100(dist=101689),MIR4431(dist=192916) | G | T | 0.00035786 | 28.2731962 | 6.0631323 | 3.11E-06 | 2.41E-06 |
| rs577415351 | 2 | 52736773 | LOC730100(dist=101718),MIR4431(dist=192887) | T | C | 0.00035786 | 28.2731962 | 6.0631323 | 3.11E-06 | 2.41E-06 |
| rs557335377 | 2 | 52662014 | LOC730100(dist=26959),MIR4431(dist=267646) | G | A | 0.00035833 | 28.2731345 | 6.0631909 | 3.12E-06 | 2.41E-06 |
| rs535448254 | 2 | 52616242 | LINC01867,LOC730100 | T | C | 0.00035845 | 28.2735878 | 6.06329161 | 3.12E-06 | 2.41E-06 |
| rs17123009 | 12 | 60175789 | SLC16A7(NM_001270623:c.*23290>0,NM_001270622:c.*2 | G | A | 0.00035952 | 28.2702854 | 6.06261943 | 3.12E-06 | 2.41E-06 |
| rs143416013 | 12 | 60099743 | SLC16A7 | T | C | 0.00035952 | 28.2710784 | 6.0628825 | 3.12E-06 | 2.41E-06 |
| rs139335791 | 12 | 60100424 | SLC16A7 | T | A | 0.00035952 | 28.2710784 | 6.0628825 | 3.12E-06 | 2.41E-06 |
| rs539714680 | 12 | 60101339 | SLC16A7 | A | G | 0.00035952 | 28.2710784 | 6.0628825 | 3.12E-06 | 2.41E-06 |
| rs73489619 | 19 | 451465 | SHC2 | G | A | 0.00036356 | 28.2655866 | 6.06180152 | 3.12E-06 | 2.41E-06 |
| rs73489621 | 19 | 451583 | SHC2 | C | G | 0.00036356 | 28.2655866 | 6.06180152 | 3.12E-06 | 2.41E-06 |
| rs374615507 | 2 | 53120357 | MIR4431(dist=190604),ASB3(dist=776760) | C | T | 0.00035774 | 28.2712515 | 6.0630206 | 3.12E-06 | 2.41E-06 |
| rs575628829 | 2 | 52736374 | LOC730100(dist=101319),MIR4431(dist=193286) | T | G | 0.00035797 | 28.2710572 | 6.06311983 | 3.12E-06 | 2.41E-06 |
| rs537393677 | 2 | 52736378 | LOC730100(dist=101323),MIR4431(dist=193282) | G | A | 0.00035797 | 28.2710572 | 6.06311983 | 3.12E-06 | 2.41E-06 |
| rs557194704 | 2 | 52600815 | LINC01867,LOC730100 | T | C | 0.00035893 | 28.2717435 | 6.06331713 | 3.12E-06 | 2.41E-06 |
| rs115998730 | 12 | 60109624 | SLC16A7 | T | C | 0.00035976 | 28.2682262 | 6.06283143 | 3.12E-06 | 2.41E-06 |
| rs147462597 | 12 | 60121592 | SLC16A7 | G | T | 0.00035976 | 28.2682262 | 6.06283143 | 3.12E-06 | 2.41E-06 |
| rs187674492 | 12 | 60122836 | SLC16A7 | T | G | 0.00035976 | 28.2682262 | 6.06283143 | 3.12E-06 | 2.41E-06 |
| rs146527019 | 12 | 60159496 | SLC16A7 | A | G | 0.00035976 | 28.2682262 | 6.06283143 | 3.12E-06 | 2.41E-06 |
| rs530144269 | 2 | 52621533 | LOC730100 | T | C | 0.00035869 | 28.2693078 | 6.06330874 | 3.13E-06 | 2.41E-06 |
| rs552629121 | 2 | 52623746 | LOC730100 | T | C | 0.00035869 | 28.2693078 | 6.06330874 | 3.13E-06 | 2.41E-06 |
| rs533251316 | 2 | 52752898 | LOC730100(dist=117843),MIR4431(dist=176762) | G | A | 0.00035738 | 28.2678348 | 6.06315558 | 3.13E-06 | 2.41E-06 |
| rs566795496 | 2 | 52754974 | LOC730100(dist=119919),MIR4431(dist=174686) | A | G | 0.00035738 | 28.2678348 | 6.06315558 | 3.13E-06 | 2.41E-06 |
| rs546326444 | 5 | 180708080 | TRIM52-AS1(dist=8772),LOC100133331(dist=42427) | T | C | 0.00041657 | 27.6531869 | 5.93140437 | 3.13E-06 | 2.41E-06 |
| rs74363876 | 2 | 52517336 | LOC730100 | G | A | 0.00035714 | 28.266951 | 6.06320579 | 3.13E-06 | 2.41E-06 |
| rs535204752 | 2 | 52629081 | LOC730100 | T | C | 0.00035833 | 28.2673555 | 6.06329899 | 3.13E-06 | 2.41E-06 |

| | | | | | | | | | | |
|---|---|---|---|---|---|---|---|---|---|---|
| rs533135336 | 2 | 52766733 | LOC730100(dist=131678),MIR4431(dist=162927) | C | T | 0.00035916 | 28.2663022 | 6.06312294 | 3.13E-06 | 2.41E-06 |
| rs560147099 | 2 | 52768261 | LOC730100(dist=133206),MIR4431(dist=161399) | G | C | 0.00035916 | 28.2663022 | 6.06312294 | 3.13E-06 | 2.41E-06 |
| rs568469982 | 2 | 52770892 | LOC730100(dist=135837),MIR4431(dist=158768) | A | G | 0.00035916 | 28.2663022 | 6.06312294 | 3.13E-06 | 2.41E-06 |
| rs535560757 | 2 | 52771898 | LOC730100(dist=136843),MIR4431(dist=157762) | T | G | 0.00035916 | 28.2663022 | 6.06312294 | 3.13E-06 | 2.41E-06 |
| rs539595685 | 2 | 52775275 | LOC730100(dist=140220),MIR4431(dist=154385) | T | A | 0.00035916 | 28.2663022 | 6.06312294 | 3.13E-06 | 2.41E-06 |
| rs573446993 | 2 | 52775400 | LOC730100(dist=140345),MIR4431(dist=154260) | G | T | 0.00035916 | 28.2663022 | 6.06312294 | 3.13E-06 | 2.41E-06 |
| rs555055715 | 2 | 52778733 | LOC730100(dist=143678),MIR4431(dist=150927) | C | T | 0.00035916 | 28.2663022 | 6.06312294 | 3.13E-06 | 2.41E-06 |
| rs573548698 | 2 | 52778763 | LOC730100(dist=143708),MIR4431(dist=150897) | C | T | 0.00035916 | 28.2663022 | 6.06312294 | 3.13E-06 | 2.41E-06 |
| rs13407120 | 2 | 52736614 | LOC730100(dist=101559),MIR4431(dist=193046) | C | T | 0.00035797 | 28.2661611 | 6.06313444 | 3.13E-06 | 2.41E-06 |
| rs117725199 | 18 | 289856 | THOC1(dist=21797),COLEC12(dist=29499) | C | A | 0.00035821 | 28.2646611 | 6.06307056 | 3.13E-06 | 2.41E-06 |
| rs113708455 | 19 | 451089 | SHC2 | T | C | 0.00036392 | 28.2585617 | 6.06189928 | 3.14E-06 | 2.41E-06 |
| rs372931574 | 2 | 52565855 | LOC730100 | A | C | 0.00035774 | 28.2639392 | 6.06319405 | 3.14E-06 | 2.41E-06 |
| rs994068702 | 11 | 120312581 | ARHGEF12 | A | G | 0.00035821 | 28.2635037 | 6.06311669 | 3.14E-06 | 2.41E-06 |
| rs558053953 | 2 | 52630672 | LOC730100 | T | C | 0.00036071 | 28.2639415 | 6.06321622 | 3.14E-06 | 2.41E-06 |
| rs112335970 | 19 | 451158 | SHC2 | T | C | 0.0003638 | 28.2576642 | 6.0618975 | 3.14E-06 | 2.41E-06 |
| rs115064953 | 19 | 451261 | SHC2 | C | T | 0.0003638 | 28.2576642 | 6.0618975 | 3.14E-06 | 2.41E-06 |
| rs371284192 | 2 | 52562748 | LOC730100 | G | T | 0.00035726 | 28.263692 | 6.06319382 | 3.14E-06 | 2.41E-06 |
| rs137932806 | 19 | 536014 | CDC34 | C | T | 0.000372 | 28.7950804 | 6.17745307 | 3.14E-06 | 2.41E-06 |
| rs547319884 | 2 | 52565681 | LOC730100 | C | T | 0.00035762 | 28.2616225 | 6.06320407 | 3.14E-06 | 2.41E-06 |
| rs369867526 | 2 | 52567276 | LOC730100 | G | C | 0.00035762 | 28.2616225 | 6.06320407 | 3.14E-06 | 2.41E-06 |
| rs370767240 | 2 | 52568339 | LOC730100 | A | C | 0.00035774 | 28.2615523 | 6.06319598 | 3.14E-06 | 2.41E-06 |
| rs542724820 | 2 | 52560729 | LOC730100 | C | T | 0.00035714 | 28.2613753 | 6.06320383 | 3.14E-06 | 2.41E-06 |
| rs367851740 | 2 | 52561853 | LOC730100 | G | T | 0.00035714 | 28.2613753 | 6.06320383 | 3.14E-06 | 2.41E-06 |
| rs565131686 | 2 | 52563049 | LOC730100 | T | C | 0.00035714 | 28.2613753 | 6.06320383 | 3.14E-06 | 2.41E-06 |
| rs531702345 | 2 | 52578740 | LOC730100 | A | C | 0.00035714 | 28.2613753 | 6.06320383 | 3.14E-06 | 2.41E-06 |
| rs545730630 | 2 | 52560097 | LOC730100 | T | C | 0.00035738 | 28.2601694 | 6.0631899 | 3.15E-06 | 2.41E-06 |
| rs553795061 | 2 | 52560104 | LOC730100 | G | A | 0.00035738 | 28.2601694 | 6.0631899 | 3.15E-06 | 2.41E-06 |
| rs368720644 | 2 | 52567528 | LOC730100 | C | G | 0.00035762 | 28.2583897 | 6.06320569 | 3.15E-06 | 2.41E-06 |
| rs368707584 | 2 | 52569011 | LOC730100 | G | C | 0.00035762 | 28.2583897 | 6.06320569 | 3.15E-06 | 2.41E-06 |
| rs7421904 | 2 | 52762547 | LOC730100(dist=127492),MIR4431(dist=167113) | G | A | 0.0003575 | 28.2580788 | 6.06316844 | 3.15E-06 | 2.41E-06 |
| rs569029246 | 2 | 52775249 | LOC730100(dist=140194),MIR4431(dist=154411) | C | A | 0.0003575 | 28.2580788 | 6.06316844 | 3.15E-06 | 2.41E-06 |
| rs374706023 | 2 | 52952647 | MIR4431(dist=22894),ASB3(dist=944470) | C | T | 0.00035893 | 28.2575758 | 6.06308577 | 3.15E-06 | 2.41E-06 |
| rs149614586 | 2 | 52559435 | LOC730100 | G | C | 0.00035738 | 28.2580344 | 6.06321 | 3.15E-06 | 2.41E-06 |
| rs554536411 | 2 | 52569404 | LOC730100 | C | T | 0.00035774 | 28.2570928 | 6.06320333 | 3.16E-06 | 2.41E-06 |
| rs375501748 | 2 | 52569709 | LOC730100 | G | A | 0.00035774 | 28.2570928 | 6.06320333 | 3.16E-06 | 2.41E-06 |
| rs373708219 | 2 | 52573401 | LOC730100 | G | A | 0.00035774 | 28.2570928 | 6.06320333 | 3.16E-06 | 2.41E-06 |
| rs112068208 | 19 | 451007 | SHC2 | G | A | 0.00036392 | 28.2505993 | 6.06188633 | 3.16E-06 | 2.41E-06 |
| rs148674242 | 2 | 52569030 | LOC730100 | G | A | 0.00035786 | 28.2555438 | 6.0631985 | 3.16E-06 | 2.41E-06 |
| rs137868799 | 2 | 55621333 | CCDC88A | C | T | 0.0003594 | 28.3414442 | 6.08164772 | 3.16E-06 | 2.41E-06 |
| rs189535388 | 2 | 52662612 | LOC730100(dist=27557),MIR4431(dist=267048) | C | T | 0.00035833 | 28.2553868 | 6.06321345 | 3.16E-06 | 2.41E-06 |
| rs566648755 | 2 | 52534240 | LOC730100 | C | T | 0.00035738 | 28.2550863 | 6.06320551 | 3.16E-06 | 2.41E-06 |
| rs372237158 | 2 | 52569628 | LOC730100 | C | T | 0.00035762 | 28.2542523 | 6.063207 | 3.16E-06 | 2.41E-06 |
| rs369944585 | 2 | 52569929 | LOC730100 | T | A | 0.00035762 | 28.2542523 | 6.063207 | 3.16E-06 | 2.41E-06 |
| rs577741617 | 2 | 52678367 | LOC730100(dist=43312),MIR4431(dist=251293) | G | A | 0.00035714 | 28.252209 | 6.06319505 | 3.17E-06 | 2.41E-06 |
| rs550501118 | 2 | 52532072 | LOC730100 | T | C | 0.0003575 | 28.251911 | 6.06320306 | 3.17E-06 | 2.41E-06 |
| rs533037143 | 2 | 52533449 | LOC730100 | C | G | 0.0003575 | 28.251911 | 6.06320306 | 3.17E-06 | 2.41E-06 |
| rs369943916 | 2 | 52533626 | LOC730100 | G | A | 0.0003575 | 28.251911 | 6.06320306 | 3.17E-06 | 2.41E-06 |
| rs112680260 | 19 | 450067 | SHC2 | T | C | 0.0003638 | 28.2465469 | 6.06205446 | 3.17E-06 | 2.41E-06 |
| rs569104524 | 2 | 52752394 | LOC730100(dist=117339),MIR4431(dist=177266) | G | A | 0.00035904 | 28.2514464 | 6.0631175 | 3.17E-06 | 2.41E-06 |
| rs551669043 | 2 | 52753497 | LOC730100(dist=118442),MIR4431(dist=176163) | G | C | 0.00035904 | 28.2514464 | 6.0631175 | 3.17E-06 | 2.41E-06 |
| rs568453194 | 2 | 52757915 | LOC730100(dist=122860),MIR4431(dist=171745) | T | C | 0.00035904 | 28.2514464 | 6.0631175 | 3.17E-06 | 2.41E-06 |
| rs535990627 | 2 | 52758087 | LOC730100(dist=123032),MIR4431(dist=171573) | T | C | 0.00035904 | 28.2514464 | 6.0631175 | 3.17E-06 | 2.41E-06 |

| | | | | | | | | | | |
|---|---|---|---|---|---|---|---|---|---|---|
| rs575466167 | 2 | 52759253 | LOC730100(dist=124198),MIR4431(dist=170407) | G | C | 0.00035904 | 28.2514464 | 6.0631175 | 3.17E-06 | 2.41E-06 |
| rs56109124 | 2 | 52771895 | LOC730100(dist=136840),MIR4431(dist=157765) | T | A | 0.00036736 | 28.2455406 | 6.0620394 | 3.17E-06 | 2.41E-06 |
| rs61369041 | 2 | 52757732 | LOC730100(dist=122677),MIR4431(dist=171928) | C | T | 0.00036772 | 28.243485 | 6.06203321 | 3.18E-06 | 2.41E-06 |
| rs568891853 | 2 | 52522576 | LOC730100 | A | T | 0.00035762 | 28.2487151 | 6.06319841 | 3.18E-06 | 2.41E-06 |
| rs566590054 | 2 | 52525433 | LOC730100 | A | T | 0.00035762 | 28.2487151 | 6.06319841 | 3.18E-06 | 2.41E-06 |
| rs574004397 | 2 | 52526286 | LOC730100 | C | T | 0.00035762 | 28.2487151 | 6.06319841 | 3.18E-06 | 2.41E-06 |
| rs557087923 | 2 | 52527447 | LOC730100 | T | G | 0.00035762 | 28.2487151 | 6.06319841 | 3.18E-06 | 2.41E-06 |
| rs367619843 | 2 | 52528524 | LOC730100 | C | T | 0.00035762 | 28.2487151 | 6.06319841 | 3.18E-06 | 2.41E-06 |
| rs562008490 | 2 | 52529152 | LOC730100 | T | G | 0.00035762 | 28.2487151 | 6.06319841 | 3.18E-06 | 2.41E-06 |
| rs574057725 | 2 | 55484418 | MTIF2 | C | G | 0.00036023 | 28.2479716 | 6.06304357 | 3.18E-06 | 2.41E-06 |
| rs74585063 | 2 | 39151832 | ARHGEF33 | G | C | 0.00072926 | 18.8657061 | 4.04927692 | 3.18E-06 | 4.05E-06 |
| rs964215441 | 1 | 108294081 | VAV3 | G | A | 0.00035298 | 28.8793593 | 6.19902223 | 3.18E-06 | 2.41E-06 |
| rs558937324 | 2 | 55358340 | RTN4 | T | C | 0.00036213 | 28.2435058 | 6.06289344 | 3.19E-06 | 2.41E-06 |
| rs572181905 | 2 | 52679204 | LOC730100(dist=44149),MIR4431(dist=250456) | A | C | 0.00035726 | 28.2435085 | 6.06320185 | 3.19E-06 | 2.41E-06 |
| rs549035362 | 2 | 52657316 | LOC730100(dist=22261),MIR4431(dist=272344) | A | G | 0.00036071 | 28.2419955 | 6.06295367 | 3.19E-06 | 2.41E-06 |
| rs533086661 | 2 | 54593020 | C2orf73(dist=4306),SPTBN1(dist=90434) | T | C | 0.00037414 | 28.2684104 | 6.06864154 | 3.19E-06 | 2.41E-06 |
| rs537133298 | 2 | 54555797 | ACYP2(dist=23362),C2orf73(dist=2274) | T | C | 0.00037307 | 28.2728523 | 6.07010723 | 3.20E-06 | 2.41E-06 |
| rs368783350 | 2 | 52928992 | MIR4431(dist=668) | T | C | 0.00036154 | 28.2292143 | 6.06076878 | 3.20E-06 | 2.41E-06 |
| rs73284906 | 8 | 82620167 | ZFAND1 | A | T | 0.00039054 | -26.475382 | 5.68466898 | 3.20E-06 | 7.23E-06 |
| rs531639262 | 2 | 52670686 | LOC730100(dist=35631),MIR4431(dist=258974) | G | A | 0.00035738 | 28.234783 | 6.06320598 | 3.21E-06 | 2.41E-06 |
| rs368598615 | 2 | 52930117 | MIR4431(dist=364) | G | A | 0.00036202 | 28.2206207 | 6.06029939 | 3.21E-06 | 2.41E-06 |
| rs759324230 | 5 | 171210528 | FGF18(dist=325898),SMIM23(dist=2348) | C | G | 0.00039256 | 28.211448 | 6.05883056 | 3.22E-06 | 2.41E-06 |
| rs146369142 | 5 | 180621561 | TRIM7(NM_203293:c.*6050>0,NM_203295:c.*6050>0,NM | C | T | 0.00038388 | 28.1864599 | 6.0542607 | 3.23E-06 | 2.41E-06 |
| rs544928443 | 2 | 52675896 | LOC730100(dist=40841),MIR4431(dist=253764) | C | T | 0.00036011 | 28.225786 | 6.06299904 | 3.23E-06 | 2.41E-06 |
| rs560255773 | 2 | 52679092 | LOC730100(dist=44037),MIR4431(dist=250568) | A | G | 0.0003575 | 28.2260325 | 6.06320744 | 3.24E-06 | 2.41E-06 |
| rs17042677 | 2 | 52507535 | LOC730100 | A | T | 0.00035786 | 28.2370349 | 6.06712942 | 3.25E-06 | 2.41E-06 |
| rs193213051 | 2 | 53000443 | MIR4431(dist=70690),ASB3(dist=896674) | G | A | 0.00036285 | 28.2121238 | 6.0620943 | 3.26E-06 | 2.41E-06 |
| rs372377537 | 2 | 52704541 | LOC730100(dist=69486),MIR4431(dist=225119) | T | A | 0.00036011 | 28.2129856 | 6.062503 | 3.26E-06 | 2.41E-06 |
| rs367596224 | 11 | 116707929 | APOA1-AS | G | A | 0.0003632 | 28.617106 | 6.14999631 | 3.27E-06 | 2.41E-06 |
| rs553929719 | 2 | 54661230 | C2orf73(dist=72516),SPTBN1(dist=22224) | G | A | 0.00039838 | 27.9390521 | 6.0045045 | 3.27E-06 | 2.41E-06 |
| rs17841752 | 19 | 536527 | CDC34 | C | T | 0.0003714 | 28.7437711 | 6.17750183 | 3.27E-06 | 2.41E-06 |
| rs565654994 | 3 | 2176571 | CNTN4-AS2 | C | A | 0.00036974 | 28.2485403 | 6.07120257 | 3.27E-06 | 2.41E-06 |
| rs143685503 | 19 | 35088376 | SCGB1B2P | C | T | 0.00242667 | 11.6185194 | 2.49719538 | 3.28E-06 | 5.36E-06 |
| rs530214521 | 2 | 52499703 | LOC730100 | G | T | 0.00034454 | 29.2162256 | 6.28040314 | 3.29E-06 | 2.41E-06 |
| rs373143962 | 2 | 52960571 | MIR4431(dist=30818),ASB3(dist=936546) | T | C | 0.00036178 | 28.2042056 | 6.06289386 | 3.29E-06 | 2.41E-06 |
| rs540445501 | 2 | 52666399 | LOC730100(dist=31344),MIR4431(dist=263261) | T | A | 0.00035833 | 28.2032656 | 6.06321787 | 3.29E-06 | 2.41E-06 |
| rs372492838 | 2 | 52710357 | LOC730100(dist=75302),MIR4431(dist=219303) | A | G | 0.00041276 | 27.9749802 | 6.01459321 | 3.30E-06 | 2.41E-06 |
| rs559783246 | 2 | 52499447 | LOC730100 | G | A | 0.00034407 | 29.2373075 | 6.28604119 | 3.30E-06 | 2.41E-06 |
| rs374739658 | 2 | 52933597 | MIR4431(dist=3844),ASB3(dist=963520) | A | G | 0.00036475 | 28.1689642 | 6.05695323 | 3.31E-06 | 2.41E-06 |
| rs147188679 | 17 | 30023108 | MIR365B(dist=120568),COPRS(dist=155776) | T | C | 0.00044141 | 26.8851537 | 5.78129071 | 3.31E-06 | 2.41E-06 |
| rs12425587 | 12 | 86003568 | ALX1(dist=308007),RASSF9(dist=194763) | A | C | 0.00045769 | 26.7424341 | 5.75063148 | 3.31E-06 | 3.96E-06 |
| rs373411394 | 2 | 52934689 | MIR4431(dist=4936),ASB3(dist=962428) | A | C | 0.00036511 | 28.1606269 | 6.05639811 | 3.32E-06 | 2.41E-06 |
| rs111405156 | 9 | 5628556 | RIC1(dist=563) | T | C | 0.0091368 | 5.39251332 | 1.15975221 | 3.32E-06 | 8.28E-06 |
| rs149566183 | 17 | 30045735 | MIR365B(dist=143195),COPRS(dist=133149) | G | A | 0.00037271 | 28.1812984 | 6.06091121 | 3.32E-06 | 2.41E-06 |
| rs765516203 | 5 | 45064862 | MRPS30(dist=249244),HCN1(dist=190190) | T | G | 0.00034657 | -26.129782 | 5.6197138 | 3.32E-06 | 5.04E-06 |
| rs76821818 | 19 | 439566 | SHC2 | C | T | 0.00036522 | 28.2440862 | 6.07555455 | 3.34E-06 | 2.41E-06 |
| rs543812413 | 2 | 52668274 | LOC730100(dist=33219),MIR4431(dist=261386) | A | G | 0.00035845 | 28.1864044 | 6.0631892 | 3.34E-06 | 2.41E-06 |
| rs564375608 | 2 | 52498780 | LOC730100 | T | A | 0.00034252 | 29.3619662 | 6.31654407 | 3.34E-06 | 2.41E-06 |
| rs111702906 | 19 | 554425 | GZMM(dist=4505),BSG(dist=16852) | C | T | 0.00037295 | 28.1740091 | 6.06127594 | 3.35E-06 | 2.41E-06 |
| rs557798528 | 2 | 52672521 | LOC730100(dist=37466),MIR4431(dist=257139) | A | C | 0.00035809 | 28.1819064 | 6.06317469 | 3.35E-06 | 2.41E-06 |
| rs566637930 | 2 | 52672633 | LOC730100(dist=37578),MIR4431(dist=257027) | G | A | 0.00035809 | 28.1819064 | 6.06317469 | 3.35E-06 | 2.41E-06 |
| rs545765809 | 2 | 52498530 | LOC730100 | T | C | 0.00034181 | 29.402205 | 6.32673675 | 3.36E-06 | 2.41E-06 |

| | | | | | | | | | | |
|---|---|---|---|---|---|---|---|---|---|---|
| rs550698104 | 2 | 52670937 | LOC730100(dist=35882),MIR4431(dist=258723) | T | C | 0.00035821 | 28.1730066 | 6.06316013 | 3.37E-06 | 2.41E-06 |
| rs370658532 | 2 | 53108117 | MIR4431(dist=178364),ASB3(dist=789000) | G | A | 0.00036237 | 28.1586183 | 6.06064285 | 3.38E-06 | 2.41E-06 |
| rs535362015 | 12 | 97739345 | NEDD1(dist=391876),RMST(dist=119454) | T | C | 0.00035488 | 28.9153429 | 6.22370059 | 3.38E-06 | 2.41E-06 |
| rs59090179 | 3 | 24063857 | NR1D2(dist=41748),LINC00691(dist=77608) | A | G | 0.00059449 | 20.1515669 | 4.33766007 | 3.39E-06 | 3.33E-06 |
| rs568953477 | 2 | 52671022 | LOC730100(dist=35967),MIR4431(dist=258638) | G | A | 0.00035845 | 28.1632426 | 6.06315309 | 3.40E-06 | 2.41E-06 |
| rs556737465 | 2 | 52497944 | LOC730100 | G | A | 0.00033991 | 29.523431 | 6.35686614 | 3.41E-06 | 2.41E-06 |
| rs554882632 | 8 | 32202615 | NRG1 | C | T | 0.00101628 | 17.6043309 | 3.7905834 | 3.41E-06 | 3.62E-06 |
| rs55823245 | 1 | 6273206 | RNF207 | C | T | 0.02060483 | 3.645322 | 0.7851738 | 3.44E-06 | 6.59E-06 |
| rs749746138 | 9 | 120852171 | TLR4(dist=372402),BRINP1(dist=1076737) | T | A | 0.00036891 | 28.3063375 | 6.09755455 | 3.45E-06 | 2.41E-06 |
| rs549967454 | 1 | 240778266 | MIR1273E | G | A | 0.00037366 | 28.1382839 | 6.06145424 | 3.45E-06 | 2.41E-06 |
| rs574154145 | 2 | 52496962 | LOC730100 | A | C | 0.00033777 | 29.6598529 | 6.39141919 | 3.47E-06 | 2.41E-06 |
| rs28693834 | 12 | 60179829 | SLC16A7(NM_001270623:c.*63690>0,NM_001270622:c.*( | G | A | 0.00037034 | 28.1252395 | 6.06109285 | 3.48E-06 | 2.41E-06 |
| rs73277208 | 8 | 82449575 | FABP12(dist=5950),IMPA1P1(dist=66544) | A | G | 0.00035702 | -26.59882 | 5.73222226 | 3.48E-06 | 4.71E-06 |
| rs146975892 | 6 | 9048496 | LOC100506207(dist=262818),TFAP2A(dist=1348420) | A | G | 0.00669622 | -6.7903392 | 1.4636728 | 3.50E-06 | 9.34E-06 |
| rs7864736 | 9 | 140189091 | TOR4A(dist=11998),NRARP(dist=4992) | G | A | 0.0765397 | 1.91261 | 0.41234379 | 3.51E-06 | 1.80E-06 |
| rs73275501 | 8 | 82446091 | FABP12(dist=2466),IMPA1P1(dist=70028) | C | T | 0.00035655 | -26.586218 | 5.73218081 | 3.52E-06 | 4.71E-06 |
| rs550145157 | 3 | 1874208 | CNTN6(dist=428916),CNTN4(dist=266279) | G | T | 0.00037842 | 28.1391176 | 6.06794612 | 3.53E-06 | 2.41E-06 |
| rs149425565 | 8 | 110172500 | TRHR(dist=40688),NUDCD1(dist=80648) | A | T | 0.00413858 | 8.1863276 | 1.76532445 | 3.53E-06 | 3.96E-06 |
| rs542228030 | 8 | 32161949 | NRG1 | G | A | 0.00107036 | 16.4923662 | 3.55667773 | 3.53E-06 | 3.62E-06 |
| rs751844754 | 3 | 161619350 | OTOL1(dist=397620),LINC01192(dist=1275681) | T | C | 0.00031745 | 28.9382256 | 6.24178245 | 3.55E-06 | 2.58E-06 |
| rs556131362 | 8 | 32188420 | NRG1 | T | G | 0.0010952 | 15.9856975 | 3.44829791 | 3.56E-06 | 3.62E-06 |
| rs899857383 | 17 | 30843917 | MYO1D | G | A | 0.00037117 | 28.2339171 | 6.09307569 | 3.59E-06 | 2.41E-06 |
| rs750296178 | 9 | 37564340 | FBXO10 | G | A | 0.00039232 | -26.377214 | 5.69306951 | 3.60E-06 | 5.90E-06 |
| rs114248084 | 19 | 450056 | SHC2 | G | A | 0.00037188 | 28.0475289 | 6.05509947 | 3.62E-06 | 2.41E-06 |
| rs534584559 | 2 | 52494745 | LOC730100 | A | C | 0.00033409 | 29.9305191 | 6.46418413 | 3.65E-06 | 2.41E-06 |
| rs113579383 | 9 | 90137915 | DAPK1 | G | A | 0.00037782 | 28.1037064 | 6.06981401 | 3.66E-06 | 2.41E-06 |
| rs372168808 | 2 | 52704357 | LOC730100(dist=69302),MIR4431(dist=225303) | T | A | 0.00036522 | 28.0546162 | 6.05925669 | 3.66E-06 | 2.41E-06 |
| rs374185367 | 2 | 53062136 | MIR4431(dist=132383),ASB3(dist=834981) | G | A | 0.00038888 | 27.8461006 | 6.01436628 | 3.66E-06 | 2.41E-06 |
| rs150294795 | 2 | 39156337 | ARHGEF33 | C | T | 0.00071952 | 18.7738986 | 4.05561424 | 3.67E-06 | 4.05E-06 |
| rs115845689 | 2 | 39155706 | ARHGEF33 | G | A | 0.0007194 | 18.7707986 | 4.05562189 | 3.69E-06 | 4.05E-06 |
| rs779800385 | 12 | 84254349 | TMTC2(dist=725702),SLC6A15(dist=998918) | G | A | 0.00048217 | 25.101365 | 5.4241085 | 3.70E-06 | 3.96E-06 |
| rs572146953 | 21 | 29221832 | LINC00113(dist=98280),LINC00314(dist=163850) | A | G | 0.00041241 | 27.9575815 | 6.04164132 | 3.70E-06 | 6.47E-06 |
| rs929332106 | 8 | 88701395 | CNBD1(dist=306440),DCAF4L2(dist=181576) | T | G | 0.00022546 | -36.619843 | 7.91439618 | 3.71E-06 | 1.25E-06 |
| rs540848381 | 2 | 52571977 | LOC730100 | A | T | 0.00036641 | 28.0398986 | 6.06116645 | 3.73E-06 | 2.41E-06 |
| rs374633957 | 2 | 53042776 | MIR4431(dist=113023),ASB3(dist=854341) | T | C | 0.00036736 | 28.0120095 | 6.05543016 | 3.73E-06 | 2.41E-06 |
| rs76744615 | 2 | 53010207 | MIR4431(dist=80454),ASB3(dist=886910) | G | A | 0.00038781 | 28.021696 | 6.05777422 | 3.73E-06 | 2.41E-06 |
| rs114962859 | 2 | 39177653 | ARHGEF33 | C | T | 0.00075042 | 18.8808769 | 4.08172296 | 3.73E-06 | 4.05E-06 |
| rs145228918 | 21 | 31124413 | GRIK1-AS1 | G | A | 0.00040064 | 27.9566355 | 6.04428254 | 3.74E-06 | 2.41E-06 |
| rs374460855 | 2 | 53108073 | MIR4431(dist=178320),ASB3(dist=789044) | G | C | 0.00038139 | 28.0264258 | 6.05954133 | 3.74E-06 | 2.41E-06 |
| rs767941193 | 7 | 117006606 | ASZ1 | T | G | 0.00038935 | -27.803729 | 6.01300961 | 3.77E-06 | 3.93E-07 |
| rs79343744 | 2 | 151551293 | LOC101929282(dist=59422),RBM43(dist=553435) | T | C | 0.00042227 | -26.307023 | 5.69119278 | 3.79E-06 | 1.89E-07 |
| rs17042683 | 2 | 52510537 | LOC730100 | A | G | 0.00036106 | 28.0107379 | 6.06357502 | 3.85E-06 | 2.41E-06 |
| rs537333512 | 2 | 52493569 | LOC730100 | G | T | 0.0003304 | 30.1269413 | 6.52423291 | 3.88E-06 | 2.41E-06 |
| rs28670976 | 3 | 105180707 | ALCAM | A | G | 0.68997053 | 1.11499452 | 0.24151734 | 3.90E-06 | 5.16E-06 |
| rs150202361 | 2 | 52704153 | LOC730100(dist=69098),MIR4431(dist=225507) | A | G | 0.0003676 | 27.9587052 | 6.05618146 | 3.90E-06 | 2.41E-06 |
| rs184772703 | 2 | 55797387 | PPP4R3B | A | G | 0.0003827 | 28.1363985 | 6.09563901 | 3.92E-06 | 2.41E-06 |
| rs78744226 | 2 | 39163630 | ARHGEF33 | A | G | 0.00071464 | 18.7179605 | 4.05660475 | 3.95E-06 | 4.05E-06 |
| rs906391415 | 5 | 173413458 | CPEB4(dist=25464),C5orf47(dist=2704) | G | A | 0.00037236 | 27.8871838 | 6.0472016 | 4.00E-06 | 2.41E-06 |
| rs144258224 | 2 | 39160588 | ARHGEF33 | G | T | 0.00071322 | 18.6966148 | 4.05674792 | 4.05E-06 | 4.05E-06 |
| rs79315504 | 2 | 39149289 | ARHGEF33 | T | C | 0.0007131 | 18.6945639 | 4.05674806 | 4.06E-06 | 4.05E-06 |
| rs79229305 | 2 | 39149783 | ARHGEF33 | C | T | 0.0007131 | 18.6945639 | 4.05674806 | 4.06E-06 | 4.05E-06 |
| rs79141371 | 2 | 39149911 | ARHGEF33 | A | C | 0.0007131 | 18.6945639 | 4.05674806 | 4.06E-06 | 4.05E-06 |

| | | | | | | | | | | |
|---|---|---|---|---|---|---|---|---|---|---|
| rs75939897 | 2 | 39161250 | ARHGEF33 | A | T | 0.0007131 | 18.6945639 | 4.05674806 | 4.06E-06 | 4.05E-06 |
| rs76593331 | 2 | 39163177 | ARHGEF33 | G | A | 0.0007131 | 18.6945639 | 4.05674806 | 4.06E-06 | 4.05E-06 |
| rs80270650 | 2 | 39163519 | ARHGEF33 | C | T | 0.0007131 | 18.6945639 | 4.05674806 | 4.06E-06 | 4.05E-06 |
| rs115901879 | 2 | 39171796 | ARHGEF33 | C | G | 0.0007131 | 18.6945639 | 4.05674806 | 4.06E-06 | 4.05E-06 |
| rs79778788 | 2 | 39145206 | MORN2(dist=35356),ARHGEF33(dist=1298) | G | A | 0.00071357 | 18.6941183 | 4.05674467 | 4.06E-06 | 4.05E-06 |
| rs73489616 | 19 | 450644 | SHC2 | G | A | 0.00037188 | 27.9225975 | 6.0598244 | 4.07E-06 | 2.41E-06 |
| rs114990055 | 2 | 39145286 | MORN2(dist=35436),ARHGEF33(dist=1218) | T | C | 0.00071488 | 18.6916846 | 4.05676182 | 4.07E-06 | 4.05E-06 |
| rs103911371 | 12 | 97055973 | CFAP54 | A | C | 0.00038602 | 27.9687217 | 6.07030835 | 4.08E-06 | 2.41E-06 |
| rs76723802 | 3 | 24482538 | THRB | T | C | 0.00087913 | 17.7928004 | 3.862213 | 4.09E-06 | 7.46E-06 |
| rs757283490 | 4 | 182401877 | LINC00290(dist=321575),TEMN3-AS1(dist=339281) | T | C | 0.00031519 | 31.5138696 | 6.84101314 | 4.09E-06 | 2.41E-06 |
| rs135440 | 22 | 44654739 | KIAA1644 | C | G | 0.88486249 | -1.6570614 | 0.35980577 | 4.12E-06 | 9.37E-06 |
| rs75888469 | 19 | 449822 | SHC2 | G | A | 0.00043261 | 27.7381098 | 6.02725509 | 4.18E-06 | 2.41E-06 |
| rs7816384 | 8 | 82512850 | FABP12(dist=69225),IMPA1P1(dist=3269) | A | G | 0.00035334 | -26.456813 | 5.75059445 | 4.21E-06 | 7.23E-06 |
| rs113676126 | 8 | 82507675 | FABP12(dist=64050),IMPA1P1(dist=8444) | A | G | 0.00035322 | -26.462727 | 5.75248849 | 4.22E-06 | 7.23E-06 |
| rs7972335 | 12 | 60144966 | SLC16A7 | C | T | 0.0004029 | 27.7649699 | 6.03722444 | 4.25E-06 | 2.41E-06 |
| rs542688940 | 12 | 59661267 | LRIG3(dist=346948),SLC16A7(dist=328554) | C | A | 0.000401 | 28.4095462 | 6.1827546 | 4.33E-06 | 2.41E-06 |
| rs112543146 | 2 | 52517531 | LOC730100 | T | G | 0.00036618 | 27.8541983 | 6.06220312 | 4.33E-06 | 2.41E-06 |
| rs533516434 | 21 | 38066225 | CLDN14(dist=117358),SIM2(dist=5196) | G | A | 0.00037081 | 27.8401802 | 6.06037285 | 4.35E-06 | 2.41E-06 |
| rs113113915 | 8 | 82677812 | CHMP4C(dist=6064),SNX16(dist=34006) | T | C | 0.00035881 | -25.941793 | 5.64720686 | 4.35E-06 | 7.23E-06 |
| rs779966965 | 9 | 120744155 | TLR4(dist=264386),BRINP1(dist=1184753) | A | G | 0.00037521 | 27.8108737 | 6.05550409 | 4.38E-06 | 2.41E-06 |
| rs74090138 | 14 | 105432498 | AHNAK2 | G | T | 0.00036843 | 26.0082657 | 5.66382667 | 4.39E-06 | 3.27E-06 |
| rs74090139 | 14 | 105432549 | AHNAK2 | C | T | 0.0003682 | 26.0063457 | 5.66416051 | 4.40E-06 | 3.27E-06 |
| rs533533423 | 8 | 32164376 | NRG1 | T | C | 0.0010706 | 16.3092555 | 3.55501962 | 4.48E-06 | 3.62E-06 |
| rs903168264 | 1 | 91913712 | HFM1(dist=43286),CDC7(dist=52692) | G | C | 0.00037687 | 28.0435295 | 6.11372878 | 4.50E-06 | 2.41E-06 |
| rs531720237 | 2 | 52703591 | LOC730100(dist=68536),MIR4431(dist=226069) | C | T | 0.00037295 | 27.7017464 | 6.04581513 | 4.61E-06 | 2.41E-06 |
| rs73285802 | 8 | 82661853 | CHMP4C | C | T | 0.00035786 | -25.960528 | 5.6687312 | 4.66E-06 | 7.23E-06 |
| rs7013201 | 8 | 82576401 | IMPA1 | C | A | 0.0003632 | -26.035979 | 5.68550146 | 4.66E-06 | 7.23E-06 |
| rs73287605 | 8 | 82662894 | CHMP4C | T | C | 0.00035845 | -25.948542 | 5.66658671 | 4.67E-06 | 7.23E-06 |
| rs61759787 | 14 | 105251426 | AKT1 | C | T | 0.00037366 | 25.9744327 | 5.67485635 | 4.71E-06 | 3.27E-06 |
| rs545222928 | 2 | 54641009 | C2orf73(dist=52295),SPTBN1(dist=42445) | G | A | 0.00042857 | 27.3592378 | 5.97837051 | 4.73E-06 | 2.41E-06 |
| rs369314523 | 2 | 52713930 | LOC730100(dist=78875),MIR4431(dist=215730) | T | C | 0.00037913 | 27.6032336 | 6.03176696 | 4.73E-06 | 2.41E-06 |
| rs75958227 | 11 | 116816591 | SIK3 | G | A | 0.00039351 | 27.6503549 | 6.04333722 | 4.75E-06 | 2.41E-06 |
| rs9775241 | 9 | 140189845 | TOR4A(dist=12752),NRARP(dist=4238) | T | G | 0.08617875 | 1.80490987 | 0.39468489 | 4.81E-06 | 2.23E-06 |
| rs59583530 | 8 | 82456832 | FABP12(dist=13207),IMPA1P1(dist=59287) | G | A | 0.00035631 | -26.122521 | 5.71278904 | 4.82E-06 | 7.23E-06 |
| rs79252187 | 11 | 36074245 | LDLRAD3 | C | T | 0.05562408 | -2.2210893 | 0.48574011 | 4.82E-06 | 3.09E-06 |
| rs138720810 | 16 | 72397879 | LINC01572 | G | A | 0.000198 | -38.536811 | 8.42924276 | 4.84E-06 | 1.25E-06 |
| rs73277219 | 8 | 82458109 | FABP12(dist=14484),IMPA1P1(dist=58010) | C | T | 0.00035643 | -26.10816 | 5.71089601 | 4.84E-06 | 7.23E-06 |
| rs542073652 | 1 | 104856346 | LOC100129138(dist=236653),LINC01676(dist=1275970) | T | G | 0.00043725 | 27.3290736 | 5.97829982 | 4.85E-06 | 2.41E-06 |
| rs59377511 | 8 | 82459205 | FABP12(dist=15580),IMPA1P1(dist=56914) | T | A | 0.00035655 | -26.105621 | 5.71090618 | 4.85E-06 | 7.23E-06 |
| rs142065631 | 14 | 105249989 | AKT1 | G | A | 0.00037152 | 25.969086 | 5.68163659 | 4.86E-06 | 3.27E-06 |
| rs73283084 | 8 | 82584902 | IMPA1 | C | T | 0.00035607 | -26.037416 | 5.69682444 | 4.87E-06 | 7.23E-06 |
| rs7727110 | 5 | 78095390 | ARSB | T | C | 0.73850927 | 1.15975356 | 0.25375 | 4.87E-06 | 3.98E-06 |
| rs73281042 | 8 | 82501372 | FABP12(dist=57747),IMPA1P1(dist=14747) | G | A | 0.00035691 | -26.016512 | 5.69381498 | 4.89E-06 | 7.23E-06 |
| rs73281040 | 8 | 82501044 | FABP12(dist=57419),IMPA1P1(dist=15075) | G | A | 0.00035714 | -26.0146 | 5.69381601 | 4.90E-06 | 7.23E-06 |
| rs75097308 | 8 | 82514241 | FABP12(dist=70616),IMPA1P1(dist=1878) | G | A | 0.00035691 | -26.005793 | 5.69192184 | 4.90E-06 | 7.23E-06 |
| rs73281051 | 8 | 82506749 | FABP12(dist=63124),IMPA1P1(dist=9370) | A | C | 0.00035702 | -26.00214 | 5.69192154 | 4.92E-06 | 7.23E-06 |
| rs58122686 | 8 | 82521885 | IMPA1P1 | G | T | 0.00035714 | -25.98427 | 5.68811346 | 4.92E-06 | 7.23E-06 |
| rs59290867 | 8 | 82527262 | IMPA1P1 | C | T | 0.00035691 | -25.984188 | 5.68811832 | 4.92E-06 | 7.23E-06 |
| rs553032007 | 6 | 93029880 | CASC6(dist=629734),EPHA7(dist=919860) | G | A | 0.00130532 | 14.7688263 | 3.23300956 | 4.92E-06 | 7.70E-06 |
| rs73281075 | 8 | 82531319 | IMPA1P1 | A | G | 0.00035679 | -25.982445 | 5.68811669 | 4.93E-06 | 7.23E-06 |
| rs73283059 | 8 | 82553731 | IMPA1P1(dist=10219),IMPA1(dist=15420) | T | G | 0.00035679 | -25.980871 | 5.6881099 | 4.93E-06 | 7.23E-06 |
| rs73281070 | 8 | 82527395 | IMPA1P1 | T | C | 0.00035667 | -25.980686 | 5.68811323 | 4.93E-06 | 7.23E-06 |

| | | | | | | | | | | |
|---|---|---|---|---|---|---|---|---|---|---|
| rs73283050 | 8 | 82544057 | IMPA1P1(dist=545) | A | G | 0.00035667 | -25.980686 | 5.68811323 | 4.93E-06 | 7.23E-06 |
| rs78567257 | 8 | 82606780 | SLC10A5 | A | T | 0.00035667 | -25.980139 | 5.68811477 | 4.94E-06 | 7.23E-06 |
| rs73283093 | 8 | 82592044 | IMPA1 | G | A | 0.00035726 | -25.979768 | 5.68806082 | 4.94E-06 | 7.23E-06 |
| rs73283079 | 8 | 82581574 | IMPA1 | C | T | 0.00035714 | -25.979666 | 5.6880732 | 4.94E-06 | 7.23E-06 |
| rs73283082 | 8 | 82582294 | IMPA1 | G | C | 0.00035714 | -25.979666 | 5.6880732 | 4.94E-06 | 7.23E-06 |
| rs61236631 | 8 | 82588662 | IMPA1 | G | C | 0.00035714 | -25.979666 | 5.6880732 | 4.94E-06 | 7.23E-06 |
| rs7013193 | 8 | 82576391 | IMPA1 | C | T | 0.00035702 | -25.979548 | 5.68808377 | 4.94E-06 | 7.23E-06 |
| rs73281034 | 8 | 82496066 | FABP12(dist=52441),IMPA1P1(dist=20053) | A | T | 0.0003575 | -26.014451 | 5.69572933 | 4.94E-06 | 7.23E-06 |
| rs7819477 | 8 | 82589317 | IMPA1 | G | A | 0.00035691 | -25.979414 | 5.68809253 | 4.94E-06 | 7.23E-06 |
| rs73283092 | 8 | 82591078 | IMPA1 | C | T | 0.00035691 | -25.979414 | 5.68809253 | 4.94E-06 | 7.23E-06 |
| rs73283066 | 8 | 82565407 | IMPA1P1(dist=21895),IMPA1(dist=3744) | T | A | 0.00035679 | -25.979263 | 5.68809948 | 4.94E-06 | 7.23E-06 |
| rs58717980 | 8 | 82573886 | IMPA1 | T | C | 0.00035679 | -25.979263 | 5.68809948 | 4.94E-06 | 7.23E-06 |
| rs73283077 | 8 | 82574734 | IMPA1 | C | A | 0.00035679 | -25.979263 | 5.68809948 | 4.94E-06 | 7.23E-06 |
| rs58765191 | 8 | 82556694 | IMPA1P1(dist=13182),IMPA1(dist=12457) | G | A | 0.00035667 | -25.979095 | 5.68810461 | 4.94E-06 | 7.23E-06 |
| rs73283071 | 8 | 82566605 | IMPA1P1(dist=23093),IMPA1(dist=2546) | T | G | 0.00035667 | -25.979095 | 5.68810461 | 4.94E-06 | 7.23E-06 |
| rs7823609 | 8 | 82567271 | IMPA1P1(dist=23759),IMPA1(dist=1880) | A | G | 0.00035667 | -25.979095 | 5.68810461 | 4.94E-06 | 7.23E-06 |
| rs73281076 | 8 | 82533560 | IMPA1P1 | C | T | 0.00035655 | -25.97891 | 5.68810794 | 4.94E-06 | 7.23E-06 |
| rs73281079 | 8 | 82533656 | IMPA1P1 | G | A | 0.00035655 | -25.97891 | 5.68810794 | 4.94E-06 | 7.23E-06 |
| rs73283045 | 8 | 82538023 | IMPA1P1 | G | T | 0.00035655 | -25.97891 | 5.68810794 | 4.94E-06 | 7.23E-06 |
| rs73283097 | 8 | 82598558 | IMPA1(NM_001144878:c.-4080>0,NM_005536:c.-47630>0 | G | T | 0.00035655 | -25.97891 | 5.68810794 | 4.94E-06 | 7.23E-06 |
| rs76392761 | 8 | 82598662 | IMPA1(dist=73) | G | A | 0.00035655 | -25.97891 | 5.68810794 | 4.94E-06 | 7.23E-06 |
| rs7830110 | 8 | 82629910 | ZFAND1 | T | C | 0.00035655 | -25.97891 | 5.68810794 | 4.94E-06 | 7.23E-06 |
| rs28436160 | 8 | 82632354 | ZFAND1 | T | C | 0.00035655 | -25.97891 | 5.68810794 | 4.94E-06 | 7.23E-06 |
| rs6984105 | 8 | 82634347 | ZFAND1(dist=808) | T | C | 0.00035655 | -25.97891 | 5.68810794 | 4.94E-06 | 7.23E-06 |
| rs10113801 | 8 | 82651596 | CHMP4C | C | T | 0.00035655 | -25.97891 | 5.68810794 | 4.94E-06 | 7.23E-06 |
| rs7813062 | 8 | 82679415 | CHMP4C(dist=7667),SNX16(dist=32403) | T | C | 0.00035655 | -25.97891 | 5.68810794 | 4.94E-06 | 7.23E-06 |
| rs60898990 | 8 | 82484344 | FABP12(dist=40719),IMPA1P1(dist=31775) | C | T | 0.0003575 | -26.021205 | 5.6975835 | 4.95E-06 | 7.23E-06 |
| rs113761827 | 8 | 82485914 | FABP12(dist=42289),IMPA1P1(dist=30205) | A | G | 0.0003575 | -26.021205 | 5.6975835 | 4.95E-06 | 7.23E-06 |
| rs61528282 | 8 | 82464804 | FABP12(dist=21179),IMPA1P1(dist=51315) | G | T | 0.00035726 | -26.053728 | 5.70523335 | 4.96E-06 | 7.23E-06 |
| rs73277233 | 8 | 82479913 | FABP12(dist=36288),IMPA1P1(dist=36206) | A | C | 0.00035762 | -26.025224 | 5.69948411 | 4.97E-06 | 7.23E-06 |
| rs111281207 | 8 | 82615846 | ZFAND1 | C | T | 0.00035797 | -25.972106 | 5.68810212 | 4.97E-06 | 7.23E-06 |
| rs74798895 | 8 | 82479044 | FABP12(dist=35419),IMPA1P1(dist=37075) | A | G | 0.00035774 | -26.021352 | 5.69949585 | 4.98E-06 | 7.23E-06 |
| rs115302759 | 6 | 91606574 | MAP3K7(dist=309554),MIR4643(dist=624804) | T | C | 0.0011885 | 14.1143765 | 3.09150805 | 4.98E-06 | 3.96E-06 |
| rs73277228 | 8 | 82463252 | FABP12(dist=19627),IMPA1P1(dist=52867) | C | T | 0.0003575 | -26.046099 | 5.7052152 | 4.99E-06 | 7.23E-06 |
| rs59175282 | 8 | 82690323 | CHMP4C(dist=18575),SNX16(dist=21495) | C | T | 0.00035726 | -25.951427 | 5.68812829 | 5.06E-06 | 7.23E-06 |
| rs191931350 | 6 | 91618301 | MAP3K7(dist=321281),MIR4643(dist=613077) | C | T | 0.0011885 | 14.0809509 | 3.08641934 | 5.06E-06 | 3.96E-06 |
| rs6920882 | 6 | 91618401 | MAP3K7(dist=321381),MIR4643(dist=612977) | T | C | 0.0011885 | 14.0809509 | 3.08641934 | 5.06E-06 | 3.96E-06 |
| rs187314727 | 8 | 109955617 | TMEM74(dist=155773),TRHR(dist=144036) | A | G | 0.00246981 | 11.0451111 | 2.4220091 | 5.11E-06 | 5.82E-06 |
| rs542586995 | 4 | 30886733 | PCDH7 | A | G | 0.00039696 | 25.5769091 | 5.61015548 | 5.14E-06 | 3.65E-06 |
| rs79053136 | 11 | 36076320 | LDLRAD3 | G | C | 0.05600226 | -2.2031499 | 0.48355099 | 5.21E-06 | 4.32E-06 |
| rs144082877 | 12 | 118374193 | KSR2 | A | G | 0.00090837 | 17.0689531 | 3.74941919 | 5.30E-06 | 4.30E-06 |
| rs574682246 | 12 | 63410783 | PPM1H(dist=82118),AVPR1A(dist=125756) | G | A | 0.00036843 | 27.5513069 | 6.05289984 | 5.32E-06 | 2.41E-06 |
| rs561792757 | 12 | 97348275 | NEDD1(dist=806) | G | C | 0.0003972 | 27.5281544 | 6.04985039 | 5.36E-06 | 2.41E-06 |
| rs567409192 | 2 | 52670336 | LOC730100(dist=35281),MIR4431(dist=259324) | C | A | 0.00038103 | 27.3963046 | 6.02686202 | 5.48E-06 | 2.41E-06 |
| rs562565330 | 12 | 127167561 | LOC100996671 | C | G | 0.00129914 | 13.6682539 | 3.00689701 | 5.48E-06 | 4.00E-06 |
| rs989602649 | 6 | 92642136 | CASC6(dist=241990),EPHA7(dist=1307604) | A | G | 0.00053197 | 23.1644651 | 5.09776531 | 5.52E-06 | 5.03E-06 |
| rs764394661 | 21 | 28775089 | ADAMTS5(dist=435650),LINC00113(dist=319609) | G | A | 0.00047778 | 23.6464293 | 5.20404598 | 5.52E-06 | 6.47E-06 |
| rs560366370 | 2 | 52669698 | LOC730100(dist=34643),MIR4431(dist=259962) | T | C | 0.00038151 | 27.379511 | 6.02684038 | 5.55E-06 | 2.41E-06 |
| rs758826592 | 21 | 28819633 | ADAMTS5(dist=480194),LINC00113(dist=275065) | C | T | 0.00048051 | 23.6406526 | 5.20393392 | 5.55E-06 | 6.47E-06 |
| rs6930752 | 6 | 101839863 | ASCC3(dist=510615),GRIK2(dist=6998) | C | A | 0.36982054 | -1.0498834 | 0.23115401 | 5.57E-06 | 3.36E-06 |
| rs147856887 | 11 | 105120267 | CARD18(dist=109806),GRIA4(dist=360533) | G | A | 0.00929368 | 5.26815124 | 1.16020741 | 5.61E-06 | 3.77E-06 |
| rs182795333 | 8 | 37436746 | LINC01605(dist=57842),ZNF703(dist=116523) | C | T | 0.00090005 | 17.1383879 | 3.77566107 | 5.65E-06 | 5.72E-06 |

| | | | | | | | | | | |
|---|---|---|---|---|---|---|---|---|---|---|
| rs17195227 | 6 | 132908951 | TAAR5(dist=780) | T | C | 0.32588543 | 1.08443573 | 0.23895086 | 5.67E-06 | 8.16E-06 |
| rs181970914 | 3 | 32430717 | CMTM8(dist=18900),CMTM7(dist=2446) | G | T | 0.00448419 | 8.17117815 | 1.80063269 | 5.68E-06 | 3.84E-06 |
| rs542193620 | 12 | 92394209 | LINC01619 | T | C | 0.00049156 | -22.048686 | 4.86345008 | 5.80E-06 | 5.39E-06 |
| rs191803878 | 12 | 85117921 | TMTC2(dist=1589274),SLC6A15(dist=135346) | G | A | 0.00047956 | 24.630283 | 5.43363951 | 5.82E-06 | 3.96E-06 |
| rs577715869 | 21 | 29023449 | ADAMTS5(dist=684010),LINC00113(dist=71249) | C | T | 0.00048479 | 23.5854072 | 5.20387631 | 5.84E-06 | 6.47E-06 |
| rs113622639 | 2 | 151575266 | LOC101929282(dist=83395),RBM43(dist=529462) | C | T | 0.00039933 | -27.596255 | 6.09133785 | 5.89E-06 | 1.89E-07 |
| rs189929690 | 18 | 2788624 | SMCHD1 | A | G | 0.00070062 | 18.6808488 | 4.12394351 | 5.90E-06 | 3.23E-06 |
| rs79012541 | 2 | 151539792 | LOC101929282(dist=47921),RBM43(dist=564936) | T | G | 0.00039006 | -27.893434 | 6.16242402 | 6.00E-06 | 1.89E-07 |
| rs181094857 | 18 | 2758416 | SMCHD1 | C | T | 0.00070383 | 18.5761161 | 4.10449937 | 6.02E-06 | 3.23E-06 |
| rs375792376 | 17 | 39602429 | KRT38(dist=4833),KRT32(dist=13336) | C | T | 0.00034966 | -27.353789 | 6.04487641 | 6.04E-06 | 3.42E-06 |
| rs105541165 | 9 | 117635668 | TNFSF15(dist=67260),TNFSF8(dist=19955) | C | A | 0.00042572 | 27.1788317 | 6.00699655 | 6.05E-06 | 2.41E-06 |
| rs771688511 | 21 | 28702859 | ADAMTS5(dist=363420),LINC00113(dist=391839) | A | C | 0.00048205 | 23.5562903 | 5.20814207 | 6.10E-06 | 6.47E-06 |
| rs143819095 | 1 | 104751586 | LOC100129138(dist=131893),LINC01676(dist=1380730) | A | C | 0.00030497 | 30.7427153 | 6.79951941 | 6.15E-06 | 2.41E-06 |
| rs184927009 | 17 | 47245198 | B4GALNT2 | C | G | 0.00276836 | -10.233304 | 2.26430828 | 6.20E-06 | 1.15E-06 |
| rs115689294 | 17 | 30047103 | MIR365B(dist=144563),COPRS(dist=131781) | A | C | 0.00041229 | 27.0209892 | 5.98031672 | 6.23E-06 | 2.41E-06 |
| rs13282027 | 8 | 25004229 | NEFL(dist=189846),DOCK5(dist=38009) | G | A | 0.33414595 | -1.050901 | 0.23274203 | 6.32E-06 | 6.73E-06 |
| rs143690756 | 9 | 112065921 | EPB41L4B | C | A | 0.0092112 | -5.3347649 | 1.18149312 | 6.32E-06 | 3.79E-06 |
| rs775045363 | 9 | 106738027 | LINC01492(dist=650712),LOC101928523(dist=23963) | G | A | 0.00015462 | -38.067645 | 8.43713254 | 6.42E-06 | 4.05E-06 |
| rs549701042 | 4 | 154740186 | SFRP2(dist=29958),DCHS2(dist=415341) | C | T | 0.00205871 | 11.0603714 | 2.45180546 | 6.45E-06 | 3.77E-06 |
| rs72885535 | 2 | 52497133 | LOC730100 | A | G | 0.00036772 | 28.6255624 | 6.3475077 | 6.49E-06 | 2.41E-06 |
| rs982243669 | 9 | 97446673 | FBP1(dist=44142),C9orf3(dist=42278) | G | A | 0.00073877 | 19.1455338 | 4.25071803 | 6.67E-06 | 3.73E-06 |
| rs12527779 | 6 | 132916548 | TAAR5(dist=5671),TAAR3P(dist=12816) | G | A | 0.32471203 | 1.07705362 | 0.239187 | 6.70E-06 | 9.99E-06 |
| rs12527780 | 6 | 132916550 | TAAR5(dist=5673),TAAR3P(dist=12814) | G | C | 0.32471821 | 1.07674318 | 0.23918696 | 6.74E-06 | 9.99E-06 |
| rs73825624 | 3 | 24400340 | THRB | T | C | 0.00381994 | 8.06500345 | 1.79182616 | 6.76E-06 | 7.63E-06 |
| rs546540293 | 17 | 54482918 | ANKFN1 | T | A | 0.00142608 | 14.8698237 | 3.30523683 | 6.83E-06 | 7.25E-06 |
| rs55634320 | 10 | 100145307 | PYROXD2 | C | T | 0.00030295 | 30.3377177 | 6.75153251 | 7.01E-06 | 2.41E-06 |
| rs183396256 | 5 | 44910440 | MRPS30(dist=94822),HCN1(dist=344612) | G | A | 0.00034395 | -27.42318 | 6.10317017 | 7.01E-06 | 5.04E-06 |
| rs541969988 | 6 | 94436294 | TSG1 | G | A | 0.00079665 | 17.6996968 | 3.93969988 | 7.03E-06 | 3.36E-06 |
| rs13377557 | 11 | 36077902 | LDLRAD3 | T | G | 0.05574507 | -2.1794325 | 0.48514651 | 7.05E-06 | 4.52E-06 |
| rs372988655 | 6 | 69675725 | ADGRB3 | C | G | 0.00042441 | 24.3063915 | 5.4136995 | 7.13E-06 | 7.72E-07 |
| rs923947970 | 6 | 93232328 | CASC6(dist=832182),EPHA7(dist=717412) | C | T | 0.00048514 | 23.7581451 | 5.29444757 | 7.21E-06 | 5.03E-06 |
| rs537236396 | 2 | 59298287 | LINC01122(dist=7386),LINC01793(dist=146556) | G | A | 0.00042334 | 26.5458278 | 5.92047376 | 7.33E-06 | 2.41E-06 |
| rs9841732 | 3 | 24698792 | MIR4792(dist=135866),RARB(dist=172022) | A | G | 0.00071322 | 18.3047362 | 4.08527128 | 7.44E-06 | 6.40E-06 |
| rs9877614 | 3 | 24703565 | MIR4792(dist=140639),RARB(dist=167249) | C | G | 0.00071322 | 18.3047362 | 4.08527128 | 7.44E-06 | 6.40E-06 |
| rs144954311 | 5 | 125358734 | LINC02240(dist=420814),LINC02039(dist=156536) | T | C | 0.00014226 | -39.437075 | 8.80198639 | 7.45E-06 | 4.05E-06 |
| rs9834142 | 3 | 24702416 | MIR4792(dist=139490),RARB(dist=168398) | A | G | 0.00071333 | 18.3027824 | 4.0852734 | 7.46E-06 | 6.40E-06 |
| rs9812320 | 3 | 24680869 | MIR4792(dist=117943),RARB(dist=189945) | G | A | 0.0007131 | 18.2980604 | 4.08426482 | 7.46E-06 | 6.40E-06 |
| rs9853786 | 3 | 24688359 | MIR4792(dist=125433),RARB(dist=182455) | C | T | 0.0007131 | 18.2980604 | 4.08426482 | 7.46E-06 | 6.40E-06 |
| rs542595676 | 7 | 42783162 | LINC01448(dist=37116),C7orf25(dist=165710) | G | A | 0.00137984 | 13.5469194 | 3.02417493 | 7.48E-06 | 6.58E-06 |
| rs9843065 | 3 | 24707247 | MIR4792(dist=144321),RARB(dist=163567) | T | C | 0.00071322 | 18.3030452 | 4.08602845 | 7.48E-06 | 6.40E-06 |
| rs531164098 | 6 | 93746493 | CASC6(dist=1346347),EPHA7(dist=203247) | A | C | 0.00049073 | 20.9304237 | 4.67491767 | 7.56E-06 | 6.66E-06 |
| rs138073410 | 7 | 101820529 | CUX1 | A | G | 0.0297503 | 2.97163805 | 0.66415833 | 7.67E-06 | 6.41E-06 |
| rs149524901 | 2 | 52545819 | LOC730100 | G | A | 0.00048681 | 25.5037335 | 5.70115362 | 7.70E-06 | 2.41E-06 |
| rs138014184 | 10 | 98524231 | PIK3AP1(dist=43952),MIR607(dist=64195) | C | G | 0.00061208 | 20.1573842 | 4.50808901 | 7.77E-06 | 7.69E-06 |
| rs13250561 | 8 | 24991003 | NEFL(dist=176620),DOCK5(dist=51235) | T | C | 0.33245353 | -1.0431891 | 0.23330872 | 7.78E-06 | 7.41E-06 |
| rs141588875 | 9 | 112054565 | EPB41L4B | C | T | 0.00924935 | -5.2432559 | 1.17334345 | 7.87E-06 | 3.79E-06 |
| rs967509050 | 6 | 93192137 | CASC6(dist=791991),EPHA7(dist=757603) | G | A | 0.00061196 | 19.1761802 | 4.29209386 | 7.90E-06 | 5.91E-06 |
| rs55999886 | 3 | 24692886 | MIR4792(dist=129960),RARB(dist=177928) | G | A | 0.00073176 | 18.2378279 | 4.08376768 | 7.97E-06 | 6.40E-06 |
| rs76967133 | 10 | 100203666 | HPS1 | T | C | 0.0004357 | 26.6687016 | 5.97307894 | 8.01E-06 | 2.41E-06 |
| rs55723486 | 3 | 24671717 | MIR4792(dist=108791),RARB(dist=199097) | G | C | 0.0007194 | 18.2205882 | 4.08384464 | 8.13E-06 | 6.40E-06 |
| rs9861176 | 3 | 24670141 | MIR4792(dist=107215),RARB(dist=200673) | C | G | 0.00073092 | 18.2087282 | 4.08134971 | 8.14E-06 | 6.40E-06 |
| rs9834456 | 3 | 24672478 | MIR4792(dist=109552),RARB(dist=198336) | A | G | 0.00071702 | 18.2186603 | 4.08393127 | 8.16E-06 | 6.40E-06 |

| | | | | | | | | | | |
|---|---|---|---|---|---|---|---|---|---|---|
| rs545970707 | 12 | 84828222 | TMTC2(dist=1299575),SLC6A15(dist=425045) | A | G | 0.00049774 | 24.1822932 | 5.42351244 | 8.24E-06 | 3.96E-06 |
| rs7637378 | 3 | 24665688 | MIR4792(dist=102762),RARB(dist=205126) | T | A | 0.00074329 | 18.1915533 | 4.08070996 | 8.28E-06 | 6.40E-06 |
| rs536590440 | 6 | 93377751 | CASC6(dist=977605),EPHA7(dist=571989) | T | C | 0.00134217 | 14.3154598 | 3.21159491 | 8.29E-06 | 7.70E-06 |
| rs369748695 | 21 | 28315900 | ADAMTS5 | C | T | 0.00054279 | 23.1380941 | 5.19409758 | 8.40E-06 | 6.47E-06 |
| rs191814814 | 7 | 43728486 | COA1 | T | C | 0.00543725 | 7.26311643 | 1.63079759 | 8.44E-06 | 8.07E-06 |
| rs6791206 | 3 | 61236328 | FHIT | A | G | 0.27760245 | -1.0957055 | 0.24619996 | 8.57E-06 | 8.28E-06 |
| rs574282721 | 17 | 54544365 | ANKFN1 | C | A | 0.0013184 | 15.404332 | 3.4695448 | 9.00E-06 | 9.69E-06 |
| rs376021354 | 2 | 53056823 | MIR4431(dist=127070),ASB3(dist=840294) | G | A | 0.00043546 | 25.8149352 | 5.81949722 | 9.17E-06 | 2.41E-06 |
| rs922706564 | 12 | 22090721 | ABCC9(dist=1093),CMAS(dist=108387) | C | A | 0.00049727 | 21.0650716 | 4.75542337 | 9.44E-06 | 8.86E-06 |
| rs59845200 | 8 | 82456271 | FABP12(dist=12646),IMPA1P1(dist=59848) | G | A | 0.0004445 | -24.402511 | 5.51528666 | 9.67E-06 | 7.23E-06 |
| rs116381596 | 14 | 105434356 | AHNAK2 | T | A | 0.00038899 | 24.7889966 | 5.62181512 | 1.04E-05 | 3.27E-06 |
| rs112782431 | 19 | 450916 | SHC2 | T | C | 0.00039767 | 26.5472642 | 6.02101288 | 1.04E-05 | 2.41E-06 |
| rs74090142 | 14 | 105434357 | AHNAK2 | T | C | 0.00038876 | 24.7851545 | 5.62193617 | 1.04E-05 | 3.27E-06 |
| rs373065827 | 17 | 39362922 | KRTAP9-1(dist=16031),KRTAP9-2(dist=19978) | G | A | 0.00037295 | -27.317475 | 6.20454844 | 1.07E-05 | 3.42E-06 |
| rs556906727 | 21 | 28894113 | ADAMTS5(dist=554674),LINC00113(dist=200585) | A | G | 0.00059674 | 22.4874383 | 5.10956632 | 1.08E-05 | 6.47E-06 |
| rs113890693 | 15 | 65328067 | MTFMT(dist=6090),SLC51B(dist=9641) | C | T | 0.02973936 | -2.9551376 | 0.67186149 | 1.09E-05 | 6.57E-06 |
| rs776691922 | 2 | 169243036 | STK39(dist=138931),CERS6(dist=69723) | A | G | 0.00221452 | 11.233049 | 2.55418871 | 1.09E-05 | 3.78E-06 |
| rs6572707 | 14 | 51358831 | ABHD12B | A | C | 0.99960946 | -25.266496 | 5.75047106 | 1.11E-05 | 8.69E-06 |
| rs74090141 | 14 | 105433831 | AHNAK2 | G | A | 0.00039375 | 24.6368164 | 5.60963333 | 1.12E-05 | 3.27E-06 |
| rs527956574 | 9 | 37465399 | ZBTB5(NM_014872:c.-228510>0) | C | A | 0.0012541 | -15.37407 | 3.50714872 | 1.17E-05 | 5.81E-06 |
| rs376675751 | 17 | 39532266 | KRT33B(dist=6214),KRT34(dist=1655) | G | A | 0.00034086 | -27.524608 | 6.30144515 | 1.25E-05 | 3.42E-06 |
| rs73225625 | 12 | 95598185 | FGD6 | C | T | 0.00124994 | 14.7409076 | 3.37576889 | 1.26E-05 | 3.72E-06 |
| rs73875514 | 4 | 176126989 | ADAM29(dist=227658),GPM6A(dist=427099) | C | G | 0.10451842 | -1.6957521 | 0.38859716 | 1.28E-05 | 8.22E-06 |
| rs12210924 | 6 | 85047569 | CEP162(dist=110216),LINC01611(dist=83182) | T | A | 0.00014678 | -42.20963 | 9.67393293 | 1.28E-05 | 5.56E-06 |
| rs112855118 | 14 | 105475614 | CDCA4(dist=296) | G | A | 0.00060661 | 20.9251714 | 4.79714096 | 1.29E-05 | 4.61E-06 |
| rs78843663 | 13 | 58778233 | LINC02338 | T | C | 0.09885417 | -1.6491634 | 0.37880304 | 1.34E-05 | 9.08E-06 |
| rs572048141 | 2 | 52593682 | LOC730100 | G | A | 0.00039815 | 26.2520137 | 6.03047099 | 1.34E-05 | 2.41E-06 |
| rs7032003 | 9 | 140194020 | NRARP(dist=63) | G | A | 0.07282291 | 1.84496945 | 0.42424983 | 1.37E-05 | 4.56E-06 |
| rs11882868 | 19 | 553179 | GZMM(dist=3259),BSG(dist=18098) | A | G | 0.00044961 | 26.1015086 | 6.00417664 | 1.38E-05 | 2.41E-06 |
| rs9922401 | 16 | 17620512 | XYLT1(dist=55774),NPIPA8(dist=791264) | G | T | 0.80929487 | 1.27123824 | 0.29243413 | 1.38E-05 | 3.30E-06 |
| rs100691141 | 1 | 224785733 | CNIH3 | G | T | 0.00014107 | -41.979115 | 9.68342368 | 1.46E-05 | 5.56E-06 |
| rs752153212 | 1 | 23924094 | ID3(dist=37809),MDS2(dist=29730) | T | C | 0.00037414 | 24.9984162 | 5.77013029 | 1.48E-05 | 8.95E-06 |
| rs140457075 | 10 | 98543172 | PIK3AP1(dist=62893),MIR607(dist=45254) | A | G | 0.00062075 | 19.502499 | 4.50282799 | 1.48E-05 | 7.69E-06 |
| rs2043080 | 2 | 3726163 | ALLC | T | C | 0.47976194 | 0.97218491 | 0.22454068 | 1.49E-05 | 7.17E-06 |
| rs916505306 | 10 | 98711486 | LCOR | A | G | 0.00063787 | 19.4636552 | 4.49649796 | 1.50E-05 | 7.69E-06 |
| rs100144516 | 10 | 98672709 | LCOR | T | G | 0.0006381 | 19.46075 | 4.49648889 | 1.50E-05 | 7.69E-06 |
| rs111567494 | 2 | 52518266 | LOC730100 | G | A | 0.00039351 | 25.9879746 | 6.01671413 | 1.57E-05 | 2.41E-06 |
| rs9922912 | 16 | 17620367 | XYLT1(dist=55629),NPIPA8(dist=791409) | C | T | 0.80919028 | 1.26060788 | 0.29227424 | 1.61E-05 | 3.59E-06 |
| rs112607013 | 19 | 451814 | SHC2 | G | C | 0.00073758 | 17.2426724 | 3.9994429 | 1.62E-05 | 8.78E-06 |
| rs190160667 | 3 | 111968572 | SLC9C1 | C | T | 0.00012515 | -43.47856 | 10.0994357 | 1.67E-05 | 5.56E-06 |
| rs956579791 | 11 | 117012983 | SIK3(dist=43852),PAFAH1B2(dist=2017) | T | C | 0.00039708 | 27.1055973 | 6.29753109 | 1.68E-05 | 2.41E-06 |
| rs191156346 | 3 | 111864399 | SLC9C1 | T | A | 0.00012681 | -41.614255 | 9.68534099 | 1.73E-05 | 5.56E-06 |
| rs116880183 | 18 | 57503119 | CCBE1(dist=138475),PMAIP1(dist=64073) | T | C | 0.01592073 | -4.1554937 | 0.96727882 | 1.74E-05 | 7.89E-06 |
| rs140521674 | 3 | 112734372 | NEPRO | A | C | 0.00012693 | -41.569882 | 9.6767655 | 1.74E-05 | 5.56E-06 |
| rs186896969 | 3 | 111887047 | SLC9C1 | G | A | 0.00012907 | -41.682919 | 9.71140222 | 1.77E-05 | 5.56E-06 |
| rs11879663 | 19 | 553147 | GZMM(dist=3227),BSG(dist=18130) | G | C | 0.00044462 | 25.7681533 | 6.00366734 | 1.77E-05 | 2.41E-06 |
| rs752935043 | 2 | 50573025 | NRXN1 | G | A | 0.00012467 | -41.550981 | 9.68232968 | 1.78E-05 | 5.56E-06 |
| rs577497738 | 8 | 32248041 | NRG1 | C | T | 0.00118434 | 15.1282904 | 3.52562743 | 1.78E-05 | 2.43E-06 |
| rs767370029 | 6 | 86041364 | TBX18(dist=567410),LOC101928820(dist=55573) | G | A | 0.0001198 | -41.510188 | 9.68763608 | 1.83E-05 | 5.56E-06 |
| rs571839631 | 6 | 53847045 | LOC101927189 | G | A | 0.00012444 | -41.492752 | 9.68570031 | 1.84E-05 | 5.56E-06 |
| rs148542272 | 2 | 178886534 | PDE11A | G | A | 0.00012657 | -41.715775 | 9.73853587 | 1.84E-05 | 5.56E-06 |
| rs144721913 | 3 | 112694576 | CD200R1(dist=639) | C | T | 0.00011944 | -41.496135 | 9.68769198 | 1.84E-05 | 5.56E-06 |

| | | | | | | | | | | |
|---|---|---|---|---|---|---|---|---|---|---|
| rs577959748 | 6 | 54037801 | MLIP | G | T | 0.00012313 | -41.490651 | 9.68762382 | 1.85E-05 | 5.56E-06 |
| rs916860201 | 6 | 53400634 | GCLC | T | C | 0.00050856 | 21.8343533 | 5.09934298 | 1.85E-05 | 9.71E-06 |
| rs118150476 | 10 | 92050053 | LINC01375(dist=332923),LOC101926942(dist=112225) | G | C | 0.00649548 | 6.49073398 | 1.5158957 | 1.85E-05 | 9.75E-06 |
| rs560150995 | 3 | 111634811 | PHLDB2 | G | T | 0.00011909 | -41.513502 | 9.69744252 | 1.86E-05 | 5.56E-06 |
| rs112883847 | 8 | 98883322 | MATN2 | C | T | 0.00030223 | -27.993953 | 6.54273779 | 1.88E-05 | 1.25E-06 |
| rs548971166 | 6 | 54235168 | TINAG | G | A | 0.00012432 | -41.411689 | 9.68420871 | 1.90E-05 | 5.56E-06 |
| rs568263181 | 2 | 179128707 | OSBPL6 | G | C | 0.00012646 | -41.655737 | 9.74228232 | 1.90E-05 | 5.56E-06 |
| rs137952554 | 3 | 114326090 | ZBTB20 | T | A | 0.00011338 | -45.759169 | 10.703721 | 1.91E-05 | 5.56E-06 |
| rs12197596 | 6 | 85165764 | LINC01611 | T | G | 0.00012717 | -41.744902 | 9.76482224 | 1.91E-05 | 5.56E-06 |
| rs540321647 | 2 | 178821554 | PDE11A | G | A | 0.00013846 | -41.282025 | 9.65669092 | 1.91E-05 | 5.56E-06 |
| rs919235896 | 12 | 84150281 | TMTC2(dist=621634),SLC6A15(dist=1102986) | T | C | 0.0006242 | 19.7919119 | 4.63391549 | 1.95E-05 | 8.62E-06 |
| rs577523040 | 6 | 54180894 | TINAG | G | T | 0.00012146 | -41.362277 | 9.68773908 | 1.96E-05 | 5.56E-06 |
| rs371400593 | 17 | 39105378 | KRT23(dist=11483),KRT39(dist=9291) | G | A | 0.00038068 | -26.423735 | 6.19573597 | 2.00E-05 | 3.42E-06 |
| rs765595000 | 2 | 50992999 | NRXN1 | G | C | 0.00012396 | -41.310317 | 9.68633173 | 2.00E-05 | 5.56E-06 |
| rs759367237 | 12 | 84138595 | TMTC2(dist=609948),SLC6A15(dist=1114672) | T | C | 0.00062871 | 19.7474301 | 4.63162498 | 2.01E-05 | 8.62E-06 |
| rs540399461 | 6 | 54264985 | TINAG(dist=10035),FAM83B(dist=446584) | A | G | 0.00013739 | -41.206719 | 9.66938949 | 2.03E-05 | 5.56E-06 |
| rs545192919 | 3 | 114435641 | ZBTB20 | C | T | 0.00012895 | -43.86464 | 10.2938217 | 2.03E-05 | 5.56E-06 |
| rs774337449 | 22 | 46881458 | CELSR1 | C | T | 0.00259056 | 9.68924636 | 2.27580892 | 2.07E-05 | 8.57E-06 |
| rs117235192 | 3 | 111687090 | PHLDB2 | T | G | 0.00012919 | -41.308894 | 9.70383 | 2.07E-05 | 5.56E-06 |
| rs186567835 | 14 | 81218210 | CEP128 | G | T | 0.00042441 | 23.985388 | 5.63564062 | 2.08E-05 | 4.64E-06 |
| rs572580183 | 1 | 225192602 | DNAH14 | G | A | 0.00013085 | -41.173082 | 9.67928945 | 2.10E-05 | 5.56E-06 |
| rs185843032 | 8 | 76277408 | CASC9(dist=86284),HNF4G(dist=42497) | C | T | 0.00335286 | -8.6080166 | 2.03394151 | 2.31E-05 | 4.02E-06 |
| rs117667963 | 9 | 110367720 | KLF4(dist=115719),ACTL7B(dist=1249149) | G | A | 0.17713489 | -1.250026 | 0.29598821 | 2.41E-05 | 9.53E-06 |
| rs188881668 | 11 | 123243276 | CLMP(dist=177263),MIR4493(dist=8872) | C | T | 0.00013941 | -40.805766 | 9.67518259 | 2.47E-05 | 5.56E-06 |
| rs185494985 | 8 | 96224286 | LINC01298 | A | C | 0.00043428 | 21.8300756 | 5.17685753 | 2.48E-05 | 3.96E-06 |
| rs746863731 | 6 | 54713225 | FAM83B | C | A | 0.00051117 | 21.3706306 | 5.07681347 | 2.56E-05 | 9.71E-06 |
| rs116542186 | 21 | 19097232 | BTG3(dist=111964),C21orf91-OT1(dist=52489) | G | A | 0.00028215 | -30.47523 | 7.26062664 | 2.70E-05 | 1.25E-06 |
| rs1420056 | 16 | 6582955 | RBFOX1 | C | A | 0.00158153 | 13.0871873 | 3.12099886 | 2.75E-05 | 3.85E-06 |
| rs369014432 | 17 | 39322980 | KRTAP4-3(dist=503) | T | C | 0.00033575 | -26.982563 | 6.47672551 | 3.10E-05 | 3.42E-06 |
| rs36183315 | 16 | 17610970 | XYLT1(dist=46232),NPIPA8(dist=800806) | G | A | 0.17886974 | -1.240368 | 0.2978357 | 3.12E-05 | 7.03E-06 |
| rs185994878 | 5 | 169872037 | KCNIP1 | T | A | 0.00048574 | 22.1880491 | 5.33539579 | 3.20E-05 | 9.97E-06 |
| rs117342505 | 7 | 49868904 | VWC2 | C | T | 0.01269586 | 4.41628485 | 1.06273583 | 3.24E-05 | 1.77E-06 |
| rs534238226 | 5 | 53681116 | LINC01033 | T | C | 0.00031638 | -26.987941 | 6.49728888 | 3.27E-05 | 1.25E-06 |
| rs770774315 | 1 | 105609809 | LOC100129138(dist=990116),LINC01676(dist=522507) | G | A | 0.00030913 | -27.47996 | 6.63484716 | 3.45E-05 | 1.25E-06 |
| rs867847786 | 15 | 60565694 | FOXB1(dist=267552),ANXA2(dist=73656) | C | T | 0.00065189 | 18.7008105 | 4.53640003 | 3.75E-05 | 4.54E-06 |
| rs758402763 | 12 | 87488732 | MGAT4C(dist=255949),LOC105369879(dist=235784) | A | C | 0.0005536 | 21.7431236 | 5.28138715 | 3.84E-05 | 3.96E-06 |
| rs11694862 | 2 | 42053055 | SLC8A1(dist=1313480),LINC01913(dist=51640) | A | G | 0.01485251 | 4.04474748 | 0.98893437 | 4.31E-05 | 7.15E-06 |
| rs557547882 | 6 | 53396939 | GCLC | G | C | 0.00057179 | 19.9738067 | 4.88522198 | 4.34E-05 | 9.71E-06 |
| rs567855665 | 9 | 86225665 | FRMD3(dist=72317),IDNK(dist=12299) | G | A | 0.00063977 | -20.737196 | 5.0777313 | 4.43E-05 | 8.60E-06 |
| rs143720809 | 13 | 24102126 | LINC00327(dist=40523),TNFRSF19(dist=42383) | G | A | 0.00070193 | 18.3234143 | 4.51711256 | 4.98E-05 | 5.53E-06 |
| rs73316791 | 10 | 98542203 | PIK3AP1(dist=61924),MIR607(dist=46223) | A | C | 0.0009376 | 15.1444567 | 3.7490487 | 5.36E-05 | 9.15E-06 |
| rs553963075 | 3 | 165040982 | LINC01322 | A | G | 0.00059318 | 18.7233514 | 4.96402992 | 0.00016207 | 8.74E-06 |
