## Supplemental_table3 for "Rare Genetic Variants Correlate with Better Processing Speed"

**Supplement Table3. SNPs replicated in the Danish combined cohort.**

Chr, Chromosome; pos, position of SNP on GRCh37 reference panel; n.obs, number of observations; caf, coding allele frequency; maf, minor allele frequency; INFO_Dkdata, imputation quality R-sq

| SNP INFO | | | | | DISCOVERY RESULTS | | | | | | | | | 1_SNPs | | | Replication RESULTS DK twins | | | | |
|---|---|---|---|---|---|---|---|---|---|---|---|---|---|---|---|---|---|---|---|---|---|
| **ExonicFunction** | **Gene** | **chr** | **pos** | **rsID** | **n.obs** | **Est** | **SE** | **Wald.Stat** | **Wald.pval** | **INFO** | **caf** | **maf** | **in_DKdata** | **INFO_Dkdata** | **replication** | **n.obs** | **maf/caf** | **Est** | **Est.SE** | **t** | **p-val** |
| intergenic | MIR4792(dist=150243),RARB(dist=157645) | 3 | 24713169 | rs7623455 | 4207 | 29.92759 | 5.15449 | 5.80612 | 6.39E-09 | IMPUTED | 0.000562 | 0.000562 | Yes, imputed | 0.470 | Yes | 1577 | 0.00030 | 10.50 | 1.20 | 8.76 | 9.08e-18 |
| intergenic | MIR4792(dist=149406),RARB(dist=158482) | 3 | 24712332 | rs9821776 | 4207 | 29.92557 | 5.15449 | 5.805727 | 6.41E-09 | IMPUTED | 0.000562 | 0.000562 | Yes, imputed | 0.396 | Yes | 1577 | 0.00040 | 9.50 | 2.15 | 4.42 | 0.000011 |
| intergenic | MIR4792(dist=149263),RARB(dist=158625) | 3 | 24712189 | rs9821587 | 4207 | 29.93104 | 5.155491 | 5.805663 | 6.41E-09 | IMPUTED | 0.000562 | 0.000562 | Yes, imputed | 0.400 | Yes | 1577 | 0.00040 | 9.48 | 2.12 | 4.48 | 0.000009 |
| intronic | THRB | 3 | 24423893 | rs75963215 | 4207 | 20.96046 | 3.799005 | 5.517355 | 3.44E-08 | IMPUTED | 0.000951 | 0.000951 | Yes, imputed | 0.308 | Yes | 1577 | 0.00085 | 8.17 | 3.72 | 2.22 | 0.028 |
| intronic | THRB | 3 | 24424415 | rs59914825 | 4207 | 20.96043 | 3.799012 | 5.517337 | 3.44E-08 | GENOTYPE | 0.000951 | 0.000951 | Yes, imputed | 0.306 | Yes | 1577 | 0.00085 | 8.28 | 3.73 | 2.22 | 0.027 |
| intronic | THRB | 3 | 24425760 | rs4266131 | 4207 | -20.8765 | 3.789505 | -5.50902 | 3.61E-08 | IMPUTED | 0.999028 | 0.000972 | Yes, imputed | 0.278 | No, INFO<0,3 | | | | | | |
| intergenic | FABP5(dist=153906),PMP2(dist=1645) | 8 | 82350918 | rs58169119 | 4207 | -28.2157 | 5.127586 | -5.50273 | 3.74E-08 | IMPUTED | 0.000471 | 0.000471 | Yes, imputed | 0.239 | No, INFO<0,3 | | | | | | |
| intronic | RARB | 3 | 25100109 | rs148022846 | 4207 | 30.55891 | 5.554728 | 5.501423 | 3.77E-08 | IMPUTED | 0.000482 | 0.000482 | Yes, imputed | 0.061 | No, INFO<0,3 | | | | | | |
| intergenic | PSAT1(dist=709376),LOC101927450(dist=95953) | 9 | 81654385 | rs146299120 | 4207 | -53.0939 | 9.674549 | -5.488 | 4.07E-08 | IMPUTED | 0.000148 | 0.000148 | Yes, imputed | 0.000 | No, INFO<0,3 | | | | | | |
| intronic | DCC | 18 | 50217262 | NA | 4207 | -51.9915 | 9.479026 | -5.4849 | 4.14E-08 | IMPUTED | 0.000163 | 0.000163 | No | NA | NA | | | | | | |
| intronic | RARB | 3 | 25228798 | rs189337466 | 4207 | 30.37702 | 5.558595 | 5.464873 | 4.63E-08 | IMPUTED | 0.000475 | 0.000475 | Yes, imputed | 0.212 | No, INFO<0,3 | | | | | | |
| intronic | RARB | 3 | 25170804 | rs78704059 | 4207 | 30.34065 | 5.552305 | 5.464514 | 4.64E-08 | IMPUTED | 0.000476 | 0.000476 | Yes, imputed | 0.474 | Yes | 1577 | 0.00114 | 12.93 | 3.58 | 3.62 | 0.0003 |
| intronic | RARB | 3 | 25206678 | rs59296535 | 4207 | 30.33341 | 5.552222 | 5.463292 | 4.67E-08 | IMPUTED | 0.000477 | 0.000477 | Yes, imputed | 0.183 | No, INFO<0,3 | | | | | | |
| intronic | RARB | 3 | 25206522 | rs74467766 | 4207 | 30.33037 | 5.552221 | 5.462745 | 4.69E-08 | IMPUTED | 0.000477 | 0.000477 | Yes, imputed | 0.177 | No, INFO<0,3 | | | | | | |
| intronic | RARB | 3 | 25194665 | rs80011850 | 4207 | 30.32872 | 5.552198 | 5.462471 | 4.70E-08 | IMPUTED | 0.000477 | 0.000477 | Yes, imputed | 0.151 | No, INFO<0,3 | | | | | | |
| intronic | RARB | 3 | 25223736 | rs77306558 | 4207 | 30.33787 | 5.553928 | 5.462416 | 4.70E-08 | IMPUTED | 0.000477 | 0.000477 | Yes, imputed | 0.181 | No, INFO<0,3 | | | | | | |
| intronic | RARB | 3 | 25217506 | rs79046847 | 4207 | 30.32667 | 5.552236 | 5.462064 | 4.71E-08 | IMPUTED | 0.000477 | 0.000477 | Yes, imputed | 0.175 | No, INFO<0,3 | | | | | | |
| intronic | RARB | 3 | 25192336 | rs77025115 | 4207 | 30.32603 | 5.552182 | 5.462003 | 4.71E-08 | IMPUTED | 0.000478 | 0.000478 | Yes, imputed | 0.254 | No, INFO<0,3 | | | | | | |
| intronic | RARB | 3 | 25212666 | rs80164536 | 4207 | 30.32529 | 5.552225 | 5.461828 | 4.71E-08 | IMPUTED | 0.000477 | 0.000477 | Yes, imputed | 0.174 | No, INFO<0,3 | | | | | | |
| intronic | RARB | 3 | 25206289 | rs80223246 | 4207 | 30.31773 | 5.552231 | 5.460459 | 4.75E-08 | IMPUTED | 0.000477 | 0.000477 | Yes, imputed | 0.177 | No, INFO<0,3 | | | | | | |
