## Supplemental_table4 for "Rare Genetic Variants Correlate with Better Processing Speed"

**Supplement Table4. Summary of the two summary scores for RARB and THRB.**

Chr, Chromosome; Start, start position of SNP on GRCh37 reference panel; End, end position of SNP on GRCh37 reference panel

| Gene | Chr | Start | End | Number of rare variants (MAF < 0.01) | Mean Score ± SD | Median [Min, Max] |
|---|---|---|---|---|---|---|
| RARB | 3 | 24829321 | 25597932 | 3029 | 0.052 ± 0.63 | 0.00 [-3.29, 12.98] |
| THRB | 3 | 24117153 | 24495708 | 1356 | 0.016 ± 0.94 | 0.00 [-3.27, 30.50] |
